# Upregulated-flotillins and sphingosine kinase 2 derail vesicular trafic to stabilize AXL and promote epithelial-mesenchymal transition

**DOI:** 10.1101/2020.05.12.090571

**Authors:** Mallory Genest, Franck Comunale, Damien Planchon, Pauline Govindin, Sophie Vacher, Ivan Bièche, Bruno Robert, Himanshu Malhotra, Andreas Schoenit, Liubov A. Tashireva, Cécile Gauthier-Rouvière, Stéphane Bodin

## Abstract

Altered endocytosis and vesicular trafficking are major players during tumorigenesis. Flotillin overexpression, a feature observed in many invasive tumors, and identified as a marker of poor prognosis, induces a deregulated endocytic and trafficking pathway called Upregulated Flotillin-Induced Trafficking (UFIT). Here, we found that, in non tumoral mammary epithelial cells, induction of the UFIT pathway promotes epithelial-to-mesenchymal transition (EMT) and accelerates the endocytosis of several transmembrane receptors, including AXL, in flotillin-positive late endosomes. AXL overexpression, frequently observed in cancer cells, is linked to EMT and metastasis formation. In flotillin-overexpressing non-tumoral mammary epithelial cells and in invasive breast carcinoma cells, we found that the UFIT-pathway-mediated AXL endocytosis allows its stabilization and depends on sphingosine-kinase 2, a lipid kinase recruited in flotillin-rich plasma membrane-domains and endosomes.

Thus, the deregulation of vesicular trafficking following flotillin upregulation, and through sphingosine kinase 2, emerges as a new mechanism of AXL overexpression and EMT-inducing signaling pathway activation.

## Introduction

Cancer progression is a multistep process involving both genetic and epigenetic changes that independently or in cooperation generate metastatic tumors. One key step in this process is the aberrant reactivation of epithelial-to-mesenchymal (EMT) transition, an essential embryonic process that is also crucial for tumor initiation and cancer cell dissemination during metastatic progression (review in ^1^). EMT induction is orchestrated by a family of EMT-inducing transcription factors (such as ZEB1/2, Snail1, Slug) that are regulated by multiple signaling pathways known to play a role during embryogenesis and also tumor development [e.g. TGFβ receptors, Receptor Tyrosine Kinases (RTKs) and the downstream signaling pathways Ras/MAPK and PI3-K/AKT, WNT, NOTCH] ^2^. These oncogenic signaling pathways are considered cancer drivers, and some of them are used as biomarkers for currently used targeted therapies, while others are potential biomarkers for therapies still under development. Because of EMT pleiotropic roles in the acquisition of invasive properties and of therapeutic resistance, it is important to identify new inducers of this cellular program.

For instance, the RTK AXL, a member of the TAM (Tyro3, AXL, Mer) family of RTKs, is an attractive candidate target for anticancer therapies (review in ^3^). AXL expression is low in normal adult tissues, but is overexpressed in several tumors, and its expression level negatively correlates with overall survival. Aberrant AXL expression is associated with EMT, invasion, metastasis formation, cell proliferation, tumor angiogenesis, reduced anti-cancer immune response, stem cell maintenance, and treatment resistance in many tumor types ^4, 5^. AXL is activated through several mechanisms, including binding to its ligand Gas6, homodimerization in conditions of overexpression, and crosstalk with other RTKs ^4^. However, the rarity of AXL genetic mutations and amplification events, despite its upregulation in many tumor types, suggests a major role for post-transcriptional/translational regulation of AXL expression.

The endosomal system is known to accumulate activated receptors and signaling molecules. It is now clear that endocytosis, which was considered for many years only a mean of limiting signal transduction, is crucial for maintaining and even generating signaling cascades ^6, 7^. Deregulation of vesicular trafficking, and particularly endocytosis, have emerged as major players during tumorigenesis ^8, 9^.

Consistently, upregulation of flotillin1 and/or 2, two proteins involved in the formation of membrane invaginations, is a common feature of many invasive tumours and a marker of poor prognosis, and is also instrumental for cancer cell invasion ^10–14^. When co-upregulated, flotillins promote plasma membrane (PM) invaginations and favor an endocytic pathway called Upregulated Flotillins-Induced trafficking (UFIT) ^15–17^. The UFIT pathway targets protein cargos to non-degradative late endosomes in which flotillins accumulate ^18–21^. Several independent studies performed in cancer cell lines reported that flotillins influence the activation of signaling pathways promoting EMT and cellular invasion and that they are scaffolding factors for several signaling molecules ^22^. Therefore, the UFIT pathway, which is at the crossroad between endocytosis and signaling, has emerged as a crucial mechanism in the regulation of oncogenic pathways.

Here, to study the contribution of flotillin-induced deregulation of the endosomal system to EMT, we reproduced the flotillin 1 and 2 co-upregulation observed in tumors in non-tumoral mammary cell lines (MCF10A and NMuMG) that naturally display low flotillin expression level. Indeed, all previous studies on flotillin role in EMT and cell invasion ^12, 13^ were carried out in cancer cell lines harboring several mutations in signaling and structural genes that promote EMT induction and invasion. This did not allow determining the exact contribution of flotillin upregulation and limited the characterization of the molecular mechanisms that are specifically altered upon flotillin upregulation. In non-tumoral cells, we found that flotillin upregulation is sufficient to promote EMT and to activate key oncogenic signaling pathways. In MCF10A cells, we identified AXL as a major contributor to EMT induced by flotillin upregulation. Using optogenetics to force flotillin oligomerization at the PM in living cells, we demonstrated that AXL is trapped in flotillin microdomains. We then found that the UFIT pathway promotes i) AXL endocytosis towards flotillin-positive late endosomes displaying the characteristics of signaling endosomes that contain activated PM receptors and also K-Ras4B; and ii) AXL stabilization, leading to its overexpression. Finally, in non-tumoral cell lines in which flotillins are upregulated and also in invasive MDA-MB-231 breast cancer cells where flotillins are endogenously overexpressed, AXL stabilization by the UFIT pathway depends on the lipid kinase sphingosine kinase 2 (SphK2) that produces the bioactive lipid sphingosine 1-phosphate (S1P), a key signaling component of the sphingolipid metabolic network during tumorigenesis.

## Results

### Flotillin upregulation in non-tumoral mammary epithelial cells is sufficient to induce EMT

To precisely determine the role of flotillin upregulation in EMT and the molecular mechanisms that are deregulated upon flotillin upregulation, we used non-tumoral epithelial cell lines (human MCF10A and murine NMuMG) in which oncogenic pathways are not activated and with low endogenous flotillin expression levels. From these two cell lines, we generated stable cell lines (MFC10AF1F2 and NMuMGF1F2) that express flotillins at levels similar to those observed in MDA-MB-231 invasive breast cancer cells (fig. S1A-C and ^18^).

First, to identify the gene expression changes and the downstream signaling pathways associated with flotillin upregulation, we compared by RNA-sequencing (RNA-seq) the expression profiles of control MCF10AmCh cells (transfected with the empty mCherry vector, mCh) and MCF10AF1F2 cells (transfected with flotillin 1-HA and flotillin 2-mCherry). Using two differential gene expression analysis pipelines (Tuxedo and DSeq2), we identified 232 upregulated and 570 downregulated genes (802 genes in total, fold change 2.8, *P*<0.05) in MCF10AF1F2 cells compared with control cells (Table S1). Gene ontology (GO) cluster analysis (fig. 1A) revealed that flotillin upregulation was associated with deregulation of genes involved in extracellular matrix, cell adhesion, migration, differentiation, apoptosis and signaling pathways, particularly MAP kinases. These processes are often deregulated during tumor development and particularly during EMT.

**Figure 1:**
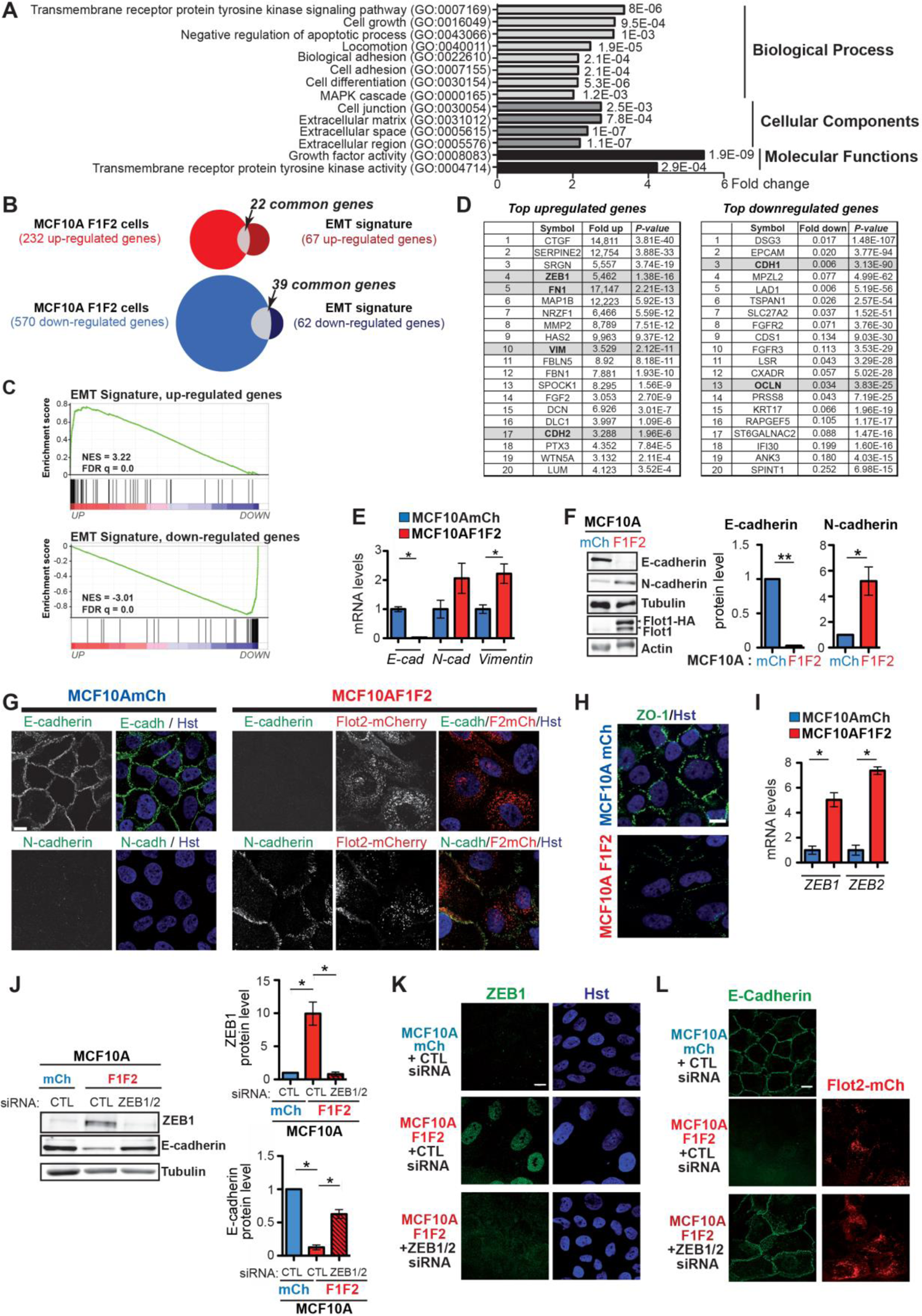
Flotillin upregulation in non-tumoral epithelial mammary MCF10A cells induces EMT. **A)** GO identification of the biological process, cellular component and molecular function terms associated with the 802 genes that were differentially expressed in MCF10AF1F2 cells compared with MCF10AmCh cells. **B)** Venn diagrams comparing the genes that we found to be up- or down-regulated (> or <2.8, p<0.05) by RNA-seq data analysis in MCF10AF1F2 *versus* MCF10AmCh cells with the EMT signature established by ^23^. **C)** Plots from gene set enrichment analysis (GSEA) showing the enriched EMT-signature genes (up-regulated, upper panel; down-regulated, lower panel) in the ranked transcriptome profile. Green lines represent enrichment profiles. NES: normalized enrichment score. FDR: false discovery rate. Each hit from the EMT signature is represented by a vertical black bar, positioned on the ranked transcriptome profile with color-coded fold change values. **D)** List of the genes with the highest fold change among the genes up- and down-regulated and found in MCF10AF1F2 cells and found also in the EMT-transcriptomic signature ^23^ P values were adjusted for multiple testing using the Benjamini-Hochberg procedure. **D)** List of the genes with the highest fold change among the genes up- and down-regulated and found in MCF10AF1F2 cells and found also in the EMT-transcriptomic signature ^23^ P values were adjusted for multiple testing using the Benjamini-Hochberg procedure. **E)** RT-qPCR analysis of the mRNA levels E-cadherin, N-cadherin and vimentin in MCF10AmCh and MCF10AF1F2 cells. Data are the mean ± SEM of four independent experiments. *P<0.05. **F)** Western blot analysis of whole MCF10AmCh and MCF10AF1F2 cell lysates to investigate the expression of E- and N-cadherin, tubulin and flotillin 1. Data are the mean ± SEM of at least four independent experiments. *P<0.05; **P<0.01. **G, H)** Confocal images showing the distribution of E- and N-cadherin, flotillin 2-mCherry and ZO-1 in MCF10AmCh and MCF10AF1F2 cells and of vimentin in NMuMGmCh and NMuMGF1F2 cells, analyzed by immunocytochemistry. Nuclei were stained with Hoechst (Hst; blue). In G, for vimentin staining, the z projection corresponds to the stacked signal of 30 plans performed every 0.3 µm. **I)** RT-qPCR analysis of the mRNA levels of ZEB1 and ZEB2 in MCF10AmCh and MCF10AF1F2 cells. Histograms show the mean ± SEM of 4 independent experiments. *P<0.05. **J)** Western blotting to assess ZEB1, E-cadherin and α-tubulin expression in cell lysates from MCF10AmCh cells and MCF10AF1F2 cells transfected with siRNAs against ZEB1/2 or control siRNA against luciferase (CTL). Histograms (mean ± SEM of 4 independent experiments) show the quantification of the amount of each protein in the indicated conditions. **K, L)** Distribution of ZEB1 (K) and E-cadherin (L) in MCF10AmCh cells and in ZEB1/2 siRNA- or CTL siRNA-transfected MCF10AF1F2 cells analyzed by immunofluorescence using specific antibodies. In K, nuclei were stained with Hoechst (Hst). Images in G,H, K and L are representative of 3 independent experiments. Scale bar: 10 µm.

**Figure S1:**
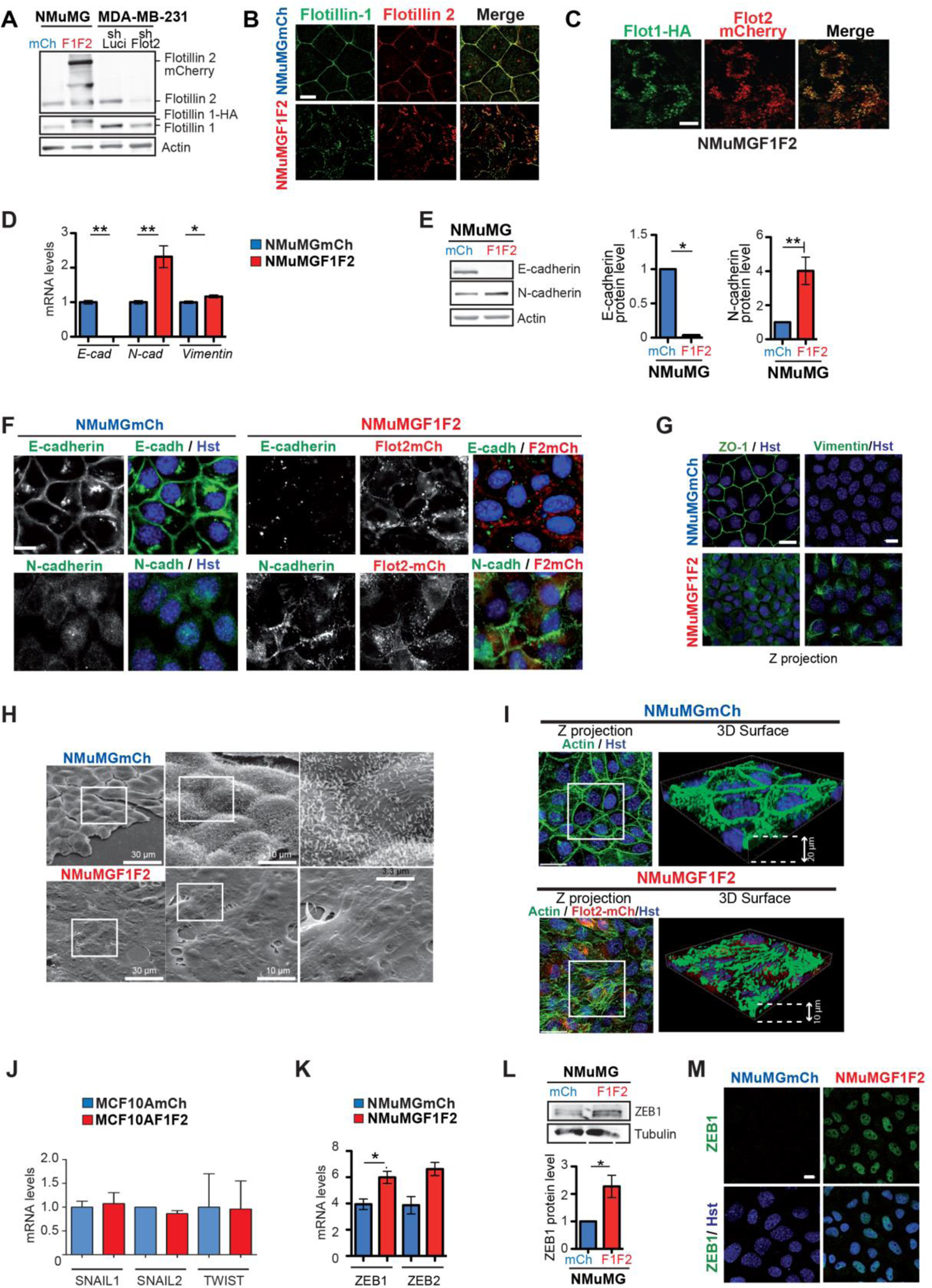
Flotillin upregulation induces EMT in the NMuMG non-tumoral murine epithelial mammary cell line. **A)** Lysates of NMuMG cells that stably express mCherry (NMuMGmCh) or both flotillin 1-HA and flotillin 2-mCherry (NMuMGF1F2) were analyzed by western blotting using antibodies against flotillin 1 and flotillin 2. In NMuMGF1F2 cells, flotillin levels were comparable to those detected in the invasive mammary tumor MDA-MB-231 cell line. Lysates from MDA-MB-231shFlot2 cells (generated using a shRNA against *FLOT2* to knockdown both flotillins ^18^) also were analyzed. Actin was used as loading control. **B)** Confocal images showing that endogenous flotillin 1 and 2 are localized at the plasma membrane in NMuMGmCh cells. In NMuMGF1F2 cells, they translocate to vesicular compartments, where they co-localize, as previously shown in MCF10AF1F2 and in MDA-MB-231 cells ^18^. **C)** Confocal images showing the co-localization of flotillin 1-HA and flotillin 2-mCherry in intracellular vesicles in NMUMGF1F2 cells. **D, K)** RT-qPCR analysis of the mRNA levels of E-cadherin, N-cadherin and vimentin (D) and of ZEB1 and ZEB2 (K) in NMuMGmCh and NMuMGF1F2 cells. Histograms show the mean ± SEM of 4 independent experiments. *P<0.05; **P<0.01. **E, L)** Western blotting performed using cell lysates from NMuMGmCh and NMuMGF1F2 cells to compare the amount of E- and N-cadherin (E), and ZEB1 (L). Histograms show the mean ± SEM of at least 4 independent experiments. *P<0.05; **P<0.01. **F, G)** Comparison of the distribution of endogenous E-, N-cadherin (F) and ZO-1 and vimentin (G) in NMuMGmCh and NMuMGF1F2 cells analyzed by immunofluorescence. Nuclei are stained with Hoechst (Hst, blue). **H)** Scanning electron microscopy images of the monolayer formed by NMuMGmCh and NMuMGF1F2 cells. **I)** Z-projection and 3D surface reconstruction of NMuMGmCh and NMuMGF1F2 cells after staining for F-actin, flotillin 2-mCherry and Hoechst (nuclei), performed with 35 plans and 22 plans every 0.3.µm for NMuMGmCh and NMuMGF1F2 respectively. **J, K)** *Characterization of the transcriptional repressors of E-cadherin expression*. RT-qPCR analysis of the mRNA levels of Snail1, Snail2, Twist in MCF10AmCh and MCF10AF1F2 cells (J) and ZEB1 and 2 in NMuMGmCh and NMuMGF1F2 cells (K). Histograms show the mean ± SEM of 3 independent experiments. **M)** Confocal images of ZEB1 localization and nuclei stained with Hoechst (Hst, blue) in NMuMGmCh and NMuMGF1F2 cells. Images inB, C, F, G, I and M are representative of cells observed in at least 3 independent experiments Scale bar: 10 µm.

To confirm that flotillin upregulation was associated with differential expression of EMT-specific genes, we compared the list of genes differentially expressed between MCF10AmCh and MCF10AF1F2 cells with the EMT transcriptional signature previously established by a meta-analysis that combined 18 independent gene expression studies on EMT induced by different treatments in different cell types ^23^. One third (22/67) of the upregulated genes and more than half (39/62) of the downregulated genes in this EMT signature overlapped with the genes deregulated upon flotillin overexpression in MCF10A cells (fig. 1B). A gene set enrichment analysis (GSEA) was performed and confirmed the enrichment of the EMT signature in our ranked transcriptome profile (fig. 1C). The analysis of the top 20 up- and down-regulated genes between MCF10AmCh and MCF10AF1F2 cells highlighted the decreased expression of E-cadherin and occludin, concomitantly with the increased expression of ZEB1, fibronectin, vimentin and N-cadherin (figure 1C).

We validated these results in MCF10AF1F2 and NMuMGF1F2 cells by RT-qPCR (fig. 1E and S1D), western blotting and immunocytochemistry analysis (fig. 1F-H and S1E-G). We confirmed the downregulation (mRNA and protein) of the hallmark epithelial marker E-cadherin (fig. 1E-G and S1D-F), of the polarity marker ZO-1 (fig. 1H and S1G), and the upregulation of the mesenchymal markers N-cadherin and vimentin (fig. 1E-G and S1D-G). Morphological analysis using scanning electron microscopy and F-actin cytoskeleton staining showed that cell polarity was lost in NMuMGF1F2 cells (flotillin upregulation) compared with NMuMGmCh cells (empty mCherry vector) (fig. S1H-I).

Several transcription factors, such as Snail1, Snail2, Twist, ZEB1 and ZEB2, can inhibit E-cadherin expression ^24^. Flotillin upregulation did not affect Snail 1/2 and Twist expression, as shown by the transcriptomic analysis and by quantification of their mRNA levels (fig. S1J). Conversely, it induced ZEB 1 and 2 mRNA expression (fig. 1I and S1K) and ZEB1 protein expression (fig. 1J and S1L)). ZEB1 localized in the nucleus (fig. 1J and S1M). Moreover, ZEB1/2 knockdown in MCF10AF1F2 cells restored E-cadherin expression (fig. 1J) and its accumulation at cell-cell contacts (fig. 1L). This demonstrated the key role of ZEB1/2 in E-cadherin downregulation upon flotillin upregulation.

In conclusion, flotillin upregulation in non-tumoral mammary epithelial cells is sufficient to induce EMT.

### Flotillin upregulation in non-tumoral mammary epithelial cells is sufficient to activate several oncogenic signaling pathways

The GO analysis of the genes that were differentially expressed between control and flotillin-upregulated MCF10A cells revealed a strong association with signal transduction pathways, particularly those involving transmembrane RTKs and the MAPK pathway (fig. 1A). Different bioinformatics analysis [DAVID, Enrich R analysis of the Kyoto Encyclopedia of Genes and Genomes (KEGG) pathways, and Genomatix] identified the PI3-K/AKT and Ras pathways, two major oncogenic signaling cascades (Table S2). In agreement, analysis of the phosphorylation profile of some kinases and of their protein substrates using the human Phospho-kinase Array revealed the increased phosphorylation of at least four main tyrosine (Y) and serine/threonine (S/T) kinases involved in oncogenic signaling pathways (ERK1/2, FAK, AKT, Fyn) and of their substrates (Hsp27, STAT) (fig. 2A). We demonstrated the activation of Ras (fig. 2B) in MCF10AF1F2 cells compared with MCF10AmCh cells, as well as ERK and PI3-K/AKT activation, as indicated by the increased ERK1/2 phosphorylation at T202/Y204 (fig. 2C, fig. S2A for NMuMG cells), the increased nuclear localization of ERK2-GFP (fig. 2D), and the increased AKT phosphorylation at S473 (fig. 2E). STAT3 phosphorylation at Y705 (fig. 2F) was also increased in MCF10AF1F2 cells compared to MCF10AmCh cells, indicating its activation upon flotillin overexpression.

**Figure 2:**
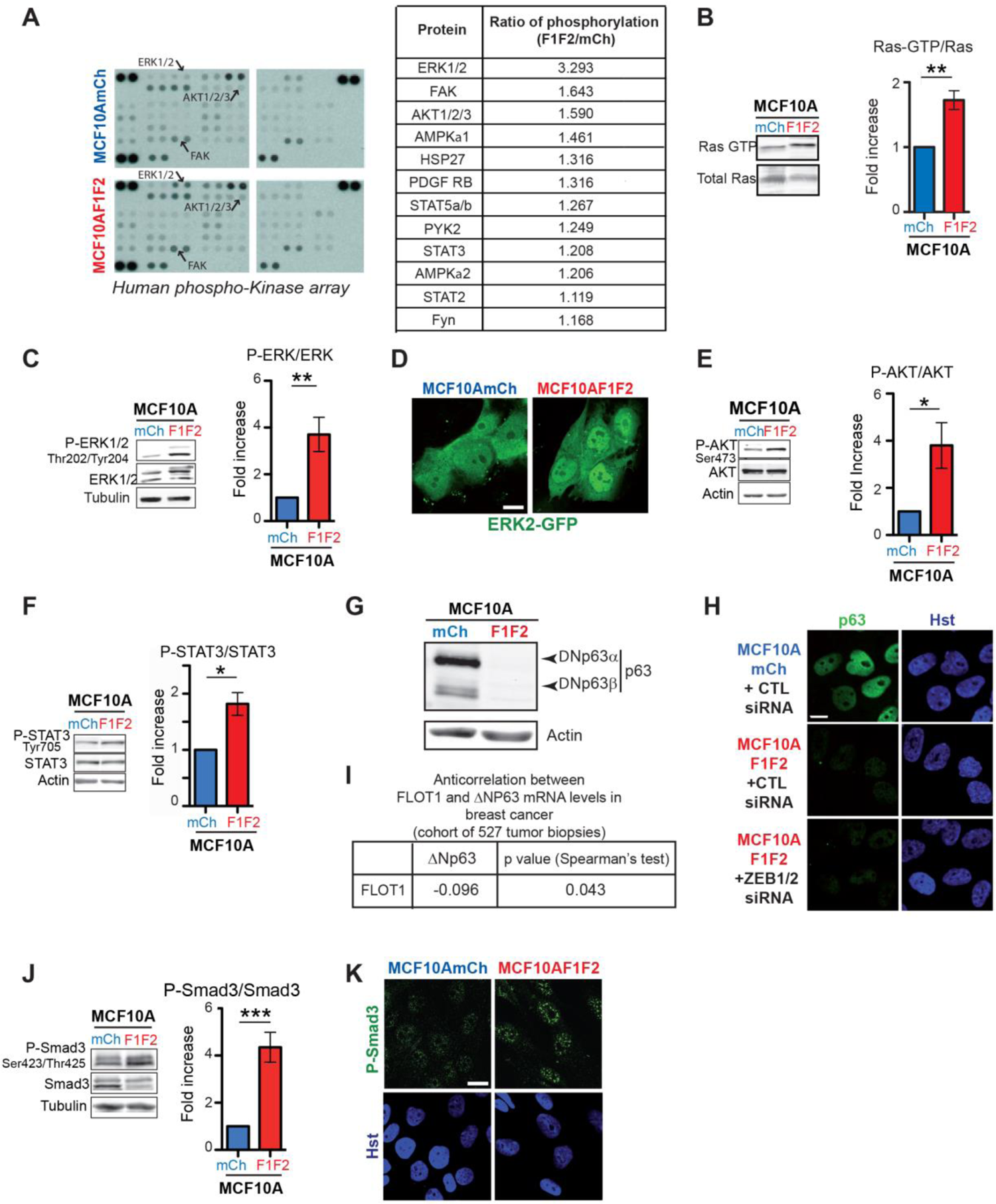
Flotillin upregulation promotes the activation of oncogenic signaling pathways. **A)** The level of active GTP-bound Ras was measured using GST-Raf-RBD in MCF10AmCh and MCF10AF1F2 cell lystaes. The histogram shows the mean value (± SEM) of Ras activity normalized to the amount of total protein calculated from 5 independent experiments. **B)** Representative example of human phospho-kinase arrays incubated with MCF10AmCh and MCF10AF1F2 cell lysates, as described by the manufacturer. The table lists the twelve kinases and kinase substrates with the highest ratio of phosphorylation change between MCF10AF1F2 and MCF10AmCh cells. Values are the mean of 2 independent experiments. **C, E, F, J)** MCF10AmCh and MCF10AF1F2 cell lysates were analyzed by western blotting to assess the phosphorylation status of the signaling proteins ERK1/2 (C), AKT (E), STAT3 (F), and Smad3 (G), with the indicated antibodies. The histograms show the mean value (± SEM) of protein phosphorylation normalized to the total amount of the considered protein in MCF10AF1F2 cells compared with control MCF10AmCh cells calculated from at least 4 independent experiments. **D)** MCF10AmCh and MCF10AF1F2 cells were transfected with an ERK2-GFP plasmid and ERK2-GFP distribution was analyzed by confocal microscopy at 36 hours post-transfection. **G) Δ**Np63α and β expression in MCF10AmCh and MCF10AF1F2 cells analyzed by western blotting using an antibody against all p63 isoforms. Characterization of the p63 isoforms allowed identifying ΔNp63α and β as the main isoforms expressed in MCF10AmCh cells (fig. S2A). **H)** Confocal microscopy images of MCF10AmCh and in ZEB1/2 siRNA- or CTL (against luciferase) siRNA-transfected MCF10AF1F2 stained by immunofluorescence using an antibody against all p63 isoforms and by Hoechst (Hst) for nuclei visualization. **I)** Analysis of the correlation between the mRNA levels of *FLOT1* and *p63* in breast cancer biopsies from 527 patients^18^. **K)** Confocal microscopy images of MCF10AmCh and MCF10AF1F2 cells stained by immunofluorescence with an antibody against Smad3 phosphorylated on S423/425 and Hoechst (Hst) staining for nuclei. In D, H and K, image are representative of 3 independent experiments. Scale bars: 10µm. *P<0.05; **P<0.01; ***P<0.001

**Figure S2:**
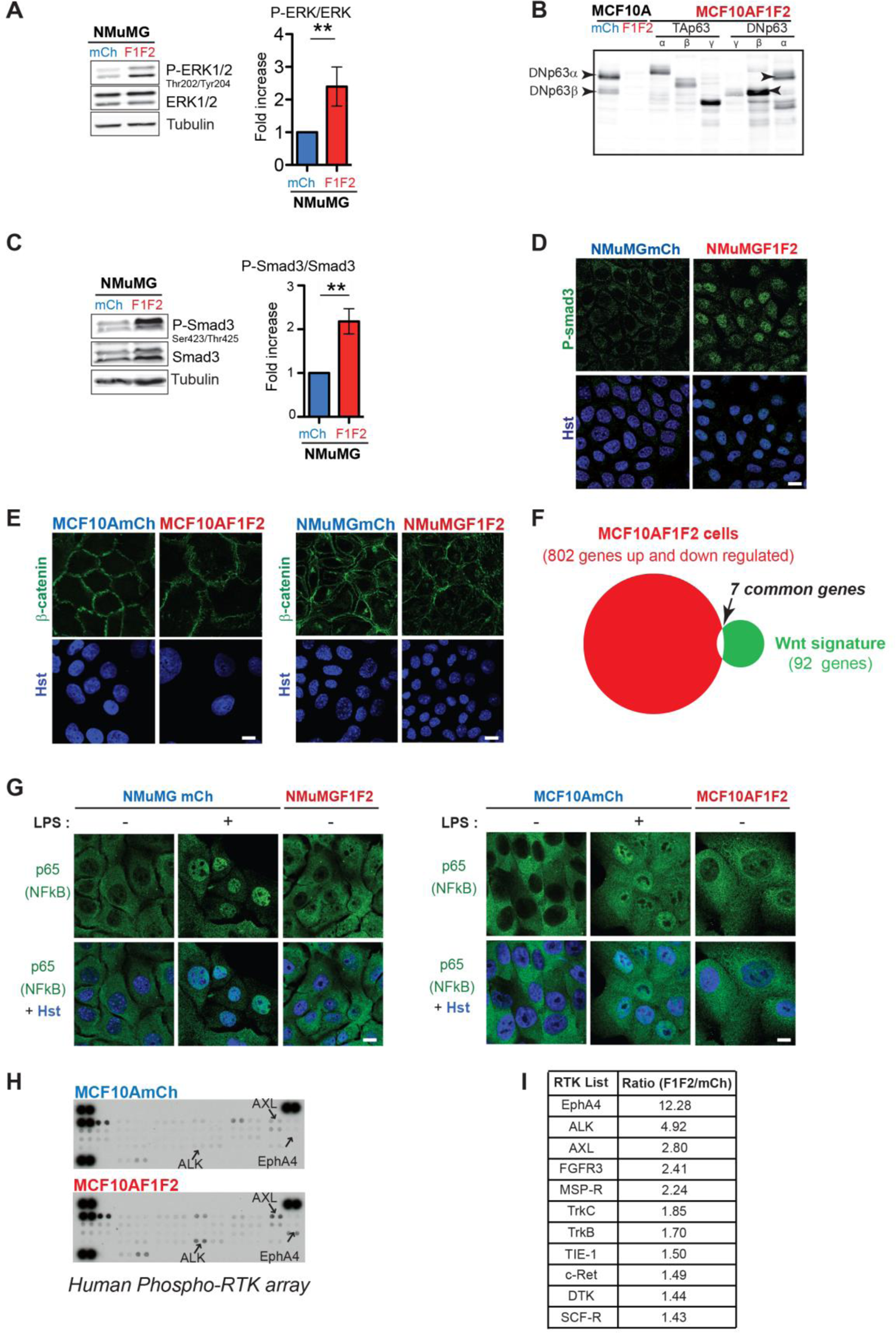
Determination of the p63 isoforms expressed in MCF10AmCh cells and downregulated in MCF10AF1F2 cells. Identification of non-activated signaling pathways downstream of flotillin upregulation in mammary epithelial cells. Screening for activated RTKs upon flotillin upregulation. **A)** *Characterization of the p63 isoforms expressed in MCF10AmCh and MCF10AF1F2 cells.* Western blot analysis of the p63 isoforms in cell lysates from MCF10AmCh, MCF10AF1F2 and MCF10AF1F2 cells transfected with different non-tagged p63 isoforms (TAp63α, β, γ or ΔNp63α, β, γ) using an antibody against all p63 isoforms. ΔNp63α and to a lesser extent ΔNp63β are the two mains isoforms expressed in MCF10A cells. The expression of ΔNp63α and of ΔNp63β was abolished in MCF10AF1F2 cells (as shown in fig. 2G). **B, C)** Cell lysates from NMuMGmCh and NMuMGF1F2 cells were analyzed by western blotting to evaluate the phosphorylation status of ERK1/2 (B) and Smad3 (C). Histograms (mean ± SEM of 4 independent experiments) show the quantification of the phosphorylation-signal normalized to the total amount of each protein. **P<0.01. **I) D)** NMuMGmCh and NMuMGF1F2 cells were labeled with a antibody against Smad3 phosphorylated on S423/425 and with Hoechst (Hst). **E, F, G)** The oncogenic NF-kB and WNT canonical signaling pathways are not activated following flotillin upregulation in MCF10A and NMuMG cells. **E)** β-catenin distribution, analyzed by immunofluorescence is similar in MCF10AmCh and MCF10AF1F2 cells and NMuMGmCh and NMuMGF1F2 cells, and no nuclear staining was detected, indicating that the WNT canonical pathway is not activated. Nuclei are stained with Hoechst (Hst). **F)** Venn diagrams illustrating the small overlap between the 802 differentially expressed genes identified by RNA-seq in MCF10AF1F2 cells *versus* MCF10AmCh cells and the 92 differentially expressed genes identified in an established WNT signature. Only 7/92 genes overlapped. **G)** Cellular distribution of the NF-kB p65 subunit, analyzed by immunofluorescence, in MCF10AmCh and MCF10AF1F2 cells and in NMuMGmCh and NMuMGF1F2 cells. In all cases, NF-kB p65 was excluded from the nucleus, indicating that the NF-kB pathway is not activated upon flotillin upregulation. As positive control, MCF10AmCh cells and NMuMGmCh cells were stimulated with lipopolysaccharide LPS (1µg/ml, 1h) to induce NF-kB p65 translocation to the nucleus. **H)** *Flotillin upregulation increases the phosphorylation level of several RTKs.* Human phospho-Receptor Tyrosine kinase arrays were incubated with 200 µg of MCF10AmCh or MCF10AF1F2 cell lysates (cultured in the presence of serum) and processed as described by the manufacturer. One array example is shown. **I)** List of the 11 RTKs with the highest increase in phosphorylation in MCF10AF1F2 compared with MCF10AmCh cells (data from 2 independent experiments). Images in D, E, G are representative of at least 3 independent experiments. Scale bar: 10 µm.

In MCF10A cells, expression of a hyperactive H-Ras mutant or its downstream effector PI3-K leads to ZEB1/2-dependent E-cadherin downregulation and EMT *via* the transcriptional repression of ΔNp63α, the predominant p63 isoform in MCF10A cells ^25^. In these cells, ΔNp63α downregulation is an EMT marker ^25, 26^. Flotillin upregulation in MCF10A cells led to the reduction of ΔNp63α and ΔNp63β expression (western blotting and immunocytochemistry, fig. 2G, H and S2B; Table S1). We excluded ZEB 1 and 2 implication in ΔNp63α downregulation through a negative feed-back loop because ZEB1 and 2 knock-down did not rescue ΔNp63α expression in MCF10AF1F2 cells (fig. 2H). Moreover, data obtained from a cohort of 527 patients with breast cancer indicated that flotillin 1 and ΔNp63 mRNA expression levels were negatively correlated (fig. 2I).

High TGF-β signaling pathway activity contributes to tumor development, notably through its ability to induce EMT ^1^, and flotillins participate in the activation of this pathway in nasopharyngeal carcinoma ^12, 13^. Here, we found that flotillin upregulation in non-tumoral cells also activated the TGF-β signaling pathway, as indicated by the significantly higher Smad3 phosphorylation at S423/T425 (fig. 2J and S2C) and the nuclear localization of phosphorylated Smad3 in MCF10AF1F2 (fig. 2K) and NMuMGF1F2 (fig. S2D) cells compared with control cells (mCh).

We never detected activation of the NF-kB and WNT canonical pathways (fig. S2E-G) although several oncogenic pathways were activated upon flotillin upregulation Altogether, these data demonstrated that flotillin upregulation in cells without oncogenic mutations (MCF10A and NMuMG cells) is sufficient to activate major oncogenic pathways, such as the Ras/ERK MAP kinase, AKT and TGF-β signaling cascades.

### Flotillin upregulation in non-tumoral mammary epithelial cells increases the expression of AXL, a key player in flotillin-induced EMT

To identify how flotillin upregulation in non-tumoral mammary cells activates these oncogenic signaling pathways, we used a Phospho-RTK array that allows the simultaneous screening of 49 tyrosine-phosphorylated RTKs. The tyrosine phosphorylation of 11 RTKs was increased by more than 1.4-fold in MCF10AF1F2 cells compared with MCF10AmCh cells grown in the presence of serum for 48 hours (fig. S2H, I). To validate these results, for the three top receptors, we compared their Y-phosphorylation status in the two cell lines by western blotting using antibodies against phosphorylation at specific Y residues.Unfortunately the available tools allowed us to validate this result only for AXL, but not for EphA4 and ALK. Specifically, the phosphorylation of AXL, the deregulation of which were reported in many invasive cancer cells has been linked to EMT ^4, 5, 27, 28^ was increased (2.53 ± 0.71-fold) in MCF10AF1F2 cells compared to control MCF10AmCh cells. Moreover, this change was of the same order of magnitude as the increase (2.57 ± 0.61-fold) in AXL protein level between these two cell lines (fig. 3A). This indicated that flotillin upregulation, increased AXL protein level but did not stimulate AXL phosphorylation *per se*, Consequently, the ratio phospho-Y702 AXL/AXL was identical between MCF10AmCh and MCF10AF1F2 cells (fig. 3A). As a control, the expression level of the transferrin receptor CD71 was comparable in both cell lines (fig. S3A). The increase in AXL level was observed in presence of serum (fig. S3B), which might contain its ligand Gas6, but also in starved cells, indicating that flotillin upregulation promotes AXL increase in MCF10A cells independently of Gas6. As aberrant AXL expression has been reported in many invasive cancer types and is linked to EMT, metastasis formation, and poor prognosis ^4, 5, 27, 28^, AXL increase in MCF10AF1F2 cells suggests that flotillin upregulation may participate in AXL overexpression in cancer. Interestingly, we found that AXL expression, which is barely detectable in normal breast tissues, was increased in breast tumor samples and was correlated with that of flotillin 1, particularly in the luminal subtype (fig. S4 A-B).

**Figure 3:**
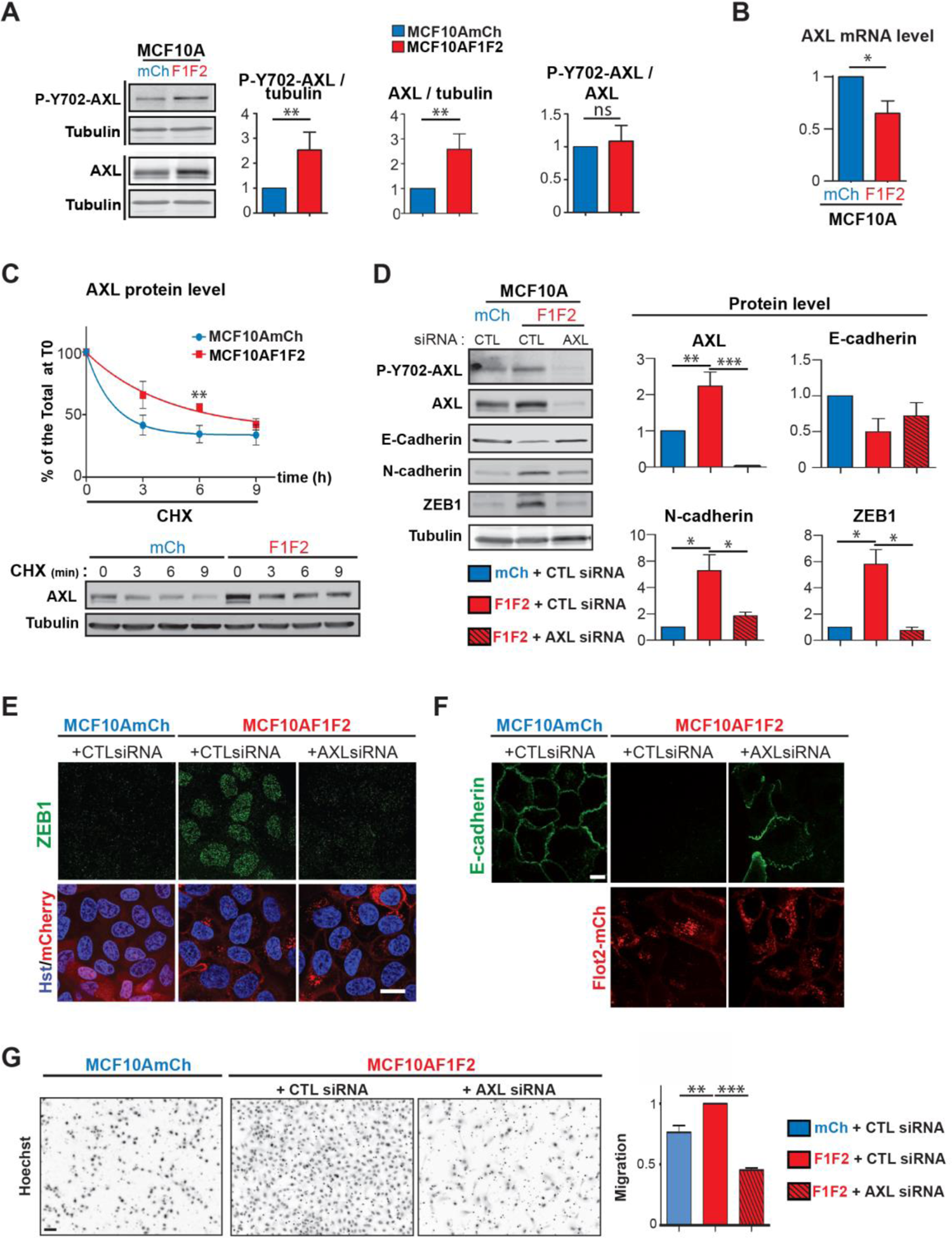
AXL is stabilized upon flotillin-upregulation and is involved in upregulated flotillins-induced EMT and cell migration. **A)** *Flotillin upregulation promotes AXL protein level increase*. Cell lysates from MCF10AmCh and MCF10AF1F2 cells were probed by western blotting with antibodies against AXL phosphorylated on Y702, total AXL and tubulin. Results are expressed as fold-increase compared with MCF10AmCh cells and are the mean ± SEM of 6 independent experiments. **B)** *Flotillin upregulation does not increase AXL mRNA level.* RT-qPCR analysis of AXL expression in MCF10AmCh and MCF10AF1F2 cells. Results are expressed relative to the level in MCF10AmCh cells and are the mean ± SEM of 4 independent experiments. **C)** *Flotillin upregulation stabilizes AXL.* MCF10A and MCF10AF1F2 cells were incubated with cycloheximide (CHX, 100µg/ml) and cell lysates collected at the indicated time points. AXL and tubulin levels were assessed by western blotting (representative western blots are shown). The graph shows the normalized AXL levels in each cell line during CHX incubation. The results are the mean ± SEM of 6 to 8 independent experiments depending on the time point, and are expressed as the percentage of AXL level at T0. CHX treatment was stopped after 9 hours because MCF10Amch cells did not survive longer treatment. The difference is significant at time 6 hours ** (*p*<0.01). **D)** *AXL knock-down in MCF10AF1F2 cells decreases ZEB1 and N-cadherin expression*. MCF10AmCh and MCF10AF1F2 cells were transfected with anti-luciferase (CTL) or AXL siRNAs and probed by western blotting with antibodies against the indicated proteins. The quantification results are the mean ± SEM of 4 independent experiments, and are expressed as fold-increase compared with control (MCF10AmCh). **E)** ZEB1 nuclear localization analyzed by immunocytochemistry and co-staining with Hoechst (Hst) and **F)** E-cadherin distribution analyzed by immunocytochemistry in MCF10AmCh and MCF10AF1F2 cells transfected with siRNA against luciferase (CTL) or AXL. Scale bar: 10 µm. **F)** *Boyden chamber migration assay.* MCF10AmCh and MCF10AF1F2 cells transfected with siRNA against luciferase (CTL) or AXL were seeded in the top of Boyden chamber inserts in serum-free medium, and serum-containing medium, acting as chemoattractant, was placed in the bottom well. Representative inverted-microscopy images of Hoechst-stained nuclei of cells that migrated through the pores. The histogram (mean ± SEM of at least four independent experiments) show the quantification of cell migration of the indicated cell lines. Scale bar: 50µm. Images in E, F, G are representative of at four independent experiments. \**p*<0.05, \*\**p*<0.01,\*\*\**p*<0.001

**Figure S3:**
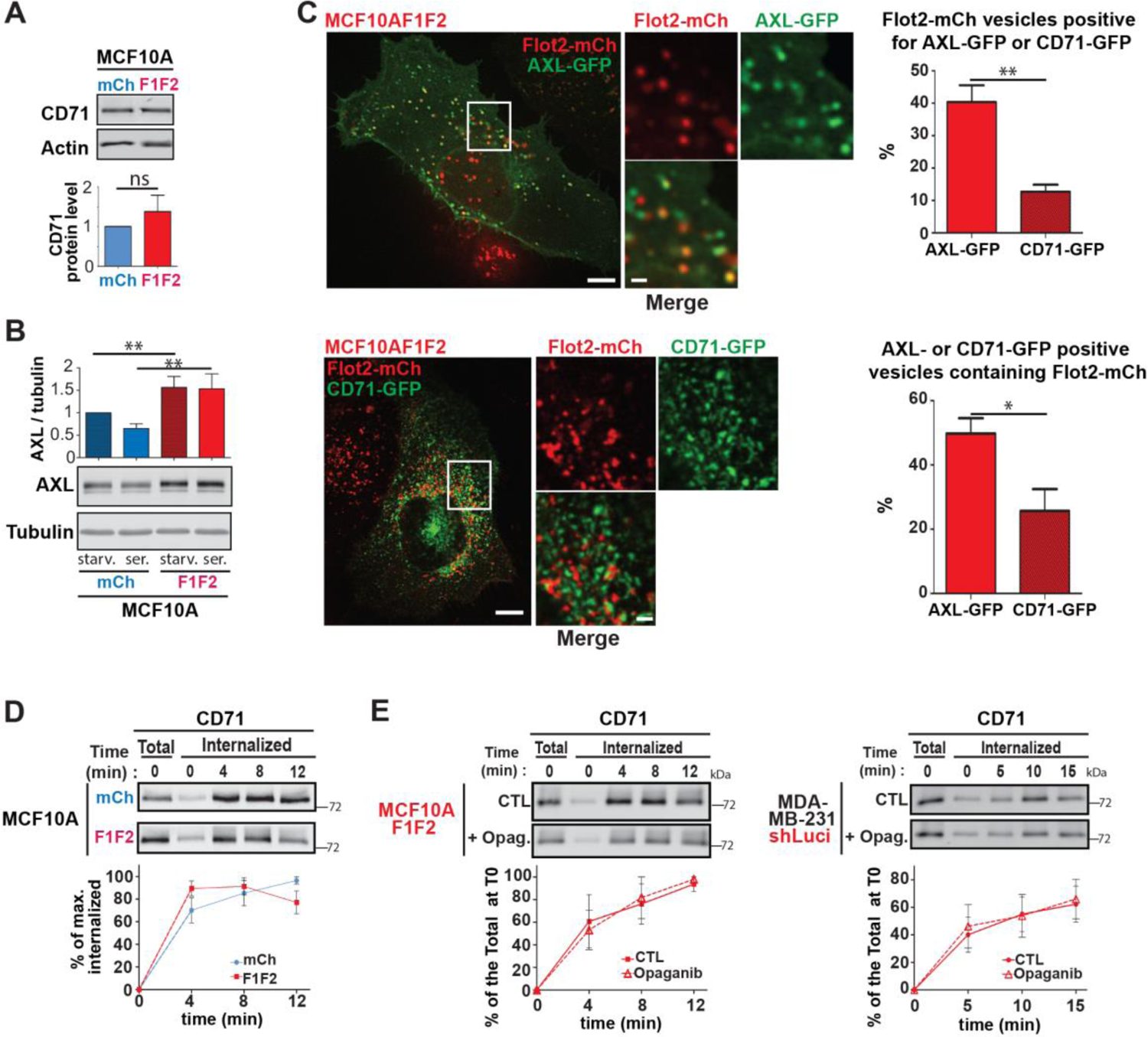
The transferrin receptor CD71 is not a cargo of the UFIT pathway. **A)** *Flotillin upregulation does not affect CD71 level.* Cell lysates of MCF10AmCh and MCF10AF1F2 cells were probed by western blotting with antibodies against CD71 and actin. Results are expressed as fold-increase compared with MCF10AmCh cells and are the mean ± SEM of 4 independent experiments. **B)** *Flotillin upregulation promotes AXL protein level increase independently from serum stimulation.* Lysates from MCF10AmCh and MCF10AF1F2 cells serum-starved for 16h (starv.) our maintained in the presence of serum (ser.) were probed by western blotting with antibodies against AXL and tubulin. Results are expressed as fold-increase compared with MCF10AmCh cells in serum-starved conditions and are the mean ± SEM of 5 independent experiments. \*\**p*<0.01 **C)** *Transferrin receptor poorly localized in in flotillin-positive late endosomes compared with AXL.* Live MCF10AF1F2 cells that express CD71-GFP or AXL GFP were imaged by spinning disk confocal microscopy. Images shown are representative of several cells in 3 independent experiments. Scale bars: 10 µm in the main image and 2 µm in the magnified images from the boxed area. The histograms show the percentage of Flotillin2-mCherry (Flot2-mCh)-positive vesicles containing AXL-GFP or CD71-GFP (upper histogram) and the percentage of AXL-GFP or CD71-GFP positive vesicles containing Flot2-mCh (lower histogram). Results are the mean ± SEM (n=690 Flot2-mCh-positive vesicles and n=746 AXL-GFP labeled vesicles in 21 AXL-GFP-expressing MCF10AF1F2 cells; and n=1511 Flot2-mCh-positive vesicles and n=1207 AXL-GFP labeled vesicles in 20 CD71-GFP-expressing MCF10AF1F2 cells). \**p*<0.05, \*\**p*<0.01 **D)** *Flotillin upregulation does not affect transferrin receptor endocytosis.* Kinetics of CD71 (transferrin receptor) internalization in MCF10AmCh and MCF10AF1F2 cells analyzed in the same experiments presented in figure 6 G. Surface proteins were labeled with biotin at 4°C and cells were incubated at 37°C for the indicated times to allow endocytosis. CD71 presence in the internalized biotinylated fraction was probed by western blotting using relevant antibodies, and quantified as the percentage of the maximum level of internalized protein. Results are expressed as the mean ± SEM of 4 independent experiments. The CD71 internalization rates were not significantly different between cell lines at 4, 8 and 12 min, p>0.1. **E)** *SphK2 inhibition has no impact on CD71 internalization in flotillin-upregulated cells.* Kinetics of CD71 internalization in MCF10AF1F2 and in MDA-MB-231shLuci cells untreated (CTL) or pre-treated with opaganib (50µM, 4h). Surface proteins were labeled with biotin at 4°C, and cells were incubated at 37°C for the indicated times to allow endocytosis. Opaganib was maintained thoughout the experiments. CD71 presence in the internalized biotinylated fraction was probed by western blotting using relevant antibodies. Results are expressed as the percentage of the maximum level of CD71 surface level at T0 and are the mean ± SEM of 4 independent experiments. In contrast to AXL (fig. 6E, F), CD71 internalization rates were not different between cell lines or between the different conditions.

**Figure S4:**
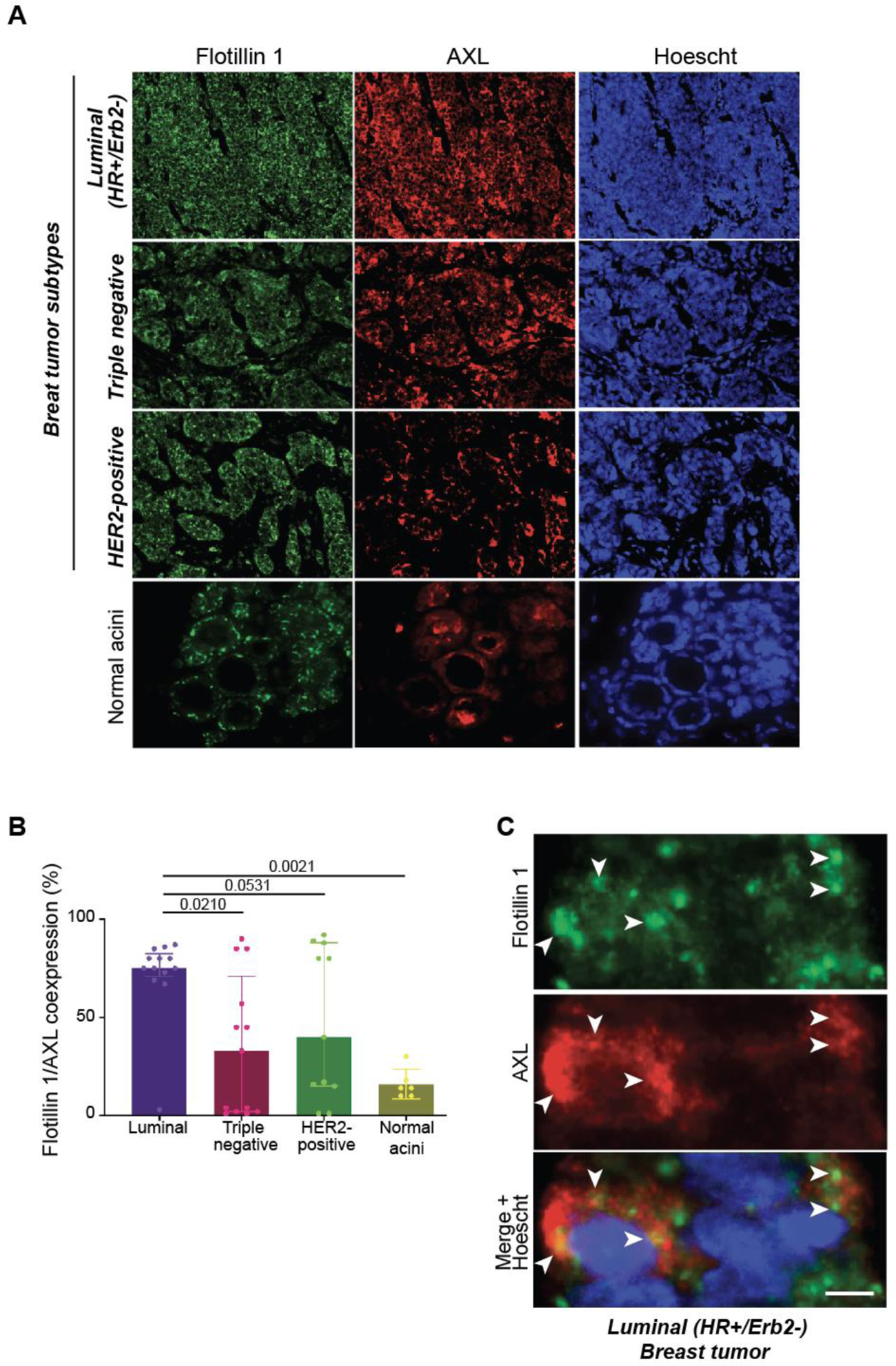
Flotillin 1 and AXL are coexpressed in invasive breast tumor cells. **A)** Immunofluorescence analysis of flotillin 1 and AXL in invasive breast tumor tissue sections in one patient sample for each breast tumor subtype and in non-tumoral breast tissue. Nuclei were stained with DAPI. Scale bar: 10µm. **B)** Immunofluorescence analysis of flotillin 1 and AXL expression in 37 breast tumor samples and 6 non-tumoral breast tissue samples. Co-expression was calculated in the different subtypes with the Vectra 3.0 Automated Quantitative Pathology Imaging System. **C)** High magnification immunofluorescence images of flotillin 1 and AXL expression in a luminal B invasive breast tumor section. Nuclei were stained with Hoechst. Arrows indicate flotillin 1 and AXL colocalization in intracellular vesicles. Scale bar: 10µm.

We then further investigated how fotillin upregulation promotes AXL level increase. This did not occur at the transcriptional level because *AXL* mRNA level was reduced in MCF10AF1F2 cells compared with MCF10AmCh cells (fig. 3B). Next, we assessed whether flotillin upregulation affected AXL stability by comparing AXL expression in MCF10AmCh and MCF10AF1F2 cells incubated with cycloheximide (CHX) to block protein synthesis. AXL expression was decreased by 50% after 1.9h and 6.8h of incubation with CHX in MCF10AmCh and in MCF10AF1F2 cells, respectively (fig. 3C). These data indicate that flotillin upregulation promotes AXL stabilization. We analyzed AXL contribution to EMT and cell migration induced by flotillin upregulation in MCF10A cells. AXL knockdown by siRNas in MCF10AF1F2 cells decreased ZEB1 expression (fig. 3D and E), reversed the E- to N-cadherin switch (fig. 3D and F) and reduced their migration (fig. 3G).

Altogether, these results showed that flotillin upregulation in non-tumoral mammary epithelial cells does not stimulate AXL activation *per se*, but promotes AXL stabilization leading to the increase of its phosphorylated form, as observed in many invasive cancer types. These results also revealed that AXL plays a central role in flotillin upregulation-induced EMT and cell migration in mammary epithelial cells.

### AXL is a cargo of the UFIT pathway that is targeted to flotillin-positive late endosomes

Deregulation of endocytosis and vesicular trafficking were shown to alter the fate of signaling receptors, affecting their expression level and downstream signaling ^7, 8, 29^. As flotillin upregulation promotes the UFIT pathway, an endocytic pathway that allows targeting cargo proteins to late endosomes, thus modifying their fate ^15, 18, 30, 31^, we asked whether AXL is a cargo of the UFIT pathway. First, to determine whether AXL was recruited to flotillin-positive microdomains at the PM, we developed an optogenetic approach based on the CRY2-CIBN system ^32^ to force flotillin oligomerization by light illumination combined with total internal reflection fluorescence (TIRF) video-microscopy (see fig. 4A for the description of the experimental set-up and fig. S4A for the validation of the flotillin 2-CIBN-mCherry association with endogenous flotillins 1 and 2). Using MCF10A cells that express flotillin 2-CIBN-mCherry, CRY2-mCitrin and AXL-Halo, upon light illumination to induce CRY2 oligomerization and binding to CIBN, we demonstrated the formation at the PM of domains that contained both flotillin 1 and 2 (fig. S5B) and AXL-Halo (fig. 4B and S5B and D, video 1). Moreover, using time-lapse imaging we could detect the concomitant endocytosis of flotillin 2-mCherry and AXL-GFP from PM sites of MCF10AF1F2 cells where both proteins initially co-accumulated (fig. 4C, video 2; fig. S5C for its characterization). In agreement, we showed that in MCF10AmCh cells, AXL was mainly at the PM. Conversely, in MCF10AF1F2 cells, AXL was still detectable at the PM in flotillin-rich regions, but was enriched in flotillin-positive vesicles (fig. 4D), that were positive for the late endosomal marker CD63 (fig. 4J). Similarly, in MDA-MB-231 mammary tumor cells, in which endogenous flotillins are overexpressed ^18^, AXL and flotillin 2 colocalized at the PM and in intracellular vesicles (fig. S5E) that were CD63-positive (fig. 4K). AXL-GFP co-trafficked with flotillin 2-mCherry in living MCF10AF1F2 cells (fig. S5F, video 3) and in living MDA-MB-231 cells (fig. S5G, video 4).

**Figure 4:**
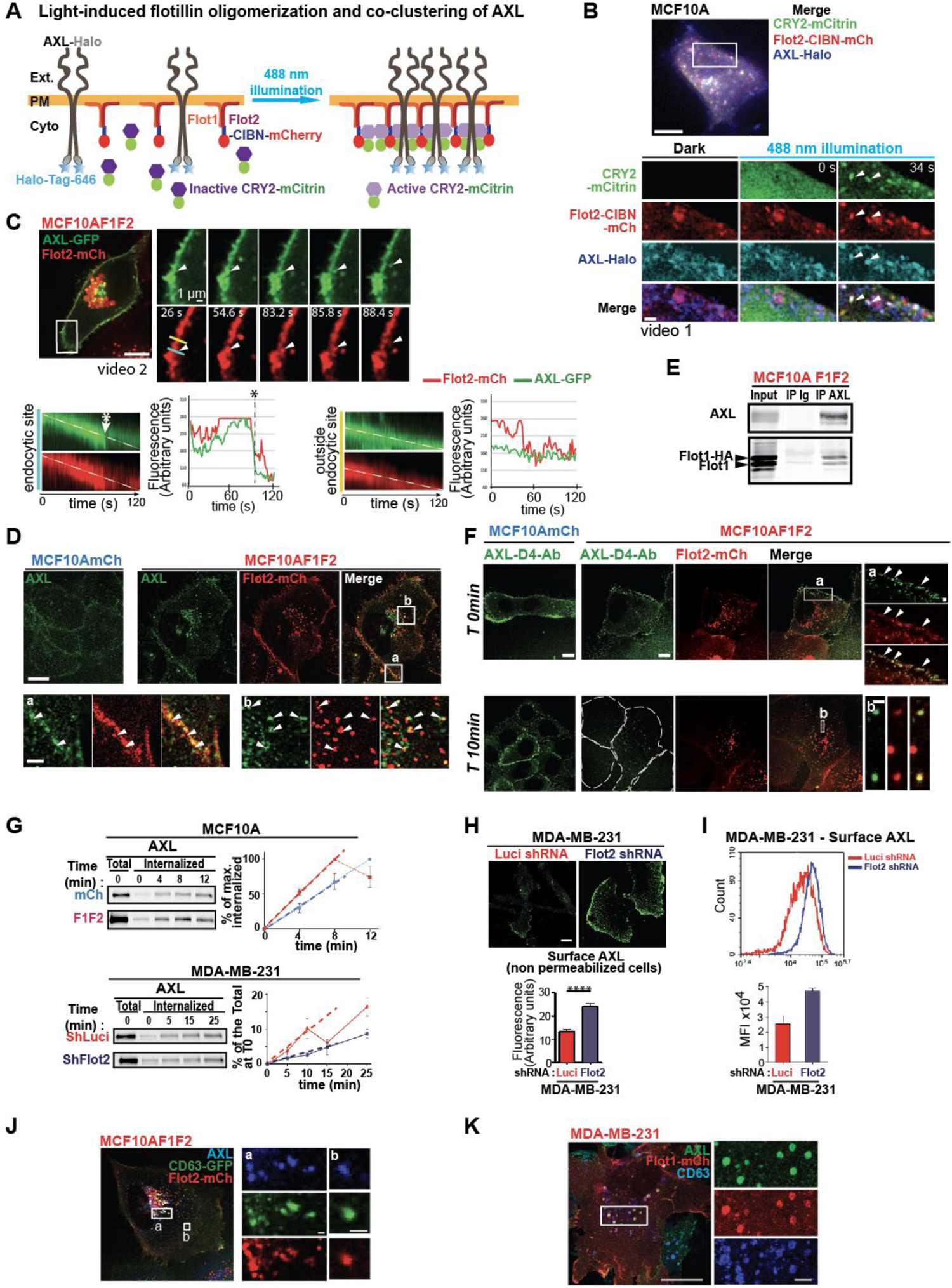
AXL is a cargo of the UFIT pathway. **A)** *Schema describing the optogenetic system used to induce flotillin oligomerization.* Flotillin 2-CIBN-mCherry, CRY2-mCitrin and AXL-Halo labeled with Halo-Tag-Janelia Fluor 646 were co-expressed in MCF10A cells that express endogenous flotillins. Light illumination (488nm) induces the oligomerization of “active” CRY2 and its binding to CIBN, allowing the oligomerization of flotillin 2-CIBN-mCherry with flotillin 1. AXL-Halo is trapped in the light-induced flotillin-microdomains. **B)** *Flotillin-oligomerization induced by a CRY2-CIBN-based optogenetic approach promotes co-clustering of AXL*. Still images from a TIRF-microscopy movie of a MCF10A cell that co-expresses CRY2-mCitrin, flotillin 2-CIBN-mCherry, and AXL-Halo labeled with Halo-Tag-Janelia Fluor 646. Representative image of a whole cell 15s after starting the 488 nm illumination to perform CRY2 binding to flotillin 2-CIBN-mCh and subsequent oligomerization (see also fig. S5 for the validation of the constructs and of the optogenetic approach). Magnified images from the boxed area taken before (dark) and after 488 nm illumination. Arrowheads indicate clusters of CRY2-mCitrin and flotillin 2-CIBN-mCherry where AXL-Halo accumulates (video 1). Images are representative of five dependent experiments. **C)** *Co-accumulation of AXL and flotillin 2 at the PM before their co-endocytosis*. Still images of a time-lapse series of one MCF10AF1F2 cell that expresses AXL-GFP to show flotillin 2/AXL co-localization at the PM and their co-endocytosis at PM sites (showed by arrowheads). Kymographs show the temporal variation of the GFP and mCherry signals along two line-scans (blue at the PM endocytic site, and yellow close to the PM endocytic site). Graphs show the intensity variation of both signals along the dotted white lines on the kymographs. The left kymograph shows the accumulation of AXL-GFP and flotillin 2-mCherry at the PM followed by a sudden concomitant drop of both signals (at around 83s, indicated by the star). The right kymograph shows that in the meantime, both AXL-GFP and flotillin 2-mCherry signals remained stable at the PM outside and close to this endocytic site (video 2). **D)** *Flotillin upregulation modifies AXL cell distribution.* Distribution of endogenous AXL in MCF10AmCh and MCF10AF1F2 cells analyzed by confocal microscopy after immunofluorescence staining. Higher magnification images of the two boxed regions show AXL colocalization with flotillin 2-mCherry at the PM (a) and in intracellular vesicles (b). White arrows indicate some vesicles positive for both AXL and flotillin 2-mCherry. **E)** *Flotillin 1 co-precipitates with AXL.* AXL was immunoprecipitated from MCF10AF1F2 cell lysates, and flotillin 1 co-immunoprecipitation was assessed by immunoblotting. Control immunoprecipitation was performed with non-relevant immunoglobulins (Ig). Results shown are representative of 3 independent experiments. **F)** *Flotillin upregulation accelerates AXL endocytosis* towards flotillin positive endosomes. MCF10AmCh and MCF10AF1F2 cells were incubated with the anti-AXL D4 antibody at 4°C followed by incubation at 37°C for 10 min to allow AXL internalization. AXL distribution and flotillin 2-mCherry signal were analyzed by immunocytochemistry. Arrowheads indicate AXL/flotillin 2 colocalization. Images are representative of at least 5 independent experiments. Scale bars are 10 µm in the main images, and 1 µm in the magnified images from the insets. **G)** *Flotillin upregulation accelerates AXL endocytosis.* Kinetics of AXL internalization in MCF10AmCh and MCF10AF1F2 cells (up), and in MDA-MB-231shLuci and MDA-MB-231shFlot2 cells (bottom). Surface proteins were labeled with biotin at 4°C, and cells were incubated at 37°C for the indicated times to allow endocytosis. AXL presence in the internalized biotinylated fraction was probed by western blotting using relevant antibodies and quantified as the percentage of the maximum level of internalized protein (reached at 12 and 8 min, in MCF10AmCh and MCF10AF1F2 cells respectively). Results are expressed as the percentage of the maximum level of surface labaled AXL at T0 and are the mean ± SEM of 4 independent experiments (all cell lines). The AXL internalization rates (dashed lines) were 8.48 ± 1.28 and 12.43 ± 0.48 %/min in MCF10AmCh and MCF10AF1F2 cells, respectively, revealing a significant 1.45-fold increase (p=0.0088) upon flotillin upregulation. Comparison of the AXL internalization rates (dashed lines) in MDA-MB-231shLuci and MDA-MB-231shFlot2 cells showed a significant 3.66-fold decrease (p=0.0009) upon flotillin downregulation. **H, I)** *AXL level at the cell surface is increased in MDA-MB-231 cells in which flotillins were knocked down.* **(H)** In fixed non-permeabilized cells, AXL expression was analyzed using the monoclonal D4 antibody against AXL extracellular domain and an Alexa488-conjugated anti-human secondary antibody. Images were acquired in the same illumination conditions for MDA-MB-shLuci cells (control) and MDA-MB-231-shFlot2cells (in which flotillins were knocked down). Using the Fiji software, the mean fluorescence intensity was quantified in each area delimited by the cell outline (n=53 MDA-MB-shLuci cells and n=41 MDA-MB-231-shFlot2cells). Values were expressed as arbitrary units and correspond to the mean ± SEM. **(I)** AXL surface staining analyzed by FACS. Fixed, non-permeabilized cells were labeled with the D9 anti-AXL antibody and a Alexa488-conjugated secondary antibody and analyzed by FACS. Results are expressed as the Mean Fluorescence Intensity (MFI) from 3 independent experiments. **J,K)** *AXL is present in flotillin- and CD63-positive vesicles.* Immunofluorescence analysis by confocal microscopy of endogenous AXL expression in MCF10AF1F2 cells that express CD63-GFP **(J)** and of endogenous AXL and CD63 in MDA-MB-231 cells that express flot1-mCherry **(K)**. Higher magnification images of the boxed regions show the presence of AXL in flotillin 2-mCherry-(J) or flotillin 1-mCherry-positive vesicles **(K)** and CD63-positive vesicles. Images are representative of three independent experiments. In D, H, J, K, scale bars are equal to 10 µm for the main images and to 2 µm for the magnified images from the insets.

**Figure S5:**
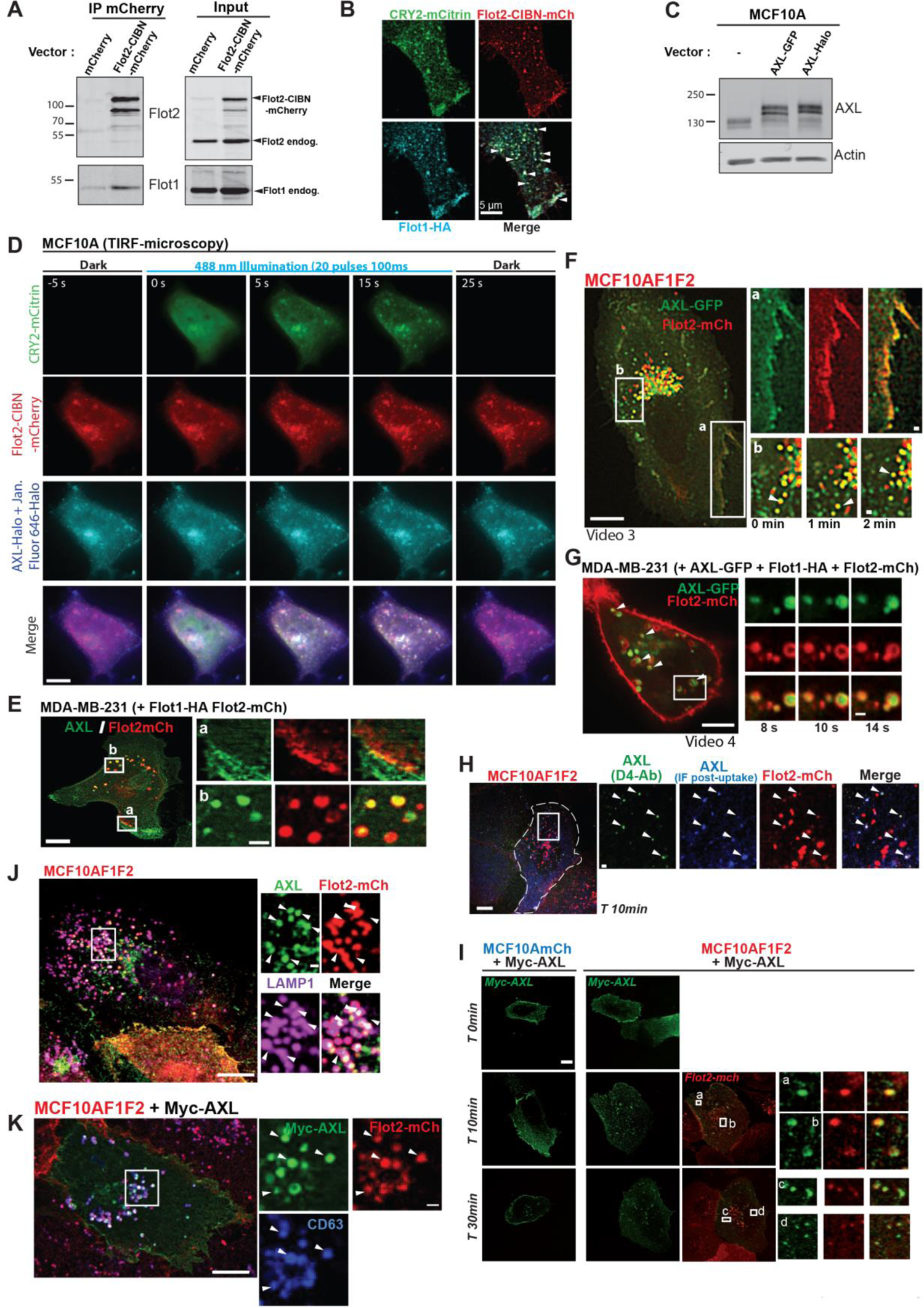
Co-distribution of AXL and flotillins at the plasma membrane and in late endosomes. **A)** *Flotillin2-CIBN-mCherry interacts with flotillin 1.* Immunoprecipitations using an antibody against mCherry (mCherry trap) and lysates from MCF10A cells that express mCherry alone or flotillin 2-CIBN-mCherry. Immunoprecipitates were then analyzed with antibodies to detect flotillin 1 and 2. Flotillin 1 was co-immunoprecipitated only in lysates from cells that express flotillin 2-CIBN-mCherry, but not mCherry alone. Results are representative of 3 independent experiments. **B)** *Light-induced flotillin 2-CIBN-mCherry microdomains contain flotillin 1.* MCF10A cells that co-express flotillin 2-CIBN-mCherry, flotillin 1-HA and CRY2-mCitrin were illuminated at 488 nm for 1 min, fixed, incubated with an anti-HA antibody and imaged by confocal microscopy. CRY2-mCitrin patches containing flotillin 2-CIBN-mCherry (white arrows) also exhibit flotillin 1-HA signal. Results are representative of 3 independent experiments. Scale bar: 5 µm. **C)** *Characterization of the AXL-GFP and AXL-Halo constructs.* MCF10A cells were transfected with empty vector (-) or plasmids encoding AXL-GFP or AXL-Halo. Expression of the proteins of the expected size (138+28=166 kDa for AXL-GFP) and (138+28=171 kDa for AXL-Halo) is shown. A doublet was observed for each tagged protein likeendogenous AXL. **D)** Still images from a TIRF-microscopy video of one MCF10A cell that co-expresses CRY2-mCitrin, flotillin 2-CIBN-mCherry and AXL-Halo labeled with Halo-Tag-Janelia Fluor 646. The cell was imaged before, during, and after 488 nm illumination (22 pulses of 100 ms each every second). **E)** Distribution of endogenous AXL in MDA-MB-231 cells analyzed by confocal microscopy after immunofluorescence staining. Higher magnification images of the two boxed regions show AXL colocalization with flotillin 2-mCherry at the PM (a) and in intracellular vesicles (b). **F)** *Co-trafficking of AXL and flotillins in MCF10AF1F2 cells.* Still images of a representative time-lapse series (video 3) of a MCF10AF1F2 cell that expresses AXL-GFP to illustrate the co-localization of AXL-GFP and flotillin 2-mCherry at the PM (box a), and in intracellular vesicles (box b) and their co-trafficking. The white arrow allows following one vesicle over time. **G)** *Co-trafficking of AXL and flotillins in MDA-MB-231 cells.* MDA-MB-231 cells were transfected with AXL-GFP, flotillin 2-mCherry and flotillin 1-HA-encoding plasmids. Live cells were imaged by spinning disk confocal microscopy to showthe co-localization (indicated by white arrows) and the co-trafficking of AXL-GFP and flotillin 2-mCherry in intracellular vesicles (video 4). **H)** MCF10AmCh and MCF10AF1F2 cells were incubated with the anti-AXL D4 antibody (against AXL extracellular domain) at 4°C followed by incubation at 37°C for 10 min to allow AXL internalization. After fixation and permeabilization, internalized AXL was labeled (green) with a FITC-conjugated secondary antibody directed against the D4 antibody. Immunofluorescence staining of AXL (blue) was also performed using an antibody against AXL cytoplasmic domain and an Alexa-633 conjugated secondary antibody. Arrowheads indicate flotillin 2-mCherry-positive vesicles in which AXL is internalized (green). **I)** MCF10AmCh and MCF10AF1F2 cells that express Myc-tagged AXL were incubated with an anti-Myc antibody at 4°C followed by incubation at 37°C for the indicated times. Myc-AXL distribution was analyzed by immunocytochemistry using an Alexa488-conjugated secondary antibody. **J)** In MCF10AF1F2 cells, endogenous AXL is localized in flotillin 2-mCherry vesicles positive for the endolysosomal marker LAMP1 (Confocal microscopy image). **K)** Myc-tagged AXL expressed in MCF10AF1F2 cells is localized in flotillin 2-mCherry vesicles positive for the late endosomal marker CD63 labelled by immunofluorescence (Confocal microscopy image). Images shown in B, D - K are representative of 3 to 5 independent experiments. In E, F, G, H, I, J, and K scale bars are equal to 10 µm in the main images and to 2 µm in the magnified images from the insets.

**Figure S6:**
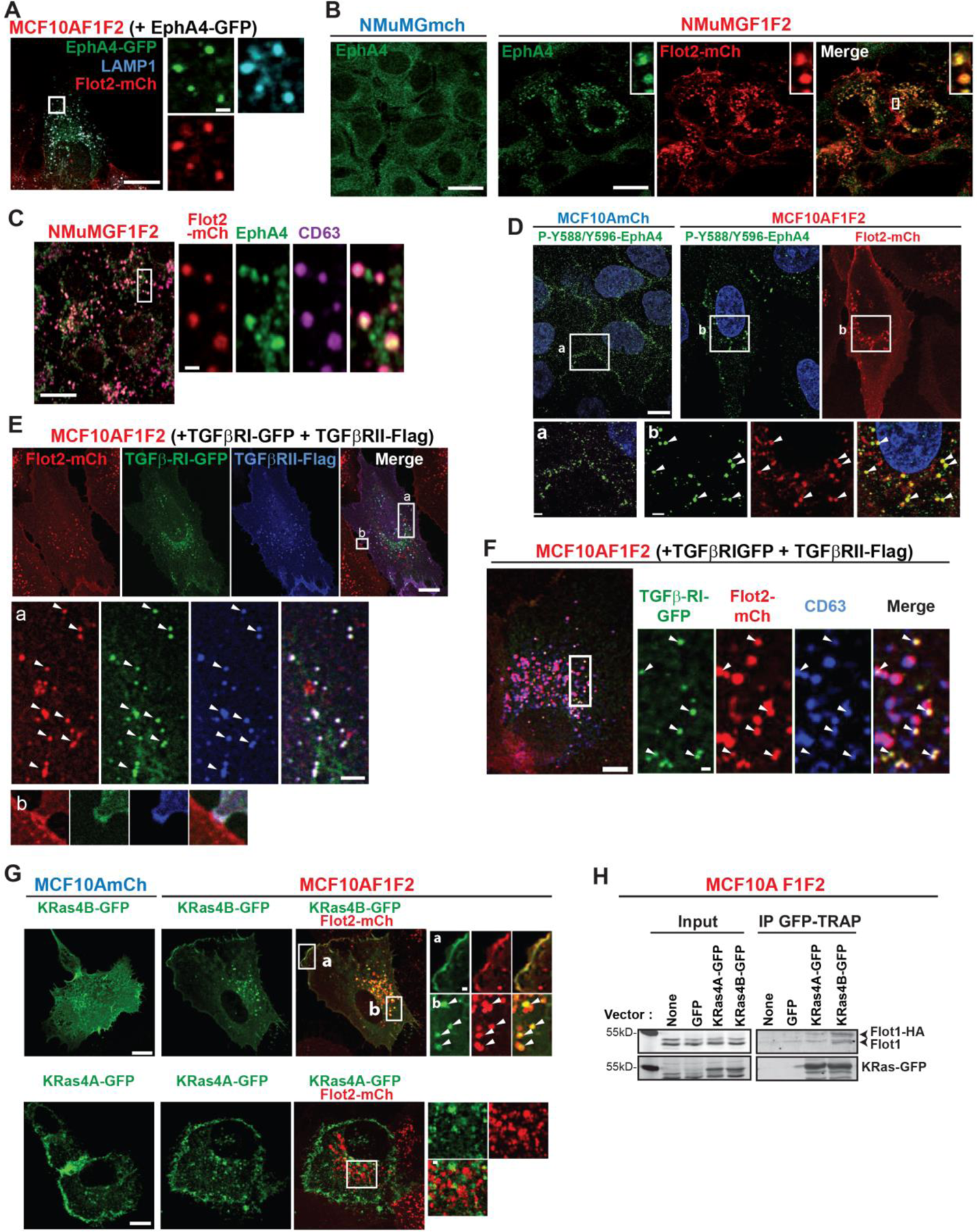
The RTK EphA4, the TGFβ receptors, and Kras4B are located in flotillin-positive late endosomes. **A)** Confocal microscopy image of one MCF10AF1F2 cell that expresses EphA4-GFP and stained for LAMP1, showing flotillin 2/EphA4 co-localization in LAMP-1-positive vesicles. **B, C)** Immunofluorescence analysis of NMuMGmCh and NMuMGF1F2 cells to detect endogenous EphA4. In NMuMGF1F2 cells, EphA4 relocated in intracellular vesicles, many of which were flotillin 2-mCherry and CD63-positive (C). **D)** MCF10AmCh and MCF10AF1F2 cells were analyzed by immunofluorescence using an antibody against EphA4 phosphorylated on Y588/Y596. In MCF10AmCh cells the signal was at cell-cell contacts (magnified image from the boxed area (a)), whereas it was concentrated in intracellular vesicles in MCF10AF1F2 cells where it co-localized with flotillin 2-mCherry (magnified images from boxed areas (b)). **E)** MCF10AF1F2 cells that co-express the TGFβ-RI (GFP tagged) and TGFβ-RII (Flag tagged) subunits were analyzed by immunofluorescence using an anti-Flag antibody and imaged by confocal microscopy. Both subunits were concomitantly found at the PM and in intracellular vesicles (white arrows) where they colocalized with flotillin 2-mCherry. **F)** MCF10AF1F2 cells that co-express TGFβ-RI-GFP and TGFβ-RII-Flag were stained for CD63 and imaged by confocal microscopy. White arrows in the magnified images of the boxed area indicate co-localization of TGFβ-RI-GFP, CD63 and flotillin 2-mCherry in intracellular vesicles. Scale bars: 10µm in the main images and 1 µm in the magnified regions. **G)** *Flotillin upregulation specifically relocates K-Ras4B to flotillin-positive endosomes.* Confocal images of MCF10AmCh and MCF10AF1F2 cells that express K-Ras4A-GFP or K-Ras4B-GFP. Unlike K-Ras4A-GFP (lower panels), K-Ras4B-GFP (upper panels) co-localized with flotillins and its distribution was modified in MCF10AF1F2 cells compared with MCF10AmCh cells (upper panel). K-Ras4B-GFP was mainly present at the PM in MCF10AmCh cells. Conversely, in MCF10AF1F2 cells, it remained visible at the PM where it co-localized with flotillin 2-mCherry (magnified images from the boxed region (a)), but it was also strongly recruited to flotillin 2-mCherry-positive vesicles and co-trafficked with flotillin 2 (magnified images of the boxed region (b)). In A, B, C, D, E, G, scale bars are equal to 10 µm in the main images and 2 µm in the magnified images from insets. In panels A-G, images shown are representative of several cells observed in 3 independent experiments. **H)** *K-Ras4B forms a complex with flotillin 1.* Lysates from MCF10AF1F2 cells that express GFP alone, K-Ras4A-GFP, or K-Ras4B-GFP were immunoprecipitated with GFP-TRAP. Immunoprecipitates were probed by western blotting using anti-flotillin 1 and anti-pan-Ras antibodies. Both K-Ras isoforms were similarly expressed and immunoprecipitated. The signal for flotillin 1-HA and flotillin 1 in precipitates from K-Ras4B-GFP-expressing cells was clearly higher than the signal observed in precipitates from K-Ras4A-GFP-expressing cells that was similar to the background signal found in precipitates from GFP-expressing cells. Results are representative of 3 independent experiments.

About 40 % of the flotillin 2-mCherry positive vesicles were positive for AXL-GFP, and half of AXL-GFP-containing vesicles were positive for flotillin 2-mCherry (fig. S3C). In contrast, vesicles positive for the membrane receptor CD71, expression of which remained unchanged upon flotillin upregulation (fig. S3A), co-localized less frequently with flotillin-positive endosomes (fig. S3C). Altogether, these findings showed that flotillins and AXL are present in the same PM domains, from where they are co-endocytosed towards intracellular vesicles where they are co-accumulated. Similarly, in breast tumor samples, flotillin 1 and AXL accumulated in intracellular vesicles in malignant cells (fig. S4C). This co-distribution of flotillins and AXL is in agreement with the detection of flotillin 1 and AXL complexes by co-immunoprecipitation experiments in MCF10AF1F2 cells (fig. 4E).

To monitor the impact of flotillin upregulation on AXL endocytosis, we performed an AXL uptake assay using the D4 antibody against the extracellular domain of human AXL, coupled to immunocytochemistry detection (see validation in fig. S5H). Straight after incubation with the D4 antibody, we could detect cell surface AXL signals in MCF10AmCh and MCF10AF1F2 cells (fig. 4F). After 10 min of internalization, the AXL-D4 signal was clearly internalized in flotillin-positive endosomes in MCF10AF1F2 cells (fig. 4F). Conversely, it was still mostly at the surface in MCF10AmCh cells. We obtained similar results using an uptake assay where AXL surface expression was monitored with an anti-Myc antibody in cells transfected with N-terminally Myc-tagged AXL (fig. S5I). These observations confirmed that AXL initially located at the PM was addressed, after endocytosis, towards flotillin-positive late endosomes. Furthermore they suggested that AXL internalization occurs faster in cells in which flotillins are upregulated. To better quantify the impact of the UFIT-pathway on AXL internalization, we used a cell surface biotinylation assay. AXL internalization rate was significantly increased in MCF10AF1F2 cells compared with MCF10AmCh cells (fig. 4G, upper panels). Reciprocally, to analyze the impact of flotillin downregulation on AXL internalization in MDA-MB-231 cells, we used the MDA-MB-231 shFlot2 cell line that we generated ^18^. As previously described, down regulation of flotillin 2 was sufficient to decrease the expression of both flotillins 1 and 2. Downregulation of flotillins in MDA-MB-231 slowed down AXL internalization (fig. 4G, lower panels) and AXL accumulated at the PM in shFlot2-MDA-MB-231 cells (fig. 6H, I). In contrast, CD71 internalization was not affected by flotillin upregulation (fig. S3D), in agreement with its low presence in flotillin-positive vesicles compared to AXL (fig. S3C), indicating that flotillin upregulation does not affect the endocytosis and trafficking of all membrane receptors.

Flotillin-positive vesicles, in which flotillins 1 and 2 are present, are LAMP-1-, Rab7a- and CD63-positive late endosomes ^18^. Furthermore, we previously characterized flotillin-positive vesicles as non degradative late endosomes on the basis of their very low staining for fluorescent clived DQ-BSA ^18^. In agreement with AXL stabilization in flotillin-upregulated cells, we found AXL in flotillin-, CD63- and LAMP1-positive vesicles in MCF10AF1F2 cells [analyzing either endogenous AXL (fig. 4J, and S5J) and Myc-tagged AXL (fig. S5K)] and in MDA-MB-231 cells (fig. 4K). These results correlate with the recent work by Zajac et al showing the presence of AXL in LAMP1-positive vesicles in mammary tumor cells ^33^. Besides, we found in flotillin-positive late endosomes other signaling transmembrane receptors and oncogenic molecules that participate in tumorigenesis, such as EphA4 and TGFβ (fig. S6A-F), and also specifically K-Ras4B (fig. S6G). Flotillin upregulation modified the distribution of only K-Ras4B but not of the other H-, N- and K-Ras4A isoforms. K-Ras4B was localized at the PM in MCF10AmCh cells, and in flotillin-positive vesicles in MCF10AF1F2 cells (fig. S6G). This correlated with the co-immunoprecipitation of K-Ras4B, but not of K-Ras4A, with flotillin 1 (fig. S6H).

Altogether, these results showed that AXL is a cargo of the UFIT pathway. Flotillin-upregulation accelerated AXL internalization and promoted AXL endocytosis toward flotillin-positive late endosomes.

### Sphingosine kinase 2 is required for AXL stabilization through the UFIT pathway

The previous results showed that AXL stabilization is correlated with its accelerated internalization and sorting towards flotillin-positive late endosomes. To identify the molecular mechanisms linking these two processes we focused on sphingolipids, metabolism of which is linked to endocytosis, and also to flotillins. Indeed, flotillins bind to sphingosine, the sphingosine-1-phosphate (S1P) precursor, and the level of S1P is reduced in cells from flotillin knock-out mice ^34^. The bioactive lipid S1P is implicated in endocytosis and membrane remodeling ^35–37^, and S1P production, induced upon sphingosine phosphorylation by sphingosine kinase (SphK) 1 or 2, has been linked to cancer progression ^38^. SphK 1/2 subcellular localization and consequently the compartmentalization of S1P production play an important role in their function ^39^. We hypothesized that flotillin rich regions of membranes can concentrate sphingosine, where SphK could be recruited to locally generate S1P.

As there is no probe to examine S1P distribution, we examined the localization of SphK isoforms in MCF10AF1F2 and MDA-MB-231 cells. We expressed GFP-tagged SphK1 and 2 because no antibody is available for immunofluorescence staining. In both cell lines, SphK1-GFP was present in the cytosol, at the PM, and in intracellular vesicles among which less than 20% were flotillin-positive (fig. 5A and B and S7A). Conversely, we detected SphK2-GFP in approximately 70% of flotillin-positive vesicles (fig. 5A-D, video 5). SphK2- and flotillin-positive vesicles also contained AXL (fig. 5C, video 6), and the late endosomal markers Rab7 (fig. S7B) and CD63 (fig. S7C). SphK2 and flotillin 2 also colocalized at discrete sites of the PM, from which we could observe the fast co-endocytosis of both proteins (fig. 5D, videos 7 and 8). Flotillin upregulation increased SphK2 recruitment to CD63-positive late endosomes (fig. 5E), without modifying its protein expression level (fig. S7E).

**Figure 5:**
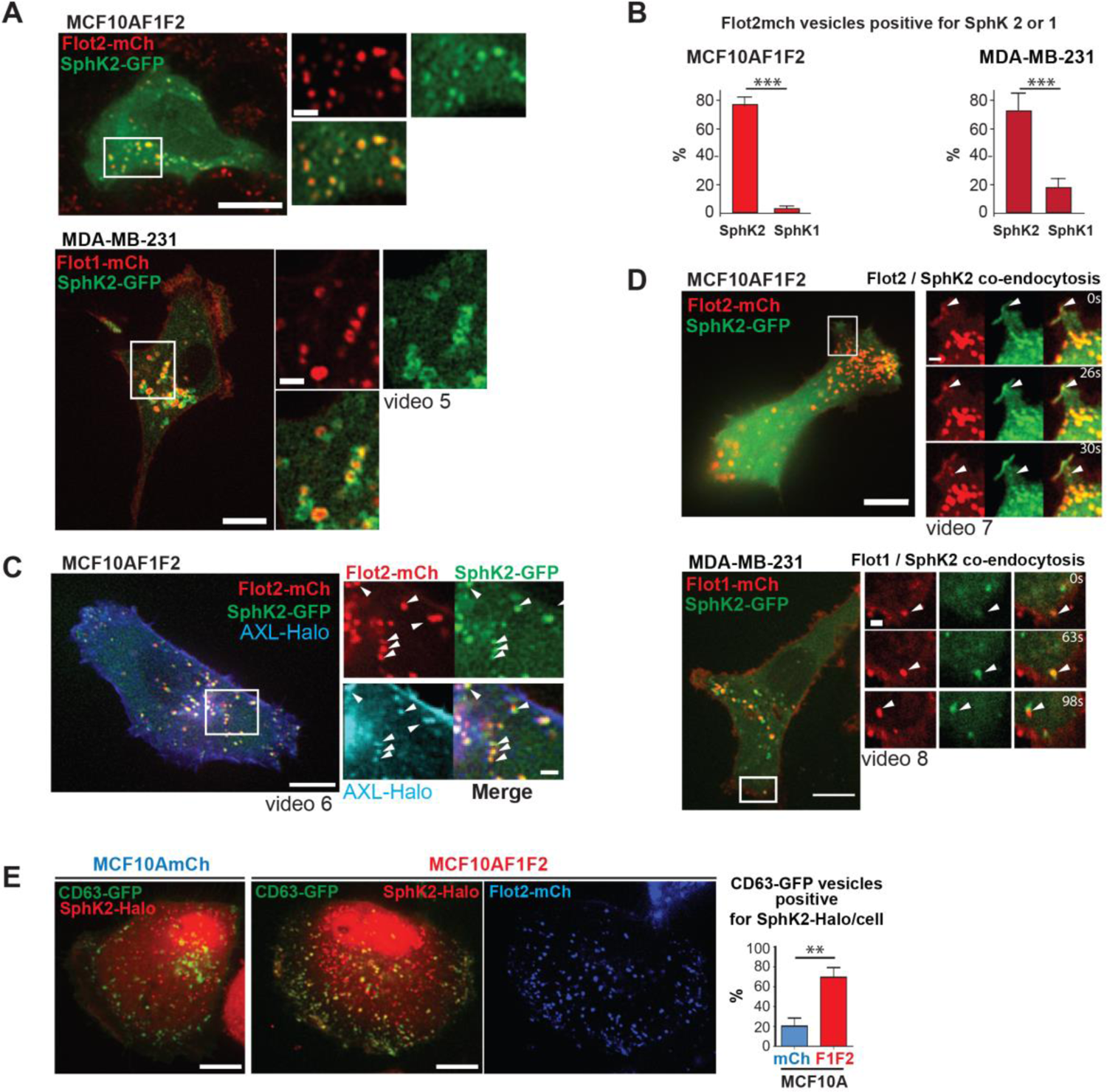
SphK2 is localized at flotillin-positive endocytic sites and at flotillin-positive late endosomes where AXL is present. **A, B, D)** *SphK2 is enriched in flotillin-positive late endosomes and co-accumulates at the PM with flotillins before endocytosis.* MCF10AF1F2 cells that express SphK2-GFP (A,B) and MDA-MB-231 cells that co-express flotillin 1-mCherry or flotillin 2-mCherry and SphK2-GFP (A,B,D) were imaged by spinning disk confocal microscopy (see also videos 5, 6, 7, 8). (**A, B**) *SphK2-GFP, but not SphK1-GFP, is abundantly present in flotillin 2-mCherry-positive vesicles* (see also fig. S7A). Quantification of flotillin 2-mCherry-positive vesicles in which SphK2-GFP or SphK1-GFP is detected (B). Results are expressed as the percentage of vesicles and are the mean ± SEM (n=300 vesicles in 10 MCF10AF1F2 cells, and n=100 vesicles in 10 MDA-MB-231 cells that co-express flotillin 2-mCherry and SPK2 or SPK1-GFP). (D) *Fast co-endocytosis of flotillin 2-mCherry and SphK2-GFP following their co-accumulation at the plasma membrane* (indicated by the arrows) in MCF10AF1F2 cells (upper panel, video 7) and MDA-MB-231 cells (lower panel, video 8). Shown are still images from spinning disk confocal videomicroscopy. **C)** *AXL is present in flotillin- and SphK2-positive vesicles.* Still image from spinning disk confocal microscopy of live MCF10AF1F2 cells that co-express SphK2-GFP and AXL-Halo labeled with Halo-tag-Janelia Fluor 646. Arrows show flotillin-positive vesicles that contain SphK2-GFP and AXL-Halo (video 6). Image shown is representative of several cells observed in 3 independent experiments. **E)** *SphK2 localization in CD63-positive endosomes is increased in MCF10AF1F2 cells.* MCF10AmCh and MCF10A F1F2 cells were co-transfected with CD63-GFP and SphK2-Halo labeled with Halo-Tag-Janelia Fluor 646. Representative images acquired by spinning disk confocal microscopy of live cells. To better visualize the CD63-GFP and SphK2-Halo co-localization, the SphK2-Halo and flotillin 2-mCherry signals were pseudocolored in red and blue, respectively. The presence of SphK2-Halo in CD63-GFP-positive vesicles was quantified (n=372 vesicles in 10 MCF10AmCh cells, and n=455 vesicles in 10 MCF10AF1F2 cells). Results are expressed as the percentage per cell of CD63-GFP labelled vesicles positive for SphK2-Halo. ***P<0.001, ** P<0.01 In A, C, D, E, Scale bars are equal to 10 µm in the main images and 2 µm in the magnified images frome the boxed areas.

Due to SphK2 enrichment in flotillin-positive membrane domains and flotillin-positive vesicles, we asked whether SphK2 was involved in AXL stabilization upon flotillin upregulation. We first incubated cells with opaganib (ABC294640), a selective SphK2 inhibitor ^34^. Few minutes after opaganib addition, we observed the dissociation of SphK2-GFP from flotillin-positive endosomes that lasted for several hours (fig. S7D, video 9). Incubation of MCF10AF1F2 or MDA-MB-231 cells with opaganib for only 4h did not have any impact on AXL expression level (fig. 6A and D). However, in MCF10AF1F2 cells incubated with opaganib for 10h, AXL level was reduced to that observed in MCF10AmCh cells (fig. 6A).

**Figure 6:**
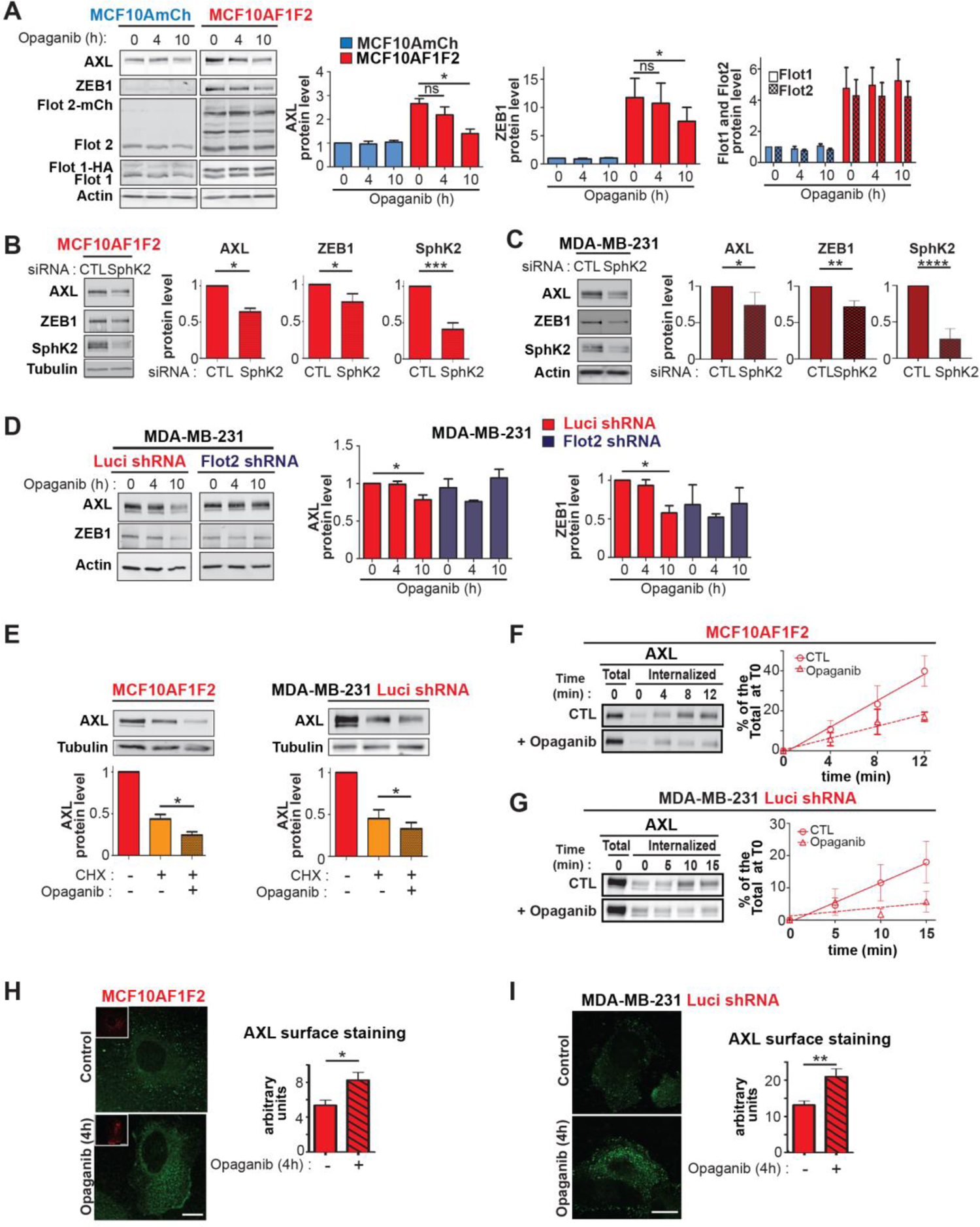
Sphingosine kinase 2 controls AXL endocytosis rate and its stabilization. **A)** *SphK2 inhibition decreases AXL and ZEB1 levels in MCF10AF1F2 cells.* MCF10AmCh and MCF10AF1F2 cells were incubated with opaganib (50 µM) for the indicated times. AXL, ZEB1, flotillin 1 and 2 were analyzed by western blotting in whole cell lysates. Results correspond to the level of each protein normalized to the actin signal, expressed as the fold increase compared to control (MCF10AmCh cells) and are the mean ± SEM of 4 independent experiments. **B,C)** *SphK2 downregulation decreases AXL and ZEB1 levels in MCF10AF1F2 and MDA-MB-231 cells.* MCF10AF1F2 and (B) and *MDA-MB-231* cells were transfected with siRNAs against luciferase (CTL) or SphK2, and siRNA efficacy was tested by western blotting with antibodies against SphK2 and also AXL, ZEB1 and tubulin. Histograms show the level of each protein normalized to the tubulin signal, expressed as fold increase compared with CTL and are the mean ± SEM of 4 independent experiments.**,D)** *SphK2 inhibition decreases AXL and ZEB1 levels in MDA-MB-231 cells.* MDA-MB-231shLuci (control) and MDA-MB-231shFlot2 cells were incubated with opaganib (50 µM) for the indicated times. AXL, ZEB1 and actin were analyzed by western blotting in whole cell lysates. The histograms show the level of each protein normalized to the actin signal, expressed as fold increase compared with control and are the mean ± SEM of 5 independent experiments. **E)** *AXL decrease upon SphK2 inhibition is not due to protein synthesis inhibition.* MCF10AF1F2 and MDA-MB-231shLuci cells were incubated or not with opaganib for 10 hours. When indicated, cycloheximide (CHX) was added 4 hours after the beginning of opaganib incubation. AXL was analyzed by western blotting in whole cell lysates. The histograms show the level of AXL normalized to the tubulin signal, expressed as fold increase compared with control (no treatment) and are the mean ± SEM of 4 independent experiments. **F, G)** *SphK2 inhibition decreases AXL internalization rate in flotillin-upregulated cells.* Kinetics of AXL internalization in MCF10AF1F2 cells, and in MDA-MB-231shLuci either untreated (CTL) or pre-treated with opaganib (50µM, 4h). Surface proteins were labeled with biotin at 4°C, and cells were incubated at 37°C for the indicated times to allow endocytosis. The presence of opaganib was maintained during the experiments. AXL presence in the internalized biotinylated fraction was probed by western blotting using relevant antibodies. Results are expressed as the percentage of the maximum level of surface labeled AXL at T0 and are the mean ± SEM of 4 independent experiments. In MCF10AF1F2 cells, the AXL internalization rates were 3.30 ± 0.66 and 1.49 ± 0.39 %/min in control and opaganib treatment conditions respectively, revealing a significant 2.2-fold decrease (p=0.0025) upon opaganib treatment. In MDA-MB-231 shLuci cells, the AXL internalization rates were 1.22 ± 0.37 and 0.26 ± 0.24 %/min in control and opaganib treatment conditions respectively, revealing a significant 4.69-fold decrease (p=0.0399) upon opaganib treatment. **H, I)** *SphK2 inhibition increases AXL level at the cell surface in flotillin-upregulated cells.* Fixed non-permeabilized cells were immunostained for AXL using the monoclonal D4 antibody against AXL extracellular domain and an Alexa488-conjugated anti-human secondary antibody. Images of control- and opaganib-treated cells (50µM, 4h) were acquired in the same conditions. Representative images are shown (for MCF10AF1F2 cells, the insets show flotillin 2-mCherry signal). The mean fluorescence intensity was quantified in the area defined by the outline of each individual cell. Results are expressed as arbitrary units and are the mean ± SEM of 23 measurements for MCF10AF1F2 cells (I), and 22 measurements for MDA-MB-231shLuci cells (J) for each condition (control and opaganib). Scale bars: 10 µm for the main images and 2 µm in the magnified images from the boxed area. *P<0.05; **P<0.01.

**Figure S7:**
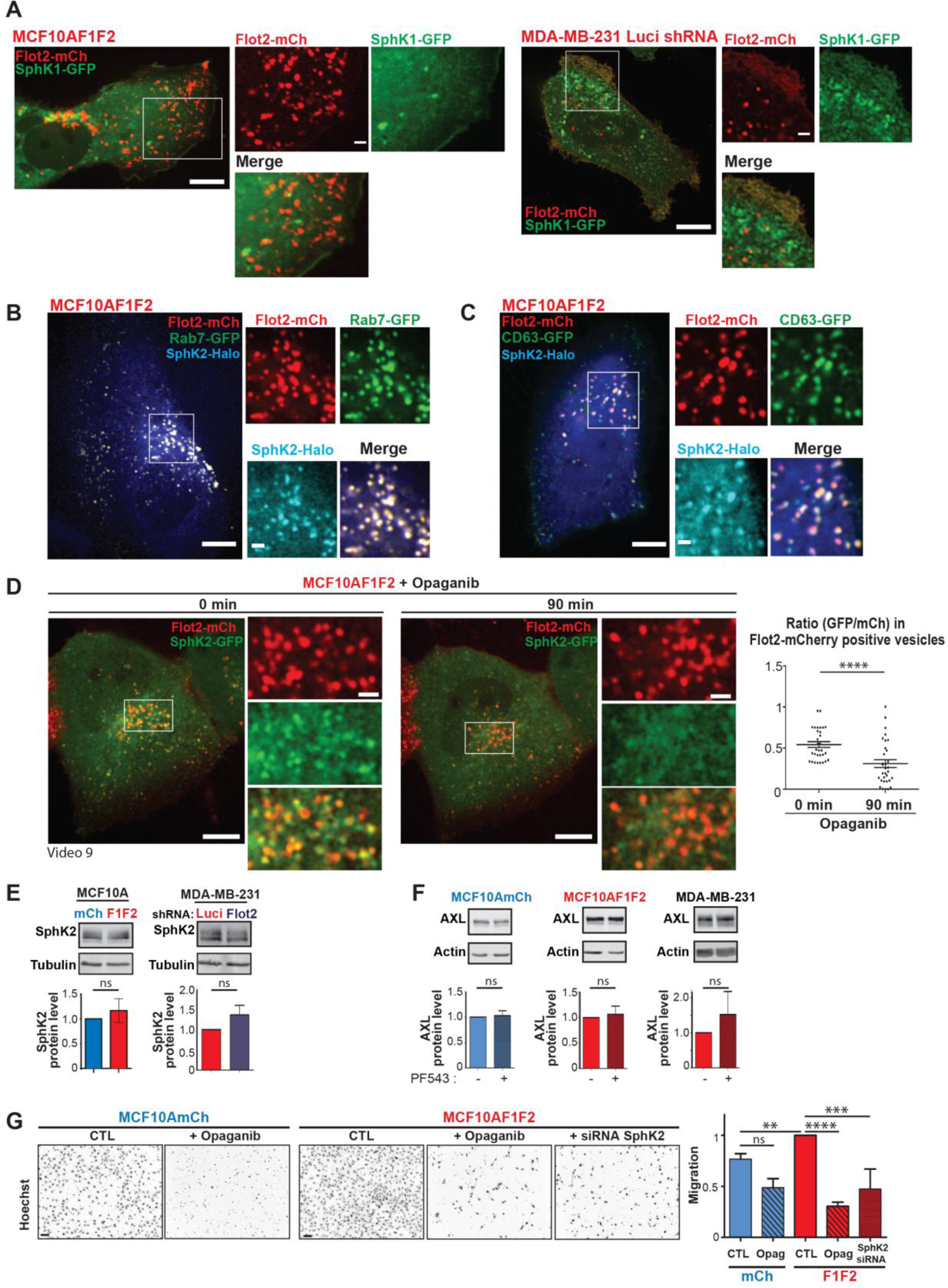
SphK1 and 2 localization and effect of SphK inhibitors. **A)** *SphK1 is barely detected in flotillin-positive endosomes.* Live MCF10AF1F2 cells that express SphK1-GFP (left), and MDA-MB-231 cells that co-express flotillin 1-mCherry and SphK1-GFP (right) were imaged by spinning disk confocal microscopy (see quantification in fig. 7B). **B, C)** *SphK2 colocalizes with flotillins in Rab7 and CD63 positive vesicles.* Still images of live MCF10AF1F2 cells that express SphK2-Halo labeled with Halo-tag-Janelia Fluor 646 and Rab7-GFP (B), or CD63-GFP (C) imaged by spinning disk confocal microscopy. Images shown are representative of 3 independent experiments. **D)** *Opaganib treatment induces the delocalization of SphK2 from flotillin-positive vesicles.* Live MCF10AF1F2 cells that express SphK2-GFP were imaged right after incubation with opaganib by spinning disk confocal microscopy every 15 min. The SPK2-GFP signal associated with flotillin 2- mCherry positive vesicles was clearly decreased after 90 min of incubation. The histogram shows the GFP/mCherry signal ratio in flotillin 2-mCherry vesicles at 0 and after 90 min of opaganib incubation (31 vesicles at T0 and 32 vesicles at T90 from 3 independent cells). **** p<0.0001 In panel A-D, scale bars are equal to 10 µm in the main images, and to 2 µm in the magnified images from the boxed area. **E)** *Flotillin level does not modulate SphK2 protein expression.* SphK2 and tubulin expression were assessed in whole cell extracts from MCF10AmCh, MCF10AF1F2, MDA-MB-231shLuci, and MDA-MB-231shFlot2 cells by western blotting using specific antibodies. The histograms show SphK2 level (normalized to tubulin) expressed as fold-increase compared with MCF10AmCh or MDA-MB-231shLuci cells, and data are the mean ± SEM of 4 independent experiments. **F)** *SphK1 inhibition does not influence AXL level in cells in which flotillins are upregulated.* MCF10AmCh, MCF10AF1F2 and MDA-MB-231 cells were incubated with the selective Sphk1 inhibitor PF543 (10 µM, 24h). AXL and actin levels were measured by western blotting in cell lysates. The histograms show AXL level, expressed as fold-increase compared with the control condition (no PF543), and data are the mean ± SEM of 4 independent experiments. **G)** *Boyden chamber migration assay.* MCF10AmCh and MCF10AF1F2 cells were seeded in the top of Boyden chamber inserts in serum-free medium, and serum-containing medium, acting as chemoattractant, was placed in the bottom well. When indicated, opaganib (50 µM) was added in the upper and lower chambers or MCF10AF1F2 were transfected with siRNAs against SphK2 for 48h prior seeding. Representative inverted-microscopy images of Hoechst-stained nuclei of cells that migrated through the pores. The histogram shows the amount of cells that migrated compared with untreated MCF10AF1F2 cells, and data are the mean ± SEM of at least 4 independent experiments. Scale bar: 50µm. **P<0.01, ****P<0.0001.

AXL was similarly decreased in MCF10F1F2 cells and MDA-MB-231 cells where SphK2 was efficiently knocked down using siRNA (fig. 6B,C). In MDA-MB-231shLuci cells that express high flotillin levels, SphK2 inhibition with opaganib for 10h also led to AXL decrease (fig. 6D). In both MCF10AF1F2 and MDA-MB-231shLuci cells, SphK2 inhibition decreased also ZEB1 level (fig. 6A-D), in agreement with our finding that its expression is dependent on AXL (fig. 3D and E). Of note, SphK2 inhibition did not alter flotillin 1 and 2 level (fig. 6A) in MCF10AmCh and MCF10AF1F2 cells (fig. 6A) or AXL level in MCF10AmCh cells (fig. 6A) and in MDA-MB-231shFlot2 cells ^18^ (fig. 6D). This indicates that SphK2 activity is required for maintaining a high level of AXL specifically in the context of flotillin upregulation. We also investigated the potential involvement of SphK1. However, in agreement with the limited localization of SphK1-GFP in flotillin-positive vesicles (fig. 5B and fig. S7A), SphK1 inhibition using PF543 had no effect on AXL levels in MCF10AF1F2 and MDA-MB-231 cells (fig. S7F).

To confirm that SphK2 was involved in AXL stabilization downstream of upregulated flotillins, we examined AXL levels in MCF10AF1F2 and MDA-MB-231shLuci cells incubated with CHX to inhibit protein synthesis. As expected CHX considerably decreased AXL level in both cell lines. This decrease was reinforced by the co-treatment with Opaganib (fig. 6E), demonstrating that SphK2 activity was involved in AXL stabilization, but not in stimulating its synthesis.

We next investigated how SphK2 was involved in AXL stabilization. As SphK2 co-accumulated with flotillins at the PM before endocytosis (fig. 5D, videos 7 and 8), we analyzed its participation in flotillin-mediated AXL endocytosis. We compared AXL internalization rate and cell surface level in cells incubated or not (control) with opaganib for only 4 hours. We chose this short treatment time when AXL decrease is not detectable yet (fig. 6A and D), because it allows a more straightforward analysis of AXL internalization and level at the cell surface. AXL internalization was significantly decreased when SphK2 was inhibited in both MCF10AF1F2 and MDA-MB-231shLuci cells (fig. 6F and G). In agreement, incubation with opaganib increased AXL expression at the cell surface in both cell lines (fig. 6H and I). This indicates that SphK2 participates in flotillin-mediated AXL endocytosis. In accordance with this finding that CD71 internalization is not influenced by flotillin level (fig. S3D), we found that it was also not altered by SphK2 inhibition (fig. S3E) confirming that it was not endocytosed through the UFIT pathway.

In conclusion, the data demonstrated that the lipid kinase SphK2, which co-localizes with flotillins at the PM and in late endosomes, possibly by locally generating S1P, is required to divert flotillin-mediated AXL endocytosis towards the UFIT pathway in order to promote its stabilization.

## Discussion

In this study, we demonstrated that flotillin upregulation in non-tumoral mammary cell lines devoid of oncogenic mutations is sufficient to activate several oncogenic signaling pathways, leading to EMT induction, a key step towards cancer cell invasion. Moreover, flotillin upregulation led to the stabilization of the RTK AXL through the UFIT pathway that promotes endocytosis of several cargos from the plasma membrane toward flotillin-positive late endosomes (see model fig. 7). The UFIT pathway is dependent on the lipid kinase SphK2, showing the role of sphingolipid metabolism on this flotillin-induced trafficking pathway. Our findings demonstrate the unprecedented role of flotillin-induced endosomal trafficking on AXL upregulation, a major feature of several invasive tumors.

**Figure 7:**
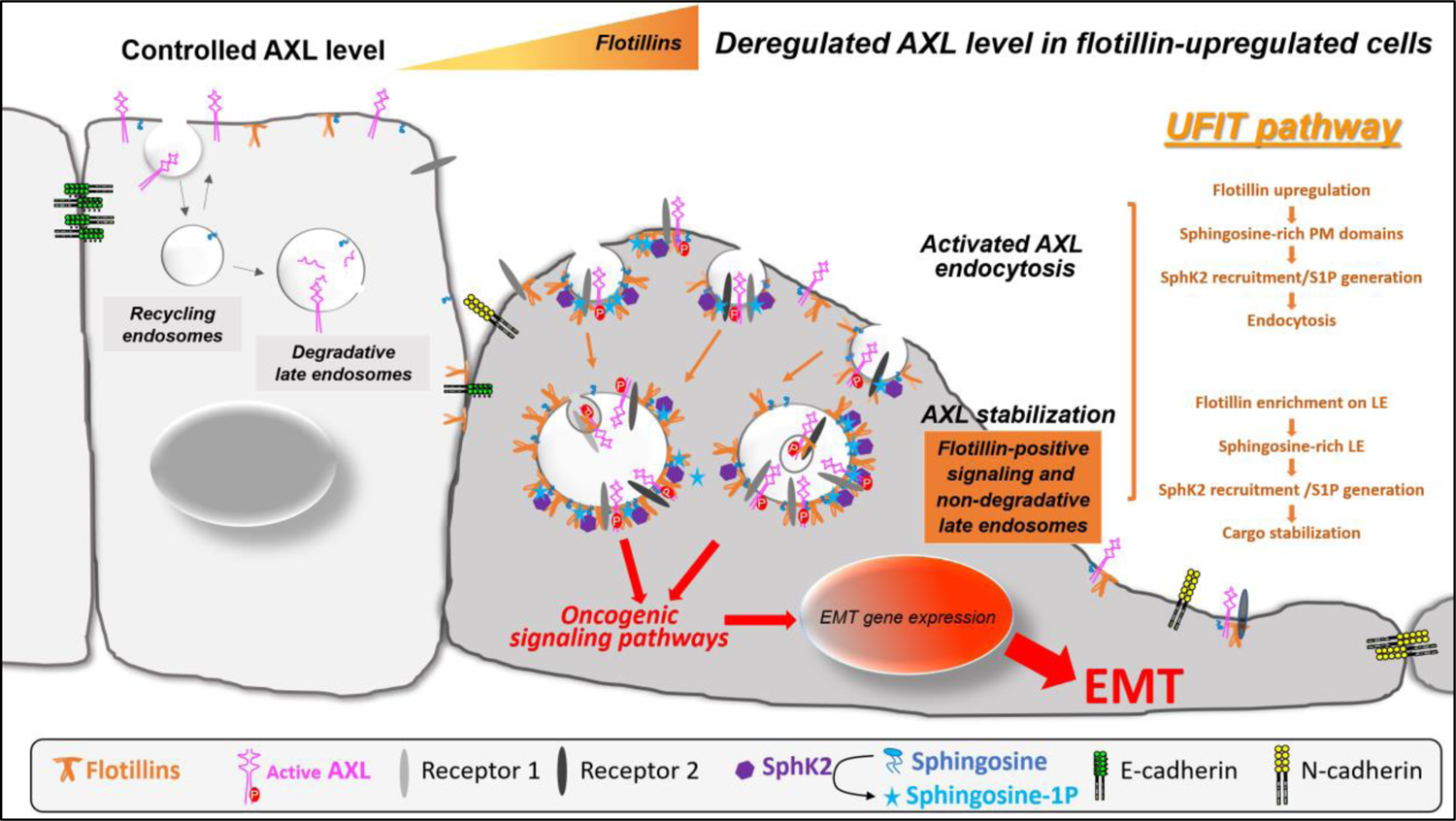
The UFIT pathway promotes AXL stabilization and EMT. Co-upregulation of flotillin 1 and 2 favors their oligomerization and the formation of flotillin-rich microdomains at the plasma membrane that initiate the UFIT pathway. At these membrane sites, signaling receptors, including the tyrosine kinase receptor AXL, are trapped. As sphingosine molecules bind to flotillins, these lipids are concentrated in flotillin-rich microdomains. This favors the recruitment of SphK2 that phosphorylates sphingosine and promotes sphingosine 1-phosphate production. Then, sphingosine 1-phosphate favors the endocytic process at the PM and most probably influences the function of late endosomes. Through the UFIT pathway AXL is delivered to flotillin-positive late endosomes, leading to its stabilization. Consequently, these flotillin-positive late endosomes favor the activation of oncogenic signaling pathways that promote EMT.

Flotillins are upregulated in many carcinoma types and also in sarcoma, and their upregulation is predictive of aggressive tumor behavior (reviewed in ^10^). Flotillin co-upregulation is necessary and sufficient to promote breast, nasopharyngeal, and rhabdomyosarcoma cancer cell invasion ^13, 18, 40^, and their specific role in metastasis formation was demonstrated in flotillin 2 knock-out mice ^14^. EMT is a crucial step in tumor initiation and cancer cell dissemination, but flotillin contribution to this process remained to be addressed ^12^. Previous studies were performed in cancer cell lines with mutations in both signaling and structural genes that promote EMT induction and invasion. This did not allow determining the exact contribution of flotillin upregulation to this crucial process and the molecular mechanisms specifically deregulated. To overcome this problem, we co-upregulated flotillins 1 and 2 (same level as observed in tumors) in immortalized non-tumoral mammary cell lines (MCF10A and NMuMG) that physiologically display low flotillin expression level. In these cells, flotillin upregulation was sufficient to promote EMT. In agreement, we found that in these models, key signaling pathways involved in EMT induction were activated. In MCFA10AF1F2 cells, the RTK AXL appeared as a key upstream actor. However, other receptors might also be co-deregulated on the basis of the results from our phospho-RTK array experiments (fig. S2H, I) and the presence in flotillin positive-endosomes of EphA4 and TGF-β-R (fig. S6), also implicated in mammary tumor development ^5, 41–43^. Flotillin upregulation in MCF10A cells led also to the activation of the Ras, ERK/MAPK, PI3-K/AKT, STAT3 and Smad3 pathways, but not NF-kB and WNT signaling cascades. Following flotillin upregulation, Ras and PI3K and the downstream signaling pathways, which are frequently activated in several cancers ^44^, were activated in MCF10A cells that do not harbor mutations in these cancer driver genes ^45^. Moreover, flotillin upregulation mimics oncogenic Ras activation that in MCF10A mammary cells has been correlated with ΔNp63α downregulation, PI3-K/AKT activation and ZEB1/2-mediated EMT^25^. These findings show that flotillin upregulation mimics the effects of cancer driver gene mutations (i.e. the activation of their downstream oncogenic signaling pathways) and acts as an EMT driver. Thus, quantification of flotillin expression level in tumors, even in the absence of cancer driver gene mutations, may help in therapeutic decision-making. Moreover, as flotillin upregulation leads to the activation of several membrane receptors and oncogenic signaling pathways, limiting their function (e.g. by inhibiting their oligomerization) could allow the simultaneously targeting of several drivers of tumor cell invasion.

We demonstrated that flotillin upregulation deregulates the trafficking of plasma membrane receptors, and identified them as new cargos of the UFIT pathway. This endocytic pathway occurs in the context of flotillin upregulation, and the proteins located in the flotillin-containing PM microdomains, which could be different in function of the cell type, are endocytosed and delivered to flotillin-positive non-degradative late endosomes ^18^. In addition to their weak degradative activity ^18^, our data suggest that these flotillin-positive vesicles have characteristics of signaling endosomes because they contained activated RTKs (fig. 4 and S6), and also the cancer driver K-Ras4B, associated with increased Ras activity upon flotillin upregulation (fig. S6G and 2B). It was previously reported that K-Ras and activated ERK1/2 are specifically recruited to Rab7-positive endosomes ^46^. Endosomes act as signaling platforms to facilitate the multiplexing of signaling molecules that in turn can affect the endosomal system by modifying the number, size and position of endosomes, and ultimately the trafficking of such signaling molecules ^6, 47^. In this context, the UFIT pathway is a key player to connect deregulation of endocytosis and signaling in cancer cells.

This is illustrated here for AXL, a cargo of the UFIT pathway, that is crucial to sustain EMT-promoting signaling upon flotillin upregulation in mammary cells. AXL is broadly expressed in a variety of solid cancers, particularly breast carcinoma, where its level is associated with high risk of metastases and poor patient prognosis ^5, 48^. AXL expression is correlated with the acquisition of a mesenchymal cell phenotype, invasive properties, and resistance to treatments. AXL is an attractive therapeutic target, and several AXL inhibitors are currently in preclinical and clinical development ^4, 49, 50^. Although AXL overexpression in tumors is attributed to post-transcriptional regulation, the underlying mechanisms are totally unknown. When overexpressed, AXL activation is ligand-independent and is the result of its oligomerization and/or association with other RTKs ^51^. Using an optogenetic system to force flotillin oligomerization upon illumination we demonstrated that AXL is trapped in light-induced flotillin clusters at the PM. Flotillin/AXL clusters are co-endocytosed, and the UFIT pathway allows AXL delivery to flotillin-positive endosomes and AXL stabilization. Thus, flotillin upregulation emerges as a key process in AXL post-transcriptional regulation. These data identify a new control mechanism in the ligand-independent regulation of AXL. Previous studies identified ligand-independent AXL stabilization at the cell surface through its association with the HER2 receptor ^51^, ligand-dependent AXL sequestration in cholesterol-enriched PM domains, or internalization ^52, 53^. To our knowledge this is the first report to describe AXL stabilization following its endocytosis. Moreover, this flotillin/AXL link is also observed in invasive breast tumor cells. Indeed, whereas AXL is very weakly expressed in non-tumoral breast lesions, its expression is broadly detected in all malignant breast tumor subtypes^51^. AXL expression appears correlated with that of flotillin 1, especially in the luminal subtype (fig. S4).

Our discoveries open a new interesting perspective for cancer therapy. Indeed, as AXL participates in the crosstalk with other RTKs (EGFR, Met,…), its inhibition could be more effective than just EGFR inhibition because it might restore sensitivity to multiple therapies ^28, 54^. Anti-AXL antibodies gave promising results in preclinical models of pancreatic and breast tumors ^49, 50, 55^. Endocytosis is dysregulated in human tumors and its inhibition increases the availability of therapeutic monoclonal antibody targets and promotes antibody-dependent cellular cytotoxicity ^56, 57^. Inhibition of flotillin function might have a strong effect on the efficacy of anti-AXL and also anti-RTK antibodies (e.g. the anti-EGFR antibody cetuximab) by increasing the membrane receptor availability at the cell surface, particularly in flotillin-positive tumors.

Flotillin upregulation-induced AXL stabilization requires the activity of SphK2, a lipid enzyme that maintains the cellular sphingolipid homeostasis, and a recognized target for cancer therapies ^58, 59^. SphK2 colocalizes with flotillin-positive domains at the PM and is enriched in flotillin-positive endosomes, in agreement with previous findings showing SphK2 localization in multivesicular endosomes in K562 leukemia cells ^60^. Flotillin binding to sphingosine and ectopic flotillin overexpression allow the formation of sphingosine domains that could favor SphK2 recruitment ^34^. We found that SphK2 inhibition or knock down, in cells with upregulated flotillins, strongly reduced AXL expression level and decreased cell invasion (fig. S7G). This is the consequence of the inhibition of upregulated flotillin-induced AXL endocytosis (fig. 6E-F). Accordingly, SphK2 inhibition did not affect AXL level in control MCF10A or in MDA-MB-231 invasive breast cells in which flotillins were knocked down. In these cells, AXL accumulated at the PM, suggesting that the UFIT pathway targets AXL to flotillin-positive endosomes to promote its escape from degradation. AXL can be degraded either at the PM or in the cytoplasm, through several mechanisms involving either metalloproteases or gamma-secretases, or the proteasome and the autophagic pathway ^61–63^. From which of these mechanisms the UFIT pathway protects AXL from degradation remains to be determined.

SphKs catalyze S1P production from sphingosine and there is a strong causal association between overactive SphK, S1P generation and cancer ^64^. SphK2 is involved in MDA-MB-453 cell migration toward epidermal growth factor ^65^ and SphK2 increase promotes oncogenesis. Moreover, results from a phase 1 clinical trial supported the use of SphK2 inhibitors in cancer treatment ^58, 59^. Showing that SphK2 inhibition impacts on AXL will help to understand the anti-tumoral function of these inhibitors and the role of SphK2 during tumor development, which is unknown. The UFIT pathway does not influence SphK2 expression level (fig. S7E), but affects its recruitment to cell membranes, at flotillin-rich endocytic sites (fig. 5D), and favors its recruitment to flotillin-positive endosomes (fig. 5E). In agreement, modifications of SphK2 subcellular localization are linked to tumorigenesis ^39, 66^. Sphingolipids are ubiquitous membrane components that are metabolized to form signaling molecules associated with cellular activities. Additional studies are required to understand how these sphingolipid modifications participate in endocytic events at the plasma membrane and how they influences endosome fusion, motion, and/or function. For example, whereas flotillin 1 directly interacts with Raf, MEK1/2 and ERK1/2 ^22^, flotillin-induced lipid modifications at late endosomes might indirectly regulate protein recruitment and/or activation. For instance, S1P allows the activation of five close G-protein coupled receptors (GPCRs), after its release in the intercellular space or directly through the outer leaflet of the plasma membrane ^67^. This leads to the subsequent induction of many downstream signaling pathways. Also, ceramide and S1P produced locally at late endosomes, again through GPCR activation, might participate in cargo sorting from the limiting late endosome membrane into their lumen ^68^.

In conclusion, this study adds new evidences for a crucial gain-of-function role of upregulated flotillins during EMT. We found that the UFIT pathway, by perturbing sphingolipid metabolism, allows the endocytosis of several PM receptors, particularly AXL, a major RTK during oncogenesis. AXL targeting to flotillin-positive late endosomes allows its stabilization and signaling during EMT.

## Methods

### Cell lines

All cell lines were authenticated, tested for contamination and cultured in a 37 °C incubator with 5% CO2. The MCF10A mammary epithelial cell line (American Type Culture Collection, ATCC CRL-10317) and the derived MCF10AmCh and MCF10AF1F2 cell lines were grown in DMEM/HAM F-12 medium supplemented with 5% horse serum (Gibco, Thermo), 10µg/ml insulin, 20ng/ml EGF, 0.5µg/ml hydrocortisone and 100ng/ml cholera toxin. The NMuMG mouse mammary epithelial cell line (ATCC CRL-1636) and the derived NMuMGmCh and NMuMG-F1F2 cell lines were grown in DMEM supplemented with 10% fetal bovine serum (FBS) (S1810-100; Biowest), 100 units/mL penicillin, 100 μg/mL streptomycin, 10 μg/mL insulin (I9278; Sigma–Aldrich). The human breast adenocarcinoma cell line MDA-MB-231 (ATCC HTB-26) and the MDA-MB-231shLuci and MDA-MB-231 shFlot2 derived cell lines (described in ^18^) were grown in DMEM with 10% FBS (Biowest, Eurobio). Retrovirus production in Phoenix cells (G. Nolan, Stanford, USA), infection and selection were performed as described ^69^. Cells were grown continuously in 1 µg/ml puromycin and 200µg/ml hygromycin.

The MCF10AF1F2 and NMuMGF1F2 cell lines that stably overexpress flotillin 1 and 2 were generated by double retroviral infection of two vectors encoding the chimeric proteins flotillin-2-mCherry and flotillin-1-HA (described in ^18^), followed by selection for resistance against puromycin and hygromycin. Cells were then sorted by flow cytometry according to the mCherry fluorescence signal.

### Plasmids and transfections

peGFP-N1-AXL was generated by subcloning the human AXL sequence (NM_021913.4) amplified by PCR using the primers 5’AAGCTTATGGCGTGGCGGTGCCCCAGG 3’ and 5’ GAATTCCGGCACCATCCTCCTGCCCTG 3’ between the *Hind*III and *Eco*RI sites in peGFP-N1. pHalo-N1-AXL was obtained by replacing GFP by the Halo-sequence in peGFP-N1-AXL between *Hind*III and *EcoR*I. To generate the pBabe-Flot2-CIBN-mCherry construct, the sequence encoding the CIBN domain was amplified by PCR using the primers 5’ GAATTCTAATGAATGGAGCTATAGGAGG 3’ and 5’ CGGATTATATTCATGTACCGGTCACACA 3’, and inserted, using the *Eco*RI and *Age*I sites, between flotillin 2 and mCherry in the pBabe-Flot2-mCherry plasmid ^18^. The peGFP-ERK2 construct was a gift from Dr H Farhan (University of Oslo, Norway). The CD63-GFP plasmid was a gift from Dr A Weaver (Vanderbilt University, Nashville, USA) ^70^. The Rab7a-GFP plasmid was described in ^18^. The pCMV3-N-Myc-human AXL (HG10279-NM) plasmid was purchased from Sino Biological. The peGFP-C1-KRas4A, peGFP-C3-KRas4B, peGFP-C3-HRas and peGFP-C3-NRas plasmids were gifts from Dr M Philips (NYU Cancer Institute, New York, USA). The phCMV3-Flot1-HA plasmid was a gift from Dr V Niggli (University of Bern, Switzerland) and pmCherry-N1-Flot2 was previously described ^18^. The plasmids encoding the different TAp63 α,β,γ and ΔNp63 α,β,γ isoforms were gifts from Dr C Caron de Fromentel (INSERM UMR590, Lyon, France) ^71^, the PCS2+chickenEphA4-GFP plasmid was a gift from Prof F Fagotto (CRBM, Montpellier, France) ^72^. The TGFβRI-GFP and TGFβRII-Flag constructs were gifts from Dr R Derynck (UCSF, San Francisco, USA) ^73^. The SphK1-GFP and SphK2-GFP constructs were gifts from Prof P de Camilli (Yale School of Medecine, New Haven, USA) ^35^. The SphK2-Halo plasmid was obtained by replacing GFP with the Halo-sequence between *Eco*RI and *Not*I. The CRY2-mCitrin plasmid was a gift from Prof W Do Heo (KAIST, Dajeon, South Korea), previously described in ^32^. The TfR(CD71)-GFP vector was purchased from Addgene. Cells were transfected with the indicated constructs using the JetPEI or JetPrime Transfection Reagents (Polyplus Transfection, Illkirch, France).

### Antibodies and reagents

Mouse antibodies used were against: actin (A5441, Sigma), flotillin 1 and flotillin 2 (BD Biosciences), E-cadherin (used for MCF10A cells, Life-technologies), E-cadherin (used for NMuMG cells, BD Biosciences), LAMP1 (used for MCF10A cells, BD Biosciences), Smad3 (ThermoFisher), vimentin and α-tubulin (Sigma), phosphorylated ERK1/2 at T202/Y204 and AKT (CST), pan-Ras (Millipore), phosphorylated Y (4G10), EphA4 (gift from Dr Greenberg, Harvard, Boston, USA) ^72^, and STAT3 (CST, #9139S). Rabbit antibodies were against flotillin 1 (F1180, Sigma), flotillin 2 (3436, CST), AXL (C89E7, CST), phosphorylated AXL at Y702 (D12B2, CST), ERK1/2 (CST), phosphorylated AKT at S473, ZEB1 (Santa Cruz), ZO-1 (Zymed), phosphorylated Smad3 at S423/425, p63, and phosphorylated EphA4 at Y588/596 (Abcam), phosphorylated STAT3 at Y705 (#9131S, CST), p65-NFκB (Santa Cruz), β-catenin (Sigma, 2206), HA-tag (Invitrogen), Flag-tag (Sigma), Myc-tag (Upstate), SPK2 (Clone D2V3G, CST). The humanized anti-AXL antibody D4 (https://patents.google.com/patent/EP3229836A1/en?oq=EP3229836A1) and the mouse anti-AXL antibody D9 ^49^ were gifts from Dr B Robert (ICRM, Montpellier, France). Alexa-488, −546, −633-conjugated secondary antibodies were from Thermo Scientific. Alexa-Fluor-488-phalloidin was from Invitrogen and Hoechst (0.1 mg/ml) was from Sigma-Aldrich. Lipopolysaccharide (LPS) (Sigma, L5668) was used at 1µg/ml for 1h. Halo-tagged proteins were labeled by incubating cells with 200nM HaloTag Ligand conjugated to Janelia Fluor 646 (Promega) for 30 min before imaging. Opaganib (ABC294640) (used at 50 µM up to 10 hours) and PF543 (used at 10 µM up to 24h) were from Selleckchem. Cycloheximide (used at 100 µg/ml for 6h) was from Sigma-Aldrich.

### RNA interference

The pSIREN-RetroQ-Flot2-shRNA (targeting both human and murine Flot2) and pSIREN-RetroQ-Luciferase shRNA vectors were previously described ^74^. Luciferase shRNA (shLuci) was used as control ^69^. For RNA-mediated interference experiments, cells were transfected using Interferin (Polyplus Transfection) with siRNA oligonucleotide duplexes (Eurogentec, Belgium). The sequences of the siRNA duplexes were: for human ZEB1, 5’ GGUAGAUGGUAAUGUAAUA 3’ matching the human ZEB1 sequence (NM_001128128.2) ^75^; for human ZEB2 5’ GCAUGUAUGCAUGUGACUU 3’ matching the human ZEB2 sequence (NM_014795.3) ^76^; for human AXL 5’ CGAAAUCCUCUAUGUCAACAU 3’ matching the human AXL sequence (NM_021913.4) ^77^; control siRNA (Eurogentec, SR-CL000-005); for human SphK2 5’ GCUGGGCUGUCCUUCAACCU 3’, matching the human SphK2 sequence (NM_020126.4) ^65^.

### Gel electrophoresis and immunoblotting

Protein concentrations in whole-cell lysates were determined with the BCA protein assay kit (Pierce). Proteins from whole cell lysates (20–60 µg) were separated on 8-10-12% polyacrylamide gels and transferred to Immobilon-FL membranes (Millipore). Primary and secondary antibodies were diluted in blocking buffer (Millipore) containing 0.1% Tween 20. Detection and analysis were performed using the Odyssey Infrared imaging system (LI-COR Biosciences). Immunoblots were quantified by densitometry using Odyssey V3.0 and Fiji.

### Immunoprecipitation

The rabbit polyclonal anti-AXL (CST, 8661) antibody (1µg) was incubated with Dynabeads Protein G (25 µl) at room temperature for 1h. Antibody coated-beads were then incubated with 1 mg of proteins from post-nuclear supernatants of the indicated cells at 4°C for 2 h, washed and loaded on SDS-PAGE. Immunoprecipitates were analyzed by western blotting.

### Phospho-RTK and phospho-kinase arrays

The Proteome Profiler Human Phospho-RTK array kit and human phospho-kinase antibody array kit (R&D Systems) were used according to the manufacturer’s instructions. Both array types were incubated with 200 µg of proteins from post-nuclear supernatants isolated from each tested cell line maintained in complete culture medium for 48h after seeding.

### Measurement of Ras activity

Cells were lysed in lysis buffer A (50 mM Tris pH 7.5, 10 mM MgCl_2_, 0.3 M NaCl, 2% (v/v) IGEPAL). Lysates were then incubated with 30 µg of Ras-GTP Binding Domain (RBD of human c-Raf kinase) fused with GST associated with beads (#RF02, Cytoskeleton) at 4°C for 1h. Prior to incubation with beads, 1/20 of each cell lysate was collected to be used as “input”. Precipitates and inputs were revealed by western blotting using an anti-pan-Ras antibody.

### AXL uptake assays

Cells were seeded in 12 mm sterile poly-lysine-coated round glass coverslips and grown to 60-70% confluence. After 15 min incubation at 4°C in serum-free medium, cells were incubated at 4°C in serum-free medium containing 250 μg/ml of D4 anti-AXL humanized antibody for 45 min. Alternatively, cells transfected with the plasmid encoding N-terminally Myc-tagged AXL (N-Myc-AXL) were incubated with the anti-Myc (9E10) antibody. Cells were extensively washed with cold serum-free medium to remove unbound antibodies and surface-bound antibodies were internalized by incubating cells at 37°C in serum-free medium for the indicated time points. Cells were then fixed and permeabilized, and the internalized primary antibody molecules were detected using the appropriate fluorescent secondary antibody (FITC-conjugated anti-human IgG for cells incubated with the D4 antibody, and Alexa488-conjugated anti-mouse IgG for cells that express N-Myc-AXL). Samples were then processed as described in the Immunofluorescence and image acquisition chapter.

### Surface biotinylation assays/internalization experiments

Cells were seeded and after 24 hours were washed with chilled PBS, incubated with biotin NHS (0.2mg/ml in PBS pH 7.9) at 4°C for 45 min, and shifted to 37°C for the indicated times. Cells were then incubated with Mesna buffer at 4°C for 15 min to remove the remaining cell-surface biotin, followed by a 10min incubation at 4°C with iodoacetamide to stop the action of Mesna. Cells were then scraped in lysis buffer (10 mM Pipes pH 7, 100 mM NaCl, 300 mM sucrose, 3 mM MgCl_2_, 0.5% (v/v) IGEPAL CA-640, 1 mM EDTA, 1 mM Na_3_VO_4_, protease inhibitor cocktail) and the post-nuclear supernatants were harvested after centrifugation (10.000rpm 4°C for 10 min). One cell plate was lysed right after biotinylation and before incubation with Mesna buffer and used to estimate the amount of total labeled surface proteins. An equal protein amount for each time point was incubated with streptavidin-coated agarose beads (at 4°C for 45 min). After two washes in PBS, beads were precipitated and resuspended in Laemmli buffer. Samples were separated by SDS-PAGE followed by western blotting with antibodies against AXL and CD71.

Results from the experiments performed using MCF10AmCh and MCF10AF1F2 cells were not expressed as the percentage of total surface labelled AXL at T0min because AXL expression level in MCF10AF1F2 cells was much strongly than in MCF10AmCh cells (see fig. 3A and 4G). In these conditions, expressing the data classically as the percentage of the maximum level of surface labeled AXL at T0min would have led to a strong underestimation of any difference in the internalization rate. Thus, to estimate correctly the internalization rate, results at each time point were expressed as the percentage of the maximum level of internalized protein (reached at 8 min for MCF10AF1F2 cells and at 12 min for MCF10A cells). Because the AXL expression level remained unchanged between untreated MDA-MB-231shLuci cells or incubated with opaganib for 4 hours and MDA-MB-231shFlot2 cells, and between untreated MCF10AF1F2 cells and those incubated with opaganib for 4 hours, results at each time point were expressed as the percentage of the maximum level of surface labeled AXL or CD71 at T0min.

### Immunofluorescence, image acquisition and analysis

Cells were fixed in 3.2% paraformaldehyde for 10 min, followed by a 2min permeabilization with 0.1% Triton X-100 or by a 10min permeabilization with 0.05% saponin, and a 15min saturation in the presence of 2% BSA. Cells were then incubated with primary and secondary antibodies. For CD63, LAMP1, and flotillin immunostaining, cells were fixed and permeabilized with acetone/methanol (v/v, 1/1) at −20°C for 2 min. Images were taken with a confocal SP5-SMD microscope (Leica) with 40×/1.3 or 63×/1.4 oil HCX PL APO CS objectives (Leica), and captured with a hybrid detector (Leica HyD) controlled using the C software. In some cases, stacks of confocal images with a step size of 0.3μm were processed with Imaris (Bitplane, Zurich, Switzerland) for visualization and volume rendering. Images were processed using Adobe Photoshop and assembled using Adobe Illustrator. For the colocalization measurements of flotillin1/2-mCherry with AXL-GFP, CD71-GFP, SphK1/2-GFP in vesicles, images were analyzed with the Fiji software and using the Icy software (http://icy.bioimageanalysis.org); the detailed protocol will be provided upon request.

### Quantification of AXL surface staining

Fixed non-permeabilized cells were immunostained using the monoclonal D4 antibody against AXL extracellular domain and an Alexa 488-conjugated anti-human secondary antibody. Images were acquired by confocal microscopy under the same illuminated conditions. Using the Fiji software, the outline of each cell was defined and the mean fluorescence intensity (taking into account the cell surface) was quantified in the area defined by the outline.

### Optogenetic and TIRF microscopy

MCF10A cells were co-transfected with vectors encoding CRY2-mCitrin, flotillin 2-CIBN-mCherry and AXL-Halo labeled with the Halo-Tag-Ligand-Janelia Fluor 646. TIRF images were acquired using an inverted Nikon microscope and a 100×/1.49 oil objective controlled using the Metamorph software (1 image /s). First, Flot2-CIBN-mCherry and AXL-Halo were imaged for 10 seconds before 488nm illumination. Then, cells were globally illuminated by 22 pulses (1 pulse/s, 100 ms time length for each pulse) of 488nm-light and CRY2-mCitrin, Flot2-CIBN-mCherry and AXL-Halo signals were simultaneously imaged for 3 min.

### Quantification of SphK2-GFP delocalization from flotillin-positive late endosomes

MCF10AF1F2 cells that express SphK2-GFP were imaged by spinning disk confocal microscopy (1 image/15 min). The Fiji software and images acquired at 0 and 90 min after opaganib (50µM) addition in the culture medium were used to define same-size individual regions of interest (ROI) surrounding one flotillin 2-mCherry-positive vesicle. For each ROI, the mean intensity corresponding to the mCherry and the GFP signals was quantified. To take into account the decrease in signal intensity due to photobleaching, quantifications were done also on three ROI of the same size, but outside flotillin 2-mCherry-positive vesicles. The mean of the obtained values was considered as background and subtracted from the GFP and mCherry signals measured in the ROI surrounding vesicles. Then, the GFP/mCherry signal ratio was individually calculated for each ROI surrounding a vesicle on images acquired at 0 and 90 min after opaganib addition.

### Time-lapse imaging experiments

Time-lapse imaging experiments were performed using an inverted Spinning disc Nikon Ti Andor CSU-X1 microscope equipped with a focus Fluor 100 × objective (NA 0.55). Time-series images were acquired with a EMCCD iXon897 Andor camera controlled by the Andor iQ3 software. Time lapse was started 24/48h after transfection. Time series of the captured images were saved as .tiff files that were compiled into .avi movies using the Fiji software.

### Scanning electron microscopy

PBS-washed cells were fixed with 2.5% glutaraldehyde in PHEM buffer, pH 7.2 at room temperature for 1 hour, and then washed in PHEM buffer. Fixed cells were dehydrated using a graded ethanol series (30-100%), followed by 10 min in graded ethanol-hexamethyldisilazane and 10 min in hexamethyldisilazane. Then, samples were sputter coated with a 10nm thick gold film and examined using a scanning electron microscope (Hitachi S4000, at CoMET, MRI-RIO Imaging, Biocampus, INM Montpellier France) and a lens detector with an acceleration voltage of 10KV at calibrated magnifications.

### RNA isolation and sequencing

Total RNA was prepared from MCF10AmCh and MCF10AF1F2 cells using the RNeasy Mini Kit (Qiagen) according to the manufacturer’s instructions. RNA quantity and quality were determined with an Agilent RNA 600 Nano Bioanalyser (Agilent Technologies), and total RNA was concentrated to 100 ng/µl. Pure RNA preparations without DNA or protein contamination were used for sequencing. Libraries, processed by Fasteris SA (Geneva, Switzerland), were prepared with the TruSeq Stranded mRNA Library Prep Kit from Illumina. Fragments were sequenced on an Illumina Hi-Seq 4000 sequencer according to the manufacturer’s instructions.

### Differential analysis of transcriptomic data

The differential gene expression analysis was carried out using three replicates per condition to compare the transcript levels between MCF10AmCh and MCF10AF1F2 cells. Two analysis methods were used: DESeq (http://bioconductor. org/packages/release/bioc/html/DESeq.html) and Tuxedo (http://cufflinks.cbcb.umd.edu/).

Genes with a *p_adj_*<0.05 were assigned as differentially expressed. The Deseq2 and Tuxedo results were compared. Only genes with an expression fold change >2.8 were considered. Using this criterion 802 up- and down-regulated genes were identified.

### Gene functional classification: GO term enrichment analysis and Gene Set Enrichment ***Analysis*.**

The PANTHER (protein annotation through evolutionary relationship, version 13.1) classification system (http://www.pantherdb.org/) was used to perform a Gene-Ontology (GO) enrichment. Enriched terms were considered statistically significant when *padj* was <0.05, and a minimum of five genes were grouped for each significant term. GO terms were categorized in three major functional groups: biological process, molecular function, and cellular component. A gene set enrichment analysis (GSEA) was performed with the GSEA software v4.1.0. The preranked method was used and the number of permutations was set to 1000. Gene

### Reverse Transcription-quantitative real-time PCR (RT-qPCR)

RNA was extracted from the indicated cell lines as described above. Then, mRNA samples were reverse transcribed to cDNA using SuperScript II Reverse Transcriptase (Thermo Fisher Scientific) and qPCR was performed using an LC480 apparatus (Roche) with the SYBR Green PCR Master Mix (Applied Biosystems). The primer sequences were: for human CDH1 forward 5’ TCATGAGTGTCCCCCGGTAT 3’ and reverse 5’ CAGCCGCTTTCAGATTTTCAT 3’; for human CDH2, forward 5’ GGCAGAAGAGAGACTGGGTC 3’ and reverse 5’ GAGGCTGGTCAGCTCCTGGC 3’; for human vimentin, Duplex QT00095795 (Quiagen); for human ZEB1, forward 5’ GCACCTGAAGAGGACCAGAG 3’ and reverse 5’ TGCATCTGGTGTTCCATTTT 3’; for human ZEB2, forward 5’ GGGAGAATTGCTTGATGGAGC 3’ and reverse 5’ TCTCGCCCGAGTGAAGCCTT 3’; for human MRPL19 (internal control), forward 5’ GGGATTTGCATTCAGAGATCAG 3’ and reverse 5’ GGAAGGGCATCTCGTAAG 3’; for murine CDH1, forward 5’ GGTCTCTTGTCCCCACA 3’ and reverse 5’ CCTGACCCACACCAAAGTCT 3’; for murine CDH2, forward 5’ AGGGTGGACGTCATTGTAGC 3’ and reverse 5’ CTGTTGGGGTCTGTCAGGAT 3’; for murine vimentin, forward 5’ AATGCTTCTCTGGCACGTCT 3’ and reverse 5’ GCTCCTGGATCTCTTCATCG 3’; for murine ZEB1, forward 5’ GCTCAGCCAGGAACCCGCAG 3’ and reverse 5’ TGGGCACCCTCTGCCACACA 3’; for murine ZEB2, forward 5’ ATGGCAACACATGGGTTTAGTGGC 3’ and reverse 5’ ATTGGACTCTGAGCAGATGGGTGT 3’; for murine MmRPL32 (internal control), forward 5’ TTAAGCGAAACTGGCGGAAAC 3’ and reverse 5’ TTGTTGCTCCCATAACCGATG 3’; for human AXL, forward 5’ GTGGGCAACCCAGGGAATATC 3’ and reverse 5’ GTACTGTCCCGTGTCGGAAAG 3’.

### RT-qPCR analysis of breast tumor samples

The mRNA levels of flotillin 1 and ΔNp63 in tumor samples from the cohort of patients with breast cancer were analyzed as described ^79^.

### Boyden chamber migration assay

Cells (5×10^4^) were seeded on the upper chamber of the Boyden chamber inserts (Corning FluoroBlok cell culture insert, 8 µm pores) in 250µl of DMEM/F12 with 1% horse serum. In the lower chamber, complete medium (750µl) was added, as chemoattractant. Opaganib (50 µM) was added in the upper and lower chambers. After 14h, cells at the bottom of the filter were fixed with 3.2% paraformaldehyde and stained with Hoechst. The nucleus of cells that had migrated through the pores was imaged with an EVOS microscope (Life technologies) using a 10x objective. Semi-automatic quantification was performed with Fiji. For each experiment, each condition was performed in triplicate.

### Patients and specimens

Patients (n=9) with invasive breast carcinoma were treated at the Cancer Research Institute, Tomsk NRMC (Tomsk, Russia). Frozen and Formalin-fixed paraffin-embedded (FFPE) tumor tissue specimens obtained during surgery were used for immunofluorescence analyses.

The histological type was defined according to the World Health Organization recommendations ^80^. Tumor grade was determined using the Bloom and Richardson grading system ^81^. The expression of estrogen receptor (ER) and progesterone receptor (PR) was scored using the HSCORE method ^82^. HER2 expression was analyzed by immunohistochemistry and calculated on a scale 0-3+, according to the ASCO/CAP guidelines ^83^. Ki-67 expression was calculated as the percentage of Ki-67-positive tumor cells relative to all tumor cells. Molecular subtypes were categorized on the basis of the primary tumor ER, PR, HER2, and Ki-67 status, according to the St Gallen recommendations ^83^. luminal A (ER^+^ and/or PR^+^, HER2^-^, and Ki-67 <20%), luminal B (ER^+^ and/or PR^+^, HER2^+/-^, and Ki-67 ≥20%), HER2^+^ (ER^-^ and PR^-^, HER2^+^), and triple-negative (ER^-^, PR^-^, HER2^-^). The study procedures were in accordance with the Helsinki Declaration (1964, amended in 1975 and 1983). This study was approved by the institutional review board; all patients signed an informed consent for voluntary participation.

### Immunofluorescence staining of patient tumor samples

Seven μm-thick FFPE tumor sections were deparaffinized, rehydrated, processed for heat-induced epitope retrieval with EDTA buffer (pH 8.0) using PT Link (Dako, Denmark), and blocked with 3% bovine serum albumin (Amresco, USA) in PBS. Expression of flotillin 1 (F1180, 1:50, Sigma) and AXL (C89E7, CST) was assessed using the BondRXm immunostainer (Leica, Germany). TSA visualization was performed with the Opal 520 and Opal 690 (Opal seven-color IHC Kit (NEL797B001KT; PerkinElmer), nuclei were stained with DAPI and samples were mounted in Fluorescence Mounting Medium (Dako, Denmark). Samples were analyzed using the Vectra 3.0 Automated Quantitative Pathology Imaging System (PerkinElmer, USA).

### Statistical analysis

All statistical analyses were performed using SPSS version 22, and graphs were generated with Prism GraphPad version 5. All data were first tested for normality using the Kolmogorov– Smirnov normality test. For experiments with n >30, the Student’s *t* test was used to identify significant differences between experimental conditions. For experiments with n <30, the non-parametric Mann-Whitney U test was used. The n value and number of independent experiments are listed in the figure legends. At least three independent experiments were performed. In the figures, the differences are significant only when indicated.

## Supporting information

Genest Video 1

Genest Video 2

Genest Video 3

Genest Video 4

Genest Video 5

Genest Video 6

Genest Video 7

Genest Video 8

Genest Video 9

Genest Tables S1 and S2

Genest Figure S1

Genest Figure S2

Genest Figure S3

Genest Figure S4

Genest Figure S5

Genest Figure S6

Genest Figure S7

## Acknowledgements

We thank Peggy Raynaud, Dominique Helmlinger Dylane Detilleux and Christelle Dantec for help in the transcriptomic analysis. We thank Dimitris Xirodimas for advices regarding experiment investigating AXL stability. We acknowledge Virginie Georget, Chantal Cazevieille, Orestis Faklaris, Sylvain de Rossi and Volker Baeker from the Montpellier Imaging Facility MRI, member of the national infrastructure France-BioImaging infrastructure. This work was supported by the “Fondation pour la Recherche Médicale” (DEQ20161136700) and the “Fondation Arc pout la recherche sur le cancer” (ARCPJA32020060002078). Montpellier Imaging Facility is supported by the French National Research Agency (ANR-10-INBS-04, “Investments for the future”. Collection of breast tumour samples was supported by the Russian Science Foundation (grant 19-75-30016). C.G.R. was supported by Institut National de la Santé et de la Recherche Médicale, M.G by the french ministry for scientific research and S.B by the University of Montpellier.

## Author contributions

C.G-R. and S.B. designed the research. M.G., F.C., D.P., P.G., H.M., A.S., C.G.R and S.B. designed and performed the experiments. B.R. generated the anti-AXL D4 and D9 antibodies. S.V. and I.B. performed the transcriptomic analysis on flotillin 1 and ΔNp63 mRNA expression from the cohort of 527 patients with breast cancer. L.A.T. performed the immunofluorescence analysis of flotillin 1 and AXL in invasive breast tumor tissue sections. M.G., S.B. and C.G-R created the figures and videos. C.G-R. and S.B. wrote the paper. All contributors discussed the results and commented on the manuscript.

## Competing interests

The authors declare no competing financial interests

## Materials &correspondence

Correspondance and material should be addressed to Stéphane Bodin and Cécile Gauthier-Rouvière

## Supplementary information

### Supplementary information on patient tumor samples

Clinicopathological characteristics of the patients who gave the breast cancer samples used for this expression analysis: *Triple negative subtype:* Patient #1: 65-year-old woman, invasive carcinoma (left side), T2N1M0, grade 3, ER−, PR−, Her2−, Ki-67-positive cells (50%), triple-negative. Lymph node-positive (1 metastasis from 6 lymph nodes). Five courses of neoadjuvant chemotherapy. Patient #2: 55-year-old woman, invasive carcinoma (left side), T1N0M1, multifocal, grade 2, ER^-^, PR^-^, Her2^-^, Ki-67-positive cells (31%), triple-negative. Lymph node negative (0 metastasis in 7 lymph nodes). Six courses of neoadjuvant chemotherapy. Patient #3: 55-year-old woman, invasive carcinoma (left side), T2N1M0, grade 2, ER-, PR-, Her2^-^, Ki-67-positive cells (31%).

Lymph node negative (0 metastasis in 7 lymph nodes). Six courses of neoadjuvant chemotherapy. Patient #4: 55-year-old woman, invasive carcinoma (left side), T1N0M0, grade 3, ER−, PR−, Her2−, Ki-67-positive cells (52%), triple-negative. Lymph node-negative (0 metastasis from 11 lymph nodes). No neoadjuvant chemotherapy. Patient #5: 48-year-old woman, invasive carcinoma (right side), T2N1M0, grade 3, ER−, PR−, Her2−, Ki-67-positive cells (37%), triple-negative. Lymph node-positive (2 metastases from 8 lymph nodes). Five courses of neoadjuvant chemotherapy. Patient #6: 48-year-old woman, invasive carcinoma (left side), T2N1M0, grade 3, ER−, PR−, Her2−, Ki-67-positive cells (80%), triple-negative. Lymph node-negative (0 metastasis from 10 lymph nodes). Six courses of neoadjuvant chemotherapy. Patient #7: 24-year-old woman, invasive carcinoma (left side), T4N0M0, grade 2, ER−, PR−, Her2−, Ki-67-positive cells (60%), triple-negative. Lymph node-negative (0 metastasis from 10 lymph nodes). Six courses of neoadjuvant chemotherapy. Patient #8: 59-year-old woman, invasive carcinoma (right side), T1N1M0, grade 2, ER−, PR−, Her2−, Ki-67-positive cells (61%), triple-negative. Lymph node-positive (1 metastasis from 8 lymph nodes). No neoadjuvant chemotherapy. Patient #9: 51-year-old woman, invasive carcinoma (right side), T2N1M0, grade 2, ER−, PR−, Her2−, Ki-67-positive cells (48%), triple-negative. Lymph node-positive (1 metastasis from 10 lymph nodes). Five courses of neoadjuvant chemotherapy.

Patient #10: 65-year-old woman, invasive carcinoma (left side), T1N1M0, grade 2, ER−, PR−, Her2−, Ki-67-positive cells (50%), triple-negative. Lymph node-positive (1 metastasis from 10 lymph nodes). No neoadjuvant chemotherapy. Patient #11: 60-year-old woman, invasive carcinoma (left side), T2N0M0, grade 2, ER−, PR−, Her2−, Ki-67-positive cells (45%), triple-negative. Lymph node-negative (0 metastasis from 8 lymph nodes). No neoadjuvant chemotherapy. Patient #12: 43-year-old woman, invasive carcinoma (left side), T2N0M0, grade 2, ER−, PR−, Her2−, Ki-67-positive cells (60%), triple-negative. Lymph node-negative (0 metastasis from 8 lymph nodes). No neoadjuvant chemotherapy. Patient #13: 54-year-old woman, invasive carcinoma (right side), T2N1M0, grade 3, ER−, PR−, Her2−, Ki-67-positive cells (60%), triple-negative. Lymph node-positive (1 metastasis from 11 lymph nodes). Six courses of neoadjuvant chemotherapy.

*Luminal subtype:* Patient #1: 49-year-old woman, invasive carcinoma (right side), T2N0M0, grade 2, ER+, PR+, Her2^-^, Ki-67-positive cells (10%), luminal. Lymph node negative (0 metastasis in 3 lymph nodes). No neoadjuvant chemotherapy. Patient #2: 66-year-old woman, invasive carcinoma (right side), T2N1M0, grade 2, ER+, PR+, Her2^-^, Ki-67-positive cells (35%), luminal. Lymph node positive (3 lymph nodes with mts from 10 lymph nodes). No neoadjuvant chemotherapy. Patient #3: 40-year-old woman, invasive carcinoma (right side), T2N1M0, grade 2, ER+, PR+, Her2^-^, Ki-67-positive cells (33%), luminal. Lymph node positive (2 lymph nodes with mts from 10 lymph nodes). No neoadjuvant chemotherapy. Patient #4: 46-year-old woman, invasive carcinoma (right side), T2N1M0, grade 2, ER+, PR-, Her2^-^, Ki-67-positive cells (12%), luminal. Lymph node-positive (5 lymph nodes with mts from 6 lymph nodes). No neoadjuvant chemotherapy. Patient #5: 57-year-old woman, invasive carcinoma (right side), T4N3M0, grade 2, ER+, PR+, Her2^-^, Ki-67-positive cells (17%), luminal. Lymph node-positive (2 lymph nodes with mts from 6 lymph nodes). Six courses of neoadjuvant chemotherapy. Patient #6: 32-year-old woman, invasive carcinoma (right side), T2N1M0, grade 2, ER+, PR+, Her2^-^, Ki-67-positive cells (10%), luminal. Lymph node-positive (6 lymph nodes with mts from 10 lymph nodes). No neoadjuvant chemotherapy. Patient #7: 60-year-old woman, invasive carcinoma (left side), T2N1M0, grade 2, ER+, PR+, Her2^-^, Ki-67-positive cells (30%), luminal. Lymph node-positive (2 lymph nodes with mts from 11 lymph nodes).

Six courses of neoadjuvant chemotherapy. Patient #8: 45-year-old woman, invasive carcinoma (right side), T2N1M0, grade 2, ER+, PR-, Her2^-^, Ki-67-positive cells (21%), luminal. Lymph node-negative (0 metastasis from 8 lymph nodes). No neoadjuvant chemotherapy. Patient #9: 36-year-old woman, invasive carcinoma (left side), T2N0M0, grade 2, ER+, PR+, Her2^-^, Ki-67-positive cells (28%), luminal. Lymph node-negative (0 metastasis from 6 lymph nodes). No neoadjuvant chemotherapy. Patient #10: 60-year-old woman, invasive carcinoma (left side), T2N0M0, grade 3, ER+, PR+, Her2^-^, Ki-67-positive cells (10%), luminal. Lymph node-negative (0 metastasis from 10 lymph nodes). No neoadjuvant chemotherapy. Patient #11: 55-year-old woman, invasive carcinoma (left side), T2N1M0, grade 3, ER+, PR+, Her2^-^, Ki-67-positive cells (25%), luminal. Lymph node-positive (1 lymph nodes with mts from 10 lymph nodes). No neoadjuvant chemotherapy. Patient #12: 56-year-old woman, invasive carcinoma (left side), T2N1M0, grade 2, ER+, PR+, Her2^-^, Ki-67-positive cells (10%), luminal. Lymph node-positive (2 lymph nodes with mts from 7 lymph nodes). No neoadjuvant chemotherapy.

Patient #13: 62-year-old woman, invasive carcinoma (right side), T1N0M0, grade 2, ER+, PR-, Her2^-^, Ki-67-positive cells (20%), luminal. Lymph node-negative (0 metastasis from 10 lymph nodes). Six courses of neoadjuvant chemotherapy. *HER2 subtype:* Patient #1: 47-year-old woman, invasive carcinoma (right side), multifocal, T4N2аM0, grade 3, ER-, PR-, Her2^+^,Ki-67-positive cells (65%). Lymph node positive (7 lymph nodes with mts from 16 lymph nodes). Six courses of neoadjuvant chemotherapy. Patient #2: 46-year-old woman, invasive carcinoma (right side), T2N0M0, grade 2, ER-, PR-, Her2^+^, Ki-67-positive cells (53%). Lymph node negative (0 metastasis in 7 lymph nodes). One course of neoadjuvant chemotherapy. Patient #3: 52-year-old woman, invasive carcinoma (left side), T2N1M0, grade 2, ER-, PR-, Her2^+^, Ki-67-positive cells (42%). Lymph node positive (6 lymph nodes with mts from 8 lymph nodes). No neoadjuvant chemotherapy. Distant metastasis in liver and lung.Patient #4: 58-year-old woman, invasive carcinoma (right side), T1N0M0, grade 2, ER-, PR-, Her2^+^, Ki-67-positive cells (35%). Lymph node-positive (6 lymph nodes with mts from 10 lymph nodes). Six courses of neoadjuvant chemotherapy. Patient #5: 37-year-old woman, invasive carcinoma (right side), T2N0M0, grade 2, ER-, PR-, Her2^+^, Ki-67-positive cells (50%). Lymph node-negative (0 lymph nodes with mts from 11 lymph nodes). Six courses of neoadjuvant chemotherapy. Patient #6: 56-year-old woman, invasive carcinoma (left side), T3N1M0, grade 3, ER-, PR-, Her2^+^, Ki-67-positive cells (34%). Lymph node-positive (1 lymph nodes with mts from 15 lymph nodes). Six courses of neoadjuvant chemotherapy. Patient #7: 64-year-old woman, invasive carcinoma (left side), T2N0M0, grade 2, ER-, PR-, Her2^+^, Ki-67-positive cells (70%). Lymph node-negative (0 lymph nodes with mts from 6 lymph nodes). Six courses of neoadjuvant chemotherapy. Patient #8: 41-year-old woman, invasive carcinoma (left side), T2N1M0, grade2, ER-, PR-, Her2^+^, Ki-67-positive cells (76%). Lymph node-negative (0 lymph nodes with mts from 8 lymph nodes). Six courses of neoadjuvant chemotherapy. Patient #9: 70-year-old woman, invasive carcinoma (right side), T2N0M0, grade 3, ER-, PR-, Her2^+^, Ki-67-positive cells (75%). Lymph node-negative (0 lymph nodes with mts from 10 lymph nodes). Six courses of neoadjuvant chemotherapy. Patient #10: 51-year-old woman, invasive carcinoma (right side), T2N1M0, grade 2, ER-, PR-, Her2^+^, Ki-67-positive cells (60%). Lymph node-positive (2 lymph nodes with mts from 6 lymph nodes). Six courses of neoadjuvant chemotherapy. Patient #11: 50-year-old woman, invasive carcinoma (left side), T2N1M0, grade

### Supplementary information on signaling receptors and proteins found in positive late endosomes (fig. S6)

#### EphA4 and TGF-β-receptors are present in flotillin-positive late endosomes

In addition to AXL, we investigated the cellular distribution of EphA4 that appears, in our phospho-RTK array experiments, at the top of the list of RTKs with increased tyrosine phosphorylation (fig. S6). We expressed EphA4-GFP in MCF10AF1F2 cells and observed its accumulation in flotillin-positive late endosomes that contained LAMP1 (fig. S6A). In MCF10A cells, detection of endogenous EphA4 by immunofluorescence using available antibodies gave very poor signals. However, the use of antibodies against P-Y588/Y596-EphA4 allowed observing a strong staining at cell-cell contacts in MCF10AmCh cells. Conversely, in MCF10AF1F2 cells, the staining decreased at the cell-cell contacts and concomitantly increased in intracellular vesicles, most of which were flotillin-positive (fig. S6D). In NMuMG cells, endogenous EphA4 was clearly detectable by immunofluorescence staining, and we observed a change in its distribution upon flotillin-upregulation. Indeed, compared with NMuMGmCh cells, NMuMGF1F2 cells exhibited a marked EphA4 signal in intracellular vesicles that were flotillin 2-mCherry- and CD63-positive (fig. S6B and C). These results strongly suggest that flotillin-upregulation stimulated EphA4 relocation, in its phosphorylated active form, ito flotillin-positive late endosomes.

As we found that flotillin-upregulation activates the TGF-β signaling pathway in MCF10A cells (fig. 2J, K) and NMuMG cells (fig. S2C, D), we examined the distribution of the TGF-β-receptor (TGF-β-R) in MCF10AF1F2 cells. Co-expression of TGF-β-RI-GFP and TGF-β-RII-Flag (the two TGF-β−R subunits) showed that both subunits co-localized with flotillins at the PM and in intracellular vesicles (fig. S6E). Moreover, we observed TGF-β-RI co-trafficking with flotillin2-mCherry in CD63-positive vesicles (fig. S6F).

Altogether, these results show that EphA4 and TGF-β-R, like AXL, are present in flotillin-microdomains at the PM and in flotillin-positive late endosomes. They suggest that these membrane compartments represent platforms from which these receptors might signal.

#### K-Ras4B localizes to flotillin-positive endosomes in MCF10AF1F2 cells

We next focused on Ras, a key protein involved in signal transmission from RTKs to the ERK/MAPK and PI3-K/AKT pathways and activated upon flotillin upregulation (fig. 2B-E, and fig. S2B). We separately expressed the GFP-tagged K-Ras4A, K-Ras4B, H-Ras and N-Ras, isoforms in MCF10AmCh and MCF10AF1F2 cells. Flotillin upregulation modified the distribution only of K-Ras4B (fig. S6G) but not of the other Ras isoforms (shown for Kras4A fig. S6G). K-Ras4B was localized at the PM in MCF10AmCh cells, and in flotillin-positive vesicles in MCF10AF1F2 cells (fig. S6G). Interestingly, this correlated with the co-immunoprecipitation of K-Ras4B, but not K-Ras4A, with flotillin 1 (fig. S6H).

### Supplementary Tables

**Table S1:** List of the 802 genes (232 upregulated and 570 downregulated genes) that were found to be differentially expressed between MCF10AmCh- and MCF10AF1F2 cells (fold change: 2.8, *P*<0.05) using the Deseq2 and Tuxedo pipelines. This analysis was performed using three replicates per condition to compare transcripts levels between MCF10AmCh and MCF10AF1F2 cells.

**Table S2:** List of the enriched GO terms and the enriched pathways obtained from the analysis of the 802 differentially expressed genes between MCF10mCh and MCF10AF1F2 cells using three distinct bioinfomatic tools (KEGG, DAVID, and Genomatix). Only the data with p value <0.001 were retained.

### Supplementary videos legends

***Video 1:*** *Flotillin oligomerization induced by a CRY2-CIBN-based optogenetic approach promotes AXL co-clustering*. Movie of the boxed area from figure 4B. MCF10A cells that co-express CRY2-mCitrin (green), flotillin 2-CIBN-mCherry (red) and AXL-Halo (blue) labeled with Halo-Tag-Janelia Fluor 646 were imaged by TIRF microscopy. First, flotillin 2-CIBN-mCherry and AXL-Halo were imaged for 8 seconds before 488nm illumination. Then, cells were globally illuminated by 20 pulses (1 pulse/s, 100 ms time length for each) of 488nm-light to stimulate and visualize CRY2. During and after illumination, flotillin 2-CIBN-mCherry and AXL-Halo were continuously imaged. Two examples of flotillin 2/AXL-positive clusters induced by CRY2 oligomerization initiated by illumination are indicated (closed arrow). Co-localization events between flotillin 2-CIBN-mCherry and AXL-Halo are also visible independently of CRY2 oligomerization before illumination and correspond mostly to AXL-containing vesicles positive for flotillin 2 passing through the 200 nm depth of field illuminated by TIRF. Thanks to the high concentration of flotillin 2-CIBN-mCh in these vesicles, CRY2 translocation could be observed. Scale bar: 2 µm.

***Video 2:*** *Co-endocytosis of flotillin 2 and AXL after their co-accumulation at the plasma membrane.* MCF10AF1F2 cells that co-express flotillin 1-HA, flotillin 2-mCherry and AXL-GFP were imaged by spinning disk confocal microscopy (1 frame/1.3 s. Scale bar: 2.5 µm). Arrows indicate the site of co-accumulation of AXL and flotillin 2 and then the emerging endocytic vesicle.

***Video 3:*** *Flotillin 2 and AXL colocalize in intracellular vesicles.* MCF10AF1F2 cells that co-express flotillin 1-HA, flotillin 2-mCherry and AXL-GFP were imaged by spinning disk confocal microscopy; 1 frame/1.3 s. Scale bar: 5 µm.

***Video 4:*** *Flotillin 2 and AXL colocalize in intracellular vesicles in tumor cells.* One MDA-MBA-MB-231 cell that co-expresses flotillin1-HA, flotillin-2-Cherry and AXL-GFP was imaged by spinning disk confocal microscopy; 1 frame/0.8 s. Scale bar: 10µm.

***Video 5:*** *SphK2 is abundant in flotillin-positive late endosomes in MDA-MB-231 cells.* Movie of the boxed area shown in figure 5A acquired by spinning disk confocal microscopy of one MDA-MB231 cell that co-expresses flotillin 1-mCherry and SphK2-GFP (1 frame/s. Scale bar: 2 µm).

***Video 6:*** *Co-localization and co-trafficking of AXL-Halo, SphK2-GFP and flotillin 2-mCherry in one MCF10AF1F2 cell.* Movie acquired by spinning disk confocal microscopy of one MCF10AF1F2 cell that co-expresses flotillin 2-mCherry (red), SphK2-GFP (green) and AXL-Halo (blue) (labeled with Halo-tag-Janelia Fluor 646) (1 frame/10 s. Scale bar: 10 µm).

***Video 7:*** *Flotillin 2-mCherry/SphK2-GFP co-endocytosis in MCF10AF1F2 cells.* Movie of the boxed area shown in figure 5D (upper panel) acquired by spinning disk confocal microscopy of one MCF10AF1F2 cell that co-expresses flotillin 2-mCherry and SphK2-GFP (1 frame/2 s). At 34s, arrows indicate sites of flotillin 2-mCherry and SphK2-GFP co-accumulation at the plasma membrane followed by endocytosis of a vesicle positive for both proteins. (Scale bar: 2µm).

***Video 8:*** *Flotillin 1-mCherry/SphK2-GFP co-endocytosis in MDA-MB-231 cells.* Movie of the boxed area shown in figure 5D (lower panel) acquired by spinning disk confocal microscopy of one MDA-MB-231 cell that co-expresses flotillin 1-mCherry and SphK2-GFP. The arrow indicates the site of flotillin 1-mCherry and SphK2-GFP co-accumulation at the plasma membrane followed by endocytosis of one vesicle positive for both proteins (1 frame/1 s). (Scale bar: 2 µm).

***Video 9:*** *Inhibition of SphK2 catalytic activity induces its delocalization from flotillin-positive endosomes*. One MCF10AF1F2 cell that co-expressing SphK2-GFP was imaged by spinning disk confocal microscopy immediately after addition of opaganib (50 µM) (1 frame/30 s. Scale bar: 10 µm). The disappearance of the SphK2-GFP signal in flotillin 2-mCherry-positive vesicles is clearly visible after 8 min and lasted for the rest of the movie (up to 28,5 min).

## Abbreviations

EMT: Epithelial-to-mesenchymal transition

PM: Plasma membrane

RTKs: Receptor Tyrosine Kinases

SphK2: Sphingosine kinase 2

S1P: Sphingosine 1-phosphate

UFIT: Upregulated Flotillin-Induced Trafficking

## Notes

### Competing Interest Statement

The authors have declared no competing interest.

### Summary of Updates

Additional data in Figure 1 and Figure 6

