## Supplementary material for "Upregulated-flotillins and sphingosine kinase 2 derail vesicular trafic to stabilize AXL and promote epithelial-mesenchymal transition": Genest Tables S1 and S2

| **Supplementary Table S1** | The differential gene expression analysis was processed on three replicates per condition to compare the levels of the transcripts between MCF10AmCh and MCF10AF1F2 cells<br>Confrontation of the results obtained from the Deseq2 and Tuxedo analyses |
|---|---|

EMT markers

| 802 common genes | test_id | gene_id | feature_type | MCF10A-mCh | MCF10A-F1F2 | MCF10A-mCh mean_1 | MCF10A-F1F2 mean_2 | log2FoldChange | pvalue | padj | significant |
|---|---|---|---|---|---|---|---|---|---|---|---|
| ARMCX3 | ENSG00000102401 | ENSG00000102401 | protein_coding | mCh_JDV-1-3-5 | F1F2_JDV-2-4-6 | 15,4465636 | 658,766641 | 5,167195894 | 6,28E-65 | 1,24E-61 | yes |
| FMN2 | ENSG00000155816 | ENSG00000155816 | protein_coding | mCh_JDV-1-3-5 | F1F2_JDV-2-4-6 | 6,095810252 | 278,6178204 | 5,085693073 | 1,29E-40 | 7,26E-38 | yes |
| ABCA8 | ENSG00000141338 | ENSG00000141338 | protein_coding | mCh_JDV-1-3-5 | F1F2_JDV-2-4-6 | 13,44623078 | 461,7690579 | 4,645926346 | 2,57E-29 | 8,04E-27 | yes |
| FRMD3 | ENSG00000172159 | ENSG00000172159 | protein_coding | mCh_JDV-1-3-5 | F1F2_JDV-2-4-6 | 18,29777519 | 397,8531995 | 4,228398312 | 7,11E-42 | 4,38E-39 | yes |
| ATP1A2 | ENSG00000018625 | ENSG00000018625 | protein_coding | mCh_JDV-1-3-5 | F1F2_JDV-2-4-6 | 8,641073242 | 236,9175498 | 4,202429862 | 3,47E-19 | 4,82E-17 | yes |
| FN1 | ENSG00000115414 | ENSG00000115414 | protein_coding | mCh_JDV-1-3-5 | F1F2_JDV-2-4-6 | 5648,306615 | 157919,9993 | 4,099905327 | 2,43E-15 | 2,21E-13 | yes |
| ANKRD1 | ENSG00000148677 | ENSG00000148677 | protein_coding | mCh_JDV-1-3-5 | F1F2_JDV-2-4-6 | 5,543040237 | 148,8739184 | 4,057892519 | 1,35E-15 | 1,29E-13 | yes |
| ATOH8 | ENSG00000168874 | ENSG00000168874 | protein_coding | mCh_JDV-1-3-5 | F1F2_JDV-2-4-6 | 40,96632357 | 702,054822 | 3,910057154 | 2,99E-37 | 1,47E-34 | yes |
| CTGF | ENSG00000118523 | ENSG00000118523 | protein_coding | mCh_JDV-1-3-5 | F1F2_JDV-2-4-6 | 197,2669763 | 3253,986731 | 3,888619963 | 5,79E-43 | 3,81E-40 | yes |
| CADM3 | ENSG00000162706 | ENSG00000162706 | protein_coding | mCh_JDV-1-3-5 | F1F2_JDV-2-4-6 | 1612,383633 | 32934,98596 | 3,876623264 | 2,63E-17 | 3,05E-15 | yes |
| C10orf90 | ENSG00000154493 | ENSG00000154493 | protein_coding | mCh_JDV-1-3-5 | F1F2_JDV-2-4-6 | 6,14981957 | 161,2178771 | 3,871716259 | 2,25E-12 | 1,43E-10 | yes |
| GNG2 | ENSG00000186469 | ENSG00000186469 | protein_coding | mCh_JDV-1-3-5 | F1F2_JDV-2-4-6 | 3,231749657 | 71,18274971 | 3,747949356 | 7,08E-13 | 4,75E-11 | yes |
| AOX1 | ENSG00000138356 | ENSG00000138356 | protein_coding | mCh_JDV-1-3-5 | F1F2_JDV-2-4-6 | 323,5224412 | 4975,164879 | 3,712311509 | 5,83E-27 | 1,48E-24 | yes |
| CNRIP1 | ENSG00000119865 | ENSG00000119865 | protein_coding | mCh_JDV-1-3-5 | F1F2_JDV-2-4-6 | 6,764275697 | 116,3980358 | 3,711238684 | 2,85E-18 | 3,56E-16 | yes |
| SERPINE2 | ENSG00000135919 | ENSG00000135919 | protein_coding | mCh_JDV-1-3-5 | F1F2_JDV-2-4-6 | 433,1458825 | 6169,794053 | 3,672945809 | 8,46E-36 | 3,88E-33 | yes |
| PDE4B | ENSG00000184588 | ENSG00000184588 | protein_coding | mCh_JDV-1-3-5 | F1F2_JDV-2-4-6 | 31,47426286 | 453,7997936 | 3,653934389 | 7,68E-30 | 2,57E-27 | yes |
| MAP1B | ENSG00000131711 | ENSG00000131711 | protein_coding | mCh_JDV-1-3-5 | F1F2_JDV-2-4-6 | 67,92048035 | 1141,081462 | 3,611633991 | 6,96E-15 | 5,92E-13 | yes |
| AFF2 | ENSG00000155966 | ENSG00000155966 | protein_coding | mCh_JDV-1-3-5 | F1F2_JDV-2-4-6 | 11,80084671 | 190,7839172 | 3,566627639 | 6E-15 | 5,17E-13 | yes |
| IFLTD1 | ENSG00000152936 | ENSG00000152936 | protein_coding | mCh_JDV-1-3-5 | F1F2_JDV-2-4-6 | 1,930806862 | 41,23724265 | 3,513020162 | 6,36E-10 | 2,83E-08 | yes |
| APCDD1L | ENSG00000198768 | ENSG00000198768 | protein_coding | mCh_JDV-1-3-5 | F1F2_JDV-2-4-6 | 5,113773767 | 109,7248173 | 3,510949844 | 1,62E-09 | 6,57E-08 | yes |
| P4HA3 | ENSG00000149380 | ENSG00000149380 | protein_coding | mCh_JDV-1-3-5 | F1F2_JDV-2-4-6 | 6,43007015 | 131,4719391 | 3,508983275 | 8E-10 | 0,000000035 | yes |
| RBP4 | ENSG00000138207 | ENSG00000138207 | protein_coding | mCh_JDV-1-3-5 | F1F2_JDV-2-4-6 | 40,93017243 | 659,6924232 | 3,49623947 | 9,03E-13 | 5,96E-11 | yes |
| HSPB8 | ENSG00000152137 | ENSG00000152137 | protein_coding | mCh_JDV-1-3-5 | F1F2_JDV-2-4-6 | 38,16909869 | 495,0896951 | 3,434031794 | 1,09E-19 | 1,55E-17 | yes |
| FLOT1 | ENSG00000137312 | ENSG00000137312 | protein_coding | mCh_JDV-1-3-5 | F1F2_JDV-2-4-6 | 1474,457542 | 16770,5792 | 3,383070267 | 2,3E-35 | 1,03E-32 | yes |
| WIPF1 | ENSG00000115935 | ENSG00000115935 | protein_coding | mCh_JDV-1-3-5 | F1F2_JDV-2-4-6 | 88,46823135 | 1033,525772 | 3,376284645 | 5,31E-27 | 1,4E-24 | yes |
| NEDD9 | ENSG00000111859 | ENSG00000111859 | protein_coding | mCh_JDV-1-3-5 | F1F2_JDV-2-4-6 | 38,75003957 | 444,7297978 | 3,362948459 | 1,02E-28 | 3E-26 | yes |
| PRSS23 | ENSG00000150687 | ENSG00000150687 | protein_coding | mCh_JDV-1-3-5 | F1F2_JDV-2-4-6 | 34,68274211 | 506,5345909 | 3,35838291 | 1,23E-11 | 7,25E-10 | yes |
| FLOT2 | ENSG00000132589 | ENSG00000132589 | protein_coding | mCh_JDV-1-3-5 | F1F2_JDV-2-4-6 | 3727,152142 | 41052,11864 | 3,349156683 | 9,89E-38 | 5,14E-35 | yes |
| SULT1B1 | ENSG00000173597 | ENSG00000173597 | protein_coding | mCh_JDV-1-3-5 | F1F2_JDV-2-4-6 | 10,21458982 | 132,920064 | 3,342652368 | 6,39E-15 | 5,48E-13 | yes |
| HAS2 | ENSG00000170961 | ENSG00000170961 | protein_coding | mCh_JDV-1-3-5 | F1F2_JDV-2-4-6 | 6,39925427 | 84,27912801 | 3,316654176 | 1,28E-13 | 9,37E-12 | yes |
| TDRD9 | ENSG00000156414 | ENSG00000156414 | protein_coding | mCh_JDV-1-3-5 | F1F2_JDV-2-4-6 | 5,437634885 | 77,14157543 | 3,307954514 | 1,93E-11 | 1,1E-09 | yes |
| TMEM100 | ENSG00000166292 | ENSG00000166292 | protein_coding | mCh_JDV-1-3-5 | F1F2_JDV-2-4-6 | 15,39767214 | 192,6982623 | 3,289319285 | 3,18E-14 | 2,52E-12 | yes |
| FBLN5 | ENSG00000140092 | ENSG00000140092 | protein_coding | mCh_JDV-1-3-5 | F1F2_JDV-2-4-6 | 67,16454469 | 768,6489892 | 3,157186529 | 1,26E-12 | 8,18E-11 | yes |
| RAB31 | ENSG00000168461 | ENSG00000168461 | protein_coding | mCh_JDV-1-3-5 | F1F2_JDV-2-4-6 | 90,58651505 | 896,1721585 | 3,14200629 | 7,33E-23 | 1,48E-20 | yes |
| MMP2 | ENSG00000087245 | ENSG00000087245 | protein_coding | mCh_JDV-1-3-5 | F1F2_JDV-2-4-6 | 176,07057 | 1920,94012 | 3,135727856 | 1,02E-13 | 7,51E-12 | yes |
| KIAA1644 | ENSG00000138944 | ENSG00000138944 | protein_coding | mCh_JDV-1-3-5 | F1F2_JDV-2-4-6 | 39,84254491 | 430,0465831 | 3,130714258 | 4,74E-14 | 3,68E-12 | yes |
| DARC | ENSG00000213088 | ENSG00000213088 | protein_coding | mCh_JDV-1-3-5 | F1F2_JDV-2-4-6 | 393,7097951 | 5368,798832 | 3,125683758 | 1,44E-08 | 0,000000488 | yes |
| SPOCK1 | ENSG00000152377 | ENSG00000152377 | protein_coding | mCh_JDV-1-3-5 | F1F2_JDV-2-4-6 | 173,2210433 | 1863,166178 | 3,052321463 | 2,83E-11 | 1,56E-09 | yes |
| CCDC18 | ENSG00000122483 | ENSG00000122483 | protein_coding | mCh_JDV-1-3-5 | F1F2_JDV-2-4-6 | 425,2598256 | 4296,139193 | 2,998968421 | 1,37E-11 | 8,02E-10 | yes |
| FBN1 | ENSG00000166147 | ENSG00000166147 | protein_coding | mCh_JDV-1-3-5 | F1F2_JDV-2-4-6 | 261,3570845 | 2545,446104 | 2,978507522 | 3,09E-12 | 1,93E-10 | yes |

| | | | | | | | | | | | |
|---|---|---|---|---|---|---|---|---|---|---|---|
| CRP | ENSG00000132693 | ENSG00000132693 | protein_coding | mCh_JDV-1-3-5 | F1F2_JDV-2-4-6 | 1,920516785 | 27,64829383 | 2,955917042 | 0,000000831 | 0,0000193 | yes |
| RPS6KA2 | ENSG00000071242 | ENSG00000071242 | protein_coding | mCh_JDV-1-3-5 | F1F2_JDV-2-4-6 | 563,9528851 | 4675,833138 | 2,951566976 | 2,09E-29 | 6,65E-27 | yes |
| CD163L1 | ENSG00000177675 | ENSG00000177675 | protein_coding | mCh_JDV-1-3-5 | F1F2_JDV-2-4-6 | 8,121786741 | 80,4513959 | 2,912544375 | 8,55E-10 | 3,73E-08 | yes |
| PRR16 | ENSG00000184838 | ENSG00000184838 | protein_coding | mCh_JDV-1-3-5 | F1F2_JDV-2-4-6 | 28,99577212 | 263,2300776 | 2,898280472 | 3,6E-12 | 2,24E-10 | yes |
| CCDC102B | ENSG00000150636 | ENSG00000150636 | protein_coding | mCh_JDV-1-3-5 | F1F2_JDV-2-4-6 | 25,01315943 | 214,007862 | 2,895058124 | 1,16E-15 | 1,12E-13 | yes |
| DCN | ENSG00000011465 | ENSG00000011465 | protein_coding | mCh_JDV-1-3-5 | F1F2_JDV-2-4-6 | 4658,716599 | 42466,77544 | 2,792132714 | 8,54E-09 | 0,000000301 | yes |
| ZCCHC24 | ENSG00000165424 | ENSG00000165424 | protein_coding | mCh_JDV-1-3-5 | F1F2_JDV-2-4-6 | 81,29742519 | 595,9287946 | 2,789356157 | 3,31E-29 | 1,02E-26 | yes |
| PCDH9 | ENSG00000184226 | ENSG00000184226 | protein_coding | mCh_JDV-1-3-5 | F1F2_JDV-2-4-6 | 161,9622191 | 1211,834047 | 2,785669614 | 1,05E-21 | 1,88E-19 | yes |
| NEGR1 | ENSG00000172260 | ENSG00000172260 | protein_coding | mCh_JDV-1-3-5 | F1F2_JDV-2-4-6 | 43,87144489 | 380,3385042 | 2,775929719 | 1,41E-09 | 5,76E-08 | yes |
| GNAZ | ENSG00000128266 | ENSG00000128266 | protein_coding | mCh_JDV-1-3-5 | F1F2_JDV-2-4-6 | 7,116502467 | 61,18787421 | 2,772682598 | 8,07E-10 | 3,52E-08 | yes |
| IRS1 | ENSG00000169047 | ENSG00000169047 | protein_coding | mCh_JDV-1-3-5 | F1F2_JDV-2-4-6 | 99,15027683 | 742,00931 | 2,746400153 | 1,46E-16 | 1,55E-14 | yes |
| ZNF229 | ENSG00000167383 | ENSG00000167383 | protein_coding | mCh_JDV-1-3-5 | F1F2_JDV-2-4-6 | 12,48727903 | 96,47685168 | 2,695084711 | 4,54E-11 | 2,42E-09 | yes |
| NR2F1 | ENSG00000175745 | ENSG00000175745 | protein_coding | mCh_JDV-1-3-5 | F1F2_JDV-2-4-6 | 27,73836289 | 203,7109186 | 2,693032805 | 7,36E-14 | 5,59E-12 | yes |
| PAPPA | ENSG00000182752 | ENSG00000182752 | protein_coding | mCh_JDV-1-3-5 | F1F2_JDV-2-4-6 | 352,837808 | 2369,854755 | 2,627272871 | 2,85E-18 | 3,56E-16 | yes |
| FAM19A2 | ENSG00000198673 | ENSG00000198673 | protein_coding | mCh_JDV-1-3-5 | F1F2_JDV-2-4-6 | 12,75206731 | 137,0535761 | 2,592242349 | 0,0000341 | 0,000479744 | yes |
| NPTXR | ENSG00000221890 | ENSG00000221890 | protein_coding | mCh_JDV-1-3-5 | F1F2_JDV-2-4-6 | 27,92611148 | 183,0293853 | 2,574555988 | 1,23E-15 | 1,19E-13 | yes |
| ANXA6 | ENSG00000197043 | ENSG00000197043 | protein_coding | mCh_JDV-1-3-5 | F1F2_JDV-2-4-6 | 244,5458087 | 1631,402584 | 2,557600974 | 2,54E-12 | 1,61E-10 | yes |
| SHISA9 | ENSG00000237515 | ENSG00000237515 | protein_coding | mCh_JDV-1-3-5 | F1F2_JDV-2-4-6 | 13,80368411 | 111,2260325 | 2,556535156 | 0,00000102 | 0,0000234 | yes |
| APOBEC3H | ENSG00000100298 | ENSG00000100298 | protein_coding | mCh_JDV-1-3-5 | F1F2_JDV-2-4-6 | 11,54123063 | 81,54617906 | 2,530839854 | 1,15E-08 | 0,000000394 | yes |
| RERG | ENSG00000134533 | ENSG00000134533 | protein_coding | mCh_JDV-1-3-5 | F1F2_JDV-2-4-6 | 234,6436621 | 1527,745115 | 2,508345004 | 4,32E-11 | 2,31E-09 | yes |
| CA8 | ENSG00000178538 | ENSG00000178538 | protein_coding | mCh_JDV-1-3-5 | F1F2_JDV-2-4-6 | 14,95809298 | 122,2816014 | 2,502494313 | 0,00000672 | 0,000120067 | yes |
| SRGN | ENSG00000122862 | ENSG00000122862 | protein_coding | mCh_JDV-1-3-5 | F1F2_JDV-2-4-6 | 702,0083045 | 4148,636462 | 2,479739035 | 2,16E-21 | 3,74E-19 | yes |
| ZEB1 | ENSG00000148516 | ENSG00000148516 | protein_coding | mCh_JDV-1-3-5 | F1F2_JDV-2-4-6 | 155,8897245 | 908,8078566 | 2,449469315 | 1,04E-18 | 1,38E-16 | yes |
| PLAC8 | ENSG00000145287 | ENSG00000145287 | protein_coding | mCh_JDV-1-3-5 | F1F2_JDV-2-4-6 | 45,00510143 | 276,6339341 | 2,434653093 | 1,04E-10 | 5,29E-09 | yes |
| RXFP1 | ENSG00000171509 | ENSG00000171509 | protein_coding | mCh_JDV-1-3-5 | F1F2_JDV-2-4-6 | 24,25981443 | 171,4976505 | 2,424501473 | 0,00000194 | 0,0000409 | yes |
| HLA-DPA1 | ENSG00000231389 | ENSG00000231389 | protein_coding | mCh_JDV-1-3-5 | F1F2_JDV-2-4-6 | 9,718333705 | 65,40980981 | 2,419539733 | 0,00000041 | 0,0000102 | yes |
| GNG11 | ENSG00000127920 | ENSG00000127920 | protein_coding | mCh_JDV-1-3-5 | F1F2_JDV-2-4-6 | 142,1347009 | 824,39926 | 2,403484661 | 2,47E-13 | 1,75E-11 | yes |
| F8 | ENSG00000185010 | ENSG00000185010 | protein_coding | mCh_JDV-1-3-5 | F1F2_JDV-2-4-6 | 96,95759845 | 529,9585619 | 2,373833491 | 1,81E-20 | 2,8E-18 | yes |
| RTN1 | ENSG00000139970 | ENSG00000139970 | protein_coding | mCh_JDV-1-3-5 | F1F2_JDV-2-4-6 | 899,1369307 | 5067,255373 | 2,371791398 | 9,7E-14 | 7,17E-12 | yes |
| SDC2 | ENSG00000169439 | ENSG00000169439 | protein_coding | mCh_JDV-1-3-5 | F1F2_JDV-2-4-6 | 11,56948758 | 70,77922221 | 2,368066644 | 2,75E-08 | 0,000000894 | yes |
| HSF2BP | ENSG00000160207 | ENSG00000160207 | protein_coding | mCh_JDV-1-3-5 | F1F2_JDV-2-4-6 | 46,86913605 | 285,6901074 | 2,360066362 | 4,51E-08 | 0,0000014 | yes |
| CAPN5 | ENSG00000149260 | ENSG00000149260 | protein_coding | mCh_JDV-1-3-5 | F1F2_JDV-2-4-6 | 17,0097901 | 110,274262 | 2,356094138 | 0,00000127 | 0,0000286 | yes |
| COL28A1 | ENSG00000215018 | ENSG00000215018 | protein_coding | mCh_JDV-1-3-5 | F1F2_JDV-2-4-6 | 131,6831176 | 914,9009163 | 2,339927092 | 0,0000163 | 0,000256197 | yes |
| SIX2 | ENSG00000170577 | ENSG00000170577 | protein_coding | mCh_JDV-1-3-5 | F1F2_JDV-2-4-6 | 21,23892983 | 126,8194066 | 2,330194919 | 7,34E-08 | 0,00000221 | yes |
| CXCR7 | ENSG00000144476 | ENSG00000144476 | protein_coding | mCh_JDV-1-3-5 | F1F2_JDV-2-4-6 | 9,970327337 | 64,24062037 | 2,299200924 | 0,00000769 | 0,000134817 | yes |
| SORBS2 | ENSG00000154556 | ENSG00000154556 | protein_coding | mCh_JDV-1-3-5 | F1F2_JDV-2-4-6 | 222,910806 | 1180,005041 | 2,282902537 | 1,41E-12 | 9,13E-11 | yes |
| PPAPDC1A | ENSG00000203805 | ENSG00000203805 | protein_coding | mCh_JDV-1-3-5 | F1F2_JDV-2-4-6 | 6,435188013 | 38,48710963 | 2,282413744 | 0,000000986 | 0,0000227 | yes |
| DUSP23 | ENSG00000158716 | ENSG00000158716 | protein_coding | mCh_JDV-1-3-5 | F1F2_JDV-2-4-6 | 488,8646558 | 2533,297089 | 2,271632954 | 2,74E-14 | 2,19E-12 | yes |
| CYR61 | ENSG00000142871 | ENSG00000142871 | protein_coding | mCh_JDV-1-3-5 | F1F2_JDV-2-4-6 | 597,0750731 | 3078,193597 | 2,261258845 | 8,8E-14 | 6,6E-12 | yes |
| ZEB2 | ENSG00000169554 | ENSG00000169554 | protein_coding | mCh_JDV-1-3-5 | F1F2_JDV-2-4-6 | 150,7940263 | 748,8305013 | 2,241006425 | 1,02E-18 | 1,36E-16 | yes |
| DKK1 | ENSG00000107984 | ENSG00000107984 | protein_coding | mCh_JDV-1-3-5 | F1F2_JDV-2-4-6 | 2069,813964 | 10674,71668 | 2,235254717 | 3,42E-11 | 1,86E-09 | yes |
| IL1RAPL2 | ENSG00000189108 | ENSG00000189108 | protein_coding | mCh_JDV-1-3-5 | F1F2_JDV-2-4-6 | 6,838865169 | 46,46966506 | 2,223085747 | 0,000127993 | 0,001422924 | yes |
| BEND6 | ENSG00000151917 | ENSG00000151917 | protein_coding | mCh_JDV-1-3-5 | F1F2_JDV-2-4-6 | 26,28328634 | 133,8944798 | 2,220919779 | 2,2E-11 | 1,24E-09 | yes |
| COL24A1 | ENSG00000171502 | ENSG00000171502 | protein_coding | mCh_JDV-1-3-5 | F1F2_JDV-2-4-6 | 18,19231548 | 103,3625337 | 2,200009175 | 0,00000444 | 0,0000837 | yes |
| ADRA1B | ENSG00000170214 | ENSG00000170214 | protein_coding | mCh_JDV-1-3-5 | F1F2_JDV-2-4-6 | 9,00364444 | 49,38502915 | 2,195013607 | 0,00000109 | 0,0000248 | yes |
| GPR173 | ENSG00000184194 | ENSG00000184194 | protein_coding | mCh_JDV-1-3-5 | F1F2_JDV-2-4-6 | 21,04875364 | 111,4219128 | 2,185365741 | 0,000000198 | 0,00000532 | yes |
| PRKAR2B | ENSG00000005249 | ENSG00000005249 | protein_coding | mCh_JDV-1-3-5 | F1F2_JDV-2-4-6 | 43,82003974 | 221,4107852 | 2,175667334 | 4,93E-09 | 0,000000182 | yes |

| | | | | | | | | | | | |
|---|---|---|---|---|---|---|---|---|---|---|---|
| SMARCA1 | ENSG00000102038 | ENSG00000102038 | protein_coding | mCh_JDV-1-3-5 | F1F2_JDV-2-4-6 | 64,50875849 | 319,8876286 | 2,165006742 | 1,26E-09 | 5,17E-08 | yes |
| PRR5L | ENSG00000135362 | ENSG00000135362 | protein_coding | mCh_JDV-1-3-5 | F1F2_JDV-2-4-6 | 69,45561296 | 353,0264817 | 2,162070287 | 4,67E-08 | 0,00000144 | yes |
| TXNRD1 | ENSG00000198431 | ENSG00000198431 | protein_coding | mCh_JDV-1-3-5 | F1F2_JDV-2-4-6 | 3427,017173 | 15840,37646 | 2,157864626 | 1,43E-22 | 2,77E-20 | yes |
| JAKMIP3 | ENSG00000188385 | ENSG00000188385 | protein_coding | mCh_JDV-1-3-5 | F1F2_JDV-2-4-6 | 10,59501919 | 56,02667508 | 2,147491857 | 0,0000019 | 0,0000401 | yes |
| PTGFR | ENSG00000122420 | ENSG00000122420 | protein_coding | mCh_JDV-1-3-5 | F1F2_JDV-2-4-6 | 582,9502357 | 3395,135467 | 2,144567524 | 0,0000591 | 0,000759315 | yes |
| SMAD6 | ENSG00000137834 | ENSG00000137834 | protein_coding | mCh_JDV-1-3-5 | F1F2_JDV-2-4-6 | 4,836027766 | 30,24159642 | 2,140216573 | 0,00018359 | 0,001931122 | yes |
| INSC | ENSG00000188487 | ENSG00000188487 | protein_coding | mCh_JDV-1-3-5 | F1F2_JDV-2-4-6 | 52,16314467 | 270,3635934 | 2,127558856 | 0,00000209 | 0,0000435 | yes |
| PTX3 | ENSG00000163661 | ENSG00000163661 | protein_coding | mCh_JDV-1-3-5 | F1F2_JDV-2-4-6 | 78,7954149 | 411,6523105 | 2,121861576 | 0,00000413 | 0,0000784 | yes |
| LMCD1 | ENSG00000071282 | ENSG00000071282 | protein_coding | mCh_JDV-1-3-5 | F1F2_JDV-2-4-6 | 9,967714054 | 54,32083092 | 2,11996361 | 0,0000201 | 0,000306085 | yes |
| AK7 | ENSG00000140057 | ENSG00000140057 | protein_coding | mCh_JDV-1-3-5 | F1F2_JDV-2-4-6 | 5,831021963 | 32,08457443 | 2,11534643 | 0,0000323 | 0,000458282 | yes |
| CHST2 | ENSG00000175040 | ENSG00000175040 | protein_coding | mCh_JDV-1-3-5 | F1F2_JDV-2-4-6 | 53,15040772 | 250,3323496 | 2,111754775 | 3,86E-10 | 1,81E-08 | yes |
| C4orf34 | ENSG00000163683 | ENSG00000163683 | protein_coding | mCh_JDV-1-3-5 | F1F2_JDV-2-4-6 | 162,1220277 | 721,7686164 | 2,101499323 | 2,86E-20 | 4,28E-18 | yes |
| SH3BGRL | ENSG00000131171 | ENSG00000131171 | protein_coding | mCh_JDV-1-3-5 | F1F2_JDV-2-4-6 | 768,4843474 | 3431,60203 | 2,100957454 | 1,1E-18 | 1,45E-16 | yes |
| TBL1X | ENSG00000101849 | ENSG00000101849 | protein_coding | mCh_JDV-1-3-5 | F1F2_JDV-2-4-6 | 168,7728861 | 787,4559859 | 2,097090103 | 6,03E-10 | 2,71E-08 | yes |
| KCNE3 | ENSG00000175538 | ENSG00000175538 | protein_coding | mCh_JDV-1-3-5 | F1F2_JDV-2-4-6 | 8,674556756 | 44,4652097 | 2,089815894 | 0,00000683 | 0,000121851 | yes |
| G6PD | ENSG00000160211 | ENSG00000160211 | protein_coding | mCh_JDV-1-3-5 | F1F2_JDV-2-4-6 | 1302,342479 | 5900,569295 | 2,077886096 | 2,36E-11 | 1,31E-09 | yes |
| GREB1L | ENSG00000141449 | ENSG00000141449 | protein_coding | mCh_JDV-1-3-5 | F1F2_JDV-2-4-6 | 82,4646061 | 387,0360267 | 2,05856985 | 0,000000166 | 0,00000456 | yes |
| IL6 | ENSG00000136244 | ENSG00000136244 | protein_coding | mCh_JDV-1-3-5 | F1F2_JDV-2-4-6 | 6,13691621 | 36,41302946 | 2,056978189 | 0,000425749 | 0,003813574 | yes |
| PMEPA1 | ENSG00000124225 | ENSG00000124225 | protein_coding | mCh_JDV-1-3-5 | F1F2_JDV-2-4-6 | 307,2572731 | 1383,435525 | 2,05563751 | 4,69E-10 | 2,16E-08 | yes |
| S100A4 | ENSG00000196154 | ENSG00000196154 | protein_coding | mCh_JDV-1-3-5 | F1F2_JDV-2-4-6 | 1175,45892 | 5540,555903 | 2,054124884 | 0,000000352 | 0,00000891 | yes |
| AKR1C2 | ENSG00000151632 | ENSG00000151632 | protein_coding | mCh_JDV-1-3-5 | F1F2_JDV-2-4-6 | 3835,966165 | 18927,35964 | 2,050593187 | 0,00000818 | 0,000141672 | yes |
| LUM | ENSG00000139329 | ENSG00000139329 | protein_coding | mCh_JDV-1-3-5 | F1F2_JDV-2-4-6 | 11,56170209 | 58,33388278 | 2,043799389 | 0,0000237 | 0,00035216 | yes |
| ZNF358 | ENSG00000198816 | ENSG00000198816 | protein_coding | mCh_JDV-1-3-5 | F1F2_JDV-2-4-6 | 85,08746581 | 386,1571383 | 2,04124677 | 1,59E-08 | 0,000000532 | yes |
| C17orf97 | ENSG00000187624 | ENSG00000187624 | protein_coding | mCh_JDV-1-3-5 | F1F2_JDV-2-4-6 | 11,22504631 | 54,34187775 | 2,040956086 | 0,00000443 | 0,0000835 | yes |
| CDO1 | ENSG00000129596 | ENSG00000129596 | protein_coding | mCh_JDV-1-3-5 | F1F2_JDV-2-4-6 | 9,960037259 | 47,23586293 | 2,038019936 | 0,00000156 | 0,0000342 | yes |
| UCHL1 | ENSG00000154277 | ENSG00000154277 | protein_coding | mCh_JDV-1-3-5 | F1F2_JDV-2-4-6 | 18,78366426 | 88,77267028 | 2,036726044 | 0,00000164 | 0,0000354 | yes |
| AKR1B1 | ENSG00000085662 | ENSG00000085662 | protein_coding | mCh_JDV-1-3-5 | F1F2_JDV-2-4-6 | 4924,63103 | 22568,69226 | 2,035808503 | 0,000000109 | 0,00000313 | yes |
| TMEM37 | ENSG00000171227 | ENSG00000171227 | protein_coding | mCh_JDV-1-3-5 | F1F2_JDV-2-4-6 | 32,60800549 | 146,4036001 | 2,023886883 | 2,47E-08 | 0,00000081 | yes |
| CDA | ENSG00000158825 | ENSG00000158825 | protein_coding | mCh_JDV-1-3-5 | F1F2_JDV-2-4-6 | 23,81261283 | 111,6461131 | 2,022545698 | 0,00000206 | 0,0000428 | yes |
| PLA2G16 | ENSG00000176485 | ENSG00000176485 | protein_coding | mCh_JDV-1-3-5 | F1F2_JDV-2-4-6 | 22,67620256 | 103,4144478 | 2,003637364 | 0,00000114 | 0,0000257 | yes |
| PDE7B | ENSG00000171408 | ENSG00000171408 | protein_coding | mCh_JDV-1-3-5 | F1F2_JDV-2-4-6 | 116,9953747 | 522,8552985 | 2,001861471 | 0,000000172 | 0,0000047 | yes |
| DLC1 | ENSG00000164741 | ENSG00000164741 | protein_coding | mCh_JDV-1-3-5 | F1F2_JDV-2-4-6 | 102,5339365 | 451,0093394 | 1,999001564 | 3,42E-08 | 0,00000109 | yes |
| EPS8 | ENSG00000151491 | ENSG00000151491 | protein_coding | mCh_JDV-1-3-5 | F1F2_JDV-2-4-6 | 1209,637336 | 5037,575332 | 1,998804743 | 5,42E-16 | 5,54E-14 | yes |
| SESN3 | ENSG00000149212 | ENSG00000149212 | protein_coding | mCh_JDV-1-3-5 | F1F2_JDV-2-4-6 | 17,0096814 | 77,0523491 | 1,997733852 | 0,000000822 | 0,0000192 | yes |
| STAC2 | ENSG00000141750 | ENSG00000141750 | protein_coding | mCh_JDV-1-3-5 | F1F2_JDV-2-4-6 | 184,1535605 | 882,6265445 | 1,990406055 | 0,0000306 | 0,000435392 | yes |
| KYNU | ENSG00000115919 | ENSG00000115919 | protein_coding | mCh_JDV-1-3-5 | F1F2_JDV-2-4-6 | 3183,157759 | 13715,15438 | 1,984587793 | 8,71E-09 | 0,000000306 | yes |
| PRUNE2 | ENSG00000106772 | ENSG00000106772 | protein_coding | mCh_JDV-1-3-5 | F1F2_JDV-2-4-6 | 13,29713792 | 69,65131212 | 1,982601395 | 0,000324923 | 0,003073683 | yes |
| MEIS3 | ENSG00000105419 | ENSG00000105419 | protein_coding | mCh_JDV-1-3-5 | F1F2_JDV-2-4-6 | 10,54629079 | 50,93925589 | 1,980541996 | 0,0000501 | 0,000660796 | yes |
| HIPK2 | ENSG00000064393 | ENSG00000064393 | protein_coding | mCh_JDV-1-3-5 | F1F2_JDV-2-4-6 | 4549,563203 | 18865,83824 | 1,974817965 | 1,84E-12 | 1,18E-10 | yes |
| APLP1 | ENSG00000105290 | ENSG00000105290 | protein_coding | mCh_JDV-1-3-5 | F1F2_JDV-2-4-6 | 16,43894451 | 73,18029385 | 1,9718335 | 0,00000148 | 0,0000329 | yes |
| SYDE1 | ENSG00000105137 | ENSG00000105137 | protein_coding | mCh_JDV-1-3-5 | F1F2_JDV-2-4-6 | 158,3834149 | 663,3457347 | 1,962777864 | 1,06E-09 | 4,48E-08 | yes |
| CITED2 | ENSG00000164442 | ENSG00000164442 | protein_coding | mCh_JDV-1-3-5 | F1F2_JDV-2-4-6 | 359,0089595 | 1584,51276 | 1,957115838 | 0,0000022 | 0,0000455 | yes |
| RNF157 | ENSG00000141576 | ENSG00000141576 | protein_coding | mCh_JDV-1-3-5 | F1F2_JDV-2-4-6 | 68,41675909 | 283,1033147 | 1,956162423 | 1,58E-10 | 7,78E-09 | yes |
| FAM129B | ENSG00000136830 | ENSG00000136830 | protein_coding | mCh_JDV-1-3-5 | F1F2_JDV-2-4-6 | 3473,944839 | 14671,27563 | 1,955944474 | 1,71E-08 | 0,000000568 | yes |
| FGF13 | ENSG00000129682 | ENSG00000129682 | protein_coding | mCh_JDV-1-3-5 | F1F2_JDV-2-4-6 | 10,32249975 | 48,18390986 | 1,951826301 | 0,0000492 | 0,000649722 | yes |
| OLFML3 | ENSG00000116774 | ENSG00000116774 | protein_coding | mCh_JDV-1-3-5 | F1F2_JDV-2-4-6 | 5,170287665 | 25,64249289 | 1,948768489 | 0,000248313 | 0,002468036 | yes |
| KB-1732A1.1 | ENSG00000253669 | ENSG00000253669 | protein_coding | mCh_JDV-1-3-5 | F1F2_JDV-2-4-6 | 20,34166419 | 91,24135067 | 1,945438739 | 0,000012 | 0,000198668 | yes |

| | | | | | | | | | | | |
|---|---|---|---|---|---|---|---|---|---|---|---|
| HKDC1 | ENSG00000156510 | ENSG00000156510 | protein_coding | mCh_JDV-1-3-5 | F1F2_JDV-2-4-6 | 9,448427553 | 53,73623388 | 1,943312348 | 0,001495587 | 0,010210951 | yes |
| AC108519.1 | ENSG00000226350 | ENSG00000226350 | protein_coding | mCh_JDV-1-3-5 | F1F2_JDV-2-4-6 | 2,241927674 | 13,50801875 | 1,93935704 | 0,002010997 | 0,012958029 | yes |
| FOXC2 | ENSG00000176692 | ENSG00000176692 | protein_coding | mCh_JDV-1-3-5 | F1F2_JDV-2-4-6 | 55,51578459 | 229,3302303 | 1,929556038 | 1,58E-08 | 0,000000528 | yes |
| ANXA5 | ENSG00000164111 | ENSG00000164111 | protein_coding | mCh_JDV-1-3-5 | F1F2_JDV-2-4-6 | 3641,390088 | 14691,57796 | 1,927099751 | 8,57E-11 | 4,4E-09 | yes |
| CCDC74A | ENSG00000163040 | ENSG00000163040 | protein_coding | mCh_JDV-1-3-5 | F1F2_JDV-2-4-6 | 3,879634947 | 20,34257076 | 1,924424673 | 0,000866847 | 0,006679227 | yes |
| MAOB | ENSG00000069535 | ENSG00000069535 | protein_coding | mCh_JDV-1-3-5 | F1F2_JDV-2-4-6 | 33,48474533 | 145,9365748 | 1,919996544 | 0,00000869 | 0,000149906 | yes |
| CCDC68 | ENSG00000166510 | ENSG00000166510 | protein_coding | mCh_JDV-1-3-5 | F1F2_JDV-2-4-6 | 117,8875316 | 473,0728782 | 1,919487618 | 1,03E-10 | 5,25E-09 | yes |
| KANK2 | ENSG00000197256 | ENSG00000197256 | protein_coding | mCh_JDV-1-3-5 | F1F2_JDV-2-4-6 | 847,3697141 | 3420,004976 | 1,916963059 | 0,000000001 | 4,26E-08 | yes |
| DNAJB4 | ENSG00000162616 | ENSG00000162616 | protein_coding | mCh_JDV-1-3-5 | F1F2_JDV-2-4-6 | 430,2330753 | 1663,310511 | 1,912332819 | 1,19E-20 | 1,88E-18 | yes |
| EHD3 | ENSG00000013016 | ENSG00000013016 | protein_coding | mCh_JDV-1-3-5 | F1F2_JDV-2-4-6 | 21,01530185 | 89,35138385 | 1,911506735 | 0,00000326 | 0,0000639 | yes |
| GYS2 | ENSG00000111713 | ENSG00000111713 | protein_coding | mCh_JDV-1-3-5 | F1F2_JDV-2-4-6 | 14,04266568 | 72,33798218 | 1,9111441 | 0,000921284 | 0,007001021 | yes |
| C12orf39 | ENSG00000134548 | ENSG00000134548 | protein_coding | mCh_JDV-1-3-5 | F1F2_JDV-2-4-6 | 166,8374414 | 806,8695546 | 1,90851256 | 0,000408293 | 0,003689031 | yes |
| PTN | ENSG00000105894 | ENSG00000105894 | protein_coding | mCh_JDV-1-3-5 | F1F2_JDV-2-4-6 | 24,36274694 | 117,0441099 | 1,903281807 | 0,000407039 | 0,00367939 | yes |
| PPM1H | ENSG00000111110 | ENSG00000111110 | protein_coding | mCh_JDV-1-3-5 | F1F2_JDV-2-4-6 | 43,44459692 | 176,7489552 | 1,89804742 | 0,000000103 | 0,00000299 | yes |
| SLC4A8 | ENSG00000050438 | ENSG00000050438 | protein_coding | mCh_JDV-1-3-5 | F1F2_JDV-2-4-6 | 13,60575415 | 66,56699338 | 1,896966498 | 0,000614988 | 0,005101114 | yes |
| ABCD2 | ENSG00000173208 | ENSG00000173208 | protein_coding | mCh_JDV-1-3-5 | F1F2_JDV-2-4-6 | 33,61813587 | 148,2399592 | 1,888224511 | 0,0000672 | 0,000841369 | yes |
| TMEM71 | ENSG00000165071 | ENSG00000165071 | protein_coding | mCh_JDV-1-3-5 | F1F2_JDV-2-4-6 | 9,759494013 | 48,2159789 | 1,871892847 | 0,001093811 | 0,008044792 | yes |
| ANGPT1 | ENSG00000154188 | ENSG00000154188 | protein_coding | mCh_JDV-1-3-5 | F1F2_JDV-2-4-6 | 310,6742077 | 1293,605714 | 1,866176797 | 0,0000129 | 0,000210249 | yes |
| SNRPN | ENSG00000128739 | ENSG00000128739 | protein_coding | mCh_JDV-1-3-5 | F1F2_JDV-2-4-6 | 400,3996673 | 1509,722548 | 1,863256094 | 5,37E-15 | 4,69E-13 | yes |
| TWIST2 | ENSG00000233608 | ENSG00000233608 | protein_coding | mCh_JDV-1-3-5 | F1F2_JDV-2-4-6 | 14,28962753 | 62,96547759 | 1,855691305 | 0,000205265 | 0,002115145 | yes |
| VSTM2L | ENSG00000132821 | ENSG00000132821 | protein_coding | mCh_JDV-1-3-5 | F1F2_JDV-2-4-6 | 243,808927 | 974,4969129 | 1,845214705 | 0,00000257 | 0,0000523 | yes |
| KCNJ15 | ENSG00000157551 | ENSG00000157551 | protein_coding | mCh_JDV-1-3-5 | F1F2_JDV-2-4-6 | 167,1937942 | 670,7185051 | 1,839029444 | 0,0000055 | 0,000100655 | yes |
| SYBU | ENSG00000147642 | ENSG00000147642 | protein_coding | mCh_JDV-1-3-5 | F1F2_JDV-2-4-6 | 385,3451729 | 1492,502731 | 1,832607385 | 0,000000245 | 0,00000645 | yes |
| MYPN | ENSG00000138347 | ENSG00000138347 | protein_coding | mCh_JDV-1-3-5 | F1F2_JDV-2-4-6 | 68,9721515 | 279,5209828 | 1,826920516 | 0,0000238 | 0,00035378 | yes |
| ZNF528 | ENSG00000167555 | ENSG00000167555 | protein_coding | mCh_JDV-1-3-5 | F1F2_JDV-2-4-6 | 67,04138988 | 249,9933731 | 1,819777628 | 5,58E-10 | 2,52E-08 | yes |
| VIM | ENSG00000026025 | ENSG00000026025 | protein_coding | mCh_JDV-1-3-5 | F1F2_JDV-2-4-6 | 8336,985934 | 30572,10484 | 1,819259526 | 3,03E-13 | 2,12E-11 | yes |
| FYN | ENSG00000010810 | ENSG00000010810 | protein_coding | mCh_JDV-1-3-5 | F1F2_JDV-2-4-6 | 34,6288415 | 137,5570203 | 1,816789924 | 0,0000127 | 0,000207815 | yes |
| STARD13 | ENSG00000133121 | ENSG00000133121 | protein_coding | mCh_JDV-1-3-5 | F1F2_JDV-2-4-6 | 523,8376895 | 1916,628161 | 1,814892464 | 5,84E-13 | 3,99E-11 | yes |
| BFSP1 | ENSG00000125864 | ENSG00000125864 | protein_coding | mCh_JDV-1-3-5 | F1F2_JDV-2-4-6 | 61,7580518 | 229,4143921 | 1,808798786 | 2,9E-09 | 0,000000113 | yes |
| SUSD5 | ENSG00000173705 | ENSG00000173705 | protein_coding | mCh_JDV-1-3-5 | F1F2_JDV-2-4-6 | 270,751231 | 992,0729321 | 1,807302159 | 3,03E-11 | 1,67E-09 | yes |
| TCEAL3 | ENSG00000196507 | ENSG00000196507 | protein_coding | mCh_JDV-1-3-5 | F1F2_JDV-2-4-6 | 81,83743228 | 302,2278049 | 1,806125554 | 0,000000001 | 4,26E-08 | yes |
| RASSF8 | ENSG00000123094 | ENSG00000123094 | protein_coding | mCh_JDV-1-3-5 | F1F2_JDV-2-4-6 | 578,0727809 | 2093,670156 | 1,80331639 | 2,48E-13 | 1,75E-11 | yes |
| IFI44L | ENSG00000137959 | ENSG00000137959 | protein_coding | mCh_JDV-1-3-5 | F1F2_JDV-2-4-6 | 566,2915674 | 2138,801297 | 1,78504016 | 0,00000171 | 0,0000367 | yes |
| NQO1 | ENSG00000181019 | ENSG00000181019 | protein_coding | mCh_JDV-1-3-5 | F1F2_JDV-2-4-6 | 5878,596709 | 20759,2157 | 1,782079393 | 3,07E-17 | 3,5E-15 | yes |
| FOXO1 | ENSG00000150907 | ENSG00000150907 | protein_coding | mCh_JDV-1-3-5 | F1F2_JDV-2-4-6 | 1779,66286 | 6390,630918 | 1,770505932 | 8,73E-10 | 3,78E-08 | yes |
| MRC2 | ENSG00000011028 | ENSG00000011028 | protein_coding | mCh_JDV-1-3-5 | F1F2_JDV-2-4-6 | 651,974327 | 2392,13913 | 1,767600538 | 0,000000251 | 0,00000659 | yes |
| HLX | ENSG00000136630 | ENSG00000136630 | protein_coding | mCh_JDV-1-3-5 | F1F2_JDV-2-4-6 | 6,828575092 | 30,3294725 | 1,765240988 | 0,001750882 | 0,011633047 | yes |
| C16orf45 | ENSG00000166780 | ENSG00000166780 | protein_coding | mCh_JDV-1-3-5 | F1F2_JDV-2-4-6 | 62,32369344 | 229,4730643 | 1,761794391 | 0,000000783 | 0,0000185 | yes |
| TRIM16L | ENSG00000108448 | ENSG00000108448 | protein_coding | mCh_JDV-1-3-5 | F1F2_JDV-2-4-6 | 267,9933819 | 944,844493 | 1,758193893 | 2,03E-11 | 1,15E-09 | yes |
| CCND1 | ENSG00000110092 | ENSG00000110092 | protein_coding | mCh_JDV-1-3-5 | F1F2_JDV-2-4-6 | 1985,828208 | 7069,382726 | 1,754499101 | 3,22E-09 | 0,000000124 | yes |
| S1PR3 | ENSG00000213694 | ENSG00000213694 | protein_coding | mCh_JDV-1-3-5 | F1F2_JDV-2-4-6 | 885,2035608 | 3239,47157 | 1,749286521 | 0,00000155 | 0,000034 | yes |
| CACNA2D3 | ENSG00000157445 | ENSG00000157445 | protein_coding | mCh_JDV-1-3-5 | F1F2_JDV-2-4-6 | 117,5633896 | 461,5660113 | 1,748076081 | 0,000185345 | 0,001942334 | yes |
| PLAG1 | ENSG00000181690 | ENSG00000181690 | protein_coding | mCh_JDV-1-3-5 | F1F2_JDV-2-4-6 | 56,16331204 | 215,9628161 | 1,746621825 | 0,0000803 | 0,00097039 | yes |
| PLEKHC1 | ENSG00000073712 | ENSG00000073712 | protein_coding | mCh_JDV-1-3-5 | F1F2_JDV-2-4-6 | 965,4712042 | 3359,560165 | 1,744361647 | 5,12E-12 | 3,08E-10 | yes |
| AKR1C3 | ENSG00000196139 | ENSG00000196139 | protein_coding | mCh_JDV-1-3-5 | F1F2_JDV-2-4-6 | 1860,571916 | 6782,96623 | 1,739404987 | 0,00000269 | 0,0000542 | yes |
| C12orf60 | ENSG00000182993 | ENSG00000182993 | protein_coding | mCh_JDV-1-3-5 | F1F2_JDV-2-4-6 | 24,87179771 | 91,89600408 | 1,731170325 | 0,0000192 | 0,000295699 | yes |
| CYP19A1 | ENSG00000137869 | ENSG00000137869 | protein_coding | mCh_JDV-1-3-5 | F1F2_JDV-2-4-6 | 12,73165021 | 50,68628793 | 1,728322446 | 0,00048711 | 0,004241897 | yes |

| | | | | | | | | | | | |
|---|---|---|---|---|---|---|---|---|---|---|---|
| CDH2 | ENSG00000170558 | ENSG00000170558 | protein_coding | mCh_JDV-1-3-5 | F1F2_JDV-2-4-6 | 624,6062848 | 2182,004611 | 1,717409862 | 5,53E-08 | 0,00000169 | yes |
| RARRES3 | ENSG00000133321 | ENSG00000133321 | protein_coding | mCh_JDV-1-3-5 | F1F2_JDV-2-4-6 | 16,62402547 | 62,02037006 | 1,706290615 | 0,00011492 | 0,001305541 | yes |
| PEPD | ENSG00000124299 | ENSG00000124299 | protein_coding | mCh_JDV-1-3-5 | F1F2_JDV-2-4-6 | 129,146691 | 442,381559 | 1,699939828 | 1,35E-08 | 0,00000046 | yes |
| BIN1 | ENSG00000136717 | ENSG00000136717 | protein_coding | mCh_JDV-1-3-5 | F1F2_JDV-2-4-6 | 850,1175993 | 2872,199257 | 1,696271896 | 2,37E-10 | 1,14E-08 | yes |
| CDKN2C | ENSG00000123080 | ENSG00000123080 | protein_coding | mCh_JDV-1-3-5 | F1F2_JDV-2-4-6 | 203,8441286 | 790,230372 | 1,696168596 | 0,000674127 | 0,00548559 | yes |
| TBX3 | ENSG00000135111 | ENSG00000135111 | protein_coding | mCh_JDV-1-3-5 | F1F2_JDV-2-4-6 | 493,5286373 | 1640,86789 | 1,690357691 | 1,42E-13 | 1,03E-11 | yes |
| TGFB2 | ENSG00000092969 | ENSG00000092969 | protein_coding | mCh_JDV-1-3-5 | F1F2_JDV-2-4-6 | 155,9560076 | 538,7647903 | 1,686300437 | 0,000000786 | 0,0000185 | yes |
| ABI3BP | ENSG00000154175 | ENSG00000154175 | protein_coding | mCh_JDV-1-3-5 | F1F2_JDV-2-4-6 | 11,56181079 | 49,7472773 | 1,68529435 | 0,004007598 | 0,022170434 | yes |
| UBE2L6 | ENSG00000156587 | ENSG00000156587 | protein_coding | mCh_JDV-1-3-5 | F1F2_JDV-2-4-6 | 317,881378 | 1064,25185 | 1,677521936 | 2,19E-09 | 8,75E-08 | yes |
| IFI35 | ENSG00000068079 | ENSG00000068079 | protein_coding | mCh_JDV-1-3-5 | F1F2_JDV-2-4-6 | 75,24547462 | 255,4625469 | 1,660029173 | 0,00000173 | 0,0000372 | yes |
| JDP2 | ENSG00000140044 | ENSG00000140044 | protein_coding | mCh_JDV-1-3-5 | F1F2_JDV-2-4-6 | 161,9320146 | 532,2112988 | 1,659332701 | 3,3E-10 | 1,56E-08 | yes |
| DPYD | ENSG00000188641 | ENSG00000188641 | protein_coding | mCh_JDV-1-3-5 | F1F2_JDV-2-4-6 | 1146,964934 | 3841,566984 | 1,657302536 | 0,000000222 | 0,00000589 | yes |
| FLVCR2 | ENSG00000119686 | ENSG00000119686 | protein_coding | mCh_JDV-1-3-5 | F1F2_JDV-2-4-6 | 204,2377102 | 685,4789726 | 1,656647774 | 0,000000354 | 0,00000894 | yes |
| CHN1 | ENSG00000128656 | ENSG00000128656 | protein_coding | mCh_JDV-1-3-5 | F1F2_JDV-2-4-6 | 321,5215244 | 1101,948413 | 1,656120445 | 0,00000814 | 0,000141163 | yes |
| MAP7D2 | ENSG00000184368 | ENSG00000184368 | protein_coding | mCh_JDV-1-3-5 | F1F2_JDV-2-4-6 | 18,15641347 | 65,94607711 | 1,65581316 | 0,000343973 | 0,003219931 | yes |
| WNT5A | ENSG00000114251 | ENSG00000114251 | protein_coding | mCh_JDV-1-3-5 | F1F2_JDV-2-4-6 | 270,3655886 | 923,9714526 | 1,647203746 | 0,000013 | 0,00021185 | yes |
| PHEX | ENSG00000102174 | ENSG00000102174 | protein_coding | mCh_JDV-1-3-5 | F1F2_JDV-2-4-6 | 60,90676084 | 204,3619528 | 1,644793939 | 0,00000176 | 0,0000377 | yes |
| RGS17 | ENSG00000091844 | ENSG00000091844 | protein_coding | mCh_JDV-1-3-5 | F1F2_JDV-2-4-6 | 26,24740695 | 90,50165155 | 1,634400585 | 0,0000626 | 0,000789795 | yes |
| BEST2 | ENSG00000039987 | ENSG00000039987 | protein_coding | mCh_JDV-1-3-5 | F1F2_JDV-2-4-6 | 15,64188028 | 63,94963498 | 1,631754451 | 0,005139863 | 0,0269103 | yes |
| C8orf85 | ENSG00000205002 | ENSG00000205002 | protein_coding | mCh_JDV-1-3-5 | F1F2_JDV-2-4-6 | 43,12324038 | 145,5341586 | 1,625978041 | 0,0000234 | 0,000348746 | yes |
| BNC2 | ENSG00000173068 | ENSG00000173068 | protein_coding | mCh_JDV-1-3-5 | F1F2_JDV-2-4-6 | 16,80663357 | 61,49314372 | 1,616182256 | 0,001369423 | 0,009495021 | yes |
| PRKCDBP | ENSG00000170955 | ENSG00000170955 | protein_coding | mCh_JDV-1-3-5 | F1F2_JDV-2-4-6 | 96,29157884 | 378,6869475 | 1,612681769 | 0,004644561 | 0,024871405 | yes |
| DEAF1 | ENSG00000177030 | ENSG00000177030 | protein_coding | mCh_JDV-1-3-5 | F1F2_JDV-2-4-6 | 325,8210097 | 1046,022722 | 1,611685683 | 5,45E-08 | 0,00000168 | yes |
| FGF2 | ENSG00000138685 | ENSG00000138685 | protein_coding | mCh_JDV-1-3-5 | F1F2_JDV-2-4-6 | 451,2144179 | 1423,767643 | 1,610483454 | 5,1E-11 | 2,7E-09 | yes |
| TIMP1 | ENSG00000102265 | ENSG00000102265 | protein_coding | mCh_JDV-1-3-5 | F1F2_JDV-2-4-6 | 15510,69078 | 49824,9189 | 1,607851649 | 0,000000141 | 0,00000396 | yes |
| GBE1 | ENSG00000114480 | ENSG00000114480 | protein_coding | mCh_JDV-1-3-5 | F1F2_JDV-2-4-6 | 1523,952972 | 4798,498229 | 1,607391514 | 6,24E-11 | 3,27E-09 | yes |
| RGL1 | ENSG00000143344 | ENSG00000143344 | protein_coding | mCh_JDV-1-3-5 | F1F2_JDV-2-4-6 | 200,70734 | 640,0406579 | 1,600632537 | 8,73E-08 | 0,00000258 | yes |
| SKAP2 | ENSG00000005020 | ENSG00000005020 | protein_coding | mCh_JDV-1-3-5 | F1F2_JDV-2-4-6 | 523,5042678 | 1676,657526 | 1,599685608 | 0,000000333 | 0,0000085 | yes |
| ISG15 | ENSG00000187608 | ENSG00000187608 | protein_coding | mCh_JDV-1-3-5 | F1F2_JDV-2-4-6 | 9,011375586 | 31,40359626 | 1,593364057 | 0,000688159 | 0,005572195 | yes |
| UGP2 | ENSG00000169764 | ENSG00000169764 | protein_coding | mCh_JDV-1-3-5 | F1F2_JDV-2-4-6 | 2027,413331 | 6495,663636 | 1,589209447 | 0,00000181 | 0,0000383 | yes |
| ADPRHL1 | ENSG00000153531 | ENSG00000153531 | protein_coding | mCh_JDV-1-3-5 | F1F2_JDV-2-4-6 | 21,83040993 | 73,3756034 | 1,584996541 | 0,000218849 | 0,002220337 | yes |
| FAM127C | ENSG00000212747 | ENSG00000212747 | protein_coding | mCh_JDV-1-3-5 | F1F2_JDV-2-4-6 | 96,93683262 | 300,372807 | 1,580930695 | 7,74E-10 | 3,39E-08 | yes |
| POLR2L | ENSG00000177700 | ENSG00000177700 | protein_coding | mCh_JDV-1-3-5 | F1F2_JDV-2-4-6 | 542,5928118 | 1735,68591 | 1,570294662 | 0,0000133 | 0,000215295 | yes |
| GLIS2 | ENSG00000126603 | ENSG00000126603 | protein_coding | mCh_JDV-1-3-5 | F1F2_JDV-2-4-6 | 133,1345212 | 421,3256487 | 1,570248943 | 0,00000294 | 0,0000585 | yes |
| NRIP3 | ENSG00000175352 | ENSG00000175352 | protein_coding | mCh_JDV-1-3-5 | F1F2_JDV-2-4-6 | 51,27344712 | 165,0966348 | 1,569043486 | 0,0000302 | 0,000433525 | yes |
| PDE6G | ENSG00000185527 | ENSG00000185527 | protein_coding | mCh_JDV-1-3-5 | F1F2_JDV-2-4-6 | 4,76666486 | 18,89620904 | 1,556907797 | 0,009485229 | 0,042577801 | yes |
| ARPC5 | ENSG00000162704 | ENSG00000162704 | protein_coding | mCh_JDV-1-3-5 | F1F2_JDV-2-4-6 | 2301,348109 | 7220,727221 | 1,552351752 | 0,00000777 | 0,000136092 | yes |
| TREM1 | ENSG00000124731 | ENSG00000124731 | protein_coding | mCh_JDV-1-3-5 | F1F2_JDV-2-4-6 | 68,85157823 | 218,6431409 | 1,549183325 | 0,0000409 | 0,00055738 | yes |
| AKR1C1 | ENSG00000187134 | ENSG00000187134 | protein_coding | mCh_JDV-1-3-5 | F1F2_JDV-2-4-6 | 3243,701675 | 11250,93847 | 1,548591486 | 0,002216996 | 0,014033299 | yes |
| RPS27L | ENSG00000185088 | ENSG00000185088 | protein_coding | mCh_JDV-1-3-5 | F1F2_JDV-2-4-6 | 1080,887546 | 3250,74335 | 1,534737181 | 8,55E-09 | 0,000000301 | yes |
| UBB | ENSG00000170315 | ENSG00000170315 | protein_coding | mCh_JDV-1-3-5 | F1F2_JDV-2-4-6 | 3148,362821 | 9815,343861 | 1,533674089 | 0,0000245 | 0,000361858 | yes |
| CMTM3 | ENSG00000140931 | ENSG00000140931 | protein_coding | mCh_JDV-1-3-5 | F1F2_JDV-2-4-6 | 46,6688011 | 145,5151928 | 1,531742714 | 0,0000262 | 0,000382898 | yes |
| CR381653.2 | ENSG00000236557 | ENSG00000236557 | protein_coding | mCh_JDV-1-3-5 | F1F2_JDV-2-4-6 | 70,98750269 | 216,3753799 | 1,524852711 | 0,00000289 | 0,0000577 | yes |
| BCL2L1 | ENSG00000171552 | ENSG00000171552 | protein_coding | mCh_JDV-1-3-5 | F1F2_JDV-2-4-6 | 1437,07371 | 4332,104871 | 1,524390187 | 0,000000288 | 0,00000748 | yes |
| NAV3 | ENSG00000067798 | ENSG00000067798 | protein_coding | mCh_JDV-1-3-5 | F1F2_JDV-2-4-6 | 66,60154807 | 235,9272004 | 1,52388799 | 0,00526998 | 0,027424185 | yes |
| FTH1 | ENSG00000167996 | ENSG00000167996 | protein_coding | mCh_JDV-1-3-5 | F1F2_JDV-2-4-6 | 26158,53662 | 80842,24846 | 1,522459017 | 0,0000267 | 0,000388855 | yes |
| EML1 | ENSG00000066629 | ENSG00000066629 | protein_coding | mCh_JDV-1-3-5 | F1F2_JDV-2-4-6 | 223,056715 | 692,5915808 | 1,521961573 | 0,000046 | 0,000614242 | yes |

| | | | | | | | | | | | |
|---|---|---|---|---|---|---|---|---|---|---|---|
| MAP1LC3B | ENSG00000140941 | ENSG00000140941 | protein_coding | mCh_JDV-1-3-5 | F1F2_JDV-2-4-6 | 867,7866565 | 2574,267068 | 1,520636272 | 2,19E-09 | 8,75E-08 | yes |
| CARD6 | ENSG00000132357 | ENSG00000132357 | protein_coding | mCh_JDV-1-3-5 | F1F2_JDV-2-4-6 | 635,8574683 | 1907,935711 | 1,509469535 | 0,00000157 | 0,0000344 | yes |
| MAPK8IP1 | ENSG00000121653 | ENSG00000121653 | protein_coding | mCh_JDV-1-3-5 | F1F2_JDV-2-4-6 | 16,0996211 | 50,58044361 | 1,508742436 | 0,000276168 | 0,002685862 | yes |
| SDSL | ENSG00000139410 | ENSG00000139410 | protein_coding | mCh_JDV-1-3-5 | F1F2_JDV-2-4-6 | 26,1394744 | 84,09380361 | 1,503595987 | 0,001020653 | 0,007588748 | yes |
| RAP2B | ENSG00000181467 | ENSG00000181467 | protein_coding | mCh_JDV-1-3-5 | F1F2_JDV-2-4-6 | 1405,894449 | 483,5140292 | -1,501423426 | 6,84E-11 | 3,54E-09 | yes |
| C16orf67 | ENSG00000131797 | ENSG00000131797 | protein_coding | mCh_JDV-1-3-5 | F1F2_JDV-2-4-6 | 442,4059497 | 147,6271164 | -1,502271944 | 0,00000365 | 0,0000701 | yes |
| FRRS1 | ENSG00000156869 | ENSG00000156869 | protein_coding | mCh_JDV-1-3-5 | F1F2_JDV-2-4-6 | 655,0381656 | 221,593662 | -1,504659789 | 0,000000085 | 0,00000252 | yes |
| STRA6 | ENSG00000137868 | ENSG00000137868 | protein_coding | mCh_JDV-1-3-5 | F1F2_JDV-2-4-6 | 280,504167 | 90,76911241 | -1,505557084 | 0,000105345 | 0,001214951 | yes |
| LGR4 | ENSG00000205213 | ENSG00000205213 | protein_coding | mCh_JDV-1-3-5 | F1F2_JDV-2-4-6 | 3230,930013 | 1108,163469 | -1,505752097 | 4,4E-11 | 2,36E-09 | yes |
| PPFIA4 | ENSG00000143847 | ENSG00000143847 | protein_coding | mCh_JDV-1-3-5 | F1F2_JDV-2-4-6 | 416,2194143 | 117,5151324 | -1,50739285 | 0,006845981 | 0,033492735 | yes |
| COL1A1 | ENSG00000108821 | ENSG00000108821 | protein_coding | mCh_JDV-1-3-5 | F1F2_JDV-2-4-6 | 267,6267255 | 87,69188354 | -1,508099689 | 0,0000249 | 0,000367597 | yes |
| SGPP2 | ENSG00000163082 | ENSG00000163082 | protein_coding | mCh_JDV-1-3-5 | F1F2_JDV-2-4-6 | 7024,351359 | 2205,108108 | -1,50998015 | 0,00052117 | 0,004461047 | yes |
| PARD6B | ENSG00000124171 | ENSG00000124171 | protein_coding | mCh_JDV-1-3-5 | F1F2_JDV-2-4-6 | 115,4633219 | 38,12085952 | -1,513465047 | 0,00000455 | 0,0000854 | yes |
| ICOSLG | ENSG00000160223 | ENSG00000160223 | protein_coding | mCh_JDV-1-3-5 | F1F2_JDV-2-4-6 | 334,8259674 | 109,4858805 | -1,516614621 | 0,0000133 | 0,000214828 | yes |
| DEF6 | ENSG00000023892 | ENSG00000023892 | protein_coding | mCh_JDV-1-3-5 | F1F2_JDV-2-4-6 | 487,5722821 | 164,8266568 | -1,517394118 | 1,81E-09 | 7,29E-08 | yes |
| TMEM171 | ENSG00000157111 | ENSG00000157111 | protein_coding | mCh_JDV-1-3-5 | F1F2_JDV-2-4-6 | 105,7504187 | 32,57303859 | -1,518809471 | 0,000804145 | 0,006309422 | yes |
| CYP3A5 | ENSG00000106258 | ENSG00000106258 | protein_coding | mCh_JDV-1-3-5 | F1F2_JDV-2-4-6 | 308,3348141 | 90,23912066 | -1,519074669 | 0,003205429 | 0,018746595 | yes |
| RNF144B | ENSG00000137393 | ENSG00000137393 | protein_coding | mCh_JDV-1-3-5 | F1F2_JDV-2-4-6 | 1007,158807 | 338,1289152 | -1,520664945 | 1,45E-08 | 0,00000049 | yes |
| SLC45A3 | ENSG00000158715 | ENSG00000158715 | protein_coding | mCh_JDV-1-3-5 | F1F2_JDV-2-4-6 | 708,0606223 | 227,9503846 | -1,520665399 | 0,0000526 | 0,000688218 | yes |
| PRRG4 | ENSG00000135378 | ENSG00000135378 | protein_coding | mCh_JDV-1-3-5 | F1F2_JDV-2-4-6 | 378,1374729 | 126,4921321 | -1,527001214 | 8,33E-09 | 0,000000294 | yes |
| CSTA | ENSG00000121552 | ENSG00000121552 | protein_coding | mCh_JDV-1-3-5 | F1F2_JDV-2-4-6 | 1688,514458 | 560,2334779 | -1,527818201 | 0,000000125 | 0,00000356 | yes |
| TMC7 | ENSG00000170537 | ENSG00000170537 | protein_coding | mCh_JDV-1-3-5 | F1F2_JDV-2-4-6 | 202,1550432 | 67,0080813 | -1,528712117 | 0,000000151 | 0,00000422 | yes |
| LAMP3 | ENSG00000078081 | ENSG00000078081 | protein_coding | mCh_JDV-1-3-5 | F1F2_JDV-2-4-6 | 131,9202289 | 38,8973001 | -1,531103568 | 0,002021048 | 0,013011857 | yes |
| PLEKHG5 | ENSG00000171680 | ENSG00000171680 | protein_coding | mCh_JDV-1-3-5 | F1F2_JDV-2-4-6 | 1910,315149 | 633,4505595 | -1,535686047 | 1,91E-08 | 0,000000635 | yes |
| C20orf96 | ENSG00000196476 | ENSG00000196476 | protein_coding | mCh_JDV-1-3-5 | F1F2_JDV-2-4-6 | 525,0658134 | 165,1910648 | -1,538117356 | 0,0000989 | 0,00115389 | yes |
| CXCL16 | ENSG00000161921 | ENSG00000161921 | protein_coding | mCh_JDV-1-3-5 | F1F2_JDV-2-4-6 | 1169,799804 | 377,5924915 | -1,538365023 | 0,00000661 | 0,000118166 | yes |
| DSE | ENSG00000111817 | ENSG00000111817 | protein_coding | mCh_JDV-1-3-5 | F1F2_JDV-2-4-6 | 6613,121466 | 2201,278434 | -1,544246849 | 9,68E-11 | 4,93E-09 | yes |
| CREG1 | ENSG00000143162 | ENSG00000143162 | protein_coding | mCh_JDV-1-3-5 | F1F2_JDV-2-4-6 | 16781,37236 | 5455,167971 | -1,546283111 | 0,000000574 | 0,0000138 | yes |
| REEP4 | ENSG00000168476 | ENSG00000168476 | protein_coding | mCh_JDV-1-3-5 | F1F2_JDV-2-4-6 | 1740,868006 | 561,4507581 | -1,546377971 | 0,00000271 | 0,0000544 | yes |
| SEMA4B | ENSG00000185033 | ENSG00000185033 | protein_coding | mCh_JDV-1-3-5 | F1F2_JDV-2-4-6 | 5052,609939 | 1660,365224 | -1,546860457 | 2,21E-08 | 0,000000729 | yes |
| CD7 | ENSG00000173762 | ENSG00000173762 | protein_coding | mCh_JDV-1-3-5 | F1F2_JDV-2-4-6 | 49,99606911 | 13,47460413 | -1,549967388 | 0,006051637 | 0,030549232 | yes |
| CPA4 | ENSG00000128510 | ENSG00000128510 | protein_coding | mCh_JDV-1-3-5 | F1F2_JDV-2-4-6 | 223,7125036 | 67,38138181 | -1,550485988 | 0,000569181 | 0,004775363 | yes |
| MARCH9 | ENSG00000139266 | ENSG00000139266 | protein_coding | mCh_JDV-1-3-5 | F1F2_JDV-2-4-6 | 388,319838 | 117,2743676 | -1,554698547 | 0,000430347 | 0,003849518 | yes |
| SALL4 | ENSG00000101115 | ENSG00000101115 | protein_coding | mCh_JDV-1-3-5 | F1F2_JDV-2-4-6 | 20,12323105 | 5,398040911 | -1,555770667 | 0,005698152 | 0,029116942 | yes |
| SILV | ENSG00000185664 | ENSG00000185664 | protein_coding | mCh_JDV-1-3-5 | F1F2_JDV-2-4-6 | 33,21483918 | 9,078788564 | -1,56200163 | 0,004079196 | 0,022459478 | yes |
| B3GNT4 | ENSG00000176383 | ENSG00000176383 | protein_coding | mCh_JDV-1-3-5 | F1F2_JDV-2-4-6 | 80,86064498 | 25,33882185 | -1,56334592 | 0,0000191 | 0,000294883 | yes |
| ZDHHC23 | ENSG00000184307 | ENSG00000184307 | protein_coding | mCh_JDV-1-3-5 | F1F2_JDV-2-4-6 | 1406,397118 | 458,8974996 | -1,564719804 | 1,25E-09 | 5,17E-08 | yes |
| DUSP9 | ENSG00000130829 | ENSG00000130829 | protein_coding | mCh_JDV-1-3-5 | F1F2_JDV-2-4-6 | 24,07228325 | 6,748552792 | -1,569206668 | 0,002290914 | 0,014401597 | yes |
| SLC24A6 | ENSG00000089060 | ENSG00000089060 | protein_coding | mCh_JDV-1-3-5 | F1F2_JDV-2-4-6 | 2062,777378 | 673,2864365 | -1,571358353 | 5,45E-11 | 2,87E-09 | yes |
| GJB5 | ENSG00000189280 | ENSG00000189280 | protein_coding | mCh_JDV-1-3-5 | F1F2_JDV-2-4-6 | 2167,469495 | 694,3894146 | -1,572153554 | 0,000000128 | 0,00000364 | yes |
| IL27RA | ENSG00000104998 | ENSG00000104998 | protein_coding | mCh_JDV-1-3-5 | F1F2_JDV-2-4-6 | 2334,286522 | 736,9516748 | -1,574497418 | 0,00000201 | 0,0000421 | yes |
| ABCA5 | ENSG00000154265 | ENSG00000154265 | protein_coding | mCh_JDV-1-3-5 | F1F2_JDV-2-4-6 | 581,2340759 | 181,6890071 | -1,576467101 | 0,00000699 | 0,000124221 | yes |
| SUN1 | ENSG00000164828 | ENSG00000164828 | protein_coding | mCh_JDV-1-3-5 | F1F2_JDV-2-4-6 | 11024,85214 | 3613,624413 | -1,576829526 | 2,61E-14 | 2,1E-12 | yes |
| MDK | ENSG00000110492 | ENSG00000110492 | protein_coding | mCh_JDV-1-3-5 | F1F2_JDV-2-4-6 | 36,52890945 | 10,46112125 | -1,577002609 | 0,001184285 | 0,008532125 | yes |
| RAPGEF3 | ENSG00000079337 | ENSG00000079337 | protein_coding | mCh_JDV-1-3-5 | F1F2_JDV-2-4-6 | 286,3096266 | 90,19986203 | -1,581101952 | 0,00000113 | 0,0000256 | yes |
| GPR126 | ENSG00000112414 | ENSG00000112414 | protein_coding | mCh_JDV-1-3-5 | F1F2_JDV-2-4-6 | 7637,142165 | 2465,099664 | -1,581802424 | 4,04E-10 | 1,88E-08 | yes |
| P2RX7 | ENSG00000089041 | ENSG00000089041 | protein_coding | mCh_JDV-1-3-5 | F1F2_JDV-2-4-6 | 45,20076684 | 11,78580018 | -1,583478654 | 0,005292512 | 0,027480084 | yes |

| | | | | | | | | | | | |
|---|---|---|---|---|---|---|---|---|---|---|---|
| C1R | ENSG00000159403 | ENSG00000159403 | protein_coding | mCh_JDV-1-3-5 | F1F2_JDV-2-4-6 | 5208,276979 | 1390,791722 | -1,584865414 | 0,003968043 | 0,022000952 | yes |
| TMEM63B | ENSG00000137216 | ENSG00000137216 | protein_coding | mCh_JDV-1-3-5 | F1F2_JDV-2-4-6 | 3750,361198 | 1190,251576 | -1,585269375 | 9,89E-08 | 0,00000289 | yes |
| CYP2J2 | ENSG00000134716 | ENSG00000134716 | protein_coding | mCh_JDV-1-3-5 | F1F2_JDV-2-4-6 | 63,26468743 | 16,98852688 | -1,593551735 | 0,00309967 | 0,018220373 | yes |
| SPTLC2 | ENSG00000100596 | ENSG00000100596 | protein_coding | mCh_JDV-1-3-5 | F1F2_JDV-2-4-6 | 3702,068406 | 1159,416098 | -1,59361652 | 0,000000486 | 0,0000119 | yes |
| KLHL17 | ENSG00000187961 | ENSG00000187961 | protein_coding | mCh_JDV-1-3-5 | F1F2_JDV-2-4-6 | 728,2318614 | 229,7810296 | -1,594182391 | 6,89E-08 | 0,00000209 | yes |
| MYB | ENSG00000118513 | ENSG00000118513 | protein_coding | mCh_JDV-1-3-5 | F1F2_JDV-2-4-6 | 114,7306749 | 32,97975373 | -1,594383363 | 0,000647794 | 0,005304113 | yes |
| CBWD6 | ENSG00000204790 | ENSG00000204790 | protein_coding | mCh_JDV-1-3-5 | F1F2_JDV-2-4-6 | 178,4265173 | 54,72077869 | -1,595283464 | 0,0000103 | 0,000173042 | yes |
| LRIG3 | ENSG00000139263 | ENSG00000139263 | protein_coding | mCh_JDV-1-3-5 | F1F2_JDV-2-4-6 | 2657,07238 | 810,6761261 | -1,598932867 | 0,000013 | 0,00021185 | yes |
| MYO1D | ENSG00000176658 | ENSG00000176658 | protein_coding | mCh_JDV-1-3-5 | F1F2_JDV-2-4-6 | 1474,879522 | 461,3950173 | -1,602299966 | 0,000000124 | 0,00000356 | yes |
| SULT1A3 | ENSG00000213599 | ENSG00000213599 | protein_coding | mCh_JDV-1-3-5 | F1F2_JDV-2-4-6 | 1161,70218 | 356,3045904 | -1,608420976 | 0,00000233 | 0,0000478 | yes |
| PCSK6 | ENSG00000140479 | ENSG00000140479 | protein_coding | mCh_JDV-1-3-5 | F1F2_JDV-2-4-6 | 1688,34685 | 533,8494531 | -1,611333694 | 1,4E-10 | 6,98E-09 | yes |
| SCART1 | ENSG00000214279 | ENSG00000214279 | protein_coding | mCh_JDV-1-3-5 | F1F2_JDV-2-4-6 | 67,04369967 | 18,52223341 | -1,6130569 | 0,001122352 | 0,008199691 | yes |
| ARTN | ENSG00000117407 | ENSG00000117407 | protein_coding | mCh_JDV-1-3-5 | F1F2_JDV-2-4-6 | 35,05299011 | 9,138567411 | -1,613453696 | 0,003212792 | 0,01876684 | yes |
| PAN2 | ENSG00000135473 | ENSG00000135473 | protein_coding | mCh_JDV-1-3-5 | F1F2_JDV-2-4-6 | 5895,883231 | 1709,402948 | -1,614864584 | 0,000196812 | 0,002046197 | yes |
| TNFRSF11A | ENSG00000141655 | ENSG00000141655 | protein_coding | mCh_JDV-1-3-5 | F1F2_JDV-2-4-6 | 327,1728459 | 94,07642036 | -1,617873938 | 0,000255336 | 0,002523053 | yes |
| IGSF3 | ENSG00000143061 | ENSG00000143061 | protein_coding | mCh_JDV-1-3-5 | F1F2_JDV-2-4-6 | 2458,695314 | 745,1010712 | -1,618615201 | 0,0000039 | 0,0000746 | yes |
| SCD | ENSG00000099194 | ENSG00000099194 | protein_coding | mCh_JDV-1-3-5 | F1F2_JDV-2-4-6 | 49932,86283 | 15132,81509 | -1,619651074 | 0,00000343 | 0,0000666 | yes |
| MRGPRX3 | ENSG00000179826 | ENSG00000179826 | protein_coding | mCh_JDV-1-3-5 | F1F2_JDV-2-4-6 | 24,11839837 | 6,115765188 | -1,622935711 | 0,003975807 | 0,022025437 | yes |
| ATP7B | ENSG00000123191 | ENSG00000123191 | protein_coding | mCh_JDV-1-3-5 | F1F2_JDV-2-4-6 | 649,8391723 | 199,9716983 | -1,623957576 | 0,000000103 | 0,00000299 | yes |
| ERMP1 | ENSG00000099219 | ENSG00000099219 | protein_coding | mCh_JDV-1-3-5 | F1F2_JDV-2-4-6 | 6123,475381 | 1921,914731 | -1,628383421 | 3,7E-12 | 2,28E-10 | yes |
| HPS4 | ENSG00000100099 | ENSG00000100099 | protein_coding | mCh_JDV-1-3-5 | F1F2_JDV-2-4-6 | 3299,688898 | 1019,794966 | -1,628393814 | 1,03E-08 | 0,000000357 | yes |
| SNN | ENSG00000184602 | ENSG00000184602 | protein_coding | mCh_JDV-1-3-5 | F1F2_JDV-2-4-6 | 492,1080226 | 152,8516741 | -1,629073687 | 1,28E-09 | 5,27E-08 | yes |
| ICAM1 | ENSG00000090339 | ENSG00000090339 | protein_coding | mCh_JDV-1-3-5 | F1F2_JDV-2-4-6 | 3142,754884 | 923,313273 | -1,62979271 | 0,0000359 | 0,000500102 | yes |
| RGS16 | ENSG00000143333 | ENSG00000143333 | protein_coding | mCh_JDV-1-3-5 | F1F2_JDV-2-4-6 | 136,8949325 | 40,05862482 | -1,631722607 | 0,0000398 | 0,000545016 | yes |
| C21orf7 | ENSG00000156265 | ENSG00000156265 | protein_coding | mCh_JDV-1-3-5 | F1F2_JDV-2-4-6 | 37,40796819 | 10,45339572 | -1,633206461 | 0,000392818 | 0,003578707 | yes |
| BIK | ENSG00000100290 | ENSG00000100290 | protein_coding | mCh_JDV-1-3-5 | F1F2_JDV-2-4-6 | 58,76019758 | 15,87021918 | -1,634081662 | 0,001100932 | 0,008082099 | yes |
| NCR3LG1 | ENSG00000188211 | ENSG00000188211 | protein_coding | mCh_JDV-1-3-5 | F1F2_JDV-2-4-6 | 2817,352784 | 854,1639967 | -1,634875923 | 0,0000004 | 0,00000999 | yes |
| STEAP2 | ENSG00000157214 | ENSG00000157214 | protein_coding | mCh_JDV-1-3-5 | F1F2_JDV-2-4-6 | 564,7137574 | 170,1776741 | -1,635235442 | 0,00000115 | 0,000026 | yes |
| RNF165 | ENSG00000141622 | ENSG00000141622 | protein_coding | mCh_JDV-1-3-5 | F1F2_JDV-2-4-6 | 509,4476023 | 154,6723989 | -1,636569456 | 0,000000232 | 0,00000615 | yes |
| MET | ENSG00000105976 | ENSG00000105976 | protein_coding | mCh_JDV-1-3-5 | F1F2_JDV-2-4-6 | 11770,26295 | 3613,210538 | -1,647017404 | 4,75E-10 | 2,19E-08 | yes |
| TACC2 | ENSG00000138162 | ENSG00000138162 | protein_coding | mCh_JDV-1-3-5 | F1F2_JDV-2-4-6 | 10223,58788 | 3041,03318 | -1,647935523 | 0,00000168 | 0,0000362 | yes |
| GFOD1 | ENSG00000145990 | ENSG00000145990 | protein_coding | mCh_JDV-1-3-5 | F1F2_JDV-2-4-6 | 912,4663568 | 281,0952889 | -1,652496379 | 5,76E-12 | 3,43E-10 | yes |
| GPRC5A | ENSG00000013588 | ENSG00000013588 | protein_coding | mCh_JDV-1-3-5 | F1F2_JDV-2-4-6 | 4151,528338 | 1135,984381 | -1,652754233 | 0,000457008 | 0,004036765 | yes |
| GP1BA | ENSG00000185245 | ENSG00000185245 | protein_coding | mCh_JDV-1-3-5 | F1F2_JDV-2-4-6 | 150,7968161 | 40,36562735 | -1,653293534 | 0,000841392 | 0,006521286 | yes |
| PRSS16 | ENSG00000112812 | ENSG00000112812 | protein_coding | mCh_JDV-1-3-5 | F1F2_JDV-2-4-6 | 462,9011998 | 140,7229396 | -1,654718989 | 2,27E-09 | 9,04E-08 | yes |
| GOLGA2B | ENSG00000120853 | ENSG00000120853 | protein_coding | mCh_JDV-1-3-5 | F1F2_JDV-2-4-6 | 133,9054752 | 38,06827493 | -1,654848914 | 0,0000708 | 0,000877757 | yes |
| RNF145 | ENSG00000145860 | ENSG00000145860 | protein_coding | mCh_JDV-1-3-5 | F1F2_JDV-2-4-6 | 16211,40836 | 4957,428118 | -1,656415356 | 8,63E-11 | 4,42E-09 | yes |
| IL2RG | ENSG00000147168 | ENSG00000147168 | protein_coding | mCh_JDV-1-3-5 | F1F2_JDV-2-4-6 | 39,48035418 | 10,82684008 | -1,660812423 | 0,000305326 | 0,002920504 | yes |
| FOXA1 | ENSG00000129514 | ENSG00000129514 | protein_coding | mCh_JDV-1-3-5 | F1F2_JDV-2-4-6 | 157,617837 | 41,60749936 | -1,660841004 | 0,00096813 | 0,007294429 | yes |
| C11orf66 | ENSG00000162148 | ENSG00000162148 | protein_coding | mCh_JDV-1-3-5 | F1F2_JDV-2-4-6 | 60,44430523 | 14,46591631 | -1,669283018 | 0,003780913 | 0,021216392 | yes |
| SH2D3A | ENSG00000125731 | ENSG00000125731 | protein_coding | mCh_JDV-1-3-5 | F1F2_JDV-2-4-6 | 597,3855417 | 177,4216765 | -1,669996434 | 7,11E-08 | 0,00000215 | yes |
| COX6B2 | ENSG00000160471 | ENSG00000160471 | protein_coding | mCh_JDV-1-3-5 | F1F2_JDV-2-4-6 | 17,57796806 | 4,036615862 | -1,675432181 | 0,004806753 | 0,025538948 | yes |
| LZTS2 | ENSG00000107816 | ENSG00000107816 | protein_coding | mCh_JDV-1-3-5 | F1F2_JDV-2-4-6 | 1583,444597 | 452,0262966 | -1,679540653 | 0,00000945 | 0,000161254 | yes |
| C3orf52 | ENSG00000114529 | ENSG00000114529 | protein_coding | mCh_JDV-1-3-5 | F1F2_JDV-2-4-6 | 402,1732789 | 119,2372559 | -1,683362298 | 6,58E-09 | 0,000000238 | yes |
| CEL | ENSG00000170835 | ENSG00000170835 | protein_coding | mCh_JDV-1-3-5 | F1F2_JDV-2-4-6 | 98,59268333 | 26,63993678 | -1,686287026 | 0,00020416 | 0,002107057 | yes |
| NBPF11 | ENSG00000203836 | ENSG00000203836 | protein_coding | mCh_JDV-1-3-5 | F1F2_JDV-2-4-6 | 104,1200439 | 28,35602364 | -1,689826987 | 0,000124142 | 0,001387148 | yes |
| RSAD2 | ENSG00000134321 | ENSG00000134321 | protein_coding | mCh_JDV-1-3-5 | F1F2_JDV-2-4-6 | 40,29301202 | 9,446245064 | -1,691053154 | 0,003381573 | 0,019495945 | yes |

| | | | | | | | | | | | |
|---|---|---|---|---|---|---|---|---|---|---|---|
| RHBDF2 | ENSG00000129667 | ENSG00000129667 | protein_coding | mCh_JDV-1-3-5 | F1F2_JDV-2-4-6 | 1183,525764 | 344,2330828 | -1,692373807 | 0,000000139 | 0,00000392 | yes |
| DAB2IP | ENSG00000136848 | ENSG00000136848 | protein_coding | mCh_JDV-1-3-5 | F1F2_JDV-2-4-6 | 1332,888543 | 383,5235981 | -1,696851579 | 0,000000525 | 0,0000127 | yes |
| MYO18A | ENSG00000196535 | ENSG00000196535 | protein_coding | mCh_JDV-1-3-5 | F1F2_JDV-2-4-6 | 5599,573047 | 1612,265149 | -1,698174256 | 0,00000039 | 0,00000974 | yes |
| SLC5A3 | ENSG00000198743 | ENSG00000198743 | protein_coding | mCh_JDV-1-3-5 | F1F2_JDV-2-4-6 | 3937,167615 | 1140,880121 | -1,699248884 | 0,000000095 | 0,00000279 | yes |
| ACCN2 | ENSG00000110881 | ENSG00000110881 | protein_coding | mCh_JDV-1-3-5 | F1F2_JDV-2-4-6 | 342,8088788 | 95,24150346 | -1,701092406 | 0,0000193 | 0,000297201 | yes |
| HOOK2 | ENSG00000095066 | ENSG00000095066 | protein_coding | mCh_JDV-1-3-5 | F1F2_JDV-2-4-6 | 820,9992202 | 233,4511518 | -1,701187216 | 0,00000177 | 0,0000378 | yes |
| GGT6 | ENSG00000167741 | ENSG00000167741 | protein_coding | mCh_JDV-1-3-5 | F1F2_JDV-2-4-6 | 31,06280021 | 8,091870419 | -1,701370538 | 0,000403499 | 0,00365365 | yes |
| AGER | ENSG00000204305 | ENSG00000204305 | protein_coding | mCh_JDV-1-3-5 | F1F2_JDV-2-4-6 | 513,3106109 | 144,302237 | -1,701619958 | 0,00000637 | 0,000114198 | yes |
| C16orf74 | ENSG00000154102 | ENSG00000154102 | protein_coding | mCh_JDV-1-3-5 | F1F2_JDV-2-4-6 | 260,8825182 | 75,16851865 | -1,70288684 | 0,000000173 | 0,00000471 | yes |
| ADAM28 | ENSG00000042980 | ENSG00000042980 | protein_coding | mCh_JDV-1-3-5 | F1F2_JDV-2-4-6 | 16,24354175 | 3,050228992 | -1,711394016 | 0,009758443 | 0,04338967 | yes |
| C1orf132 | ENSG00000203709 | ENSG00000203709 | protein_coding | mCh_JDV-1-3-5 | F1F2_JDV-2-4-6 | 384,6280422 | 91,85630924 | -1,714679723 | 0,001910052 | 0,012443401 | yes |
| FUT6 | ENSG00000156413 | ENSG00000156413 | protein_coding | mCh_JDV-1-3-5 | F1F2_JDV-2-4-6 | 34,70067725 | 9,101290022 | -1,714870945 | 0,00019501 | 0,002031751 | yes |
| SERPINF2 | ENSG00000167711 | ENSG00000167711 | protein_coding | mCh_JDV-1-3-5 | F1F2_JDV-2-4-6 | 122,2363523 | 32,41407841 | -1,715955446 | 0,000132483 | 0,001469522 | yes |
| KIAA1161 | ENSG00000164976 | ENSG00000164976 | protein_coding | mCh_JDV-1-3-5 | F1F2_JDV-2-4-6 | 1483,158794 | 422,1381394 | -1,717990658 | 0,000000165 | 0,00000455 | yes |
| NLRP3 | ENSG00000162711 | ENSG00000162711 | protein_coding | mCh_JDV-1-3-5 | F1F2_JDV-2-4-6 | 42,8327858 | 9,843640851 | -1,726934243 | 0,002457132 | 0,015194791 | yes |
| KIAA1467 | ENSG00000084444 | ENSG00000084444 | protein_coding | mCh_JDV-1-3-5 | F1F2_JDV-2-4-6 | 872,5419122 | 249,3145247 | -1,729137898 | 7,94E-09 | 0,000000282 | yes |
| FAAH2 | ENSG00000165591 | ENSG00000165591 | protein_coding | mCh_JDV-1-3-5 | F1F2_JDV-2-4-6 | 443,8441417 | 105,7483422 | -1,729339394 | 0,001507972 | 0,010282241 | yes |
| MYO1B | ENSG00000128641 | ENSG00000128641 | protein_coding | mCh_JDV-1-3-5 | F1F2_JDV-2-4-6 | 7901,009644 | 2262,307264 | -1,734627845 | 9,86E-10 | 0,000000042 | yes |
| KIF21B | ENSG00000116852 | ENSG00000116852 | protein_coding | mCh_JDV-1-3-5 | F1F2_JDV-2-4-6 | 76,11965552 | 18,7014261 | -1,734963155 | 0,00075883 | 0,006030603 | yes |
| P2RX5 | ENSG00000083454 | ENSG00000083454 | protein_coding | mCh_JDV-1-3-5 | F1F2_JDV-2-4-6 | 516,4802217 | 150,4113438 | -1,735151386 | 5,39E-14 | 4,15E-12 | yes |
| GUCY1A3 | ENSG00000164116 | ENSG00000164116 | protein_coding | mCh_JDV-1-3-5 | F1F2_JDV-2-4-6 | 38,87867009 | 9,467008852 | -1,73610346 | 0,000835865 | 0,006491192 | yes |
| TIAF1 | ENSG00000221995 | ENSG00000221995 | protein_coding | mCh_JDV-1-3-5 | F1F2_JDV-2-4-6 | 297,7807561 | 80,95892565 | -1,736227842 | 0,00000853 | 0,000147398 | yes |
| GJB3 | ENSG00000188910 | ENSG00000188910 | protein_coding | mCh_JDV-1-3-5 | F1F2_JDV-2-4-6 | 2970,654415 | 848,956396 | -1,738265008 | 6,96E-10 | 3,07E-08 | yes |
| IGF1R | ENSG00000140443 | ENSG00000140443 | protein_coding | mCh_JDV-1-3-5 | F1F2_JDV-2-4-6 | 5312,62395 | 1512,508525 | -1,740830694 | 1,34E-09 | 0,000000055 | yes |
| STARD5 | ENSG00000172345 | ENSG00000172345 | protein_coding | mCh_JDV-1-3-5 | F1F2_JDV-2-4-6 | 105,2820208 | 28,40928336 | -1,741016732 | 0,0000115 | 0,000190472 | yes |
| NR6A1 | ENSG00000148200 | ENSG00000148200 | protein_coding | mCh_JDV-1-3-5 | F1F2_JDV-2-4-6 | 170,8936834 | 47,21669296 | -1,745193629 | 0,000000522 | 0,0000127 | yes |
| TPRG1 | ENSG00000188001 | ENSG00000188001 | protein_coding | mCh_JDV-1-3-5 | F1F2_JDV-2-4-6 | 185,4812702 | 47,90752174 | -1,746451338 | 0,000113162 | 0,001290772 | yes |
| MAPK15 | ENSG00000181085 | ENSG00000181085 | protein_coding | mCh_JDV-1-3-5 | F1F2_JDV-2-4-6 | 47,24490499 | 11,83133437 | -1,746613133 | 0,000326093 | 0,003081799 | yes |
| COBL | ENSG00000106078 | ENSG00000106078 | protein_coding | mCh_JDV-1-3-5 | F1F2_JDV-2-4-6 | 996,368349 | 262,7809951 | -1,749274184 | 0,0000337 | 0,000474676 | yes |
| SERPINA3 | ENSG00000196136 | ENSG00000196136 | protein_coding | mCh_JDV-1-3-5 | F1F2_JDV-2-4-6 | 35564,46069 | 8107,407116 | -1,751932223 | 0,001876281 | 0,012272012 | yes |
| CORO1A | ENSG00000102879 | ENSG00000102879 | protein_coding | mCh_JDV-1-3-5 | F1F2_JDV-2-4-6 | 55,92181238 | 12,23259292 | -1,754985617 | 0,002750091 | 0,016651594 | yes |
| ZNF165 | ENSG00000197279 | ENSG00000197279 | protein_coding | mCh_JDV-1-3-5 | F1F2_JDV-2-4-6 | 144,1195488 | 37,53201229 | -1,755726868 | 0,0000479 | 0,000635443 | yes |
| ZNF488 | ENSG00000165388 | ENSG00000165388 | protein_coding | mCh_JDV-1-3-5 | F1F2_JDV-2-4-6 | 64,55261817 | 15,64520926 | -1,755827933 | 0,000607147 | 0,005044562 | yes |
| SPINT2 | ENSG00000167642 | ENSG00000167642 | protein_coding | mCh_JDV-1-3-5 | F1F2_JDV-2-4-6 | 12051,17355 | 3373,137164 | -1,758593521 | 3,53E-09 | 0,000000134 | yes |
| STMN3 | ENSG00000197457 | ENSG00000197457 | protein_coding | mCh_JDV-1-3-5 | F1F2_JDV-2-4-6 | 116,3248712 | 32,08404315 | -1,758594606 | 0,000000113 | 0,00000326 | yes |
| SERPINA1 | ENSG00000197249 | ENSG00000197249 | protein_coding | mCh_JDV-1-3-5 | F1F2_JDV-2-4-6 | 6055,869678 | 1628,549957 | -1,761129351 | 0,00000302 | 0,0000598 | yes |
| CPT1B | ENSG00000205560 | ENSG00000205560 | protein_coding | mCh_JDV-1-3-5 | F1F2_JDV-2-4-6 | 3831,668473 | 961,6012104 | -1,763293396 | 0,000183017 | 0,001926118 | yes |
| POU5F1 | ENSG00000204531 | ENSG00000204531 | protein_coding | mCh_JDV-1-3-5 | F1F2_JDV-2-4-6 | 93,57884832 | 25,00758019 | -1,767937197 | 0,00000307 | 0,0000607 | yes |
| EPGN | ENSG00000182585 | ENSG00000182585 | protein_coding | mCh_JDV-1-3-5 | F1F2_JDV-2-4-6 | 2178,804997 | 573,0501667 | -1,769422499 | 0,0000115 | 0,000191082 | yes |
| IL17RD | ENSG00000144730 | ENSG00000144730 | protein_coding | mCh_JDV-1-3-5 | F1F2_JDV-2-4-6 | 334,7318939 | 90,22980598 | -1,771845355 | 0,000000799 | 0,0000188 | yes |
| KIAA0247 | ENSG00000100647 | ENSG00000100647 | protein_coding | mCh_JDV-1-3-5 | F1F2_JDV-2-4-6 | 3917,860746 | 1065,029545 | -1,773606236 | 0,000000171 | 0,00000467 | yes |
| SFN | ENSG00000175793 | ENSG00000175793 | protein_coding | mCh_JDV-1-3-5 | F1F2_JDV-2-4-6 | 4595,733316 | 1156,884215 | -1,775728439 | 0,000100167 | 0,001166824 | yes |
| APOE | ENSG00000130203 | ENSG00000130203 | protein_coding | mCh_JDV-1-3-5 | F1F2_JDV-2-4-6 | 23,82564751 | 4,388090035 | -1,776539204 | 0,00624726 | 0,031217317 | yes |
| COMTD1 | ENSG00000165644 | ENSG00000165644 | protein_coding | mCh_JDV-1-3-5 | F1F2_JDV-2-4-6 | 132,1342778 | 35,77411015 | -1,776734563 | 0,000000203 | 0,00000544 | yes |
| SOX7 | ENSG00000171056 | ENSG00000171056 | protein_coding | mCh_JDV-1-3-5 | F1F2_JDV-2-4-6 | 1033,963097 | 290,8738382 | -1,780875591 | 6,6E-14 | 5,03E-12 | yes |
| PRKCQ | ENSG00000065675 | ENSG00000065675 | protein_coding | mCh_JDV-1-3-5 | F1F2_JDV-2-4-6 | 34,64675402 | 8,096264456 | -1,783031973 | 0,0006207 | 0,005133395 | yes |
| PPP4R4 | ENSG00000119698 | ENSG00000119698 | protein_coding | mCh_JDV-1-3-5 | F1F2_JDV-2-4-6 | 24,04386325 | 5,38819029 | -1,788582444 | 0,001121138 | 0,008199691 | yes |

| | | | | | | | | | | | |
|---|---|---|---|---|---|---|---|---|---|---|---|
| RASSF9 | ENSG00000198774 | ENSG00000198774 | protein_coding | mCh_JDV-1-3-5 | F1F2_JDV-2-4-6 | 104,3464347 | 28,02212562 | -1,789417491 | 0,000000125 | 0,00000356 | yes |
| IL7R | ENSG00000168685 | ENSG00000168685 | protein_coding | mCh_JDV-1-3-5 | F1F2_JDV-2-4-6 | 185,4853554 | 41,67553501 | -1,791478958 | 0,001125781 | 0,00821258 | yes |
| FAM83H-AS1 | ENSG00000203499 | ENSG00000203499 | protein_coding | mCh_JDV-1-3-5 | F1F2_JDV-2-4-6 | 630,5655612 | 167,7441854 | -1,794433493 | 0,000000326 | 0,00000836 | yes |
| CLDN23 | ENSG00000253958 | ENSG00000253958 | protein_coding | mCh_JDV-1-3-5 | F1F2_JDV-2-4-6 | 68,85163259 | 16,88495151 | -1,797579405 | 0,000111984 | 0,001278803 | yes |
| WNT10B | ENSG00000169884 | ENSG00000169884 | protein_coding | mCh_JDV-1-3-5 | F1F2_JDV-2-4-6 | 79,19458117 | 19,64348518 | -1,801692519 | 0,0000596 | 0,000764398 | yes |
| ICAM5 | ENSG00000105376 | ENSG00000105376 | protein_coding | mCh_JDV-1-3-5 | F1F2_JDV-2-4-6 | 328,6277455 | 77,49850618 | -1,807247957 | 0,000295253 | 0,002836523 | yes |
| CD36 | ENSG00000135218 | ENSG00000135218 | protein_coding | mCh_JDV-1-3-5 | F1F2_JDV-2-4-6 | 46,79704205 | 9,118866169 | -1,810165546 | 0,003439663 | 0,019734438 | yes |
| CSF1R | ENSG00000182578 | ENSG00000182578 | protein_coding | mCh_JDV-1-3-5 | F1F2_JDV-2-4-6 | 344,1324627 | 88,94998145 | -1,810292323 | 0,00000215 | 0,0000447 | yes |
| PCP2 | ENSG00000174788 | ENSG00000174788 | protein_coding | mCh_JDV-1-3-5 | F1F2_JDV-2-4-6 | 27,09599854 | 5,38819029 | -1,812071291 | 0,002833827 | 0,017019417 | yes |
| SLC16A7 | ENSG00000118596 | ENSG00000118596 | protein_coding | mCh_JDV-1-3-5 | F1F2_JDV-2-4-6 | 1123,337235 | 296,0457578 | -1,812075139 | 0,000000147 | 0,0000041 | yes |
| SLC37A2 | ENSG00000134955 | ENSG00000134955 | protein_coding | mCh_JDV-1-3-5 | F1F2_JDV-2-4-6 | 1459,742381 | 332,5254171 | -1,819334573 | 0,000489337 | 0,004250038 | yes |
| GM2A | ENSG00000196743 | ENSG00000196743 | protein_coding | mCh_JDV-1-3-5 | F1F2_JDV-2-4-6 | 10776,98326 | 2847,689152 | -1,825281222 | 1,08E-08 | 0,000000372 | yes |
| GAL3ST4 | ENSG00000197093 | ENSG00000197093 | protein_coding | mCh_JDV-1-3-5 | F1F2_JDV-2-4-6 | 26,85402323 | 5,712768968 | -1,825826058 | 0,00117845 | 0,008508422 | yes |
| P2RY1 | ENSG00000169860 | ENSG00000169860 | protein_coding | mCh_JDV-1-3-5 | F1F2_JDV-2-4-6 | 25,04150246 | 4,426429972 | -1,831720672 | 0,004561815 | 0,024530233 | yes |
| KAZN | ENSG00000189337 | ENSG00000189337 | protein_coding | mCh_JDV-1-3-5 | F1F2_JDV-2-4-6 | 533,4564831 | 144,9897389 | -1,83272634 | 1,33E-15 | 1,28E-13 | yes |
| SLC4A3 | ENSG00000114923 | ENSG00000114923 | protein_coding | mCh_JDV-1-3-5 | F1F2_JDV-2-4-6 | 488,229393 | 123,8130738 | -1,835971607 | 0,00000151 | 0,0000333 | yes |
| LRIG1 | ENSG00000144749 | ENSG00000144749 | protein_coding | mCh_JDV-1-3-5 | F1F2_JDV-2-4-6 | 1736,10715 | 443,4486625 | -1,839337465 | 0,000000478 | 0,0000117 | yes |
| SLC35F3 | ENSG00000183780 | ENSG00000183780 | protein_coding | mCh_JDV-1-3-5 | F1F2_JDV-2-4-6 | 76,1195242 | 17,32469385 | -1,840660055 | 0,000280561 | 0,002721886 | yes |
| ACOT11 | ENSG00000162390 | ENSG00000162390 | protein_coding | mCh_JDV-1-3-5 | F1F2_JDV-2-4-6 | 780,3965958 | 205,929705 | -1,841168962 | 4,59E-10 | 2,12E-08 | yes |
| LYST | ENSG00000143669 | ENSG00000143669 | protein_coding | mCh_JDV-1-3-5 | F1F2_JDV-2-4-6 | 2003,096317 | 518,0487705 | -1,841180499 | 5,93E-08 | 0,00000181 | yes |
| TIMP3 | ENSG00000100234 | ENSG00000100234 | protein_coding | mCh_JDV-1-3-5 | F1F2_JDV-2-4-6 | 7632,01506 | 2007,292563 | -1,845292077 | 4,9E-10 | 2,23E-08 | yes |
| STC1 | ENSG00000159167 | ENSG00000159167 | protein_coding | mCh_JDV-1-3-5 | F1F2_JDV-2-4-6 | 1618,975637 | 416,8772833 | -1,849094362 | 4,03E-08 | 0,00000127 | yes |
| CNTNAP3B | ENSG00000154529 | ENSG00000154529 | protein_coding | mCh_JDV-1-3-5 | F1F2_JDV-2-4-6 | 88,58866419 | 22,62794747 | -1,849364023 | 0,000000136 | 0,00000384 | yes |
| PLAU | ENSG00000122861 | ENSG00000122861 | protein_coding | mCh_JDV-1-3-5 | F1F2_JDV-2-4-6 | 5035,385216 | 1262,232071 | -1,851187592 | 0,00000128 | 0,0000287 | yes |
| ELOVL7 | ENSG00000164181 | ENSG00000164181 | protein_coding | mCh_JDV-1-3-5 | F1F2_JDV-2-4-6 | 590,699148 | 156,8676654 | -1,853390407 | 3,77E-13 | 2,61E-11 | yes |
| DDX26B | ENSG00000165359 | ENSG00000165359 | protein_coding | mCh_JDV-1-3-5 | F1F2_JDV-2-4-6 | 901,6442672 | 233,8780995 | -1,854476689 | 3,14E-09 | 0,000000121 | yes |
| CACNB4 | ENSG00000182389 | ENSG00000182389 | protein_coding | mCh_JDV-1-3-5 | F1F2_JDV-2-4-6 | 53,23496116 | 12,82757186 | -1,856163392 | 0,0000162 | 0,000255326 | yes |
| DUOX1 | ENSG00000137857 | ENSG00000137857 | protein_coding | mCh_JDV-1-3-5 | F1F2_JDV-2-4-6 | 1326,224839 | 353,0290638 | -1,85684326 | 1,28E-14 | 1,07E-12 | yes |
| ALDH2 | ENSG00000111275 | ENSG00000111275 | protein_coding | mCh_JDV-1-3-5 | F1F2_JDV-2-4-6 | 31,785515 | 5,753377847 | -1,857940671 | 0,003177521 | 0,018622517 | yes |
| ATP13A4 | ENSG00000127249 | ENSG00000127249 | protein_coding | mCh_JDV-1-3-5 | F1F2_JDV-2-4-6 | 11,63122805 | 1,675621832 | -1,858418363 | 0,006522604 | 0,032258283 | yes |
| MAPKBP1 | ENSG00000137802 | ENSG00000137802 | protein_coding | mCh_JDV-1-3-5 | F1F2_JDV-2-4-6 | 2736,087094 | 701,9984883 | -1,859838998 | 1,45E-08 | 0,00000049 | yes |
| PPARGC1B | ENSG00000155846 | ENSG00000155846 | protein_coding | mCh_JDV-1-3-5 | F1F2_JDV-2-4-6 | 1618,265444 | 417,9504582 | -1,85996469 | 3,09E-09 | 0,00000012 | yes |
| SLC2A1 | ENSG00000117394 | ENSG00000117394 | protein_coding | mCh_JDV-1-3-5 | F1F2_JDV-2-4-6 | 4573,226603 | 1200,685929 | -1,862689443 | 4,46E-12 | 2,72E-10 | yes |
| KRT14 | ENSG00000186847 | ENSG00000186847 | protein_coding | mCh_JDV-1-3-5 | F1F2_JDV-2-4-6 | 1248,824265 | 258,6055263 | -1,865596756 | 0,000927304 | 0,007043299 | yes |
| SLC4A11 | ENSG00000088836 | ENSG00000088836 | protein_coding | mCh_JDV-1-3-5 | F1F2_JDV-2-4-6 | 483,310465 | 117,5206726 | -1,867100783 | 0,0000052 | 0,0000961 | yes |
| KLHDC8B | ENSG00000185909 | ENSG00000185909 | protein_coding | mCh_JDV-1-3-5 | F1F2_JDV-2-4-6 | 165,6769545 | 32,37906997 | -1,867742045 | 0,001685976 | 0,01128923 | yes |
| IGSF9B | ENSG00000080854 | ENSG00000080854 | protein_coding | mCh_JDV-1-3-5 | F1F2_JDV-2-4-6 | 350,6339156 | 86,30501762 | -1,868313853 | 0,00000177 | 0,0000378 | yes |
| FYB | ENSG00000082074 | ENSG00000082074 | protein_coding | mCh_JDV-1-3-5 | F1F2_JDV-2-4-6 | 107,8177864 | 26,71062879 | -1,876955662 | 0,000000379 | 0,00000951 | yes |
| CASP10 | ENSG00000003400 | ENSG00000003400 | protein_coding | mCh_JDV-1-3-5 | F1F2_JDV-2-4-6 | 92,13151204 | 22,97555888 | -1,877003391 | 0,000000143 | 0,00000402 | yes |
| MARCO | ENSG00000019169 | ENSG00000019169 | protein_coding | mCh_JDV-1-3-5 | F1F2_JDV-2-4-6 | 121,1716421 | 26,349304 | -1,878802525 | 0,000294742 | 0,002833001 | yes |
| CPNE2 | ENSG00000140848 | ENSG00000140848 | protein_coding | mCh_JDV-1-3-5 | F1F2_JDV-2-4-6 | 1414,845257 | 370,443629 | -1,883271278 | 9,05E-16 | 8,89E-14 | yes |
| MFAP5 | ENSG00000197614 | ENSG00000197614 | protein_coding | mCh_JDV-1-3-5 | F1F2_JDV-2-4-6 | 50,87489694 | 9,171982037 | -1,886437175 | 0,002421249 | 0,015019965 | yes |
| FAM131B | ENSG00000159784 | ENSG00000159784 | protein_coding | mCh_JDV-1-3-5 | F1F2_JDV-2-4-6 | 53,23805449 | 11,10974733 | -1,888937951 | 0,000500499 | 0,00431847 | yes |
| IL17RE | ENSG00000163701 | ENSG00000163701 | protein_coding | mCh_JDV-1-3-5 | F1F2_JDV-2-4-6 | 1426,315933 | 353,5158298 | -1,904816636 | 9,03E-09 | 0,000000316 | yes |
| MEI1 | ENSG00000167077 | ENSG00000167077 | protein_coding | mCh_JDV-1-3-5 | F1F2_JDV-2-4-6 | 17,9403762 | 3,389052327 | -1,906405426 | 0,001224676 | 0,008727532 | yes |
| RNF122 | ENSG00000133874 | ENSG00000133874 | protein_coding | mCh_JDV-1-3-5 | F1F2_JDV-2-4-6 | 42,96396614 | 9,490041584 | -1,907192099 | 0,0000807 | 0,000972943 | yes |
| NRARP | ENSG00000198435 | ENSG00000198435 | protein_coding | mCh_JDV-1-3-5 | F1F2_JDV-2-4-6 | 128,9589877 | 31,7928791 | -1,909996124 | 1,14E-08 | 0,000000393 | yes |

| | | | | | | | | | | | |
|---|---|---|---|---|---|---|---|---|---|---|---|
| MYO5B | ENSG00000167306 | ENSG00000167306 | protein_coding | mCh_JDV-1-3-5 | F1F2_JDV-2-4-6 | 824,4511112 | 189,343831 | -1,912716569 | 0,0000133 | 0,000214828 | yes |
| ETS2 | ENSG00000157557 | ENSG00000157557 | protein_coding | mCh_JDV-1-3-5 | F1F2_JDV-2-4-6 | 4613,660516 | 1084,967039 | -1,914428501 | 0,00000253 | 0,0000516 | yes |
| GPR141 | ENSG00000187037 | ENSG00000187037 | protein_coding | mCh_JDV-1-3-5 | F1F2_JDV-2-4-6 | 163,8980486 | 35,71568155 | -1,91617779 | 0,0000958 | 0,001123463 | yes |
| RHBDL1 | ENSG00000103269 | ENSG00000103269 | protein_coding | mCh_JDV-1-3-5 | F1F2_JDV-2-4-6 | 70,31467119 | 14,17808374 | -1,920490347 | 0,000470002 | 0,004129367 | yes |
| CNTD2 | ENSG00000105219 | ENSG00000105219 | protein_coding | mCh_JDV-1-3-5 | F1F2_JDV-2-4-6 | 27,42241859 | 5,041641428 | -1,922522344 | 0,001272477 | 0,008987041 | yes |
| NBEAL2 | ENSG00000160796 | ENSG00000160796 | protein_coding | mCh_JDV-1-3-5 | F1F2_JDV-2-4-6 | 4615,400006 | 1155,918946 | -1,923377999 | 4,86E-12 | 2,93E-10 | yes |
| LBX2 | ENSG00000179528 | ENSG00000179528 | protein_coding | mCh_JDV-1-3-5 | F1F2_JDV-2-4-6 | 46,44258246 | 8,786030691 | -1,925021473 | 0,00097909 | 0,007362952 | yes |
| NUP210 | ENSG00000132182 | ENSG00000132182 | protein_coding | mCh_JDV-1-3-5 | F1F2_JDV-2-4-6 | 4257,916249 | 1060,816131 | -1,928473574 | 8,41E-12 | 5E-10 | yes |
| SLC44A3 | ENSG00000143036 | ENSG00000143036 | protein_coding | mCh_JDV-1-3-5 | F1F2_JDV-2-4-6 | 285,8355947 | 70,09133284 | -1,93392095 | 3,65E-10 | 1,73E-08 | yes |
| RAB11FIP4 | ENSG00000131242 | ENSG00000131242 | protein_coding | mCh_JDV-1-3-5 | F1F2_JDV-2-4-6 | 338,8886996 | 77,44553416 | -1,934725721 | 0,00000503 | 0,0000934 | yes |
| TLL1 | ENSG00000038295 | ENSG00000038295 | protein_coding | mCh_JDV-1-3-5 | F1F2_JDV-2-4-6 | 194,8068555 | 42,1396162 | -1,952419715 | 0,0000346 | 0,000485004 | yes |
| PLAT | ENSG00000104368 | ENSG00000104368 | protein_coding | mCh_JDV-1-3-5 | F1F2_JDV-2-4-6 | 5796,480219 | 1388,794449 | -1,954093251 | 2,34E-09 | 9,31E-08 | yes |
| LEAP2 | ENSG00000164406 | ENSG00000164406 | protein_coding | mCh_JDV-1-3-5 | F1F2_JDV-2-4-6 | 311,1386561 | 72,18207045 | -1,955871026 | 0,000000276 | 0,00000717 | yes |
| ZFP57 | ENSG00000204644 | ENSG00000204644 | protein_coding | mCh_JDV-1-3-5 | F1F2_JDV-2-4-6 | 43,5911852 | 9,433594226 | -1,956052195 | 0,0000253 | 0,000371837 | yes |
| SLC46A1 | ENSG00000076351 | ENSG00000076351 | protein_coding | mCh_JDV-1-3-5 | F1F2_JDV-2-4-6 | 4437,398805 | 1049,775343 | -1,957942117 | 1,49E-08 | 0,000000503 | yes |
| MYO10 | ENSG00000145555 | ENSG00000145555 | protein_coding | mCh_JDV-1-3-5 | F1F2_JDV-2-4-6 | 1859,804106 | 431,018491 | -1,959971769 | 0,000000214 | 0,00000569 | yes |
| PRODH | ENSG00000100033 | ENSG00000100033 | protein_coding | mCh_JDV-1-3-5 | F1F2_JDV-2-4-6 | 496,4300174 | 56,23400913 | -1,961307465 | 0,005637764 | 0,028916287 | yes |
| IL1RAP | ENSG00000196083 | ENSG00000196083 | protein_coding | mCh_JDV-1-3-5 | F1F2_JDV-2-4-6 | 2224,156197 | 524,3757278 | -1,962135198 | 1,48E-08 | 0,0000005 | yes |
| C2orf48 | ENSG00000163009 | ENSG00000163009 | protein_coding | mCh_JDV-1-3-5 | F1F2_JDV-2-4-6 | 28,39397933 | 5,407891532 | -1,962929948 | 0,000447939 | 0,003974454 | yes |
| MAP7 | ENSG00000135525 | ENSG00000135525 | protein_coding | mCh_JDV-1-3-5 | F1F2_JDV-2-4-6 | 791,6916575 | 184,4597995 | -1,969117416 | 3,68E-08 | 0,00000117 | yes |
| C10orf57 | ENSG00000122378 | ENSG00000122378 | protein_coding | mCh_JDV-1-3-5 | F1F2_JDV-2-4-6 | 3273,509356 | 801,6337867 | -1,972738553 | 5,56E-16 | 5,62E-14 | yes |
| CCDC88C | ENSG00000015133 | ENSG00000015133 | protein_coding | mCh_JDV-1-3-5 | F1F2_JDV-2-4-6 | 1412,869915 | 344,7574787 | -1,977577835 | 5,38E-16 | 5,53E-14 | yes |
| HES7 | ENSG00000179111 | ENSG00000179111 | protein_coding | mCh_JDV-1-3-5 | F1F2_JDV-2-4-6 | 139,4768358 | 31,33570447 | -1,97884138 | 0,000000803 | 0,0000189 | yes |
| MALL | ENSG00000144063 | ENSG00000144063 | protein_coding | mCh_JDV-1-3-5 | F1F2_JDV-2-4-6 | 438,4894307 | 77,05501013 | -1,979436053 | 0,000950064 | 0,007188969 | yes |
| TMC5 | ENSG00000103534 | ENSG00000103534 | protein_coding | mCh_JDV-1-3-5 | F1F2_JDV-2-4-6 | 283,4157285 | 59,24295412 | -1,984059355 | 0,0000357 | 0,000497248 | yes |
| CHST6 | ENSG00000183196 | ENSG00000183196 | protein_coding | mCh_JDV-1-3-5 | F1F2_JDV-2-4-6 | 88,8637969 | 20,22475074 | -1,987221819 | 9,23E-08 | 0,00000272 | yes |
| SPINT1 | ENSG00000166145 | ENSG00000166145 | protein_coding | mCh_JDV-1-3-5 | F1F2_JDV-2-4-6 | 7356,494438 | 1786,050524 | -1,987589839 | 6,33E-17 | 6,98E-15 | yes |
| CCL5 | ENSG00000161570 | ENSG00000161570 | protein_coding | mCh_JDV-1-3-5 | F1F2_JDV-2-4-6 | 24,53521891 | 4,045403936 | -1,98928464 | 0,001262605 | 0,008930105 | yes |
| KIAA1543 | ENSG00000076826 | ENSG00000076826 | protein_coding | mCh_JDV-1-3-5 | F1F2_JDV-2-4-6 | 275,1131725 | 63,45702501 | -1,990148462 | 1,08E-08 | 0,000000373 | yes |
| SDR16C5 | ENSG00000170786 | ENSG00000170786 | protein_coding | mCh_JDV-1-3-5 | F1F2_JDV-2-4-6 | 462,6611808 | 96,79708044 | -1,991013388 | 0,0000269 | 0,000391535 | yes |
| FRMD5 | ENSG00000171877 | ENSG00000171877 | protein_coding | mCh_JDV-1-3-5 | F1F2_JDV-2-4-6 | 48,55851112 | 9,800375604 | -1,993012526 | 0,0000722 | 0,000889575 | yes |
| IL12A | ENSG00000168811 | ENSG00000168811 | protein_coding | mCh_JDV-1-3-5 | F1F2_JDV-2-4-6 | 77,08859386 | 16,87630729 | -2,010954924 | 0,000000599 | 0,0000144 | yes |
| AP3B2 | ENSG00000103723 | ENSG00000103723 | protein_coding | mCh_JDV-1-3-5 | F1F2_JDV-2-4-6 | 72,73858649 | 15,23250626 | -2,014576251 | 0,00000899 | 0,000154746 | yes |
| MFSD2A | ENSG00000168389 | ENSG00000168389 | protein_coding | mCh_JDV-1-3-5 | F1F2_JDV-2-4-6 | 6188,911467 | 1386,015562 | -2,016786966 | 3,31E-08 | 0,00000106 | yes |
| TMTC2 | ENSG00000179104 | ENSG00000179104 | protein_coding | mCh_JDV-1-3-5 | F1F2_JDV-2-4-6 | 535,7113641 | 122,9821765 | -2,017453015 | 3,03E-10 | 1,45E-08 | yes |
| RAET1L | ENSG00000155918 | ENSG00000155918 | protein_coding | mCh_JDV-1-3-5 | F1F2_JDV-2-4-6 | 22,8125325 | 2,712468203 | -2,023156585 | 0,003319661 | 0,019238438 | yes |
| CTSL2 | ENSG00000136943 | ENSG00000136943 | protein_coding | mCh_JDV-1-3-5 | F1F2_JDV-2-4-6 | 449,0900974 | 79,98926115 | -2,027463885 | 0,000383945 | 0,003515723 | yes |
| GRB7 | ENSG00000141738 | ENSG00000141738 | protein_coding | mCh_JDV-1-3-5 | F1F2_JDV-2-4-6 | 835,4318218 | 181,4603177 | -2,02803838 | 0,000000346 | 0,00000879 | yes |
| PLEKHG3 | ENSG00000126822 | ENSG00000126822 | protein_coding | mCh_JDV-1-3-5 | F1F2_JDV-2-4-6 | 2086,072511 | 486,2790773 | -2,036567917 | 1,08E-15 | 1,06E-13 | yes |
| FAM134B | ENSG00000154153 | ENSG00000154153 | protein_coding | mCh_JDV-1-3-5 | F1F2_JDV-2-4-6 | 727,0961304 | 165,5135231 | -2,04013467 | 1,88E-11 | 1,07E-09 | yes |
| RIMS3 | ENSG00000117016 | ENSG00000117016 | protein_coding | mCh_JDV-1-3-5 | F1F2_JDV-2-4-6 | 75,70321544 | 14,92096585 | -2,040928794 | 0,000033 | 0,000465477 | yes |
| GALNT14 | ENSG00000158089 | ENSG00000158089 | protein_coding | mCh_JDV-1-3-5 | F1F2_JDV-2-4-6 | 527,9156804 | 102,8392791 | -2,048027106 | 0,0000395 | 0,000541775 | yes |
| C9orf61 | ENSG00000135063 | ENSG00000135063 | protein_coding | mCh_JDV-1-3-5 | F1F2_JDV-2-4-6 | 59,33618365 | 12,11371035 | -2,049651691 | 0,00000503 | 0,0000934 | yes |
| IL22RA1 | ENSG00000142677 | ENSG00000142677 | protein_coding | mCh_JDV-1-3-5 | F1F2_JDV-2-4-6 | 149,9588046 | 32,53349226 | -2,053259425 | 3,54E-08 | 0,00000113 | yes |
| IP6K3 | ENSG00000161896 | ENSG00000161896 | protein_coding | mCh_JDV-1-3-5 | F1F2_JDV-2-4-6 | 127,1081417 | 20,14527065 | -2,056467529 | 0,0007903 | 0,006214629 | yes |
| AL353791.1 | ENSG00000227921 | ENSG00000227921 | protein_coding | mCh_JDV-1-3-5 | F1F2_JDV-2-4-6 | 42,45218427 | 7,758503668 | -2,063471293 | 0,000109603 | 0,001255381 | yes |
| DDR1 | ENSG00000204580 | ENSG00000204580 | protein_coding | mCh_JDV-1-3-5 | F1F2_JDV-2-4-6 | 11497,8192 | 2596,202152 | -2,072833157 | 1,26E-14 | 1,06E-12 | yes |

| | | | | | | | | | | | |
|---|---|---|---|---|---|---|---|---|---|---|---|
| THSD1 | ENSG00000136114 | ENSG00000136114 | protein_coding | mCh_JDV-1-3-5 | F1F2_JDV-2-4-6 | 593,5670763 | 132,9071743 | -2,073009802 | 6,53E-13 | 4,43E-11 | yes |
| TNFRSF25 | ENSG00000215788 | ENSG00000215788 | protein_coding | mCh_JDV-1-3-5 | F1F2_JDV-2-4-6 | 1372,595051 | 299,0761209 | -2,076902602 | 6,64E-10 | 2,94E-08 | yes |
| PLXNB1 | ENSG00000164050 | ENSG00000164050 | protein_coding | mCh_JDV-1-3-5 | F1F2_JDV-2-4-6 | 15455,9382 | 3170,241554 | -2,078271589 | 0,000000886 | 0,0000205 | yes |
| CBWD3 | ENSG00000196873 | ENSG00000196873 | protein_coding | mCh_JDV-1-3-5 | F1F2_JDV-2-4-6 | 63,98650087 | 11,83625968 | -2,07881562 | 0,0000589 | 0,000757804 | yes |
| GLS2 | ENSG00000135423 | ENSG00000135423 | protein_coding | mCh_JDV-1-3-5 | F1F2_JDV-2-4-6 | 553,7449755 | 113,5069456 | -2,082679303 | 0,000000674 | 0,0000161 | yes |
| CPNE4 | ENSG00000196353 | ENSG00000196353 | protein_coding | mCh_JDV-1-3-5 | F1F2_JDV-2-4-6 | 25,36040877 | 3,379201706 | -2,086855758 | 0,001405837 | 0,00970997 | yes |
| WDR21B | ENSG00000182308 | ENSG00000182308 | protein_coding | mCh_JDV-1-3-5 | F1F2_JDV-2-4-6 | 68,88949566 | 13,51840065 | -2,089044487 | 0,00000563 | 0,000102822 | yes |
| LMTK3 | ENSG00000142235 | ENSG00000142235 | protein_coding | mCh_JDV-1-3-5 | F1F2_JDV-2-4-6 | 106,1621122 | 20,53682243 | -2,092206884 | 0,00000906 | 0,000155256 | yes |
| C6orf105 | ENSG00000111863 | ENSG00000111863 | protein_coding | mCh_JDV-1-3-5 | F1F2_JDV-2-4-6 | 414,7759495 | 76,695423 | -2,094957583 | 0,0000397 | 0,000544104 | yes |
| HOOK1 | ENSG00000134709 | ENSG00000134709 | protein_coding | mCh_JDV-1-3-5 | F1F2_JDV-2-4-6 | 1412,847256 | 317,9556772 | -2,097552704 | 8,91E-20 | 1,27E-17 | yes |
| TGFA | ENSG00000163235 | ENSG00000163235 | protein_coding | mCh_JDV-1-3-5 | F1F2_JDV-2-4-6 | 1678,406402 | 363,7994264 | -2,098159554 | 3,87E-11 | 2,1E-09 | yes |
| SCD5 | ENSG00000145284 | ENSG00000145284 | protein_coding | mCh_JDV-1-3-5 | F1F2_JDV-2-4-6 | 2431,695152 | 522,4601415 | -2,100163863 | 2,24E-10 | 1,08E-08 | yes |
| IL4R | ENSG00000077238 | ENSG00000077238 | protein_coding | mCh_JDV-1-3-5 | F1F2_JDV-2-4-6 | 6201,252343 | 1394,051011 | -2,107789322 | 2,54E-23 | 5,51E-21 | yes |
| PGF | ENSG00000119630 | ENSG00000119630 | protein_coding | mCh_JDV-1-3-5 | F1F2_JDV-2-4-6 | 76,3997974 | 14,86186212 | -2,114905262 | 0,00000236 | 0,0000482 | yes |
| COBLL1 | ENSG00000082438 | ENSG00000082438 | protein_coding | mCh_JDV-1-3-5 | F1F2_JDV-2-4-6 | 1138,203338 | 249,480823 | -2,117636646 | 7,79E-16 | 7,76E-14 | yes |
| C1orf38 | ENSG00000130775 | ENSG00000130775 | protein_coding | mCh_JDV-1-3-5 | F1F2_JDV-2-4-6 | 340,7883057 | 66,75592765 | -2,124210719 | 0,00000103 | 0,0000237 | yes |
| SLPI | ENSG00000124107 | ENSG00000124107 | protein_coding | mCh_JDV-1-3-5 | F1F2_JDV-2-4-6 | 9983,409798 | 2009,798211 | -2,12662929 | 0,000000109 | 0,00000313 | yes |
| ABCC2 | ENSG00000023839 | ENSG00000023839 | protein_coding | mCh_JDV-1-3-5 | F1F2_JDV-2-4-6 | 14182,1226 | 2949,05009 | -2,13023379 | 1,07E-09 | 4,49E-08 | yes |
| IRF5 | ENSG00000128604 | ENSG00000128604 | protein_coding | mCh_JDV-1-3-5 | F1F2_JDV-2-4-6 | 584,0312646 | 124,0308942 | -2,131947789 | 5,61E-12 | 3,35E-10 | yes |
| EDA | ENSG00000158813 | ENSG00000158813 | protein_coding | mCh_JDV-1-3-5 | F1F2_JDV-2-4-6 | 318,8116245 | 69,33846093 | -2,132405253 | 8,72E-17 | 9,45E-15 | yes |
| CNTNAP3 | ENSG00000106714 | ENSG00000106714 | protein_coding | mCh_JDV-1-3-5 | F1F2_JDV-2-4-6 | 294,4637555 | 60,96754602 | -2,136658617 | 9,23E-10 | 3,98E-08 | yes |
| FGF11 | ENSG00000161958 | ENSG00000161958 | protein_coding | mCh_JDV-1-3-5 | F1F2_JDV-2-4-6 | 133,4987273 | 26,67122631 | -2,141936806 | 5,52E-08 | 0,00000169 | yes |
| AC148477.2 | ENSG00000204589 | ENSG00000204589 | protein_coding | mCh_JDV-1-3-5 | F1F2_JDV-2-4-6 | 39,85274891 | 5,067474376 | -2,158725602 | 0,000891162 | 0,006834555 | yes |
| GOLGA2P5 | ENSG00000238105 | ENSG00000238105 | protein_coding | mCh_JDV-1-3-5 | F1F2_JDV-2-4-6 | 187,1017489 | 37,79241808 | -2,158953855 | 2,19E-09 | 8,75E-08 | yes |
| DQX1 | ENSG00000144045 | ENSG00000144045 | protein_coding | mCh_JDV-1-3-5 | F1F2_JDV-2-4-6 | 264,7522614 | 45,79389525 | -2,159313655 | 0,0000329 | 0,000465461 | yes |
| CARD10 | ENSG00000100065 | ENSG00000100065 | protein_coding | mCh_JDV-1-3-5 | F1F2_JDV-2-4-6 | 774,9313838 | 147,6204233 | -2,170362399 | 0,000000372 | 0,00000936 | yes |
| DSP | ENSG00000096696 | ENSG00000096696 | protein_coding | mCh_JDV-1-3-5 | F1F2_JDV-2-4-6 | 43036,60846 | 9281,809755 | -2,170759902 | 5,55E-27 | 1,44E-24 | yes |
| PP632 | ENSG00000116883 | ENSG00000116883 | protein_coding | mCh_JDV-1-3-5 | F1F2_JDV-2-4-6 | 400,7683574 | 83,01755476 | -2,172481571 | 4,37E-13 | 3,01E-11 | yes |
| S100P | ENSG00000163993 | ENSG00000163993 | protein_coding | mCh_JDV-1-3-5 | F1F2_JDV-2-4-6 | 606,3265267 | 91,63935666 | -2,180746869 | 0,000203292 | 0,002102276 | yes |
| KLK1 | ENSG00000167748 | ENSG00000167748 | protein_coding | mCh_JDV-1-3-5 | F1F2_JDV-2-4-6 | 12,78565953 | 1,351574428 | -2,180816841 | 0,001227893 | 0,008744141 | yes |
| SIRPA | ENSG00000198053 | ENSG00000198053 | protein_coding | mCh_JDV-1-3-5 | F1F2_JDV-2-4-6 | 1928,391421 | 387,3688715 | -2,183433934 | 1,7E-10 | 8,29E-09 | yes |
| RNF128 | ENSG00000133135 | ENSG00000133135 | protein_coding | mCh_JDV-1-3-5 | F1F2_JDV-2-4-6 | 36,01204145 | 5,398572184 | -2,186181569 | 0,00018141 | 0,001911249 | yes |
| C1orf113 | ENSG00000214193 | ENSG00000214193 | protein_coding | mCh_JDV-1-3-5 | F1F2_JDV-2-4-6 | 989,8543641 | 201,9914562 | -2,192217599 | 3,68E-13 | 2,56E-11 | yes |
| HSD11B2 | ENSG00000176387 | ENSG00000176387 | protein_coding | mCh_JDV-1-3-5 | F1F2_JDV-2-4-6 | 108,7668285 | 18,24000128 | -2,19285687 | 0,0000262 | 0,000382898 | yes |
| SYTL1 | ENSG00000142765 | ENSG00000142765 | protein_coding | mCh_JDV-1-3-5 | F1F2_JDV-2-4-6 | 1700,468283 | 324,3857737 | -2,193282985 | 5,98E-08 | 0,00000182 | yes |
| MYCBPAP | ENSG00000136449 | ENSG00000136449 | protein_coding | mCh_JDV-1-3-5 | F1F2_JDV-2-4-6 | 35,01448832 | 5,073993507 | -2,197998991 | 0,000217193 | 0,002209209 | yes |
| F11R | ENSG00000158769 | ENSG00000158769 | protein_coding | mCh_JDV-1-3-5 | F1F2_JDV-2-4-6 | 15205,69897 | 3182,465049 | -2,202906988 | 8,52E-23 | 1,7E-20 | yes |
| ST6GAL1 | ENSG00000073849 | ENSG00000073849 | protein_coding | mCh_JDV-1-3-5 | F1F2_JDV-2-4-6 | 2581,92623 | 530,243012 | -2,204870986 | 2,42E-16 | 2,52E-14 | yes |
| FRMD4B | ENSG00000114541 | ENSG00000114541 | protein_coding | mCh_JDV-1-3-5 | F1F2_JDV-2-4-6 | 1329,111989 | 276,7423465 | -2,210634367 | 3,84E-23 | 8,21E-21 | yes |
| TNFSF10 | ENSG00000121858 | ENSG00000121858 | protein_coding | mCh_JDV-1-3-5 | F1F2_JDV-2-4-6 | 534,5288201 | 89,88807338 | -2,223280036 | 0,00000995 | 0,00016841 | yes |
| RAPGEFL1 | ENSG00000108352 | ENSG00000108352 | protein_coding | mCh_JDV-1-3-5 | F1F2_JDV-2-4-6 | 2014,167578 | 391,2427688 | -2,223689148 | 1,66E-10 | 8,16E-09 | yes |
| GBP6 | ENSG00000183347 | ENSG00000183347 | protein_coding | mCh_JDV-1-3-5 | F1F2_JDV-2-4-6 | 28,34533701 | 3,041440918 | -2,223901181 | 0,000902635 | 0,006909111 | yes |
| PODXL | ENSG00000128567 | ENSG00000128567 | protein_coding | mCh_JDV-1-3-5 | F1F2_JDV-2-4-6 | 2272,121325 | 460,8696029 | -2,229900263 | 2,94E-18 | 3,65E-16 | yes |
| CYP27B1 | ENSG00000111012 | ENSG00000111012 | protein_coding | mCh_JDV-1-3-5 | F1F2_JDV-2-4-6 | 798,7767685 | 159,5013286 | -2,230134126 | 1,43E-14 | 1,19E-12 | yes |
| NOTCH1 | ENSG00000148400 | ENSG00000148400 | protein_coding | mCh_JDV-1-3-5 | F1F2_JDV-2-4-6 | 2980,559383 | 591,3476987 | -2,23042525 | 1,72E-13 | 1,24E-11 | yes |
| RCSD1 | ENSG00000198771 | ENSG00000198771 | protein_coding | mCh_JDV-1-3-5 | F1F2_JDV-2-4-6 | 61,77103213 | 10,77744313 | -2,235878763 | 0,00000124 | 0,0000279 | yes |
| NTSR1 | ENSG00000101188 | ENSG00000101188 | protein_coding | mCh_JDV-1-3-5 | F1F2_JDV-2-4-6 | 110,4117143 | 19,57226189 | -2,242263592 | 0,000000424 | 0,0000105 | yes |

| | | | | | | | | | | | |
|---|---|---|---|---|---|---|---|---|---|---|---|
| YBX2 | ENSG00000006047 | ENSG00000006047 | protein_coding | mCh_JDV-1-3-5 | F1F2_JDV-2-4-6 | 71,1674884 | 11,46614682 | -2,246971069 | 0,0000142 | 0,000228701 | yes |
| S100A2 | ENSG00000196754 | ENSG00000196754 | protein_coding | mCh_JDV-1-3-5 | F1F2_JDV-2-4-6 | 28638,22279 | 5213,595658 | -2,256276159 | 2,31E-08 | 0,000000759 | yes |
| C20orf82 | ENSG00000101230 | ENSG00000101230 | protein_coding | mCh_JDV-1-3-5 | F1F2_JDV-2-4-6 | 142,1527448 | 26,08436032 | -2,260221005 | 6,62E-09 | 0,000000239 | yes |
| PTGS1 | ENSG00000095303 | ENSG00000095303 | protein_coding | mCh_JDV-1-3-5 | F1F2_JDV-2-4-6 | 846,8858408 | 162,2442279 | -2,267179695 | 9,89E-13 | 6,51E-11 | yes |
| EPHB2 | ENSG00000133216 | ENSG00000133216 | protein_coding | mCh_JDV-1-3-5 | F1F2_JDV-2-4-6 | 834,2538482 | 136,4062863 | -2,268649356 | 0,00000506 | 0,0000938 | yes |
| TTC9 | ENSG00000133985 | ENSG00000133985 | protein_coding | mCh_JDV-1-3-5 | F1F2_JDV-2-4-6 | 96,70586307 | 13,92086559 | -2,27036063 | 0,000072 | 0,000888171 | yes |
| STEAP4 | ENSG00000127954 | ENSG00000127954 | protein_coding | mCh_JDV-1-3-5 | F1F2_JDV-2-4-6 | 88,64828964 | 14,5350145 | -2,270978462 | 0,00000379 | 0,0000726 | yes |
| DHRS13 | ENSG00000167536 | ENSG00000167536 | protein_coding | mCh_JDV-1-3-5 | F1F2_JDV-2-4-6 | 444,8503773 | 86,11066157 | -2,276360508 | 1,5E-15 | 1,41E-13 | yes |
| HAP1 | ENSG00000173805 | ENSG00000173805 | protein_coding | mCh_JDV-1-3-5 | F1F2_JDV-2-4-6 | 191,925145 | 35,10032624 | -2,277513811 | 1,5E-09 | 6,09E-08 | yes |
| PLEKHH1 | ENSG00000054690 | ENSG00000054690 | protein_coding | mCh_JDV-1-3-5 | F1F2_JDV-2-4-6 | 525,570616 | 95,41378494 | -2,278536417 | 3,56E-09 | 0,000000135 | yes |
| TMEM51 | ENSG00000171729 | ENSG00000171729 | protein_coding | mCh_JDV-1-3-5 | F1F2_JDV-2-4-6 | 519,1751032 | 96,89834741 | -2,285491796 | 1,64E-11 | 9,52E-10 | yes |
| AGR2 | ENSG00000106541 | ENSG00000106541 | protein_coding | mCh_JDV-1-3-5 | F1F2_JDV-2-4-6 | 38,30264317 | 5,02739677 | -2,294647619 | 0,00012465 | 0,00139071 | yes |
| CORO2B | ENSG00000103647 | ENSG00000103647 | protein_coding | mCh_JDV-1-3-5 | F1F2_JDV-2-4-6 | 66,98980817 | 11,2133227 | -2,295477382 | 0,000000565 | 0,0000136 | yes |
| EPPK1 | ENSG00000227184 | ENSG00000227184 | protein_coding | mCh_JDV-1-3-5 | F1F2_JDV-2-4-6 | 87,72796189 | 14,50811901 | -2,301281218 | 0,000000708 | 0,0000168 | yes |
| SLC2A9 | ENSG00000109667 | ENSG00000109667 | protein_coding | mCh_JDV-1-3-5 | F1F2_JDV-2-4-6 | 1083,889046 | 211,448227 | -2,30449501 | 6,52E-26 | 1,59E-23 | yes |
| GRB14 | ENSG00000115290 | ENSG00000115290 | protein_coding | mCh_JDV-1-3-5 | F1F2_JDV-2-4-6 | 265,0597319 | 49,3769162 | -2,306491832 | 3,44E-13 | 2,4E-11 | yes |
| PCTK3 | ENSG00000117266 | ENSG00000117266 | protein_coding | mCh_JDV-1-3-5 | F1F2_JDV-2-4-6 | 2578,802765 | 461,4008775 | -2,314474902 | 3,95E-10 | 1,85E-08 | yes |
| GSDMB | ENSG00000073605 | ENSG00000073605 | protein_coding | mCh_JDV-1-3-5 | F1F2_JDV-2-4-6 | 408,6477428 | 66,07721843 | -2,317097382 | 0,00000107 | 0,0000245 | yes |
| PNLIPRP3 | ENSG00000203837 | ENSG00000203837 | protein_coding | mCh_JDV-1-3-5 | F1F2_JDV-2-4-6 | 2271,622346 | 140,3896564 | -2,317186228 | 0,001309291 | 0,00918785 | yes |
| LINGO2 | ENSG00000174482 | ENSG00000174482 | protein_coding | mCh_JDV-1-3-5 | F1F2_JDV-2-4-6 | 36,40273827 | 5,74072701 | -2,319369334 | 0,00000197 | 0,0000414 | yes |
| IFI30 | ENSG00000216490 | ENSG00000216490 | protein_coding | mCh_JDV-1-3-5 | F1F2_JDV-2-4-6 | 1579,539328 | 297,6643501 | -2,327700913 | 1,24E-18 | 1,6E-16 | yes |
| JAG1 | ENSG00000101384 | ENSG00000101384 | protein_coding | mCh_JDV-1-3-5 | F1F2_JDV-2-4-6 | 3267,905181 | 600,1259992 | -2,327937695 | 1,35E-13 | 9,8E-12 | yes |
| CD14 | ENSG00000170458 | ENSG00000170458 | protein_coding | mCh_JDV-1-3-5 | F1F2_JDV-2-4-6 | 743,1759413 | 117,721448 | -2,328751583 | 0,00000154 | 0,0000338 | yes |
| KCNQ5 | ENSG00000185760 | ENSG00000185760 | protein_coding | mCh_JDV-1-3-5 | F1F2_JDV-2-4-6 | 683,4163882 | 126,3102435 | -2,330943344 | 6,62E-15 | 5,66E-13 | yes |
| TP53I11 | ENSG00000175274 | ENSG00000175274 | protein_coding | mCh_JDV-1-3-5 | F1F2_JDV-2-4-6 | 341,7369357 | 64,26726764 | -2,334090987 | 1,81E-19 | 2,53E-17 | yes |
| LPAR5 | ENSG00000184574 | ENSG00000184574 | protein_coding | mCh_JDV-1-3-5 | F1F2_JDV-2-4-6 | 152,5196251 | 23,73268564 | -2,347516182 | 0,00000136 | 0,0000302 | yes |
| ADARB2 | ENSG00000185736 | ENSG00000185736 | protein_coding | mCh_JDV-1-3-5 | F1F2_JDV-2-4-6 | 45,07225414 | 6,726051334 | -2,348197146 | 0,0000043 | 0,0000811 | yes |
| FAM84B | ENSG00000168672 | ENSG00000168672 | protein_coding | mCh_JDV-1-3-5 | F1F2_JDV-2-4-6 | 2092,791275 | 369,9765341 | -2,348250974 | 2,36E-11 | 1,31E-09 | yes |
| SP6 | ENSG00000189120 | ENSG00000189120 | protein_coding | mCh_JDV-1-3-5 | F1F2_JDV-2-4-6 | 194,4604171 | 33,14005949 | -2,349733526 | 3,36E-09 | 0,000000129 | yes |
| SLC47A2 | ENSG00000180638 | ENSG00000180638 | protein_coding | mCh_JDV-1-3-5 | F1F2_JDV-2-4-6 | 781,9716065 | 118,4653439 | -2,350924768 | 0,00000318 | 0,0000626 | yes |
| C17orf47 | ENSG00000181013 | ENSG00000181013 | protein_coding | mCh_JDV-1-3-5 | F1F2_JDV-2-4-6 | 22,75832839 | 2,693829508 | -2,356402047 | 0,000113997 | 0,001297291 | yes |
| SIRPB2 | ENSG00000196209 | ENSG00000196209 | protein_coding | mCh_JDV-1-3-5 | F1F2_JDV-2-4-6 | 166,5544057 | 26,03616976 | -2,356787389 | 0,000000744 | 0,0000176 | yes |
| LPCAT3 | ENSG00000172197 | ENSG00000172197 | protein_coding | mCh_JDV-1-3-5 | F1F2_JDV-2-4-6 | 341,5572535 | 59,64977364 | -2,359435813 | 4,02E-11 | 2,17E-09 | yes |
| SEMA6B | ENSG00000167680 | ENSG00000167680 | protein_coding | mCh_JDV-1-3-5 | F1F2_JDV-2-4-6 | 65,93334527 | 10,77638058 | -2,367728887 | 2,46E-08 | 0,000000805 | yes |
| PTHLH | ENSG00000087494 | ENSG00000087494 | protein_coding | mCh_JDV-1-3-5 | F1F2_JDV-2-4-6 | 35,15580479 | 5,092632201 | -2,368854548 | 0,00000513 | 0,0000949 | yes |
| KCNS3 | ENSG00000170745 | ENSG00000170745 | protein_coding | mCh_JDV-1-3-5 | F1F2_JDV-2-4-6 | 366,0199027 | 63,73419264 | -2,38572968 | 8,14E-13 | 5,43E-11 | yes |
| MYBPHL | ENSG00000221986 | ENSG00000221986 | protein_coding | mCh_JDV-1-3-5 | F1F2_JDV-2-4-6 | 23,16467318 | 3,036515607 | -2,387364125 | 0,0000201 | 0,000305937 | yes |
| TXNDC3 | ENSG00000086288 | ENSG00000086288 | protein_coding | mCh_JDV-1-3-5 | F1F2_JDV-2-4-6 | 36,26909859 | 3,364425775 | -2,394086024 | 0,000317409 | 0,003015619 | yes |
| SMPDL3B | ENSG00000130768 | ENSG00000130768 | protein_coding | mCh_JDV-1-3-5 | F1F2_JDV-2-4-6 | 737,5657214 | 127,7063244 | -2,399206943 | 1,69E-13 | 1,22E-11 | yes |
| PERP | ENSG00000112378 | ENSG00000112378 | protein_coding | mCh_JDV-1-3-5 | F1F2_JDV-2-4-6 | 38388,62004 | 6910,61156 | -2,404412737 | 4,9E-23 | 1,02E-20 | yes |
| SLITRK4 | ENSG00000179542 | ENSG00000179542 | protein_coding | mCh_JDV-1-3-5 | F1F2_JDV-2-4-6 | 170,5879523 | 29,38635088 | -2,407847195 | 9,29E-14 | 6,94E-12 | yes |
| AIF1L | ENSG00000126878 | ENSG00000126878 | protein_coding | mCh_JDV-1-3-5 | F1F2_JDV-2-4-6 | 398,6939242 | 67,91421324 | -2,417547005 | 2,43E-13 | 1,73E-11 | yes |
| TMC6 | ENSG00000141524 | ENSG00000141524 | protein_coding | mCh_JDV-1-3-5 | F1F2_JDV-2-4-6 | 129,5609435 | 19,51088922 | -2,425426268 | 0,000000148 | 0,00000413 | yes |
| NCCRP1 | ENSG00000188505 | ENSG00000188505 | protein_coding | mCh_JDV-1-3-5 | F1F2_JDV-2-4-6 | 33,07089591 | 3,388521054 | -2,427247124 | 0,000134791 | 0,001491777 | yes |
| MAPK13 | ENSG00000156711 | ENSG00000156711 | protein_coding | mCh_JDV-1-3-5 | F1F2_JDV-2-4-6 | 472,4209016 | 81,02387338 | -2,431489911 | 7,59E-16 | 7,6E-14 | yes |
| LIPG | ENSG00000101670 | ENSG00000101670 | protein_coding | mCh_JDV-1-3-5 | F1F2_JDV-2-4-6 | 1012,737709 | 139,989059 | -2,431879613 | 0,00000318 | 0,0000626 | yes |
| EOMES | ENSG00000163508 | ENSG00000163508 | protein_coding | mCh_JDV-1-3-5 | F1F2_JDV-2-4-6 | 36,5953647 | 3,690067 | -2,435172806 | 0,000134474 | 0,001489095 | yes |

| | | | | | | | | | | | |
|---|---|---|---|---|---|---|---|---|---|---|---|
| BBOX1 | ENSG00000129151 | ENSG00000129151 | protein_coding | mCh_JDV-1-3-5 | F1F2_JDV-2-4-6 | 157,6333081 | 17,84086783 | -2,45249042 | 0,0000507 | 0,000667888 | yes |
| RAB17 | ENSG00000124839 | ENSG00000124839 | protein_coding | mCh_JDV-1-3-5 | F1F2_JDV-2-4-6 | 1360,74404 | 220,1080764 | -2,457888379 | 1,16E-11 | 6,85E-10 | yes |
| TNNT1 | ENSG00000105048 | ENSG00000105048 | protein_coding | mCh_JDV-1-3-5 | F1F2_JDV-2-4-6 | 231,3447054 | 36,21544631 | -2,458084168 | 9,47E-10 | 4,07E-08 | yes |
| WNK4 | ENSG00000126562 | ENSG00000126562 | protein_coding | mCh_JDV-1-3-5 | F1F2_JDV-2-4-6 | 104,4779546 | 12,74862305 | -2,466420082 | 0,0000149 | 0,000238295 | yes |
| ANK3 | ENSG00000151150 | ENSG00000151150 | protein_coding | mCh_JDV-1-3-5 | F1F2_JDV-2-4-6 | 1427,021039 | 239,7971955 | -2,473285923 | 3,27E-18 | 4,03E-16 | yes |
| C6orf141 | ENSG00000197261 | ENSG00000197261 | protein_coding | mCh_JDV-1-3-5 | F1F2_JDV-2-4-6 | 571,0598176 | 94,87564078 | -2,473305978 | 4,76E-16 | 4,92E-14 | yes |
| GJB2 | ENSG00000165474 | ENSG00000165474 | protein_coding | mCh_JDV-1-3-5 | F1F2_JDV-2-4-6 | 938,0602852 | 157,9502942 | -2,480991218 | 3,52E-20 | 5,19E-18 | yes |
| ERBB3 | ENSG00000065361 | ENSG00000065361 | protein_coding | mCh_JDV-1-3-5 | F1F2_JDV-2-4-6 | 2134,197833 | 334,7637401 | -2,484632581 | 4,02E-11 | 2,17E-09 | yes |
| SYK | ENSG00000165025 | ENSG00000165025 | protein_coding | mCh_JDV-1-3-5 | F1F2_JDV-2-4-6 | 174,7328059 | 26,37726204 | -2,492998709 | 1,36E-09 | 5,57E-08 | yes |
| GALNTL4 | ENSG00000110328 | ENSG00000110328 | protein_coding | mCh_JDV-1-3-5 | F1F2_JDV-2-4-6 | 919,6130053 | 146,3383739 | -2,495813524 | 5,82E-13 | 3,98E-11 | yes |
| NTF4 | ENSG00000225950 | ENSG00000225950 | protein_coding | mCh_JDV-1-3-5 | F1F2_JDV-2-4-6 | 38,79405318 | 3,347380901 | -2,496986402 | 0,000148889 | 0,001621728 | yes |
| DAPP1 | ENSG00000070190 | ENSG00000070190 | protein_coding | mCh_JDV-1-3-5 | F1F2_JDV-2-4-6 | 346,2800134 | 54,86829443 | -2,500453911 | 6,19E-13 | 4,21E-11 | yes |
| CCND2 | ENSG00000118971 | ENSG00000118971 | protein_coding | mCh_JDV-1-3-5 | F1F2_JDV-2-4-6 | 1405,821607 | 227,4313757 | -2,504690428 | 8,13E-16 | 8,06E-14 | yes |
| UGT8 | ENSG00000174607 | ENSG00000174607 | protein_coding | mCh_JDV-1-3-5 | F1F2_JDV-2-4-6 | 999,8652583 | 160,3159305 | -2,510797146 | 3,41E-15 | 3,03E-13 | yes |
| ARL4C | ENSG00000188042 | ENSG00000188042 | protein_coding | mCh_JDV-1-3-5 | F1F2_JDV-2-4-6 | 110,2853983 | 13,62690132 | -2,513186968 | 0,00000412 | 0,0000783 | yes |
| VSNL1 | ENSG00000163032 | ENSG00000163032 | protein_coding | mCh_JDV-1-3-5 | F1F2_JDV-2-4-6 | 323,7039852 | 47,98593929 | -2,515672733 | 1,12E-09 | 4,65E-08 | yes |
| EXPH5 | ENSG00000110723 | ENSG00000110723 | protein_coding | mCh_JDV-1-3-5 | F1F2_JDV-2-4-6 | 465,2599508 | 73,16938068 | -2,519612051 | 9,32E-14 | 6,94E-12 | yes |
| CD9 | ENSG00000010278 | ENSG00000010278 | protein_coding | mCh_JDV-1-3-5 | F1F2_JDV-2-4-6 | 7391,298785 | 1207,175575 | -2,532931981 | 3,87E-23 | 8,21E-21 | yes |
| SLC29A2 | ENSG00000174669 | ENSG00000174669 | protein_coding | mCh_JDV-1-3-5 | F1F2_JDV-2-4-6 | 971,481406 | 154,0978311 | -2,53982826 | 3,37E-17 | 3,82E-15 | yes |
| EFNB1 | ENSG00000090776 | ENSG00000090776 | protein_coding | mCh_JDV-1-3-5 | F1F2_JDV-2-4-6 | 2061,634386 | 334,6796919 | -2,542325809 | 1,5E-23 | 3,44E-21 | yes |
| KCNK6 | ENSG00000099337 | ENSG00000099337 | protein_coding | mCh_JDV-1-3-5 | F1F2_JDV-2-4-6 | 192,5526224 | 29,81450493 | -2,543719291 | 3,34E-14 | 2,64E-12 | yes |
| MST1R | ENSG00000164078 | ENSG00000164078 | protein_coding | mCh_JDV-1-3-5 | F1F2_JDV-2-4-6 | 2624,712208 | 424,0887249 | -2,544340961 | 1,85E-22 | 3,54E-20 | yes |
| PRG2 | ENSG00000186652 | ENSG00000186652 | protein_coding | mCh_JDV-1-3-5 | F1F2_JDV-2-4-6 | 50,5328107 | 6,429430698 | -2,552348704 | 0,000000501 | 0,0000122 | yes |
| CHI3L1 | ENSG00000133048 | ENSG00000133048 | protein_coding | mCh_JDV-1-3-5 | F1F2_JDV-2-4-6 | 1265,328721 | 16,83463586 | -2,553174705 | 0,000528577 | 0,004508335 | yes |
| PPP1R14C | ENSG00000198729 | ENSG00000198729 | protein_coding | mCh_JDV-1-3-5 | F1F2_JDV-2-4-6 | 663,6654844 | 98,38840126 | -2,554261554 | 2,13E-11 | 1,21E-09 | yes |
| GSDMA | ENSG00000167914 | ENSG00000167914 | protein_coding | mCh_JDV-1-3-5 | F1F2_JDV-2-4-6 | 33,59270961 | 4,060179867 | -2,561856242 | 0,00000133 | 0,0000297 | yes |
| LTB4R | ENSG00000213903 | ENSG00000213903 | protein_coding | mCh_JDV-1-3-5 | F1F2_JDV-2-4-6 | 3117,204847 | 434,0187269 | -2,562687228 | 5,53E-09 | 0,000000203 | yes |
| CEACAM1 | ENSG00000079385 | ENSG00000079385 | protein_coding | mCh_JDV-1-3-5 | F1F2_JDV-2-4-6 | 65,98969611 | 9,104621512 | -2,573831177 | 3,24E-09 | 0,000000125 | yes |
| SLC6A11 | ENSG00000132164 | ENSG00000132164 | protein_coding | mCh_JDV-1-3-5 | F1F2_JDV-2-4-6 | 163,7445536 | 23,69980229 | -2,57709668 | 3,32E-11 | 1,81E-09 | yes |
| LLGL2 | ENSG00000073350 | ENSG00000073350 | protein_coding | mCh_JDV-1-3-5 | F1F2_JDV-2-4-6 | 689,9462206 | 98,80244984 | -2,593236789 | 2,22E-11 | 1,24E-09 | yes |
| FGR | ENSG00000000938 | ENSG00000000938 | protein_coding | mCh_JDV-1-3-5 | F1F2_JDV-2-4-6 | 173,9416066 | 17,45969794 | -2,596960056 | 0,000016 | 0,000252105 | yes |
| CXCR2 | ENSG00000180871 | ENSG00000180871 | protein_coding | mCh_JDV-1-3-5 | F1F2_JDV-2-4-6 | 36,95787243 | 3,031590297 | -2,598057924 | 0,0000603 | 0,000771667 | yes |
| ARHGEF4 | ENSG00000136002 | ENSG00000136002 | protein_coding | mCh_JDV-1-3-5 | F1F2_JDV-2-4-6 | 5269,821473 | 777,6967157 | -2,598329594 | 6,19E-14 | 4,73E-12 | yes |
| CES1 | ENSG00000198848 | ENSG00000198848 | protein_coding | mCh_JDV-1-3-5 | F1F2_JDV-2-4-6 | 78,05788093 | 8,800275349 | -2,607872028 | 0,00000212 | 0,0000441 | yes |
| CXCR1 | ENSG00000163464 | ENSG00000163464 | protein_coding | mCh_JDV-1-3-5 | F1F2_JDV-2-4-6 | 744,1463057 | 4,729182314 | -2,609600895 | 0,000397084 | 0,00361423 | yes |
| TRAF3IP3 | ENSG00000009790 | ENSG00000009790 | protein_coding | mCh_JDV-1-3-5 | F1F2_JDV-2-4-6 | 644,2083725 | 96,71196046 | -2,613200246 | 9,22E-18 | 1,1E-15 | yes |
| S100A9 | ENSG00000163220 | ENSG00000163220 | protein_coding | mCh_JDV-1-3-5 | F1F2_JDV-2-4-6 | 7268,159407 | 817,6460212 | -2,618873032 | 0,00000195 | 0,0000409 | yes |
| SYTL2 | ENSG00000137501 | ENSG00000137501 | protein_coding | mCh_JDV-1-3-5 | F1F2_JDV-2-4-6 | 213,338327 | 28,73506843 | -2,631722003 | 3,68E-10 | 1,73E-08 | yes |
| METRNL | ENSG00000176845 | ENSG00000176845 | protein_coding | mCh_JDV-1-3-5 | F1F2_JDV-2-4-6 | 70,08580314 | 7,076994233 | -2,659355811 | 0,00000401 | 0,0000765 | yes |
| GPC4 | ENSG00000076716 | ENSG00000076716 | protein_coding | mCh_JDV-1-3-5 | F1F2_JDV-2-4-6 | 768,8953976 | 109,4502063 | -2,664576882 | 5,5E-16 | 5,59E-14 | yes |
| GDPD3 | ENSG00000102886 | ENSG00000102886 | protein_coding | mCh_JDV-1-3-5 | F1F2_JDV-2-4-6 | 154,5254245 | 17,8845205 | -2,665051157 | 0,000000199 | 0,00000535 | yes |
| SLC3A2 | ENSG00000168003 | ENSG00000168003 | protein_coding | mCh_JDV-1-3-5 | F1F2_JDV-2-4-6 | 20755,19499 | 3020,842168 | -2,674182222 | 2,88E-21 | 4,86E-19 | yes |
| CATSPERB | ENSG00000133962 | ENSG00000133962 | protein_coding | mCh_JDV-1-3-5 | F1F2_JDV-2-4-6 | 65,04853906 | 6,071968668 | -2,677532832 | 0,00000772 | 0,00013535 | yes |
| CLMN | ENSG00000165959 | ENSG00000165959 | protein_coding | mCh_JDV-1-3-5 | F1F2_JDV-2-4-6 | 1181,970786 | 165,4844978 | -2,678055685 | 2,21E-15 | 2,02E-13 | yes |
| ITGA2 | ENSG00000164171 | ENSG00000164171 | protein_coding | mCh_JDV-1-3-5 | F1F2_JDV-2-4-6 | 8221,477816 | 1166,908869 | -2,678876826 | 3,46E-17 | 3,9E-15 | yes |
| PLLP | ENSG00000102934 | ENSG00000102934 | protein_coding | mCh_JDV-1-3-5 | F1F2_JDV-2-4-6 | 530,496469 | 67,77548557 | -2,685757506 | 4,77E-10 | 2,19E-08 | yes |
| CYP4F3 | ENSG00000186529 | ENSG00000186529 | protein_coding | mCh_JDV-1-3-5 | F1F2_JDV-2-4-6 | 136,1113152 | 16,49967529 | -2,688975312 | 9,97E-09 | 0,000000346 | yes |

| | | | | | | | | | | | |
|---|---|---|---|---|---|---|---|---|---|---|---|
| GPR110 | ENSG00000153292 | ENSG00000153292 | protein_coding | mCh_JDV-1-3-5 | F1F2_JDV-2-4-6 | 303,5368175 | 37,41086077 | -2,692826397 | 3,5E-09 | 0,000000134 | yes |
| PLB1 | ENSG00000163803 | ENSG00000163803 | protein_coding | mCh_JDV-1-3-5 | F1F2_JDV-2-4-6 | 57,73168814 | 7,111471406 | -2,693491295 | 2,72E-09 | 0,000000107 | yes |
| PRR15L | ENSG00000167183 | ENSG00000167183 | protein_coding | mCh_JDV-1-3-5 | F1F2_JDV-2-4-6 | 670,5716788 | 82,0824417 | -2,719742013 | 1,02E-09 | 4,33E-08 | yes |
| SOX9 | ENSG00000125398 | ENSG00000125398 | protein_coding | mCh_JDV-1-3-5 | F1F2_JDV-2-4-6 | 266,8006981 | 28,39943274 | -2,719747942 | 0,000000365 | 0,00000921 | yes |
| ATP10B | ENSG00000118322 | ENSG00000118322 | protein_coding | mCh_JDV-1-3-5 | F1F2_JDV-2-4-6 | 95,65700439 | 11,51433737 | -2,721093173 | 2,4E-09 | 9,52E-08 | yes |
| ESYT3 | ENSG00000158220 | ENSG00000158220 | protein_coding | mCh_JDV-1-3-5 | F1F2_JDV-2-4-6 | 150,7552392 | 19,97738321 | -2,723146755 | 8,23E-14 | 6,2E-12 | yes |
| ACP5 | ENSG00000102575 | ENSG00000102575 | protein_coding | mCh_JDV-1-3-5 | F1F2_JDV-2-4-6 | 12,12487088 | 0,333366752 | -2,730887296 | 0,000123343 | 0,001379774 | yes |
| TNFSF15 | ENSG00000181634 | ENSG00000181634 | protein_coding | mCh_JDV-1-3-5 | F1F2_JDV-2-4-6 | 53,51265281 | 6,069699725 | -2,74156278 | 1,26E-08 | 0,000000432 | yes |
| SH3RF2 | ENSG00000156463 | ENSG00000156463 | protein_coding | mCh_JDV-1-3-5 | F1F2_JDV-2-4-6 | 1201,189626 | 157,1767977 | -2,747443293 | 2,02E-14 | 1,64E-12 | yes |
| PRSS12 | ENSG00000164099 | ENSG00000164099 | protein_coding | mCh_JDV-1-3-5 | F1F2_JDV-2-4-6 | 1697,504162 | 230,2822537 | -2,758975326 | 1,98E-20 | 3,03E-18 | yes |
| SFRP1 | ENSG00000104332 | ENSG00000104332 | protein_coding | mCh_JDV-1-3-5 | F1F2_JDV-2-4-6 | 23008,31029 | 2940,878357 | -2,760944434 | 1,8E-13 | 1,29E-11 | yes |
| SLC27A3 | ENSG00000143554 | ENSG00000143554 | protein_coding | mCh_JDV-1-3-5 | F1F2_JDV-2-4-6 | 2851,713382 | 388,0183815 | -2,763312045 | 7,52E-22 | 1,39E-19 | yes |
| ALS2CL | ENSG00000178038 | ENSG00000178038 | protein_coding | mCh_JDV-1-3-5 | F1F2_JDV-2-4-6 | 11275,92618 | 1413,403905 | -2,76897767 | 1,18E-12 | 7,73E-11 | yes |
| CASZ1 | ENSG00000130940 | ENSG00000130940 | protein_coding | mCh_JDV-1-3-5 | F1F2_JDV-2-4-6 | 466,1278542 | 61,11853244 | -2,780567675 | 1,02E-17 | 1,19E-15 | yes |
| UNC5B | ENSG00000107731 | ENSG00000107731 | protein_coding | mCh_JDV-1-3-5 | F1F2_JDV-2-4-6 | 1919,870948 | 212,4186107 | -2,780942171 | 9,78E-09 | 0,00000034 | yes |
| MARVELD2 | ENSG00000152939 | ENSG00000152939 | protein_coding | mCh_JDV-1-3-5 | F1F2_JDV-2-4-6 | 1176,524976 | 154,7259416 | -2,801310832 | 5,48E-21 | 8,93E-19 | yes |
| BLNK | ENSG00000095585 | ENSG00000095585 | protein_coding | mCh_JDV-1-3-5 | F1F2_JDV-2-4-6 | 118,485067 | 10,76107338 | -2,810945041 | 0,00000077 | 0,0000182 | yes |
| ITGA6 | ENSG00000091409 | ENSG00000091409 | protein_coding | mCh_JDV-1-3-5 | F1F2_JDV-2-4-6 | 29302,90789 | 3843,585814 | -2,818893403 | 1,59E-23 | 3,6E-21 | yes |
| RORC | ENSG00000143365 | ENSG00000143365 | protein_coding | mCh_JDV-1-3-5 | F1F2_JDV-2-4-6 | 389,1906582 | 34,29273998 | -2,82276497 | 0,00000108 | 0,0000247 | yes |
| TACSTD2 | ENSG00000184292 | ENSG00000184292 | protein_coding | mCh_JDV-1-3-5 | F1F2_JDV-2-4-6 | 7891,852699 | 972,8355544 | -2,829711092 | 2,62E-15 | 2,36E-13 | yes |
| ANXA8L2 | ENSG00000186807 | ENSG00000186807 | protein_coding | mCh_JDV-1-3-5 | F1F2_JDV-2-4-6 | 109,9953286 | 12,54080185 | -2,83022631 | 5,93E-11 | 3,11E-09 | yes |
| STAT4 | ENSG00000138378 | ENSG00000138378 | protein_coding | mCh_JDV-1-3-5 | F1F2_JDV-2-4-6 | 169,3049129 | 19,25806511 | -2,835560303 | 4,78E-11 | 2,53E-09 | yes |
| TGM1 | ENSG00000092295 | ENSG00000092295 | protein_coding | mCh_JDV-1-3-5 | F1F2_JDV-2-4-6 | 1601,272337 | 166,7246763 | -2,859042178 | 3,52E-09 | 0,000000134 | yes |
| SBSN | ENSG00000189001 | ENSG00000189001 | protein_coding | mCh_JDV-1-3-5 | F1F2_JDV-2-4-6 | 160,7427229 | 2,375238688 | -2,867231141 | 0,0000893 | 0,00106176 | yes |
| SAA2 | ENSG00000134339 | ENSG00000134339 | protein_coding | mCh_JDV-1-3-5 | F1F2_JDV-2-4-6 | 4936,765946 | 459,3624579 | -2,871595677 | 0,000000107 | 0,00000309 | yes |
| ANO9 | ENSG00000185101 | ENSG00000185101 | protein_coding | mCh_JDV-1-3-5 | F1F2_JDV-2-4-6 | 2107,330476 | 234,2301444 | -2,876215044 | 1,48E-11 | 8,61E-10 | yes |
| GALNT3 | ENSG00000115339 | ENSG00000115339 | protein_coding | mCh_JDV-1-3-5 | F1F2_JDV-2-4-6 | 3273,792719 | 399,8902552 | -2,884185769 | 1,34E-19 | 1,89E-17 | yes |
| CDS1 | ENSG00000163624 | ENSG00000163624 | protein_coding | mCh_JDV-1-3-5 | F1F2_JDV-2-4-6 | 2372,612116 | 302,1414831 | -2,889373796 | 2,24E-32 | 9,03E-30 | yes |
| FZD4 | ENSG00000174804 | ENSG00000174804 | protein_coding | mCh_JDV-1-3-5 | F1F2_JDV-2-4-6 | 1080,098919 | 111,6123017 | -2,890460623 | 9,66E-10 | 4,14E-08 | yes |
| ZBED2 | ENSG00000177494 | ENSG00000177494 | protein_coding | mCh_JDV-1-3-5 | F1F2_JDV-2-4-6 | 465,3149698 | 50,39927434 | -2,901647756 | 1,65E-11 | 9,52E-10 | yes |
| HR | ENSG00000168453 | ENSG00000168453 | protein_coding | mCh_JDV-1-3-5 | F1F2_JDV-2-4-6 | 947,4245307 | 115,9422067 | -2,90440517 | 4,35E-23 | 9,14E-21 | yes |
| KCNC4 | ENSG00000116396 | ENSG00000116396 | protein_coding | mCh_JDV-1-3-5 | F1F2_JDV-2-4-6 | 85,01769982 | 9,471934163 | -2,912978758 | 1,75E-13 | 1,25E-11 | yes |
| FRAS1 | ENSG00000138759 | ENSG00000138759 | protein_coding | mCh_JDV-1-3-5 | F1F2_JDV-2-4-6 | 1162,241317 | 126,9920012 | -2,918381101 | 1,63E-12 | 1,05E-10 | yes |
| PRRG2 | ENSG00000126460 | ENSG00000126460 | protein_coding | mCh_JDV-1-3-5 | F1F2_JDV-2-4-6 | 212,3881117 | 23,60622139 | -2,928914041 | 4,16E-14 | 3,26E-12 | yes |
| S100A8 | ENSG00000143546 | ENSG00000143546 | protein_coding | mCh_JDV-1-3-5 | F1F2_JDV-2-4-6 | 5048,130306 | 423,5803833 | -2,944288111 | 0,000000137 | 0,00000386 | yes |
| CELSR2 | ENSG00000143126 | ENSG00000143126 | protein_coding | mCh_JDV-1-3-5 | F1F2_JDV-2-4-6 | 6343,206797 | 733,131675 | -2,946884213 | 5,47E-19 | 7,45E-17 | yes |
| SAA1 | ENSG00000173432 | ENSG00000173432 | protein_coding | mCh_JDV-1-3-5 | F1F2_JDV-2-4-6 | 13271,02642 | 1244,592723 | -2,954562244 | 3,83E-09 | 0,000000144 | yes |
| VIPR1 | ENSG00000114812 | ENSG00000114812 | protein_coding | mCh_JDV-1-3-5 | F1F2_JDV-2-4-6 | 53,90609424 | 5,069068196 | -2,965262208 | 1,05E-09 | 4,45E-08 | yes |
| ITGB6 | ENSG00000115221 | ENSG00000115221 | protein_coding | mCh_JDV-1-3-5 | F1F2_JDV-2-4-6 | 2651,906247 | 307,034124 | -2,965510973 | 1,34E-21 | 2,35E-19 | yes |
| MACC1 | ENSG00000183742 | ENSG00000183742 | protein_coding | mCh_JDV-1-3-5 | F1F2_JDV-2-4-6 | 169,8223699 | 13,16919542 | -2,968201323 | 0,000000274 | 0,00000716 | yes |
| CCDC64 | ENSG00000135127 | ENSG00000135127 | protein_coding | mCh_JDV-1-3-5 | F1F2_JDV-2-4-6 | 2579,510705 | 79,84863117 | -2,969686665 | 0,0000304 | 0,000435278 | yes |
| KCNK5 | ENSG00000164626 | ENSG00000164626 | protein_coding | mCh_JDV-1-3-5 | F1F2_JDV-2-4-6 | 1054,980124 | 113,6924185 | -2,972088321 | 1,89E-14 | 1,54E-12 | yes |
| SERINC2 | ENSG00000168528 | ENSG00000168528 | protein_coding | mCh_JDV-1-3-5 | F1F2_JDV-2-4-6 | 5076,469399 | 579,3420066 | -2,9860792 | 5,39E-22 | 1E-19 | yes |
| PALMD | ENSG00000099260 | ENSG00000099260 | protein_coding | mCh_JDV-1-3-5 | F1F2_JDV-2-4-6 | 31,59758074 | 2,374707414 | -2,997345157 | 0,000000126 | 0,00000358 | yes |
| SPINK5 | ENSG00000133710 | ENSG00000133710 | protein_coding | mCh_JDV-1-3-5 | F1F2_JDV-2-4-6 | 150,0410483 | 15,56746684 | -3,002350912 | 6,15E-14 | 4,73E-12 | yes |
| NOTCH3 | ENSG00000074181 | ENSG00000074181 | protein_coding | mCh_JDV-1-3-5 | F1F2_JDV-2-4-6 | 455,426986 | 43,10098444 | -3,004827581 | 1,7E-10 | 8,29E-09 | yes |
| AFAP1L2 | ENSG00000169129 | ENSG00000169129 | protein_coding | mCh_JDV-1-3-5 | F1F2_JDV-2-4-6 | 94,46112733 | 8,486609839 | -3,005276458 | 1,61E-09 | 6,52E-08 | yes |

| | | | | | | | | | | | |
|---|---|---|---|---|---|---|---|---|---|---|---|
| C6orf205 | ENSG00000204544 | ENSG00000204544 | protein_coding | mCh_JDV-1-3-5 | F1F2_JDV-2-4-6 | 51,70602433 | 2,365388067 | -3,008225671 | 0,00000733 | 0,000129245 | yes |
| KRT6B | ENSG00000185479 | ENSG00000185479 | protein_coding | mCh_JDV-1-3-5 | F1F2_JDV-2-4-6 | 94,57967606 | 8,466908597 | -3,018961516 | 9,56E-10 | 0,000000041 | yes |
| CDC42BPG | ENSG00000171219 | ENSG00000171219 | protein_coding | mCh_JDV-1-3-5 | F1F2_JDV-2-4-6 | 1276,231564 | 126,5242312 | -3,02229953 | 1,66E-12 | 1,06E-10 | yes |
| P2RY2 | ENSG00000175591 | ENSG00000175591 | protein_coding | mCh_JDV-1-3-5 | F1F2_JDV-2-4-6 | 205,2994813 | 20,01958591 | -3,04708056 | 7,49E-13 | 5,01E-11 | yes |
| AQP3 | ENSG00000165272 | ENSG00000165272 | protein_coding | mCh_JDV-1-3-5 | F1F2_JDV-2-4-6 | 1147,891557 | 109,8706696 | -3,047380167 | 4,49E-12 | 2,73E-10 | yes |
| DMKN | ENSG00000161249 | ENSG00000161249 | protein_coding | mCh_JDV-1-3-5 | F1F2_JDV-2-4-6 | 1841,136486 | 200,4696256 | -3,05110018 | 5,88E-23 | 1,21E-20 | yes |
| PPFIBP2 | ENSG00000166387 | ENSG00000166387 | protein_coding | mCh_JDV-1-3-5 | F1F2_JDV-2-4-6 | 323,0896058 | 33,0287586 | -3,07140655 | 3,92E-17 | 4,39E-15 | yes |
| NHS | ENSG00000188158 | ENSG00000188158 | protein_coding | mCh_JDV-1-3-5 | F1F2_JDV-2-4-6 | 213,9064323 | 21,04658166 | -3,090726592 | 2,07E-15 | 1,91E-13 | yes |
| CLDN7 | ENSG00000181885 | ENSG00000181885 | protein_coding | mCh_JDV-1-3-5 | F1F2_JDV-2-4-6 | 5861,696293 | 605,5328121 | -3,097475658 | 1,53E-20 | 2,38E-18 | yes |
| TMEM40 | ENSG00000088726 | ENSG00000088726 | protein_coding | mCh_JDV-1-3-5 | F1F2_JDV-2-4-6 | 337,1632369 | 35,05479205 | -3,098470705 | 7,86E-22 | 1,44E-19 | yes |
| LPAR3 | ENSG00000171517 | ENSG00000171517 | protein_coding | mCh_JDV-1-3-5 | F1F2_JDV-2-4-6 | 53,94232236 | 4,397940656 | -3,110411772 | 4,68E-10 | 2,16E-08 | yes |
| A2ML1 | ENSG00000166535 | ENSG00000166535 | protein_coding | mCh_JDV-1-3-5 | F1F2_JDV-2-4-6 | 341,788409 | 8,109446567 | -3,113297497 | 0,0000129 | 0,000210578 | yes |
| SORL1 | ENSG00000137642 | ENSG00000137642 | protein_coding | mCh_JDV-1-3-5 | F1F2_JDV-2-4-6 | 3158,343399 | 308,0014314 | -3,118282434 | 1,91E-16 | 2E-14 | yes |
| FAM49A | ENSG00000197872 | ENSG00000197872 | protein_coding | mCh_JDV-1-3-5 | F1F2_JDV-2-4-6 | 40,07423015 | 3,041440918 | -3,120773909 | 2,74E-09 | 0,000000108 | yes |
| SLC2A3 | ENSG00000059804 | ENSG00000059804 | protein_coding | mCh_JDV-1-3-5 | F1F2_JDV-2-4-6 | 407,2695979 | 34,80182409 | -3,125243318 | 4,7E-11 | 2,5E-09 | yes |
| CDH16 | ENSG00000166589 | ENSG00000166589 | protein_coding | mCh_JDV-1-3-5 | F1F2_JDV-2-4-6 | 54,30511096 | 2,707542893 | -3,14399996 | 0,000000825 | 0,0000192 | yes |
| MAPK4 | ENSG00000141639 | ENSG00000141639 | protein_coding | mCh_JDV-1-3-5 | F1F2_JDV-2-4-6 | 74,58749403 | 4,712668713 | -3,148330727 | 7,94E-08 | 0,00000238 | yes |
| STYK1 | ENSG00000060140 | ENSG00000060140 | protein_coding | mCh_JDV-1-3-5 | F1F2_JDV-2-4-6 | 101,4513725 | 8,787093238 | -3,151053785 | 2,27E-12 | 1,44E-10 | yes |
| SFTPD | ENSG00000133661 | ENSG00000133661 | protein_coding | mCh_JDV-1-3-5 | F1F2_JDV-2-4-6 | 42,96390268 | 1,684409906 | -3,171022667 | 0,0000018 | 0,0000383 | yes |
| COL13A1 | ENSG00000197467 | ENSG00000197467 | protein_coding | mCh_JDV-1-3-5 | F1F2_JDV-2-4-6 | 67,05184301 | 5,106876859 | -3,174720619 | 4,79E-10 | 2,19E-08 | yes |
| CA2 | ENSG00000104267 | ENSG00000104267 | protein_coding | mCh_JDV-1-3-5 | F1F2_JDV-2-4-6 | 748,0926422 | 17,66897377 | -3,180315607 | 0,00000768 | 0,000134817 | yes |
| CD247 | ENSG00000198821 | ENSG00000198821 | protein_coding | mCh_JDV-1-3-5 | F1F2_JDV-2-4-6 | 62,43986892 | 1,694260527 | -3,188489225 | 0,0000047 | 0,0000879 | yes |
| SDC1 | ENSG00000115884 | ENSG00000115884 | protein_coding | mCh_JDV-1-3-5 | F1F2_JDV-2-4-6 | 11707,52699 | 1110,363044 | -3,201659264 | 1,26E-20 | 1,98E-18 | yes |
| KLK14 | ENSG00000129437 | ENSG00000129437 | protein_coding | mCh_JDV-1-3-5 | F1F2_JDV-2-4-6 | 79,71403073 | 6,069168451 | -3,215746178 | 6,45E-11 | 3,36E-09 | yes |
| CGB7 | ENSG00000196337 | ENSG00000196337 | protein_coding | mCh_JDV-1-3-5 | F1F2_JDV-2-4-6 | 58,23079756 | 2,693298235 | -3,228538796 | 0,000000383 | 0,0000096 | yes |
| RAPGEF5 | ENSG00000136237 | ENSG00000136237 | protein_coding | mCh_JDV-1-3-5 | F1F2_JDV-2-4-6 | 428,529171 | 39,01178864 | -3,241630327 | 8,1E-20 | 1,17E-17 | yes |
| DHRS3 | ENSG00000162496 | ENSG00000162496 | protein_coding | mCh_JDV-1-3-5 | F1F2_JDV-2-4-6 | 1542,367824 | 106,5056383 | -3,243702688 | 1,38E-09 | 5,65E-08 | yes |
| IL1B | ENSG00000125538 | ENSG00000125538 | protein_coding | mCh_JDV-1-3-5 | F1F2_JDV-2-4-6 | 477,1689157 | 40,10387597 | -3,257176685 | 5,49E-15 | 4,78E-13 | yes |
| LRG1 | ENSG00000171236 | ENSG00000171236 | protein_coding | mCh_JDV-1-3-5 | F1F2_JDV-2-4-6 | 222,7723654 | 17,20861149 | -3,262053817 | 3,64E-12 | 2,26E-10 | yes |
| SERPINB2 | ENSG00000197632 | ENSG00000197632 | protein_coding | mCh_JDV-1-3-5 | F1F2_JDV-2-4-6 | 266,9809103 | 2,037477899 | -3,267859756 | 0,00000735 | 0,000129565 | yes |
| FST | ENSG00000134363 | ENSG00000134363 | protein_coding | mCh_JDV-1-3-5 | F1F2_JDV-2-4-6 | 125,5776516 | 7,766760468 | -3,285285547 | 4,05E-09 | 0,000000151 | yes |
| B4GALNT3 | ENSG00000139044 | ENSG00000139044 | protein_coding | mCh_JDV-1-3-5 | F1F2_JDV-2-4-6 | 179,0056558 | 15,22917477 | -3,290003411 | 1E-17 | 1,18E-15 | yes |
| ABCA12 | ENSG00000144452 | ENSG00000144452 | protein_coding | mCh_JDV-1-3-5 | F1F2_JDV-2-4-6 | 4674,31605 | 451,2037999 | -3,301067528 | 2,29E-54 | 2,66E-51 | yes |
| RHOV | ENSG00000104140 | ENSG00000104140 | protein_coding | mCh_JDV-1-3-5 | F1F2_JDV-2-4-6 | 644,4186353 | 51,95239441 | -3,303057986 | 4,57E-15 | 4,01E-13 | yes |
| LMAN1L | ENSG00000140506 | ENSG00000140506 | protein_coding | mCh_JDV-1-3-5 | F1F2_JDV-2-4-6 | 46,83079732 | 1,666833758 | -3,325775933 | 0,000000372 | 0,00000936 | yes |
| CKMT1A | ENSG00000223572 | ENSG00000223572 | protein_coding | mCh_JDV-1-3-5 | F1F2_JDV-2-4-6 | 1246,755003 | 99,98028816 | -3,32786988 | 6,07E-16 | 6,11E-14 | yes |
| CRABP2 | ENSG00000143320 | ENSG00000143320 | protein_coding | mCh_JDV-1-3-5 | F1F2_JDV-2-4-6 | 823,9318433 | 46,05589952 | -3,328986446 | 1,17E-08 | 0,000000401 | yes |
| RDH16 | ENSG00000139547 | ENSG00000139547 | protein_coding | mCh_JDV-1-3-5 | F1F2_JDV-2-4-6 | 98,82159223 | 6,410792003 | -3,330323478 | 2,16E-10 | 1,04E-08 | yes |
| SLC2A12 | ENSG00000146411 | ENSG00000146411 | protein_coding | mCh_JDV-1-3-5 | F1F2_JDV-2-4-6 | 328,5221184 | 29,71638614 | -3,33149705 | 6,37E-32 | 2,51E-29 | yes |
| ALDH1A3 | ENSG00000184254 | ENSG00000184254 | protein_coding | mCh_JDV-1-3-5 | F1F2_JDV-2-4-6 | 1541,3549 | 111,7283887 | -3,332238112 | 2,75E-12 | 1,73E-10 | yes |
| SYT8 | ENSG00000149043 | ENSG00000149043 | protein_coding | mCh_JDV-1-3-5 | F1F2_JDV-2-4-6 | 7799,184458 | 572,0053814 | -3,342864465 | 6,77E-13 | 4,56E-11 | yes |
| IL1A | ENSG00000115008 | ENSG00000115008 | protein_coding | mCh_JDV-1-3-5 | F1F2_JDV-2-4-6 | 1495,115322 | 126,2550026 | -3,345168112 | 3,88E-21 | 6,44E-19 | yes |
| BCAM | ENSG00000187244 | ENSG00000187244 | protein_coding | mCh_JDV-1-3-5 | F1F2_JDV-2-4-6 | 394,3371728 | 30,67814646 | -3,372009053 | 1,34E-16 | 1,43E-14 | yes |
| GRHL1 | ENSG00000134317 | ENSG00000134317 | protein_coding | mCh_JDV-1-3-5 | F1F2_JDV-2-4-6 | 328,8522161 | 27,00884325 | -3,372891951 | 7,05E-21 | 1,12E-18 | yes |
| LEMD1 | ENSG00000186007 | ENSG00000186007 | protein_coding | mCh_JDV-1-3-5 | F1F2_JDV-2-4-6 | 41,45243005 | 2,018307931 | -3,389983403 | 7,62E-09 | 0,000000272 | yes |
| HES2 | ENSG00000069812 | ENSG00000069812 | protein_coding | mCh_JDV-1-3-5 | F1F2_JDV-2-4-6 | 992,3899669 | 71,86323606 | -3,397850417 | 2,5E-14 | 2,02E-12 | yes |
| LRP4 | ENSG00000134569 | ENSG00000134569 | protein_coding | mCh_JDV-1-3-5 | F1F2_JDV-2-4-6 | 615,797214 | 35,71511081 | -3,404622103 | 6,14E-10 | 2,75E-08 | yes |

| | | | | | | | | | | | |
|---|---|---|---|---|---|---|---|---|---|---|---|
| ENTPD2 | ENSG00000054179 | ENSG00000054179 | protein_coding | mCh_JDV-1-3-5 | F1F2_JDV-2-4-6 | 3432,461106 | 198,103537 | -3,408358376 | 6,57E-10 | 2,92E-08 | yes |
| GYLTL1B | ENSG00000165905 | ENSG00000165905 | protein_coding | mCh_JDV-1-3-5 | F1F2_JDV-2-4-6 | 87,43218555 | 5,721025768 | -3,448656818 | 2,98E-13 | 2,1E-11 | yes |
| MDFI | ENSG00000112559 | ENSG00000112559 | protein_coding | mCh_JDV-1-3-5 | F1F2_JDV-2-4-6 | 508,178848 | 40,63415075 | -3,459224811 | 8,81E-27 | 2,2E-24 | yes |
| ATP8B1 | ENSG00000081923 | ENSG00000081923 | protein_coding | mCh_JDV-1-3-5 | F1F2_JDV-2-4-6 | 418,9362749 | 30,80142307 | -3,477116792 | 4,73E-19 | 6,48E-17 | yes |
| CBLC | ENSG00000142273 | ENSG00000142273 | protein_coding | mCh_JDV-1-3-5 | F1F2_JDV-2-4-6 | 240,4276405 | 17,94763083 | -3,48660937 | 4,45E-21 | 7,32E-19 | yes |
| ST6GALNAC2 | ENSG00000070731 | ENSG00000070731 | protein_coding | mCh_JDV-1-3-5 | F1F2_JDV-2-4-6 | 562,526976 | 40,4923793 | -3,495066709 | 1,12E-18 | 1,47E-16 | yes |
| SLC7A5 | ENSG00000103257 | ENSG00000103257 | protein_coding | mCh_JDV-1-3-5 | F1F2_JDV-2-4-6 | 66819,64573 | 4509,567079 | -3,499981281 | 2,13E-15 | 1,95E-13 | yes |
| SAA4 | ENSG00000148965 | ENSG00000148965 | protein_coding | mCh_JDV-1-3-5 | F1F2_JDV-2-4-6 | 235,9510451 | 9,752185047 | -3,562128364 | 3,68E-09 | 0,000000139 | yes |
| LEPREL1 | ENSG00000090530 | ENSG00000090530 | protein_coding | mCh_JDV-1-3-5 | F1F2_JDV-2-4-6 | 2404,889561 | 187,3891299 | -3,564021194 | 5,24E-43 | 3,57E-40 | yes |
| GPR143 | ENSG00000101850 | ENSG00000101850 | protein_coding | mCh_JDV-1-3-5 | F1F2_JDV-2-4-6 | 332,5446639 | 22,28579264 | -3,575174212 | 8,2E-19 | 1,1E-16 | yes |
| SLC1A3 | ENSG00000079215 | ENSG00000079215 | protein_coding | mCh_JDV-1-3-5 | F1F2_JDV-2-4-6 | 4581,367957 | 341,4112555 | -3,607249868 | 1,92E-38 | 1,05E-35 | yes |
| ADAP1 | ENSG00000105963 | ENSG00000105963 | protein_coding | mCh_JDV-1-3-5 | F1F2_JDV-2-4-6 | 166,9974948 | 9,793856473 | -3,624682673 | 3,9E-15 | 3,44E-13 | yes |
| FUT1 | ENSG00000174951 | ENSG00000174951 | protein_coding | mCh_JDV-1-3-5 | F1F2_JDV-2-4-6 | 157,4912128 | 9,500423478 | -3,635462741 | 1,65E-16 | 1,74E-14 | yes |
| PTPRF | ENSG00000142949 | ENSG00000142949 | protein_coding | mCh_JDV-1-3-5 | F1F2_JDV-2-4-6 | 9517,827261 | 638,0957077 | -3,663515438 | 1,43E-25 | 3,44E-23 | yes |
| CGN | ENSG00000143375 | ENSG00000143375 | protein_coding | mCh_JDV-1-3-5 | F1F2_JDV-2-4-6 | 395,8391352 | 16,22488099 | -3,673757323 | 3,08E-10 | 1,47E-08 | yes |
| MPZL2 | ENSG00000149573 | ENSG00000149573 | protein_coding | mCh_JDV-1-3-5 | F1F2_JDV-2-4-6 | 7562,442829 | 555,9218798 | -3,683314692 | 2,28E-65 | 4,99E-62 | yes |
| CKMT1B | ENSG00000237289 | ENSG00000237289 | protein_coding | mCh_JDV-1-3-5 | F1F2_JDV-2-4-6 | 655,6707017 | 39,12723533 | -3,68978026 | 3,71E-18 | 4,55E-16 | yes |
| CDH3 | ENSG00000062038 | ENSG00000062038 | protein_coding | mCh_JDV-1-3-5 | F1F2_JDV-2-4-6 | 20539,85567 | 1479,750345 | -3,694168083 | 8,15E-55 | 1,07E-51 | yes |
| SEMA3A | ENSG00000075213 | ENSG00000075213 | protein_coding | mCh_JDV-1-3-5 | F1F2_JDV-2-4-6 | 346,1356627 | 17,95468124 | -3,699874635 | 8,2E-14 | 6,2E-12 | yes |
| ARHGAP25 | ENSG00000163219 | ENSG00000163219 | protein_coding | mCh_JDV-1-3-5 | F1F2_JDV-2-4-6 | 166,2217126 | 7,130110101 | -3,72044782 | 2,22E-11 | 1,24E-09 | yes |
| IRF6 | ENSG00000117595 | ENSG00000117595 | protein_coding | mCh_JDV-1-3-5 | F1F2_JDV-2-4-6 | 9947,308814 | 637,2367432 | -3,737713006 | 7,59E-28 | 2,14E-25 | yes |
| C1orf106 | ENSG00000163362 | ENSG00000163362 | protein_coding | mCh_JDV-1-3-5 | F1F2_JDV-2-4-6 | 1588,634012 | 95,14735648 | -3,738072882 | 5,74E-21 | 9,28E-19 | yes |
| FGF1 | ENSG00000113578 | ENSG00000113578 | protein_coding | mCh_JDV-1-3-5 | F1F2_JDV-2-4-6 | 66,51441747 | 2,036946626 | -3,743626594 | 1,19E-09 | 4,94E-08 | yes |
| TNNI1 | ENSG00000159173 | ENSG00000159173 | protein_coding | mCh_JDV-1-3-5 | F1F2_JDV-2-4-6 | 73,72078603 | 3,708174421 | -3,757476111 | 2,9E-15 | 2,6E-13 | yes |
| FGFR2 | ENSG00000066468 | ENSG00000066468 | protein_coding | mCh_JDV-1-3-5 | F1F2_JDV-2-4-6 | 335,8670497 | 20,93262439 | -3,802537131 | 8,95E-33 | 3,76E-30 | yes |
| CD82 | ENSG00000085117 | ENSG00000085117 | protein_coding | mCh_JDV-1-3-5 | F1F2_JDV-2-4-6 | 3205,295437 | 199,0472548 | -3,809457248 | 1,16E-32 | 4,77E-30 | yes |
| TMCC3 | ENSG00000057704 | ENSG00000057704 | protein_coding | mCh_JDV-1-3-5 | F1F2_JDV-2-4-6 | 267,8289719 | 12,54678971 | -3,819476847 | 1,48E-14 | 1,23E-12 | yes |
| TMEM125 | ENSG00000179178 | ENSG00000179178 | protein_coding | mCh_JDV-1-3-5 | F1F2_JDV-2-4-6 | 127,7764623 | 6,41026073 | -3,826117396 | 5,32E-17 | 5,9E-15 | yes |
| BTBD11 | ENSG00000151136 | ENSG00000151136 | protein_coding | mCh_JDV-1-3-5 | F1F2_JDV-2-4-6 | 482,2389972 | 30,12430767 | -3,828998131 | 7,05E-38 | 3,76E-35 | yes |
| RNF152 | ENSG00000176641 | ENSG00000176641 | protein_coding | mCh_JDV-1-3-5 | F1F2_JDV-2-4-6 | 540,3179526 | 24,91173035 | -3,834654015 | 1,76E-14 | 1,45E-12 | yes |
| ARHGEF16 | ENSG00000130762 | ENSG00000130762 | protein_coding | mCh_JDV-1-3-5 | F1F2_JDV-2-4-6 | 883,6496011 | 52,38508634 | -3,84303153 | 1,79E-29 | 5,8E-27 | yes |
| TNNT2 | ENSG00000118194 | ENSG00000118194 | protein_coding | mCh_JDV-1-3-5 | F1F2_JDV-2-4-6 | 97,34112115 | 3,392383817 | -3,858294675 | 1,66E-11 | 9,52E-10 | yes |
| LAMA3 | ENSG00000053747 | ENSG00000053747 | protein_coding | mCh_JDV-1-3-5 | F1F2_JDV-2-4-6 | 41260,89034 | 2570,198362 | -3,869786916 | 5,5E-48 | 4,52E-45 | yes |
| NTF4 | ENSG00000167744 | ENSG00000167744 | protein_coding | mCh_JDV-1-3-5 | F1F2_JDV-2-4-6 | 220,0729687 | 9,397379384 | -3,873086775 | 5,04E-14 | 3,9E-12 | yes |
| EPN3 | ENSG00000049283 | ENSG00000049283 | protein_coding | mCh_JDV-1-3-5 | F1F2_JDV-2-4-6 | 180,1575327 | 8,758603923 | -3,882089087 | 9,19E-18 | 1,1E-15 | yes |
| KRT17 | ENSG00000128422 | ENSG00000128422 | protein_coding | mCh_JDV-1-3-5 | F1F2_JDV-2-4-6 | 13306,24335 | 692,7266222 | -3,904798748 | 1,1E-21 | 1,96E-19 | yes |
| AZGP1 | ENSG00000160862 | ENSG00000160862 | protein_coding | mCh_JDV-1-3-5 | F1F2_JDV-2-4-6 | 326,5949214 | 14,84322343 | -3,941083701 | 2,77E-17 | 3,2E-15 | yes |
| AIM1L | ENSG00000176092 | ENSG00000176092 | protein_coding | mCh_JDV-1-3-5 | F1F2_JDV-2-4-6 | 162,2820902 | 4,731307408 | -3,97950359 | 1,82E-11 | 1,04E-09 | yes |
| PVRL1 | ENSG00000110400 | ENSG00000110400 | protein_coding | mCh_JDV-1-3-5 | F1F2_JDV-2-4-6 | 9209,159367 | 457,8702556 | -4,059963107 | 3,75E-30 | 1,27E-27 | yes |
| IGSF9 | ENSG00000085552 | ENSG00000085552 | protein_coding | mCh_JDV-1-3-5 | F1F2_JDV-2-4-6 | 316,3589405 | 15,59262466 | -4,097063485 | 6,74E-34 | 2,96E-31 | yes |
| KRT15 | ENSG00000171346 | ENSG00000171346 | protein_coding | mCh_JDV-1-3-5 | F1F2_JDV-2-4-6 | 26874,61981 | 734,9729437 | -4,107300261 | 3,68E-12 | 2,27E-10 | yes |
| HSH2D | ENSG00000196684 | ENSG00000196684 | protein_coding | mCh_JDV-1-3-5 | F1F2_JDV-2-4-6 | 220,4529001 | 8,785499418 | -4,1101114 | 6,22E-19 | 8,4E-17 | yes |
| KLK7 | ENSG00000169035 | ENSG00000169035 | protein_coding | mCh_JDV-1-3-5 | F1F2_JDV-2-4-6 | 393,9480211 | 1,351574428 | -4,112193387 | 8,28E-09 | 0,000000293 | yes |
| ST14 | ENSG00000149418 | ENSG00000149418 | protein_coding | mCh_JDV-1-3-5 | F1F2_JDV-2-4-6 | 7359,463444 | 372,272933 | -4,113264822 | 8,79E-42 | 5,25E-39 | yes |
| CXADR | ENSG00000154639 | ENSG00000154639 | protein_coding | mCh_JDV-1-3-5 | F1F2_JDV-2-4-6 | 1495,278628 | 70,80215056 | -4,11993465 | 1,43E-30 | 5,02E-28 | yes |
| CRB3 | ENSG00000130545 | ENSG00000130545 | protein_coding | mCh_JDV-1-3-5 | F1F2_JDV-2-4-6 | 178,6828182 | 6,419580077 | -4,127696961 | 1,2E-16 | 1,29E-14 | yes |
| SERPINF1 | ENSG00000132386 | ENSG00000132386 | protein_coding | mCh_JDV-1-3-5 | F1F2_JDV-2-4-6 | 359,3362762 | 12,1383369 | -4,140697811 | 1,94E-15 | 1,81E-13 | yes |

| | | | | | | | | | | | |
|---|---|---|---|---|---|---|---|---|---|---|---|
| GPM6A | ENSG00000150625 | ENSG00000150625 | protein_coding | mCh_JDV-1-3-5 | F1F2_JDV-2-4-6 | 128,974006 | 4,027827788 | -4,156972729 | 5,62E-15 | 4,86E-13 | yes |
| TNNI2 | ENSG00000130598 | ENSG00000130598 | protein_coding | mCh_JDV-1-3-5 | F1F2_JDV-2-4-6 | 3981,459657 | 157,3150288 | -4,162584968 | 4,24E-20 | 6,2E-18 | yes |
| DSC2 | ENSG00000134755 | ENSG00000134755 | protein_coding | mCh_JDV-1-3-5 | F1F2_JDV-2-4-6 | 3762,927684 | 178,2088423 | -4,217092515 | 9,36E-47 | 7,1E-44 | yes |
| EPS8L1 | ENSG00000131037 | ENSG00000131037 | protein_coding | mCh_JDV-1-3-5 | F1F2_JDV-2-4-6 | 265,5490451 | 10,13760512 | -4,255993557 | 2,32E-23 | 5,09E-21 | yes |
| PADI2 | ENSG00000117115 | ENSG00000117115 | protein_coding | mCh_JDV-1-3-5 | F1F2_JDV-2-4-6 | 243,9704931 | 5,773610362 | -4,266302414 | 2,58E-13 | 1,82E-11 | yes |
| CLDN4 | ENSG00000189143 | ENSG00000189143 | protein_coding | mCh_JDV-1-3-5 | F1F2_JDV-2-4-6 | 565,7391245 | 14,82617855 | -4,275367057 | 3,14E-14 | 2,5E-12 | yes |
| ANKRD22 | ENSG00000152766 | ENSG00000152766 | protein_coding | mCh_JDV-1-3-5 | F1F2_JDV-2-4-6 | 181,5225487 | 4,374907924 | -4,323957061 | 1,83E-14 | 1,5E-12 | yes |
| GRAMD2 | ENSG00000175318 | ENSG00000175318 | protein_coding | mCh_JDV-1-3-5 | F1F2_JDV-2-4-6 | 483,3841396 | 18,86226314 | -4,371399841 | 8,12E-34 | 3,48E-31 | yes |
| HBEGF | ENSG00000113070 | ENSG00000113070 | protein_coding | mCh_JDV-1-3-5 | F1F2_JDV-2-4-6 | 847,6160599 | 30,89567909 | -4,376710882 | 4,2E-27 | 1,12E-24 | yes |
| CAPN8 | ENSG00000203697 | ENSG00000203697 | protein_coding | mCh_JDV-1-3-5 | F1F2_JDV-2-4-6 | 146,9424083 | 4,707212129 | -4,392897281 | 2,88E-22 | 5,46E-20 | yes |
| TRPV4 | ENSG00000111199 | ENSG00000111199 | protein_coding | mCh_JDV-1-3-5 | F1F2_JDV-2-4-6 | 166,8479988 | 5,407360258 | -4,429533364 | 3,74E-24 | 8,79E-22 | yes |
| C1orf116 | ENSG00000182795 | ENSG00000182795 | protein_coding | mCh_JDV-1-3-5 | F1F2_JDV-2-4-6 | 2224,840299 | 57,49674466 | -4,455634214 | 3,04E-17 | 3,49E-15 | yes |
| SEMA6A | ENSG00000092421 | ENSG00000092421 | protein_coding | mCh_JDV-1-3-5 | F1F2_JDV-2-4-6 | 925,7168536 | 26,67494522 | -4,466564773 | 3,42E-20 | 5,07E-18 | yes |
| XDH | ENSG00000158125 | ENSG00000158125 | protein_coding | mCh_JDV-1-3-5 | F1F2_JDV-2-4-6 | 849,4287849 | 33,45744392 | -4,48696208 | 2,65E-57 | 3,73E-54 | yes |
| IL1RN | ENSG00000136689 | ENSG00000136689 | protein_coding | mCh_JDV-1-3-5 | F1F2_JDV-2-4-6 | 169,7736189 | 1,684941179 | -4,510235334 | 4,64E-12 | 2,81E-10 | yes |
| PRSS8 | ENSG00000052344 | ENSG00000052344 | protein_coding | mCh_JDV-1-3-5 | F1F2_JDV-2-4-6 | 1618,530288 | 51,98807798 | -4,517700704 | 2,62E-27 | 7,19E-25 | yes |
| LSR | ENSG00000105699 | ENSG00000105699 | protein_coding | mCh_JDV-1-3-5 | F1F2_JDV-2-4-6 | 2457,721838 | 81,68225036 | -4,529172062 | 9,17E-31 | 3,29E-28 | yes |
| CLDN1 | ENSG00000163347 | ENSG00000163347 | protein_coding | mCh_JDV-1-3-5 | F1F2_JDV-2-4-6 | 4157,190623 | 117,4261776 | -4,577907559 | 1,9E-23 | 4,22E-21 | yes |
| PLA2G4E | ENSG00000188089 | ENSG00000188089 | protein_coding | mCh_JDV-1-3-5 | F1F2_JDV-2-4-6 | 131,6123236 | 3,366019596 | -4,581466142 | 8,99E-22 | 1,63E-19 | yes |
| CLCA2 | ENSG00000137975 | ENSG00000137975 | protein_coding | mCh_JDV-1-3-5 | F1F2_JDV-2-4-6 | 20930,82336 | 580,4422047 | -4,603243905 | 1,06E-23 | 2,46E-21 | yes |
| NFASC | ENSG00000163531 | ENSG00000163531 | protein_coding | mCh_JDV-1-3-5 | F1F2_JDV-2-4-6 | 126,8841606 | 2,689435471 | -4,638460864 | 3,8E-19 | 5,25E-17 | yes |
| SLC6A14 | ENSG00000087916 | ENSG00000087916 | protein_coding | mCh_JDV-1-3-5 | F1F2_JDV-2-4-6 | 119,3026662 | 2,369782104 | -4,655580306 | 1,77E-18 | 2,27E-16 | yes |
| S100A14 | ENSG00000189334 | ENSG00000189334 | protein_coding | mCh_JDV-1-3-5 | F1F2_JDV-2-4-6 | 15604,86762 | 488,1683784 | -4,66435841 | 5,59E-37 | 2,69E-34 | yes |
| SLC27A2 | ENSG00000140284 | ENSG00000140284 | protein_coding | mCh_JDV-1-3-5 | F1F2_JDV-2-4-6 | 1529,638387 | 48,59076885 | -4,750034081 | 1,23E-54 | 1,52E-51 | yes |
| TINAGL1 | ENSG00000142910 | ENSG00000142910 | protein_coding | mCh_JDV-1-3-5 | F1F2_JDV-2-4-6 | 1641,546393 | 47,87942451 | -4,750556494 | 1,1E-37 | 5,58E-35 | yes |
| MMP28 | ENSG00000129270 | ENSG00000129270 | protein_coding | mCh_JDV-1-3-5 | F1F2_JDV-2-4-6 | 1718,945849 | 46,20688594 | -4,767020448 | 3,79E-31 | 1,41E-28 | yes |
| GLDC | ENSG00000178445 | ENSG00000178445 | protein_coding | mCh_JDV-1-3-5 | F1F2_JDV-2-4-6 | 758,7324121 | 21,02847423 | -4,790289453 | 4,58E-36 | 2,15E-33 | yes |
| COL17A1 | ENSG00000065618 | ENSG00000065618 | protein_coding | mCh_JDV-1-3-5 | F1F2_JDV-2-4-6 | 31039,39645 | 933,9905024 | -4,805614123 | 1,83E-51 | 1,8E-48 | yes |
| OCLN | ENSG00000197822 | ENSG00000197822 | protein_coding | mCh_JDV-1-3-5 | F1F2_JDV-2-4-6 | 865,5127222 | 20,26815983 | -4,848062829 | 1,42E-27 | 3,93E-25 | yes |
| DENND1C | ENSG00000205744 | ENSG00000205744 | protein_coding | mCh_JDV-1-3-5 | F1F2_JDV-2-4-6 | 300,560825 | 7,054492775 | -4,8794712 | 1,88E-30 | 6,52E-28 | yes |
| FGFBP1 | ENSG00000137440 | ENSG00000137440 | protein_coding | mCh_JDV-1-3-5 | F1F2_JDV-2-4-6 | 7911,462018 | 171,5416999 | -4,924745494 | 3,67E-27 | 9,91E-25 | yes |
| GRIP1 | ENSG00000155974 | ENSG00000155974 | protein_coding | mCh_JDV-1-3-5 | F1F2_JDV-2-4-6 | 489,8689888 | 10,8048699 | -4,962860919 | 6,34E-31 | 2,32E-28 | yes |
| FAM83B | ENSG00000168143 | ENSG00000168143 | protein_coding | mCh_JDV-1-3-5 | F1F2_JDV-2-4-6 | 619,9729199 | 15,5630728 | -5,015255109 | 1,72E-50 | 1,62E-47 | yes |
| DSC3 | ENSG00000134762 | ENSG00000134762 | protein_coding | mCh_JDV-1-3-5 | F1F2_JDV-2-4-6 | 14072,0409 | 375,980249 | -5,049670168 | 1,78E-80 | 5,01E-77 | yes |
| NGFR | ENSG00000064300 | ENSG00000064300 | protein_coding | mCh_JDV-1-3-5 | F1F2_JDV-2-4-6 | 318,7600199 | 3,051291539 | -5,113355956 | 9,92E-18 | 1,17E-15 | yes |
| PI3 | ENSG00000124102 | ENSG00000124102 | protein_coding | mCh_JDV-1-3-5 | F1F2_JDV-2-4-6 | 1341,850833 | 16,58128047 | -5,152375041 | 4,29E-20 | 6,23E-18 | yes |
| PCDH1 | ENSG00000156453 | ENSG00000156453 | protein_coding | mCh_JDV-1-3-5 | F1F2_JDV-2-4-6 | 1942,999783 | 42,3229547 | -5,156016684 | 1,18E-45 | 8,6E-43 | yes |
| TSPAN1 | ENSG00000117472 | ENSG00000117472 | protein_coding | mCh_JDV-1-3-5 | F1F2_JDV-2-4-6 | 6396,687679 | 141,1588145 | -5,215269937 | 1,69E-57 | 2,57E-54 | yes |
| NIPAL4 | ENSG00000172548 | ENSG00000172548 | protein_coding | mCh_JDV-1-3-5 | F1F2_JDV-2-4-6 | 660,9341162 | 8,423112077 | -5,391864371 | 5,82E-27 | 1,48E-24 | yes |
| GPR56 | ENSG00000205336 | ENSG00000205336 | protein_coding | mCh_JDV-1-3-5 | F1F2_JDV-2-4-6 | 1611,681155 | 29,01730056 | -5,411137625 | 2,61E-50 | 2,34E-47 | yes |
| TMC4 | ENSG00000167608 | ENSG00000167608 | protein_coding | mCh_JDV-1-3-5 | F1F2_JDV-2-4-6 | 1492,835404 | 14,58267379 | -5,603489156 | 2,8E-26 | 6,92E-24 | yes |
| EPCAM | ENSG00000119888 | ENSG00000119888 | protein_coding | mCh_JDV-1-3-5 | F1F2_JDV-2-4-6 | 4607,461169 | 80,97848304 | -5,62835111 | 7,64E-98 | 3,77E-94 | yes |
| TP63 | ENSG00000073282 | ENSG00000073282 | protein_coding | mCh_JDV-1-3-5 | F1F2_JDV-2-4-6 | 1764,990239 | 25,58725193 | -5,62953791 | 4,64E-47 | 3,66E-44 | yes |
| DSG3 | ENSG00000134757 | ENSG00000134757 | protein_coding | mCh_JDV-1-3-5 | F1F2_JDV-2-4-6 | 26114,95687 | 398,8502861 | -5,837983358 | 1,5E-111 | 1,48E-107 | yes |
| F5 | ENSG00000198734 | ENSG00000198734 | protein_coding | mCh_JDV-1-3-5 | F1F2_JDV-2-4-6 | 237,8190744 | 1,347180391 | -5,987773464 | 1,26E-29 | 4,14E-27 | yes |
| PTAFR | ENSG00000169403 | ENSG00000169403 | protein_coding | mCh_JDV-1-3-5 | F1F2_JDV-2-4-6 | 1675,680274 | 21,62398445 | -6,02699928 | 3,3E-100 | 2,17E-96 | yes |
| TSPAN18 | ENSG00000157570 | ENSG00000157570 | protein_coding | mCh_JDV-1-3-5 | F1F2_JDV-2-4-6 | 465,9389689 | 4,384758545 | -6,032376684 | 1,94E-45 | 1,37E-42 | yes |

| | | | | | | | | | | | |
|---|---|---|---|---|---|---|---|---|---|---|---|
| AP1M2 | ENSG00000129354 | ENSG00000129354 | protein_coding | mCh_JDV-1-3-5 | F1F2_JDV-2-4-6 | 638,0352508 | 5,734739153 | -6,140217273 | 4,3E-49 | 3,69E-46 | yes |
| CAMK2B | ENSG00000058404 | ENSG00000058404 | protein_coding | mCh_JDV-1-3-5 | F1F2_JDV-2-4-6 | 1156,223461 | 9,124322753 | -6,200392969 | 6,3E-43 | 4,01E-40 | yes |
| FGFR3 | ENSG00000068078 | ENSG00000068078 | protein_coding | mCh_JDV-1-3-5 | F1F2_JDV-2-4-6 | 2476,526583 | 14,26408296 | -6,218393944 | 9,13E-32 | 3,53E-29 | yes |
| B3GNT3 | ENSG00000179913 | ENSG00000179913 | protein_coding | mCh_JDV-1-3-5 | F1F2_JDV-2-4-6 | 860,0114355 | 3,721887805 | -6,910993566 | 1,6E-53 | 1,75E-50 | yes |
| BDKRB2 | ENSG00000168398 | ENSG00000168398 | protein_coding | mCh_JDV-1-3-5 | F1F2_JDV-2-4-6 | 1871,677943 | 11,18483338 | -6,966142498 | 2,22E-97 | 8,75E-94 | yes |
| LAD1 | ENSG00000159166 | ENSG00000159166 | protein_coding | mCh_JDV-1-3-5 | F1F2_JDV-2-4-6 | 1991,430933 | 6,754009376 | -7,268695098 | 3,16E-59 | 5,19E-56 | yes |
| CDH1 | ENSG00000039068 | ENSG00000039068 | protein_coding | mCh_JDV-1-3-5 | F1F2_JDV-2-4-6 | 17574,64659 | 78,84529915 | -7,30681257 | 9,5E-94 | 3,13E-90 | yes |
| FUT3 | ENSG00000171124 | ENSG00000171124 | protein_coding | mCh_JDV-1-3-5 | F1F2_JDV-2-4-6 | 616,1862306 | 1,008888329 | -7,526540292 | 1,8E-52 | 1,87E-49 | yes |
| ESRP1 | ENSG00000104413 | ENSG00000104413 | protein_coding | mCh_JDV-1-3-5 | F1F2_JDV-2-4-6 | 3026,065506 | 12,15151901 | -7,52832695 | 1,89E-119 | 3,73E-115 | yes |
| FXYD3 | ENSG00000089356 | ENSG00000089356 | protein_coding | mCh_JDV-1-3-5 | F1F2_JDV-2-4-6 | 4296,903118 | 12,82264655 | -7,636068291 | 7,36E-78 | 1,82E-74 | yes |

### Supplementary Table S2

**List of the enriched GO terms and the enriched pathways obtained from the analysis of the 802 differentially expressed genes between MCF10mCh and MCF10AF1F2 using three distinct bioinfomatic tools (KEGG, DAVID, and Genomatix)**

| KEGG Enrichr pathways | KEGG David | Genomatix |
|---|---|---|
| Cell adhesion molecules (CAMs)_Homo sapiens_hsa04514 | hsa04151:PI3K-Akt signaling pathway | Extracellular matrix organization |
| PI3K-Akt signaling pathway_Homo sapiens_hsa04151 | hsa04514:Cell adhesion molecules (CAMs) | Cell-Cell communication |
| Proteoglycans in cancer_Homo sapiens_hsa05205 | hsa05205:Proteoglycans in cancer | Cell junction organization |
| Hematopoietic cell lineage_Homo sapiens_hsa04640 | hsa04640:Hematopoietic cell lineage | a6b1 and a6b4 Integrin signaling |
| Pathways in cancer_Homo sapiens_hsa05200 | hsa05200:Pathways in cancer | Beta5 beta6 beta7 and beta8 integrin cell surface interactions |
| Rap1 signaling pathway_Homo sapiens_hsa04015 | hsa04015:Rap1 signaling pathway | amb2 Integrin signaling |
| Malaria_Homo sapiens_hsa05144 | hsa05144:Malaria | Alpha6Beta4Integrin |
| Inflammatory bowel disease (IBD)_Homo sapiens_hsa05321 | hsa04060:Cytokine-cytokine receptor interaction | EGFR1 |
| Insulin resistance_Homo sapiens_hsa04931 | hsa05321:Inflammatory bowel disease (IBD) | Syndecan-2-mediated signaling events |
| AGE-RAGE signaling pathway in diabetic complications_Homo sapiens_hsa04933 | hsa04931:Insulin resistance | Direct p53 effectors |
| Ras signaling pathway_Homo sapiens_hsa04014 | hsa04014:Ras signaling pathway | FGF signaling pathway |
| Cytokine-cytokine receptor interaction_Homo sapiens_hsa04060 | hsa04512:ECM-receptor interaction | Signaling by MST1 |
| Prion diseases_Homo sapiens_hsa05020 | hsa05323:Rheumatoid arthritis | Transport of small molecules |
| | hsa05230:Central carbon metabolism in cancer | Visual phototransduction |
| | hsa00601:Glycosphingolipid biosynthesis - lacto and neolacto series | Cytokine Signaling in Immune system |
| | hsa04750:Inflammatory mediator regulation of TRP channels | SHP2 signaling |
| | hsa04610:Complement and coagulation cascades | Beta2 integrin cell surface interactions |
| | hsa05412:Arrhythmogenic right ventricular cardiomyopathy (ARVC) | Angiopoietin receptor Tie2-mediated signaling |
| | hsa05218:Melanoma | Hemostasis |
| | hsa05202:Transcriptional misregulation in cancer | Validated transcriptional targets of TAp63 isoforms |
| | hsa04064:NF-kappa B signaling pathway | ion channels and their functional role in vascular endothelium |
| | hsa04144:Endocytosis | Signaling by Retinoic Acid |
| | | Beta3 integrin cell surface interactions |
| | | Blood coagulation |
| | | Beta1 integrin cell surface interactions |
| | | Plasminogen activating cascade |
| | | Validated transcriptional targets of deltaNp63 isoforms |
| | | ATF-2 transcription factor network |
| | | Endogenous TLR signaling |
| | | IL27-mediated signaling events |
| | | Validated transcriptional targets of AP1 family members Fra1 and Fra2 |
| | | signal transduction through il1r |
| | | inhibition of matrix metalloproteinases |
| | | IL-7 |
| | | PI3K/AKT Signaling in Cancer |

**David version 6.8**

https://david.ncifcrf.gov/home.jsp

DAVID users may publish or otherwise publicly disclose the results of using DAVID. Please acknowledge DAVID in your publications by citing the following two references:

Huang DW, Sherman BT, Lempicki RA. Systematic and integrative analysis of large gene lists using DAVID Bioinformatics Resources. Nature Protoc. 2009;4(1):44-57.

Huang DW, Sherman BT, Lempicki RA. Bioinformatics enrichment tools: paths toward the comprehensive functional analysis of large gene lists. Nucleic Acids Res. 2009;37(1):1-13.

**KEGG Pathway DataBase**

https://www.kegg.jp/kegg/pathway.html

Please cite the following article(s) when using KEGG.

Kanehisa, Furumichi, M., Tanabe, M., Sato, Y., and Morishima, K.; KEGG: new perspectives on genomes, pathways, diseases and drugs. Nucleic Acids Res. 45, D353-D361 (2017).

Kanehisa, M., Sato, Y., Kawashima, M., Furumichi, M., and Tanabe, M.; KEGG as a reference resource for gene and protein annotation. Nucleic Acids Res. 44, D457-D462 (2016).

Kanehisa, M. and Goto, S.; KEGG: Kyoto Encyclopedia of Genes and Genomes. Nucleic Acids Res. 28, 27-30 (2000).

**Genomatix**

http://www.genomatix.de/

Genomatix Software suite version 3.10
