## Supplementary figures and images for "Upregulated-flotillins and sphingosine kinase 2 derail vesicular trafic to stabilize AXL and promote epithelial-mesenchymal transition"

### Genest Figure S1

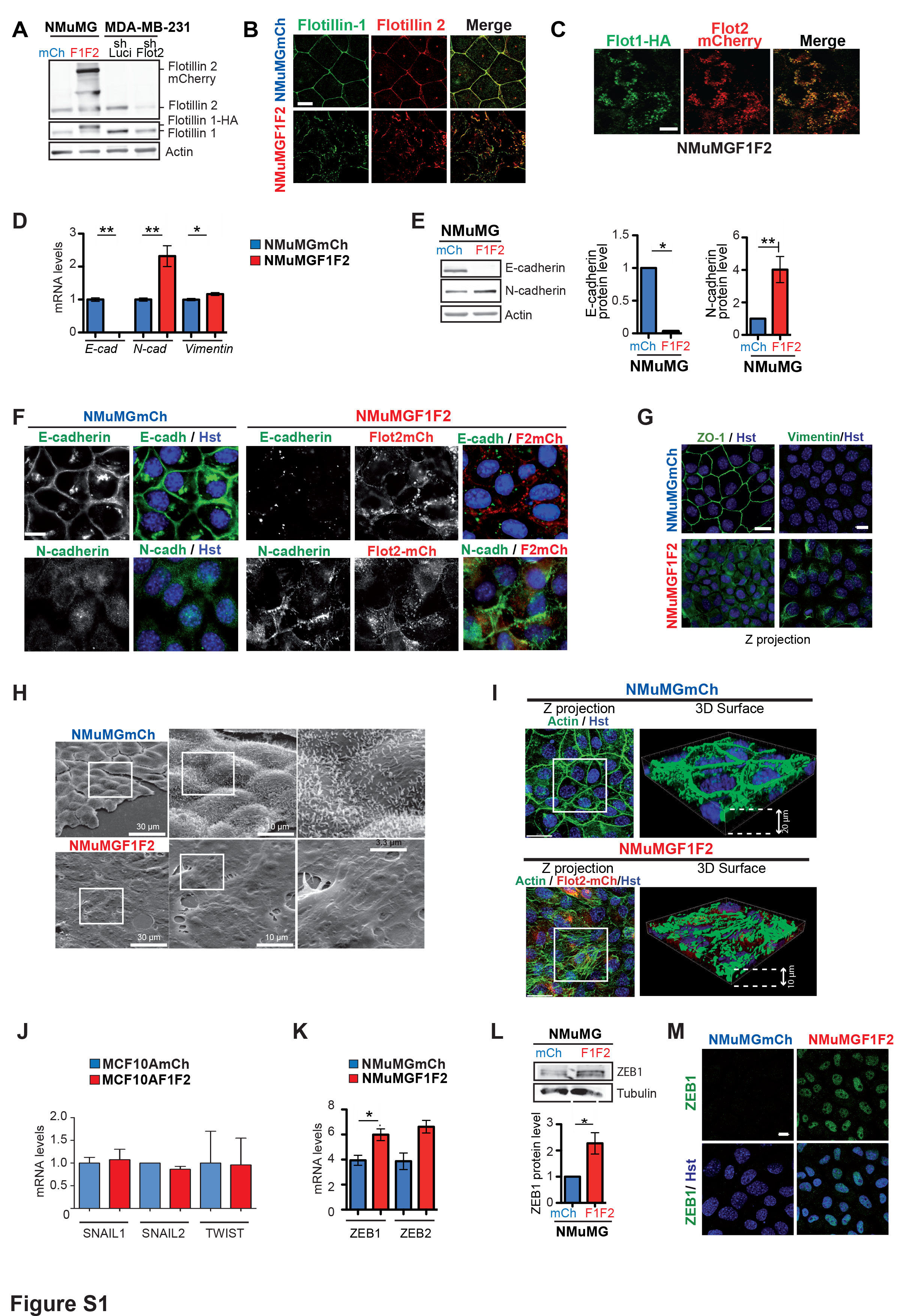

### Genest Figure S2

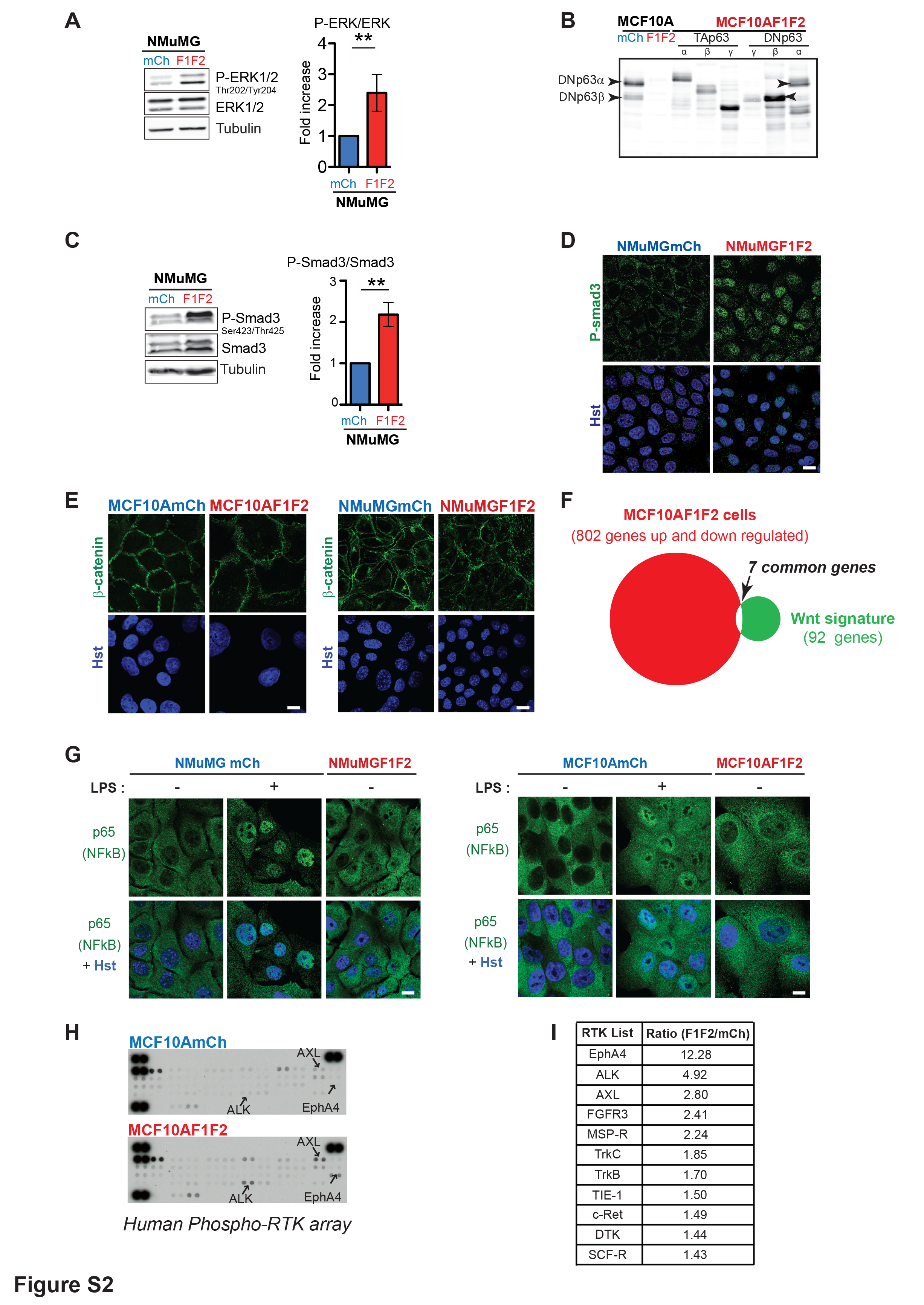

### Genest Figure S3

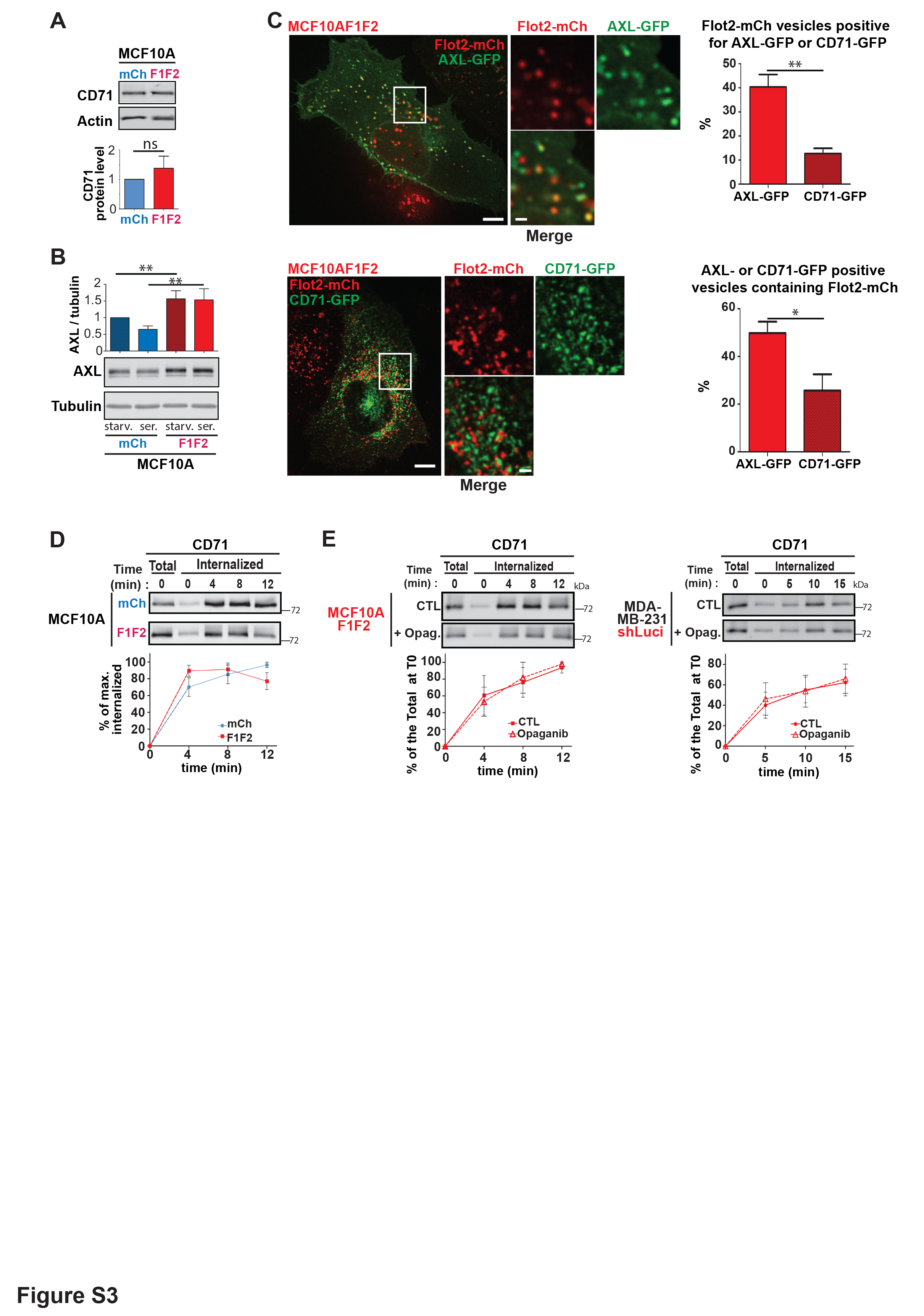

### Genest Figure S4

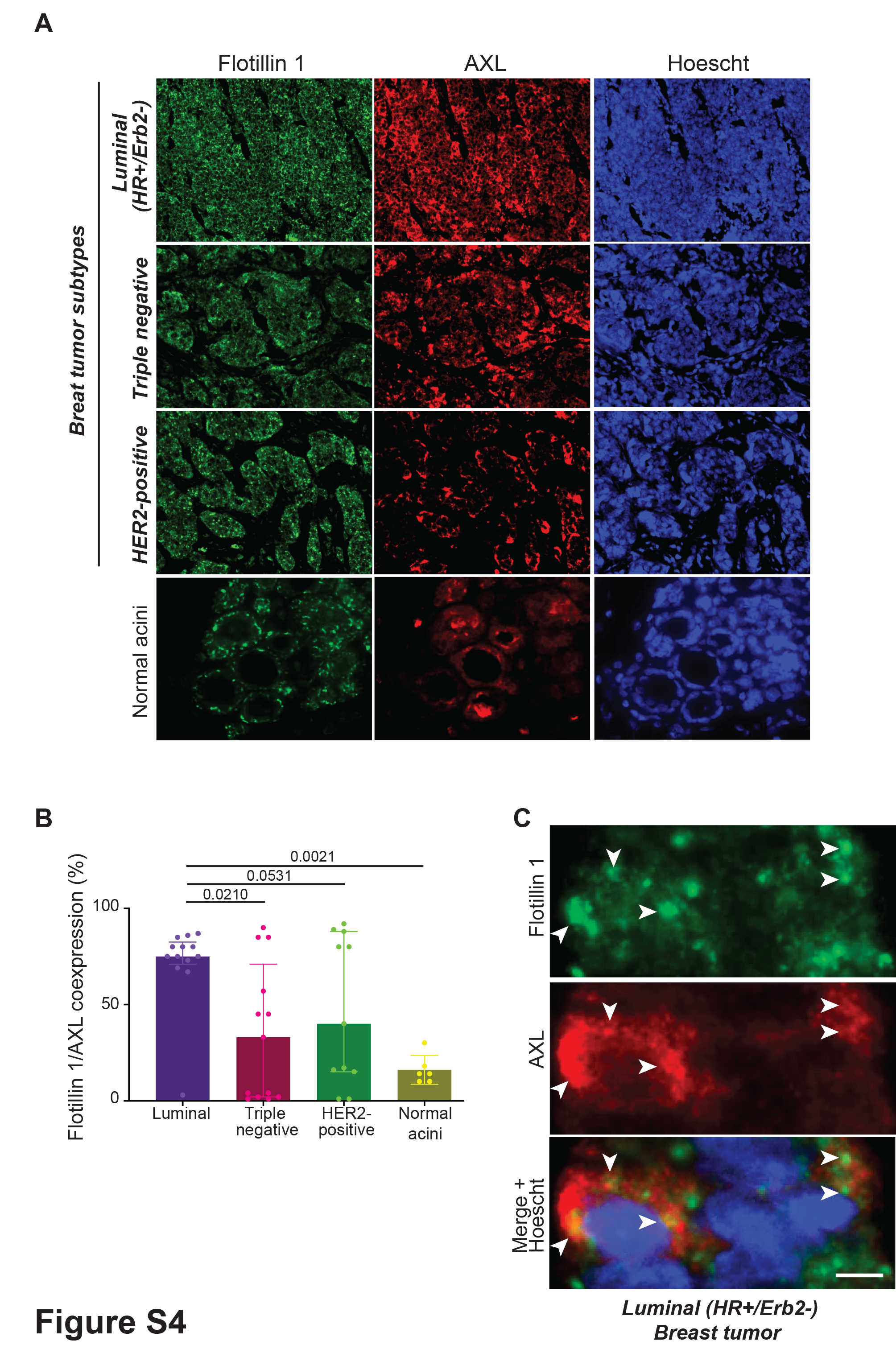

### Genest Figure S5

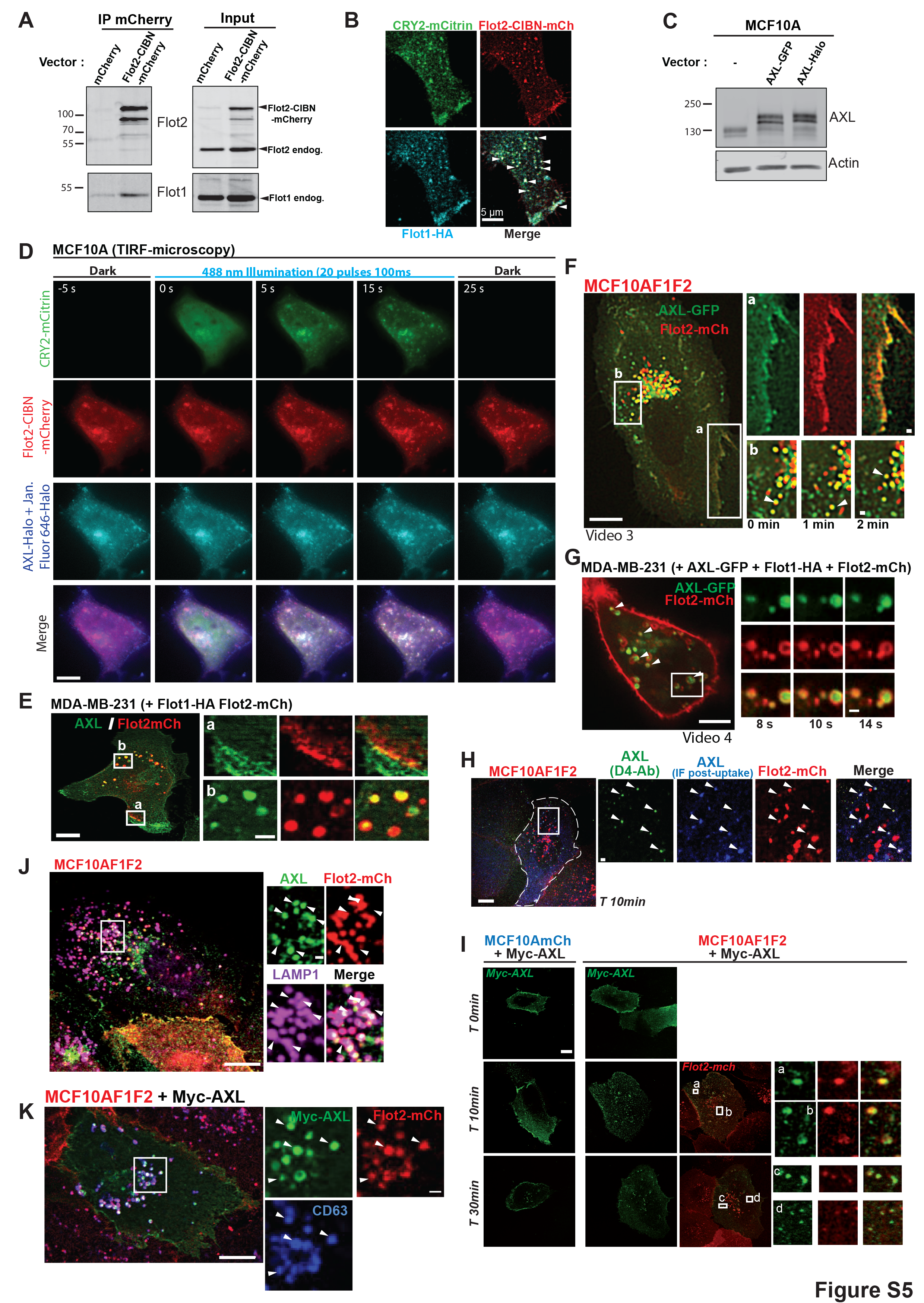

### Genest Figure S6

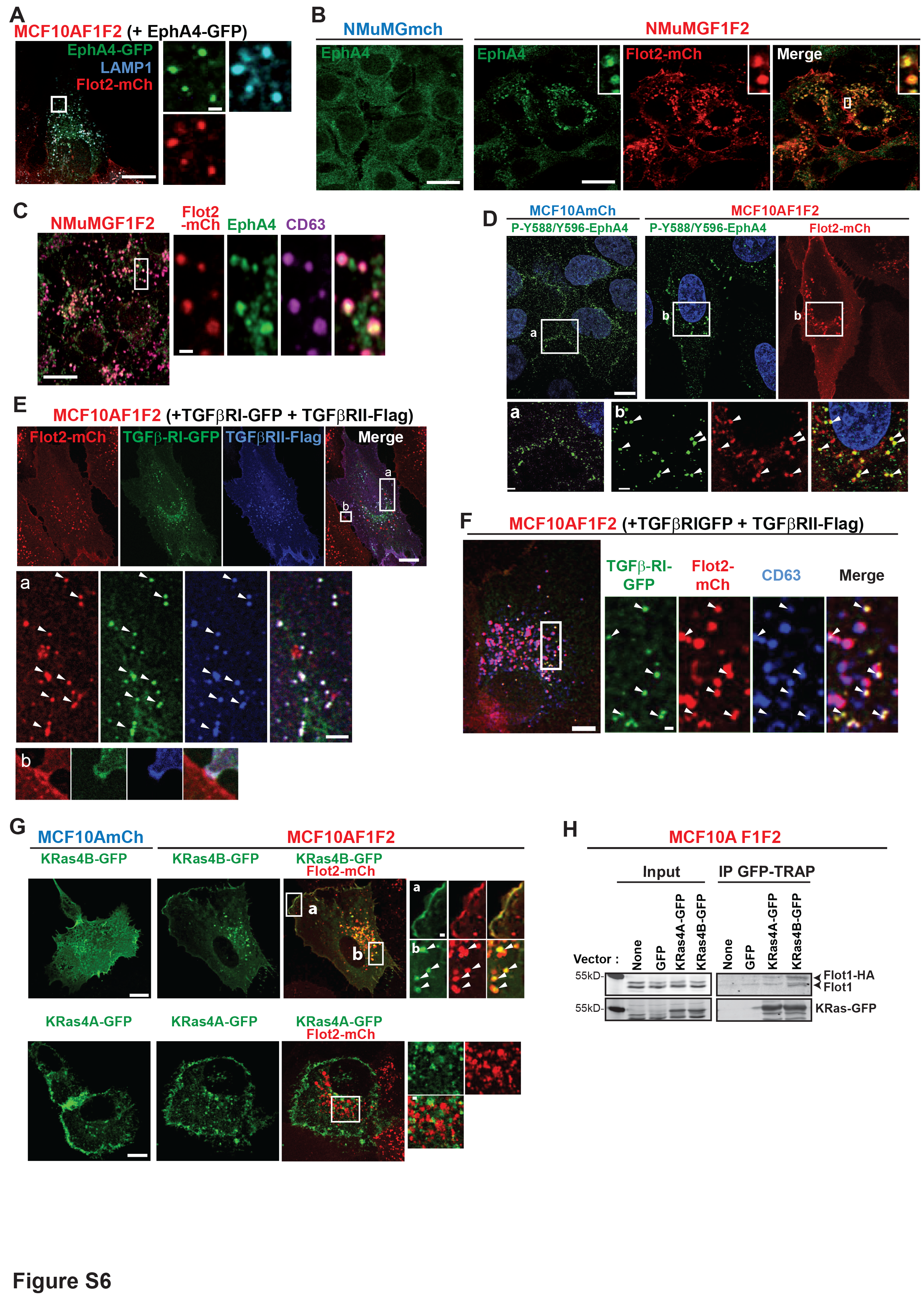

### Genest Figure S7

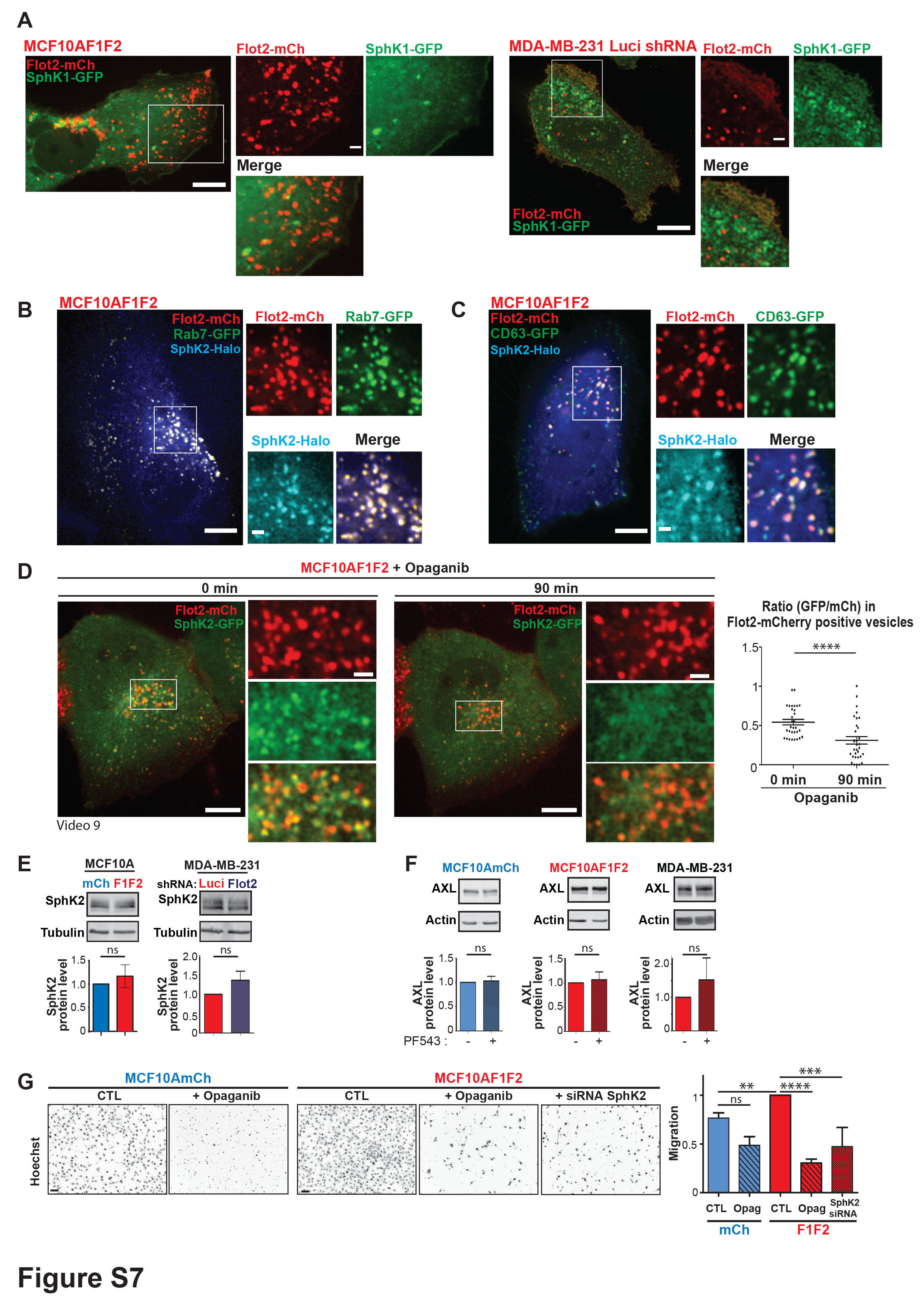
